# Comparative genomics and phylogenomics of the Mustelinae lineage (Mustelidae, Carnivora)

**DOI:** 10.1101/2025.07.01.662473

**Authors:** Azamat A. Totikov, Andrey A. Tomarovsky, Polina L. Perelman, Tatiana M. Bulyonkova, Natalia A. Serdyukova, Aliya R. Yakupova, David Mohr, Daniel W. Foerster, Jose Horacio Grau Jipoulou, Violetta R. Beklemisheva, Mikhail Sidorov, Inês Miranda, Liliana Farelo, Alexei V. Abramov, Ksenia Krasheninnikova, Anna S. Mukhacheva, Victor V. Panov, Elena Balanovska, Nikolay Cherkasov, Karol Zub, Alan F. Scott, José Melo-Ferreira, Innokentiy M. Okhlopkov, Anna Zhuk, Klaus-Peter Koepfli, Alexander S. Graphodatsky, Sergei Kliver

**Author notes:** /.

## Abstract

Mustelinae are among the most diverse and taxonomically complex subfamilies within the Mustelidae, yet their evolutionary history and genetic diversity remain largely unexplored at the whole-genome level. Here, we present the first comprehensive comparative and phylogenomic study of this lineage, integrating nuclear and mitochondrial genomes from ten species across the Holarctic and Indomalayan realms. Our dataset includes two novel genome assemblies (*Mustela strigidorsa*, *M. sibirica*) and an improved genome for *M. nivalis*, enabling robust cross-species analyses of genome size, chromosomal evolution, genetic diversity, and demographic history. We uncover striking inter-and intraspecific variation in genome-wide heterozygosity and genome size, with evidence of marked homozygosity in some Asian lineages (*M. eversmanii*, *M. sibirica*, *M. strigidorsa*) and remarkable genetic diversity in widespread species such as *M. nivalis* and *M. erminea*. Phylogenomic results support the previously suggested split of *M. richardsonii* from *M. erminea*, but we found no evidence for speciation within *M. nivalis*. Ancestral reconstruction of chromosomal rearrangements revealed key chromosomal fissions that shaped the Mustelinae radiation, including early events predating the divergence of modern *Mustela* species. The results confirmed the ancestral karyotype of *Mustela* (2n=44) and the Mustelinae (2n=42). Finally, demographic reconstructions exposed species-specific responses to Quaternary climatic cycles, ranging from long-term resilience in *M. nivalis* to repeated population bottlenecks in *M. putorius* and *M. sibirica*. Collectively, our findings establish a genomic foundation for future evolutionary and conservation genomic research on this emblematic Mustelidae lineage.

## Introduction

The Mustelinae lineage represents the most numerous and diverse group of small mammals within the Mustelidae family, comprising at least 20 recognized species in the genera *Mustela* and *Neogale* (Figure 1). Members of this subfamily range from the cosmopolitan *M. nivalis*, the smallest member of the order Carnivora, to the tropical and poorly understood *M. strigidorsa*. Their remarkable species richness and high ecological adaptability have enabled them to inhabit a wide range of environments throughout the Holarctic region (Figure 1) (Macdonald et al. 2017). While most Mustelinae species inhabit the Palearctic and Nearctic regions, some are restricted to the Neotropics (*N. africana*, *N. felipei*) or the Indomalayan zone (*M. kathiah*, *M. strigidorsa*, *M. nudipes* and *M. lutreolina*). Many of these species are exploited by humans to varying degrees, whether in the fur industry (*M. sibirica*, *M. putorius*, *M. eversmanii*, *M. erminea*, and *N. vison*), in agriculture for rodent population control (for example, *M. erminea*), in traditional Asian medicine (for example, *M. strigidorsa*), as domestic pets, or even as model organisms for studying various viral diseases (*M. putorius furo*) (Belser et al. 2011; Harrington et al. 2017; Hiller and Vantassel 2022).

**Figure 1.**
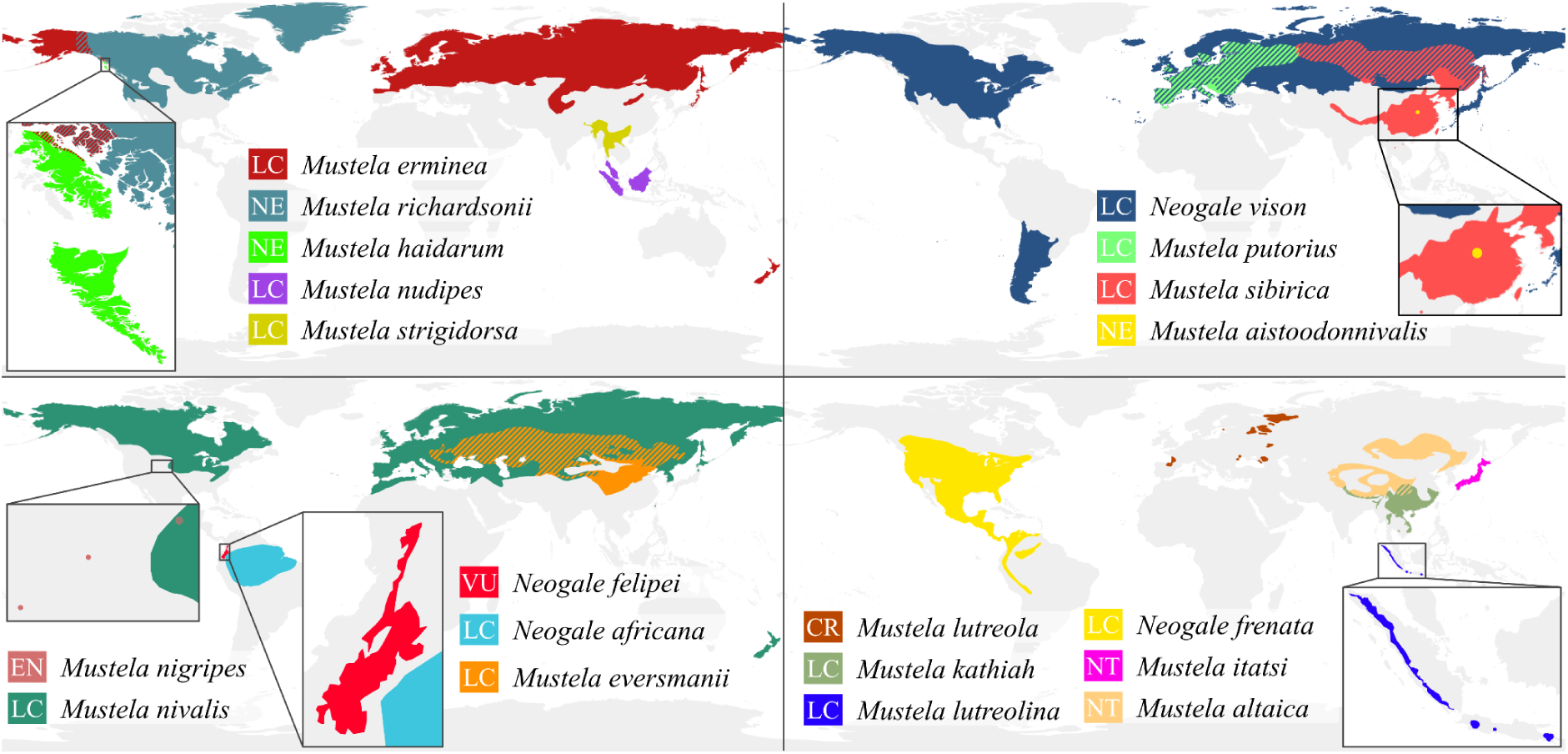
Geographic range and global conservation status of Mustelinae species. Ranges for *M. richardsonii*, *M. haidarum* and *M. aistoodonnivalis* are based on previous studies (Colella et al. 2018; Colella et al. 2021; Liu et al. 2023). Ranges for all other species were obtained from the IUCN Red List of Threatened Species (version 2025-1) (IUCN 2025). Global conservation statuses according to IUCN data: LC – Least Concern, NT – Near Threatened, VU – Vulnerable, EN – Endangered, CR – Critically Endangered, NE – Not Evaluated

Research on Mustelinae species has often focused only on describing their morphological traits, emphasizing physical characteristics such as body size, skull shape, fur patterns and other distinctive features (Kitchener et al. 2017). However, these studies inevitably encountered and brought to light the challenges of studying this subfamily, with factors such as varying degrees of sexual dimorphism, age-related variability and the occurrence of interspecific hybrids complicating accurate taxonomic delineation (Kitchener et al. 2017). As a result, researchers increasingly turned to molecular and cytogenetic approaches to gain deeper insight into the evolution of the subfamily. For example, molecular studies revealed that several species within the Mustelinae exhibit obligate embryonic diapause (for example, *M. erminea*, *N. vison,* and *N. frenata*), a reproductive strategy that ensures postnatal development occurs under the most favorable environmental conditions for offspring survival (Thom et al. 2004; Fenelon et al. 2017). Cytogenetic research elucidated the complex karyotypic evolution in Mustelinae, characterized by multiple chromosomal rearrangements, including fusions that produced the *N. vison* karyotype (2n=30) and both fissions and fusions that generated a karyotype resembling that of *M. erminea* (2n=44) (Graphodatsky et al. 1989). Moreover, it has become evident that substantial differences in the number of chromosomal arms among *Mustela* species, despite similar or identical diploid numbers (2n), cannot be explained solely by unequal translocations or pericentric inversions, which typically alter chromosome morphology (Graphodatsky et al. 1989). Karyotype evolution within *Mustela* has involved the emergence of additional, entirely heterochromatic arms in the karyotypes of certain species, through an increase in heterochromatic material on the short arms of originally acrocentric chromosomes (Graphodatsky et al. 2002). While Mustelinae species share some similarities in heterochromatin localization, they differ in the size of heterochromatic blocks and, consequently, in total heterochromatin content, which affects the genome size (Graphodatsky et al. 1977). Despite data limitations, these and other cytogenetic studies have laid the groundwork for understanding the complex phylogenetic relationships within this diverse group of mammals. Notably, even before the advent of sequencing technologies, cytogenetic studies had already indicated the need to separate *Neogale* from *Mustela* within the Mustelinae subfamily, based on karyotypic evolution from a *Martes*-like ancestor to a *Mustela*-like lineage (Graphodatsky et al. 1989).

In recent decades, the systematics of Mustelidae species has undergone significant revisions and changes, primarily based on nuclear and/or mitochondrial data (Koepfli, Deere, et al. 2008; Yu et al. 2011; Sato et al. 2012; Law et al. 2018; Hassanin et al. 2021). Stepping beyond morphological markers has enabled a reevaluation of the traditional classification of Mustelidae [19], leading to the division of the five previously recognized subfamilies into eight (Koepfli, Deere, et al. 2008; Yu et al. 2011). Subsequent studies focusing on Mustelinae, along with extensive taxonomic discussions (Harding and Smith 2009; Hassanin et al. 2021; Patterson et al. 2021), helped to refine the boundaries between the genera *Mustela* and *Neogale*. Species-level classification remains challenging due to the rapid evolutionary diversification within this group (Koepfli, Kanchanasaka, et al. 2008; Law et al. 2018; Hassanin et al. 2021), which has resulted in a number of morphologically indistinguishable species, as well as hard-to-resolve nodes on phylogenetic trees (Koepfli, Deere, et al. 2008). This complexity is further exacerbated by hybridization among some species (Cabria et al. 2011; Cserkész et al. 2021), which often leads to ambiguous phylogenetic relationships (Etherington et al. 2022). These challenges make it difficult to determine the total number of species within Mustelinae, or their phylogenetic relationships, with no current scientific consensus on either the number of species or subspecies. Despite these difficulties, a notable taxonomic revision has been the recognition of *M. itatsi* as taxonomically distinct from the closely related *M. sibirica*, previously classified as a single taxonomic unit (Abramov 2000). Similarly, taxonomic reassessment has successfully resolved the classification of *M. erminea.* Whole-genome and mitochondrial data (Colella et al. 2018; Colella et al. 2021) revealed that North American *M. erminea* actually comprises three distinct species: *M. erminea*, *M. richardsonii* and *M. haidarum*.

Currently, mitochondrial genome assemblies are available for nearly every species within the Mustelinae, with the exception of *M. lutreolina*, *M. aistoodonnivalis*, *N. felipei* and *N. africana*. However, the availability of whole-genome data is far less complete. Genome assemblies are absent for thirteen species (*M. strigidorsa*, *M. sibirica*, *M. altaica*, *M. aistoodonnivalis*, *M. itatsi*, *M. kathiah*, *M. lutreolina*, *M. nudipes*, *M. haidarum*, *M. richardsonii*, *N. frenata*, *N. felipei* and *N. africana*). Among the remaining species, chromosome-level genome assemblies were published for only six species (*M. nivalis*, *M. erminea*, *M. lutreola*, *M. nigripes*, *M. putorius furo* and *N. vison*), and scaffold-level genome assemblies exist for only two others (*M. eversmanii* and *M. putorius*). Notably, the genome assembly for *M. nivalis* was published very recently (mMusNiv2.hap1.1, GCA_964662115.1). Despite the increasing volume of available genomic data, its distribution across the Mustelinae lineage remains uneven (Supplementary Table ST1), underscoring the need for further sequencing and genome assembly efforts for the less-studied species.

Comparative analysis based on the integration of nuclear and mitochondrial genome data represents a powerful tool for studying evolutionary processes, enabling the identification of both common features and key differences between related species. This approach is particularly significant in the context of conservation research, where understanding genetic diversity, phylogenetic relationships and demographic history is crucial for developing effective biodiversity conservation strategies. In this study, we compiled an extensive dataset, comprising 10 genome assemblies, 50 resequencing samples and 149 mitochondrial genomes. This dataset includes three new genomic assemblies, two of which were generated by us and one improved from a publicly available draft genome assembly, as well as 9 samples newly resequenced as part of this project. Using this comprehensive dataset, we aimed to: (1) assess and compare whole-genome levels of genetic diversity among the species; (2) determine interspecific and intraspecific phylogenetic relationships; (3) reconstruct population history; (4) evaluate genome size variation within and between species; and (5) reconstruct synteny blocks to identify structural rearrangements among genomes. Thus, our study contributes to a deeper understanding of genetic diversity and evolutionary processes within the Mustelinae, while also offering valuable guidance for the development of conservation strategies and the preservation of genetic diversity in these species.

## Results

### Chromosome-level genome assemblies and connection with karyotypes

We improved the publicly available genome assembly of *M. nivalis* (GCA_019141155.1) to chromosome-level and generated the first chromosome-level assembly for *M. strigidorsa*. The *M. nivalis* assembly includes 21 chromosomal scaffolds (chromosomes) (Supplementary Figure SF1A), which correspond to the published karyotype of *M. nivalis* (2n=42) (Graphodatsky et al. 2020), and chromosome numbering follows this karyotype. For *M. strigidorsa* we observed an additional pair of chromosomes (Supplementary Figure SF1B), with 2n=44, similar to *M. erminea* (Graphodatsky et al. 2020). The difference in karyotype number between *M. nivalis* and *M. strigidorsa* + *M. erminea* is due to the fusion of chr14 and chr16 in the former species relative to the latter two species (Supplementary Figure SF1C). No discrepancies between pairwise whole-genome alignments (WGAs) and available cytogenetic data (Supplementary Tables ST2, <u>ST3</u>) (Graphodatsky et al. 2002; Graphodatsky et al. 2020) were detected for the *M. nivalis* assembly. A one-to-one synteny between *M. strigidorsa* and *M. erminea* chromosome scaffolds was observed (Supplementary Figure SF1C); thus we decided to use the same chromosome numbering for both species.

The genome assemblies of *M. strigidorsa* and *M. nivalis* have total lengths of 2.42 Gbp and 2.45 Gbp, respectively (Supplementary Table ST4). Notably, the assembly lengths differ from the estimates based on 23-mers: 3.21 Gbp for *M. nivalis* and 3.34 Gbp for *M. strigidorsa* (Supplementary Table ST5). The sizes of the assemblies are comparable to the lengths of the assemblies of the other species included in our study, ranging from 2.41 Gbp for the *M. sibirica* to 2.68 Gbp for the *N. vison* (Supplementary Table ST4). The scaffold N50s, 115.1 Mbp (*M. strigidorsa*) and 138.4 Mbp (*M. nivalis*) are also comparable to the values for other assemblies (130.15 - 220.3 Mbp, depending on the species) (Supplementary Table ST4). The BUSCO analysis using the Mammalia odb10 dataset (2024-01-08, total BUSCOs = 9226) on the obtained genome assemblies of *M. strigidorsa* and *M. nivalis* showed 94.6% and 96% complete and single-copy BUSCOs, with only 3.9% and 2.9% missing BUSCOs, respectively. Chromosome-level genome assemblies of other Mustelinae species also demonstrated high quality, with complete and single-copy BUSCOs ranging from 94.4% to 96.4%, and missing BUSCOs ranging from 2.6% to 3.8% (Supplementary Figure SF2 and Supplementary Table ST6).

### Genome sizes and repeats content

We observed significant variation of genome sizes across the analyzed species and samples (Figure 2; Supplementary Table ST5). The estimates ranged from 2.16 Gbp for *M. putorius* (ERR7260426) to 3.35 Gbp for *M. nivalis* (T100). The greatest variation was detected within *M. putorius* and *M. richardsonii*. The *M. putorius* samples formed two distinct groups: one ranging from 2.16 Gbp to 2.3 Gbp and another from 2.53 Gbp to 2.63 Gbp. Among *M. richardsonii* samples, one outlier exhibited a notably small genome size of 2.29 Gbp, while the remaining samples ranged from 2.58 Gbp to 2.79 Gbp. The smallest differences in genome size were observed between *N. vison* individuals.

**Figure 2.**
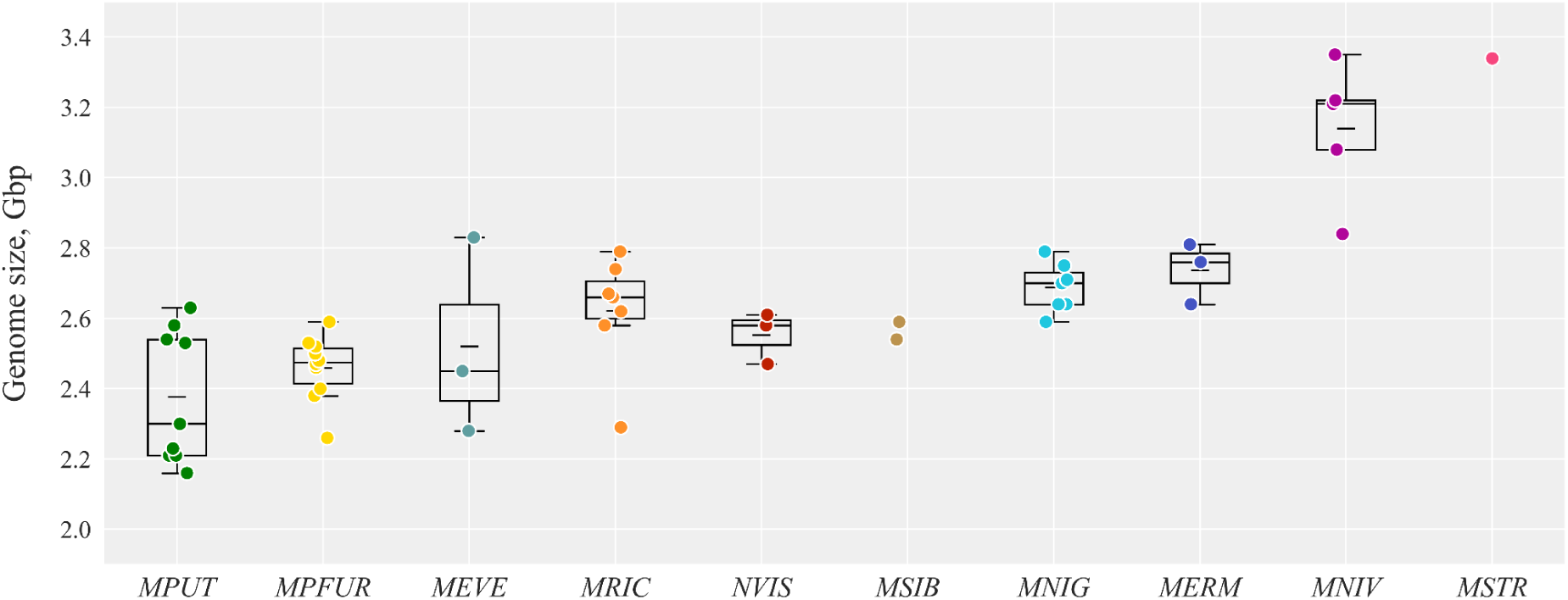
Estimation of genome sizes for Mustelinae species based on k-mer analysis. Short horizontal lines within boxplots represent the mean value and long lines indicate the median value. Species abbreviation: MPUT – *Mustela putorius*, MPFUR – *Mustela putorius furo*, MEVE – *Mustela eversmanii*, MRIC – *Mustela richardsonii*, NVIS – *Neogale vison*, MSIB – *Mustela sibirica*, MNIG – *Mustela nigripes*, MERM – *Mustela erminea*, MNIV – *Mustela nivalis*, MSTR – *Mustela strigidorsa*. Detailed data are provided in Supplementary Table ST5

An analysis of transposable elements (TE) in the genome assemblies revealed TE content ranging from 35.85% to 39.6% (Supplementary Table ST7). Most TEs belonged to the SINE and LINE superfamilies. Minimal SINE and LINE content were observed in the *M. putorius furo* genome with 9.61% and 19.02%, respectively. Kimura distance-based copy divergence profiles showed similar patterns (same number and location of the peaks) for assemblies across the seven species which included three or more individuals (Supplementary Figure SF3).

### Genome rearrangements

We reconstructed a macro-level synteny map for the available genomes of the Mustelinae as well as other species within the Mustelidae family (Figure 3). This revealed highly conserved syntenic segments, mostly maintaining centromere positions across species. However, some exceptions indicate evolutionary shifts in centromere positions within the genus *Mustela* (Supplementary Figure SF4). For example, centromere positions in homologous chromosomes of *M. strigidorsa* and *M. erminea* differ from those in other *Mustela* species. A centromeric shift transformed subtelocentric MSTR 5 and MERM 5 to acrocentric MNIV 17 and MLUT 13, or even to acrocentric MPFUR 16 and MNIG 13 (chromosome numbers are used throughout). Notably, these changes in centromere position occurred without apparent chromosomal rearrangements. A similar pattern was observed for MERM 12 and its homologs: MSTR 12 and MERM 12 are acrocentrics, whereas MNIV 13, MLUT 9, MPFUR 10 and MNIG 9 are submetacentrics.

**Figure 3.**
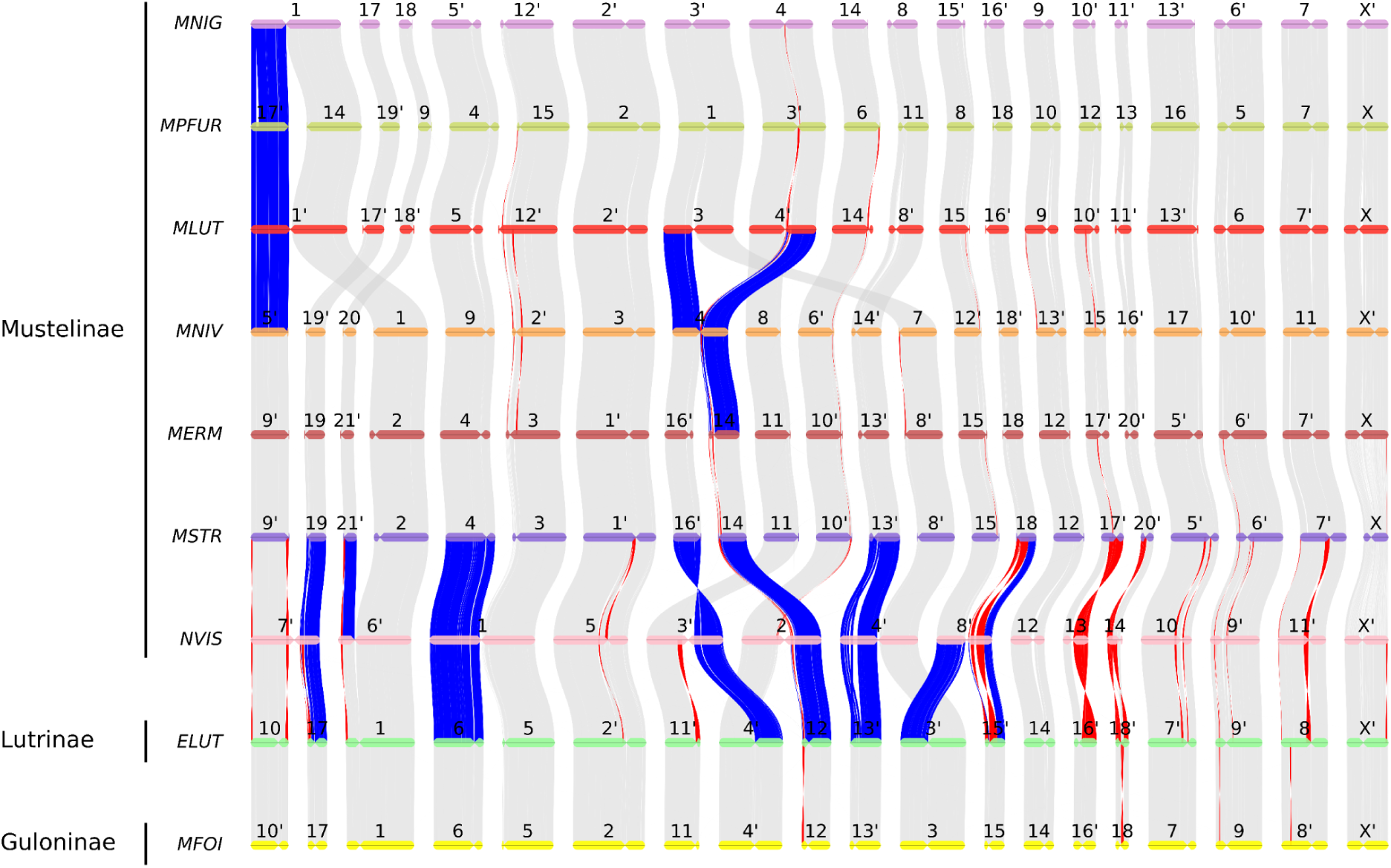
Macro-level synteny map among species within the Mustelinae and Mustelidae. Chromosomes labeled by primes (‘) were reverse complemented. Fusions and fissions are highlighted in blue. Inversions larger than 1 Mbp are highlighted in red. Abbreviation: MFOI – *Martes foina*, ELUT – *Enhydra lutris*, NVIS – *Neogale vison*, MSTR – *Mustela strigidorsa*, MERM – *Mustela erminea*, MNIV – *Mustela nivalis*, MLUT – *Mustela lutreola*, MPFUR – *Mustela putorius furo*, MNIG – *Mustela nigripes*. Macro-level synteny (gray lines) is shown between chromosomes (horizontal colored lines) of 7 species of Mustelinae, one Lutrinae (ELUT) and one Guloninae (MFOI) species. Centromeric positions are tentative, approximately drawn based on the analysis of comparative chromosome painting maps and G-banded karyotypes

Despite the highly conserved synteny among Mustelinae species, we identified a number of structural rearrangements in the genomes of *Mustela* species, including new inversions and all previously described translocations. Some of the inversions are likely lineage-specific. For example, inversions between MNIV 12, 13 and 15, and their homologs MLUT 15, 9 and 10 likely distinguish the ferret-like (MLUT, MPFUR, MNIG) lineage from other *Mustela* species (MSTR, MERM, MNIV). The sizes of these inversions are approximately 6.89 Mbp, 2.23 Mbp, and 13.19 Mbp, respectively. At the same time, NVIS stands out significantly due to the presence of a large number of translocations and species-specific inversions (Figure 3) even if multiple fissions and fusions of chromosomes are not considered. Expectedly, distinct heterochromatic arms observed in cytogenetic studies (Graphodatsky et al. 1976; Graphodatsky et al. 1977; Graphodatsky et al. 1989; Graphodatsky et al. 2002) on the chromosomes for some *Mustela* taxa are absent in current genome assemblies. This discrepancy is evident from the reconstructed syntenic blocks. For instance, the large heterochromatic arm on MNIV 1 and MERM 9 is missing.

### Heterozygosity level and Runs of Homozygosity

We detected significant variation in mean heterozygosity at both the interspecific and intraspecific levels (Figure 4; Supplementary <u>File</u>; Supplementary Table ST8). Among all the species we analyzed, the highest intraspecific variation was observed in those with samples collected from multiple localities across their geographic range: *M. putorius* (0.26–0.8 SNP/kbp), *M. eversmanii* (0.54–0.62 SNP/kbp), *M. sibirica* (0.85 and 1.11 SNP/kbp), *M. richardsonii* (1.28–2.94 SNP/kbp), *M. erminea* (1.87, 2.1 and 2.14 SNP/kbp) and *M. nivalis* (2.23–2.9 SNP/kbp). Species showing low intraspecific variation in mean heterozygosity include *M. nigripes* (0.02–0.04 SNP/kbp), which is expected due to a relatively recent critical bottleneck (Wisely et al. 2002); *M. putorius furo* (0.13–0.25 SNP/kbp), a domesticated species; and *N. vison* (0.85, 0.86 and 0.86 SNP/kbp), for which samples were collected from an experimental fur farm (Manakhov et al. 2020; Bergeron et al. 2023). *M. strigidorsa* is represented by a single sample with mean heterozygosity of 0.62 SNP/kbp.

**Figure 4.**
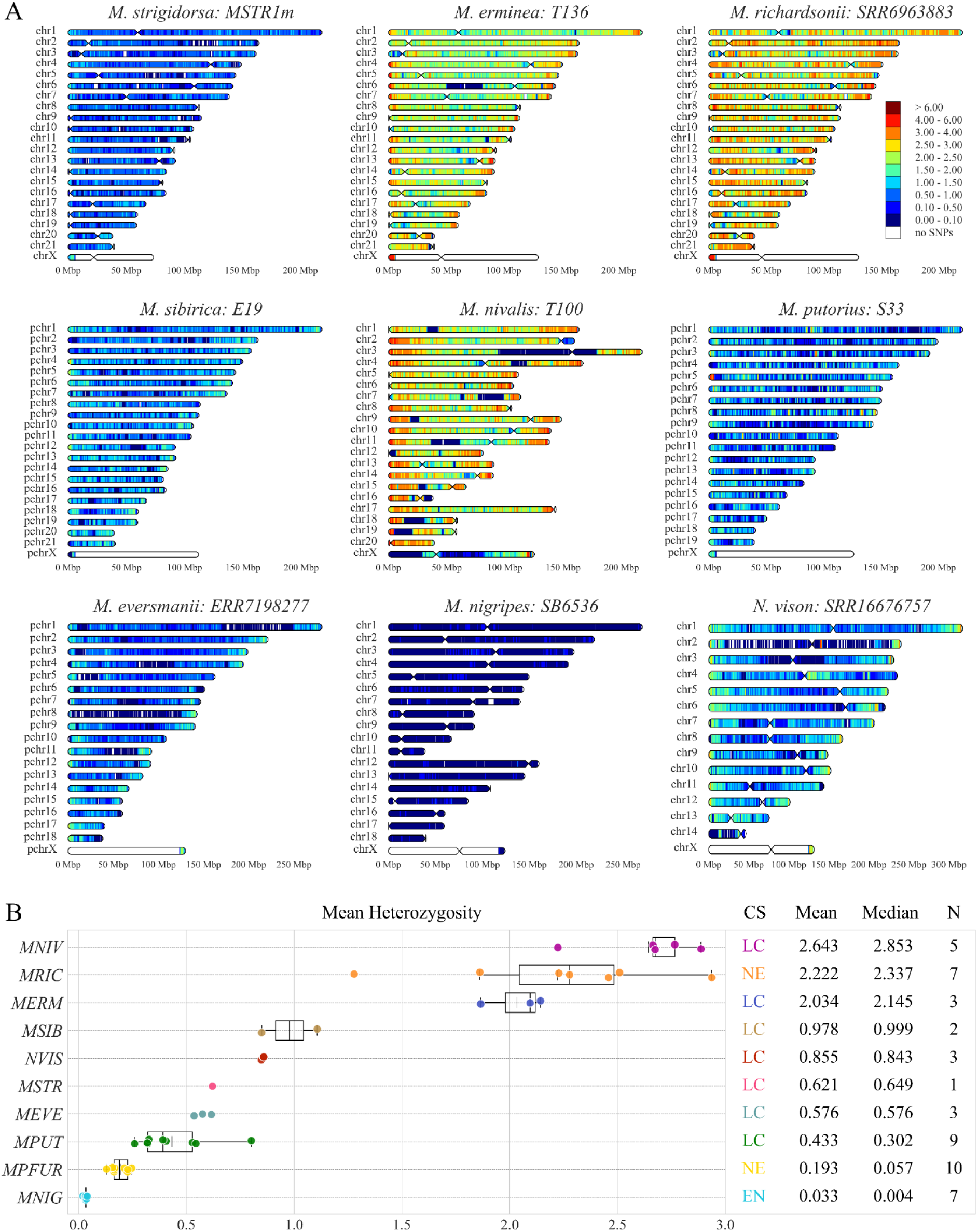
Heterozygosity of Mustelinae species. (A) Distribution of the heterozygous SNP density on the chromosomes of Mustelinae species. Heterozygous SNPs were counted in 1 Mbp windows with 100 kbp steps and scaled to SNPs/kbp (represented by color scale on the left, ranging from dark blue (extremely low heterozygosity) to brown (very high heterozygosity), with 0 to 0.1 and over 6 heterozygous SNPs per 1 kbp respectively). Centromeres are not indicated for pseudochromosome-level assemblies, where chromosomes are labeled with the prefix “pchr”. Each heatmap represents a single individual selected as a representative of the corresponding species. Distribution of heterozygous SNP density on chromosomes for the other samples is presented in the Supplementary <u>File</u>. (B) Intra-and interspecific heterozygosity among Mustelinae species. The values at the top of the figure reflect the intraspecies mean and median heterozygosity. Short vertical lines within boxplots represent the mean value and long lines indicate the median value. Each dot corresponds to a separate individual. Species abbreviation: MNIG – *Mustela nigripes*, MPFUR – *Mustela putorius furo*, MPUT – *Mustela putorius*, MEVE – *Mustela eversmanii*, MSTR – *Mustela strigidorsa*, NVIS – *Neogale vison*, MSIB – *Mustela sibirica*, MERM – *Mustela erminea*, MRIC – *Mustela richardsonii*, MNIV – *Mustela nivalis*. Species global conservation status (CS) according to IUCN data: EN – Endangered, NE – Not Evaluated, LC – Least Concern. N – number of individuals. Detailed data are provided in Supplementary Table ST8

The two species with the lowest heterozygosity (*M. nigripes* and *M. putorius furo*) display many short (<1 Mbp), long (>=1 Mbp) and ultra long (>=10 Mbp) RoH (<u>R</u>uns of <u>H</u>omozygosity) on all chromosomes (Figure 5; Supplementary Figure SF5), which encompass a significant fraction of their genomes (Supplementary Table ST9). In the *M. nigripes* individuals, such regions cover 2.1-2.2 Gbp out of 2.5 Gbp of the assembly (199-414 RoH) and 77.83-91.81% of the total RoH length is contained in the ultra long category. In *M. putorius furo*, RoH are shorter and cover a smaller (1.48-1.88 Gbp, 64.9-82.05%) fraction of the assembly – ultra long RoH harbor only 18.09-34.44% of the cumulative RoH length.

**Figure 5.**
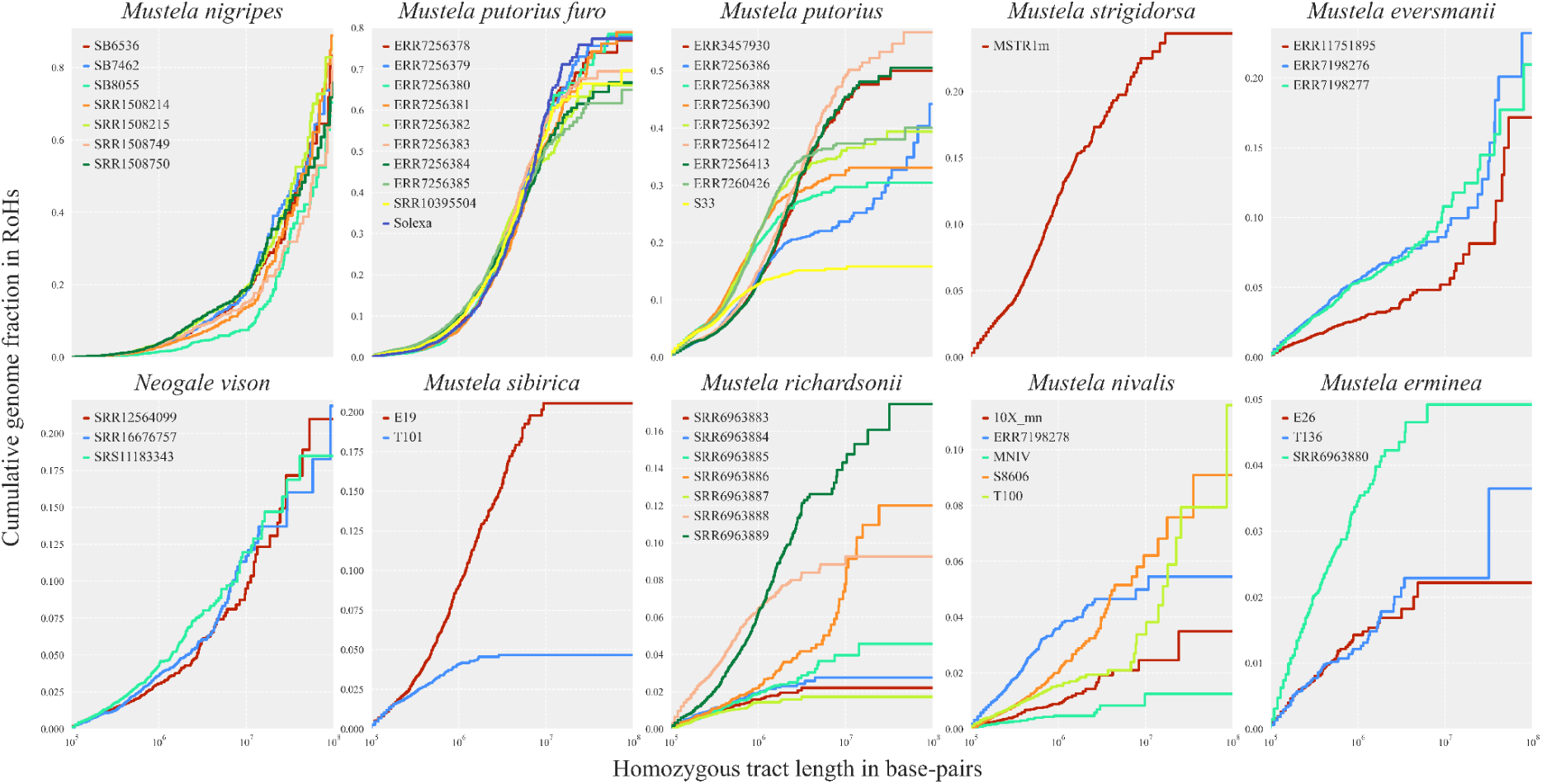
The Runs of Homozygosity (RoH) content in the Mustelinae samples by species. Cumulative distribution plots of RoH. Homozygous tract lengths and cumulative genome fraction in RoH are represented on X and Y axes. Tracts are ordered from shortest to longest. X chromosomes were excluded from all samples. Detailed data are provided in Supplementary Table ST9

*M. putorius* has the highest variation in total RoH length (Figure 5): from 371.8 Mbp (S33) to 1.4 Gbp (ERR7256386). In the samples gathered from European mainland populations (S33, ERR7256388, ERR7256390, ERR7256392 and ERR7260426), RoH encompass from 15.84% to 39.96% of the assembly (2.5 Gbp) and most are short (<1 Mbp; Supplementary Table ST9). Sample ERR7256386 (Spain) stands out from the other samples by its RoH content (1.4 Gbp in total), with a major contribution from ultra long RoH (60.1 %). The remaining *M. putorius* samples ERR3457930, ERR7256412 and ERR7256413 (UK) have a comparable fraction of RoH (49.87-52.75% of the assembly, or 1.2 Gbp). The *M. eversmanii* samples have 335-631 RoH (23.47-27.84% of the assembly), but most of their lengths (57.35-77.9%) are contained in 9-13 ultra long RoH.

The samples from the highly heterozygous species (Figure 4, Supplementary Table ST8) such as *M. richardsonii*, *M. erminea* and *M. nivalis,* are almost devoid of RoH, with a few exceptions (Supplementary Table ST9). For example, in the *M. nivalis* samples T100 and S8606, RoH encompass 270 Mbp and 131 Mbp of the assembly (2.5 Gbp) and more than half of the chromosomes of the female T100 sample contain at least one long RoH. Of these, seven RoH even exceed 10 Mbp: chr1 (10.2 Mbp), chr4 (14.1 Mbp), chr19 (16 Mbp), chr18 (18 Mbp), chr7 (22.5 Mbp), chr11 (25.5 Mbp), and chr3 (85 Mbp). Despite the overall high mean heterozygosity, we found such ultra long RoH in the *M. richardsonii* samples SRR6963885 (1 RoH, 14.2 Mbp), SRR6963886 (7 RoH, 97.4 Mbp) and SRR6963889 (4 RoH, 72.5 Mbp).

### Phylogenetic trees

#### Nuclear tree, speciation and ancestral rearrangements

We reconstructed a ML (<u>M</u>aximum <u>L</u>ikelihood) phylogenetic tree (Figure 6) using positions that vary between species (including heterozygous ones) from 6,599 single-copy protein-coding genes. The inferred topology showed a clear differentiation of the currently recognized species, with a bootstrap support value of 100 for all nodes at and above species level (Figure 6). Just as expected, *N. vison* appeared as a sister group to the genus *Mustela*. *M. strigidorsa*, the only tropical species involved in the analysis, diverged first within *Mustela*. Next, stoats (*M. richardsonii* and *M. erminea*), *M. nivalis, M. sibirica, M. lutreola* and, finally, the ferret lineage (*M. eversmannii*, *M. nigripes*, *M. putorius* + *M. p. furo*) split sequentially from the stem.

**Figure 6.**
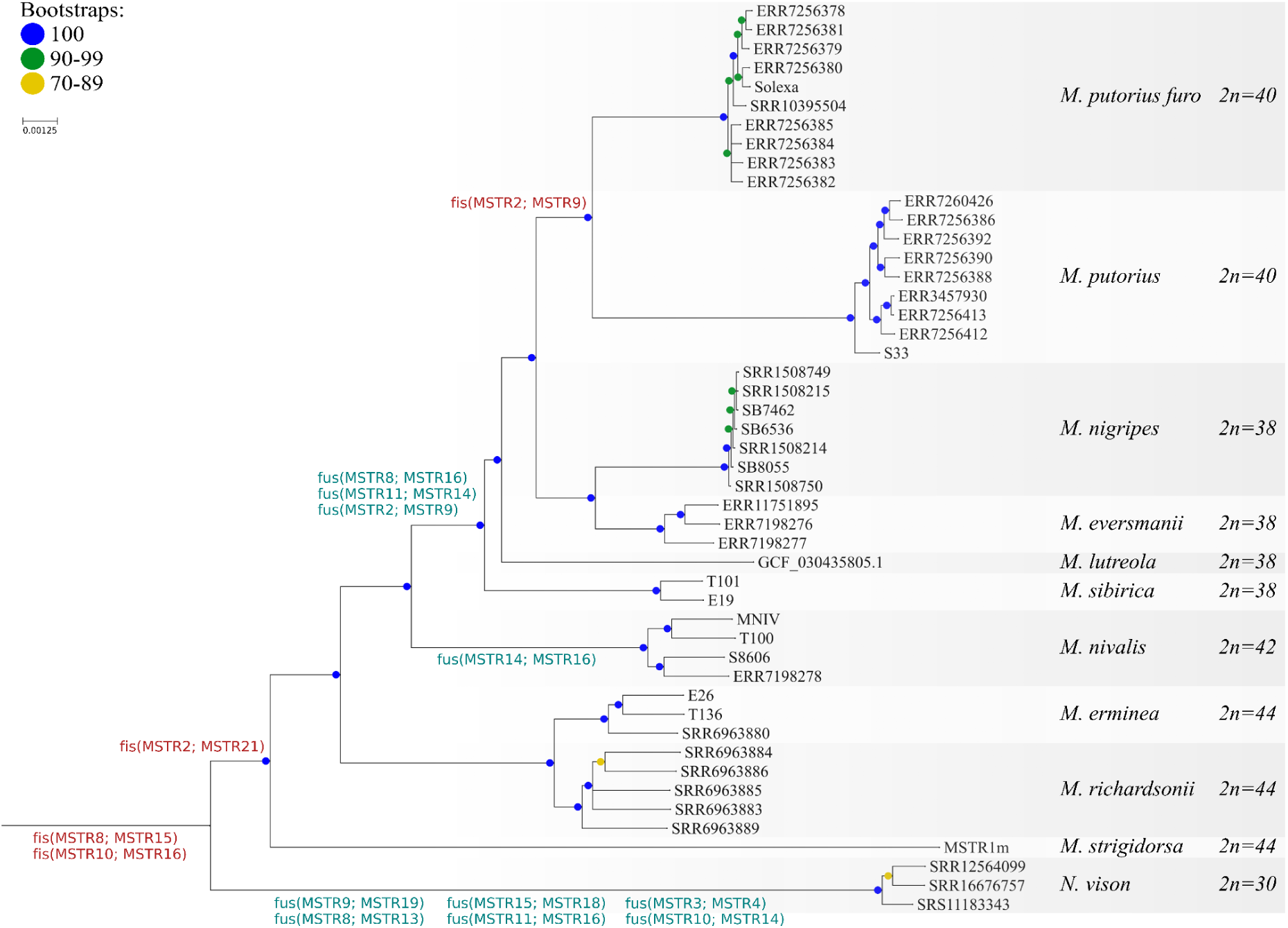
Maximum likelihood phylogram based on nuclear phylogenomic data of analyzed Mustelinae species. Only autosome data was used for the reconstruction, and the tree was rooted on *Martes foina* (not shown, mfoi.min_150.pseudohap2.1_HiC, DNAZoo). Labels near branches correspond to ancestral fissions (red) and fusions (green). The translocations were encoded relative to *M. strigidorsa* chromosomes, as they represent the chromosome-level syntenic blocks. Note a difference in designation between fissions and fusions. For example, fis(MSTR8, MSTR15) means a fission of ancestral chromosome to homologs of MSTR8 and MSTR15, and fus(MSTR14; MSTR16) means a fusion of homologs of MSTR14 and MSTR16 to a new chromosome

To test the hypothesis that there is species-level divergence within *M. nivalis,* we compared distances between and within sister species (Supplementary Figure SF6): *M. nivalis* (9.77×10^-4^ – 1.16×10^-3^), *M. erminea* (1.08×10^-3^ – 1.19×10^-3^), *M. richardsonii* (1.47×10^-3^ – 1.66×10^-3^), *M. eversmanii* (5.56×10^-4^ – 9.12×10^-4^), *M. erminea*-*M. richardsonii* (2.30×10^-3^ – 2.66×10^-3^) and *M. eversmanii*-*M. nigripes* (2.60×10^-3^ – 2.80×10^-3^). We found a clear differentiation between inter-and intraspecific distances (Figure 7). Distances within *M. nivalis* are relatively small and clearly fall within the intraspecific category: its mean (1.05×10^-3^) does not significantly differ from the *M. erminea* mean (1.15×10^-3^, two-sided Mann-Whitney test p-value 0.095) and is smaller than the mean for *M. richardsonii* (1.55×10^-3^, one-sided Mann-Whitney test p-value 0.00068). At the same time, our comparison confirms the previous split of *M. richardsonii* and *M. erminea* as the mean distance between them is similar to the distance between *M. eversmani* and *M. nigripes* (2.47×10^-3^ vs 2.72×10^-3^).

**Figure 7.**
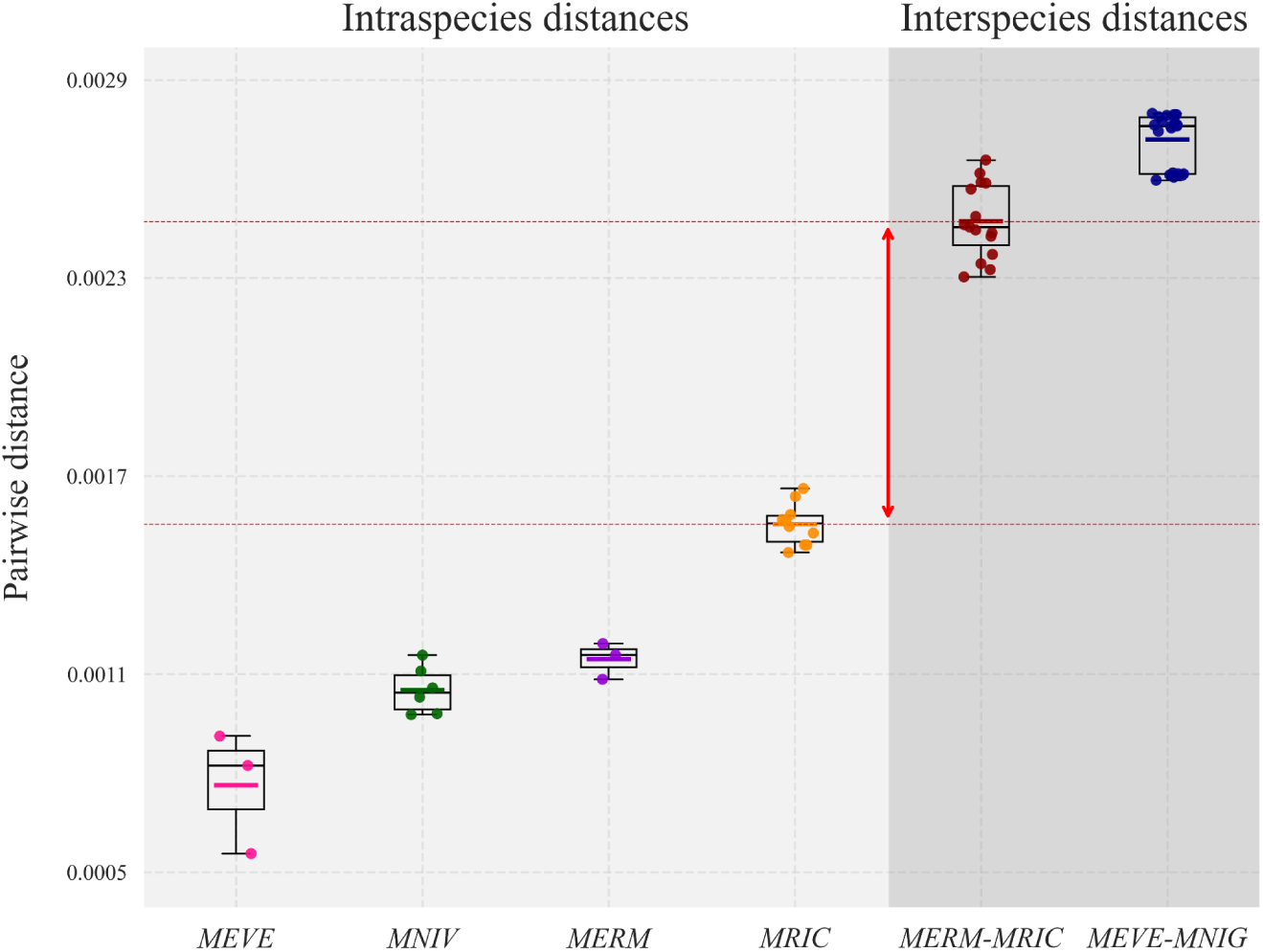
Intraspecific and interspecific pairwise distances among Mustelinae species. Colored and black horizontal lines within each boxplot represent the mean and median values, respectively. Red dashed lines and arrows indicate the gap between intra-and interspecific distances. Species abbreviation: MEVE – *Mustela eversmanii*, MNIV – *Mustela nivalis*, MERM – *Mustela erminea*, MRIC – *Mustela richardsonii*, MNIG – *Mustela nigripes*

We also reconstructed a species tree using the coalescent-based approach (Supplementary Figure SF7). The inferred topology was completely congruent with the concatenated ML tree (Figure 6). All internal nodes at and above the species level showed high posterior probabilities (pp1=1.0). However, quartet support values (q1) varied across the tree (Supplementary Figure SF8, Supplementary Table ST10). The lowest support was observed for node 9 (MRCA of *M. sibirica* and *M. lutreola)*, with only 47.7% of quartets supporting the main topology (q2=30.2%, q3=21.9%). A higher, but still reduced, quartet support was detected for node 2 (MRCA of *Neogale* and *Mustela*, q1=61.7%), which corresponds to the MRCA (Most Recent Common Ancestor) of the *Mustela* and *Neogale* (*N. vison*). All other interspecific nodes were supported by a high fraction of the quartets (q1=72.8–88.7%).

We mapped the detected Robertsonian translocations onto the phylogenetic tree (Figure 6) and performed an ancestral reconstruction of chromosomal fission and fusion events (Supplementary Figure SF9). Our results are consistent with previous cytogenetic study (Graphodatsky et al. 2002). However, inclusion of the *M. strigidorsa* assembly, generated in this study, resulted in a better relative placement of the fission event fis(MSTR2; MSTR21). Specifically, while the earlier study suggested that fis(MSTR2; MSTR21) occurred after the divergence of *Neogale* but before the split of *M. erminea*, our phylogenetic placement of *M. strigidorsa*, as a basal lineage within *Mustela*, indicates that the event likely occurred even earlier, prior to the diversification of extant *Mustela* species.

#### Mitochondrial tree

We reconstructed a ML mitochondrial phylogenetic tree using 149 mitochondrial genomes including all the samples from the nuclear tree except for the *M. p. furo, N. vison* and *M. nigripes* (Supplementary Table ST11). We obtained a well-resolved topology with 100% bootstrap support for all nodes above species level (Figure 8). At the intraspecific level we observed a complex genetic structure composed of multiple clades, likely reflecting both distinct populations and subspecies. For example, *M. sibirica* samples formed two well-resolved major clades, each of which further subdivided into two subclades. Positions of most species were the same on the mitochondrial (Figure 8), nuclear (Figure 6) and species trees (Supplementary Figure SF7), with a few exceptions. For two samples (colored in Figure 8) we observed mitonuclear discordance: the mitochondrial genome (1906) of *M. eversmanii* individual ERR11751895 (male) clustered together with *M. putorius* in the mitochondrial tree. Similarly, sample ERR7260426, identified as *M. putorius* (male), appears to have *M. lutreola* mtDNA (BK069848).

**Figure 8.**
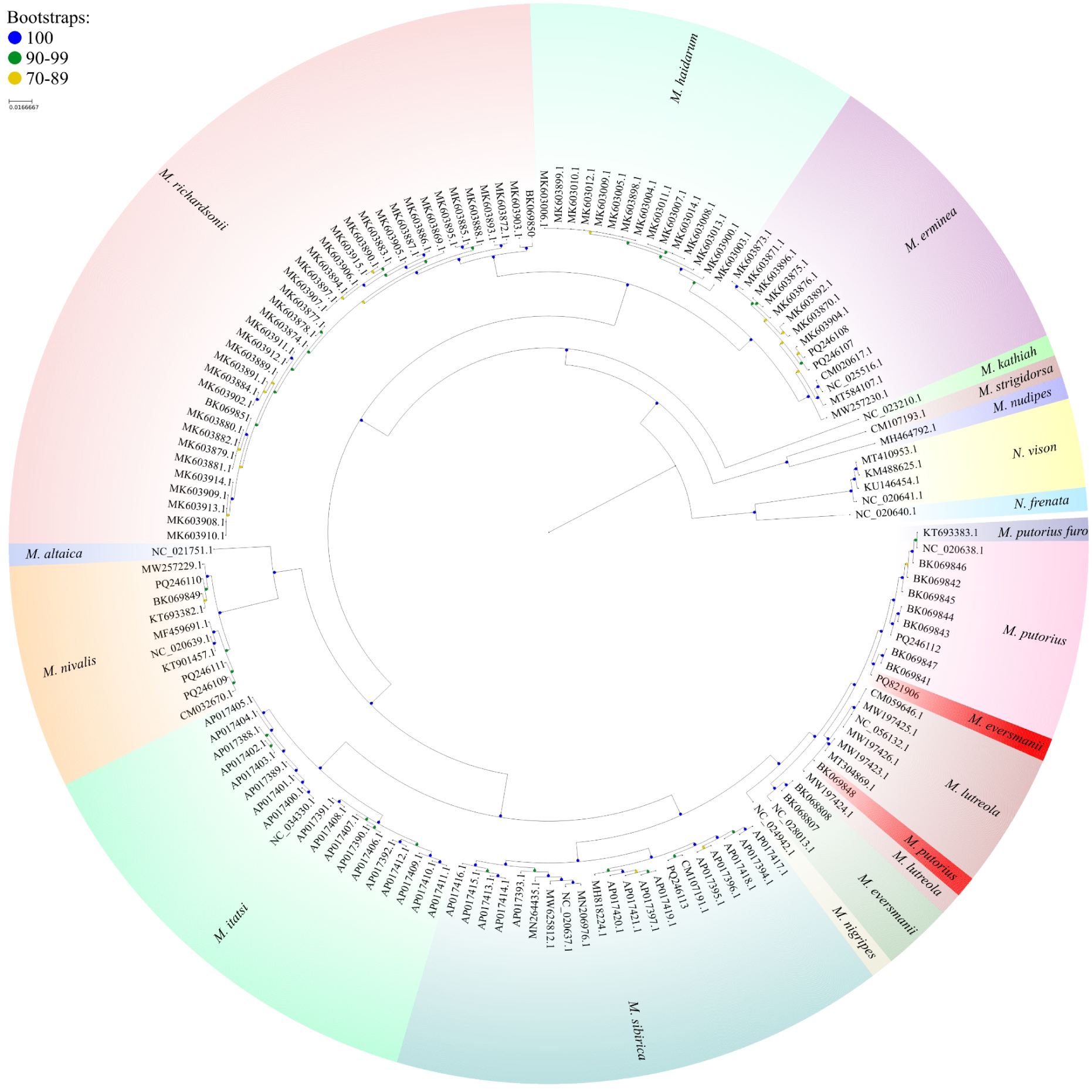
Phylogenetic tree constructed using the maximum likelihood method based on complete mitochondrial genomes (15,793 bp). The tree was rooted on *Martes foina* (not shown, NC_020643.1). Samples with mitonuclear discordance are highlighted in red

#### Demographic history

We analyzed the historical effective population size (N_e_) dynamics based on whole-genome data from various *Mustela* species using the Pairwise Sequentially Markovian Coalescent model (PSMC) (Li and Durbin 2011). The results revealed pronounced differences in N_e_ trajectories, reflecting the distinct demographic histories of each species (Figure 9).

**Figure 9.**
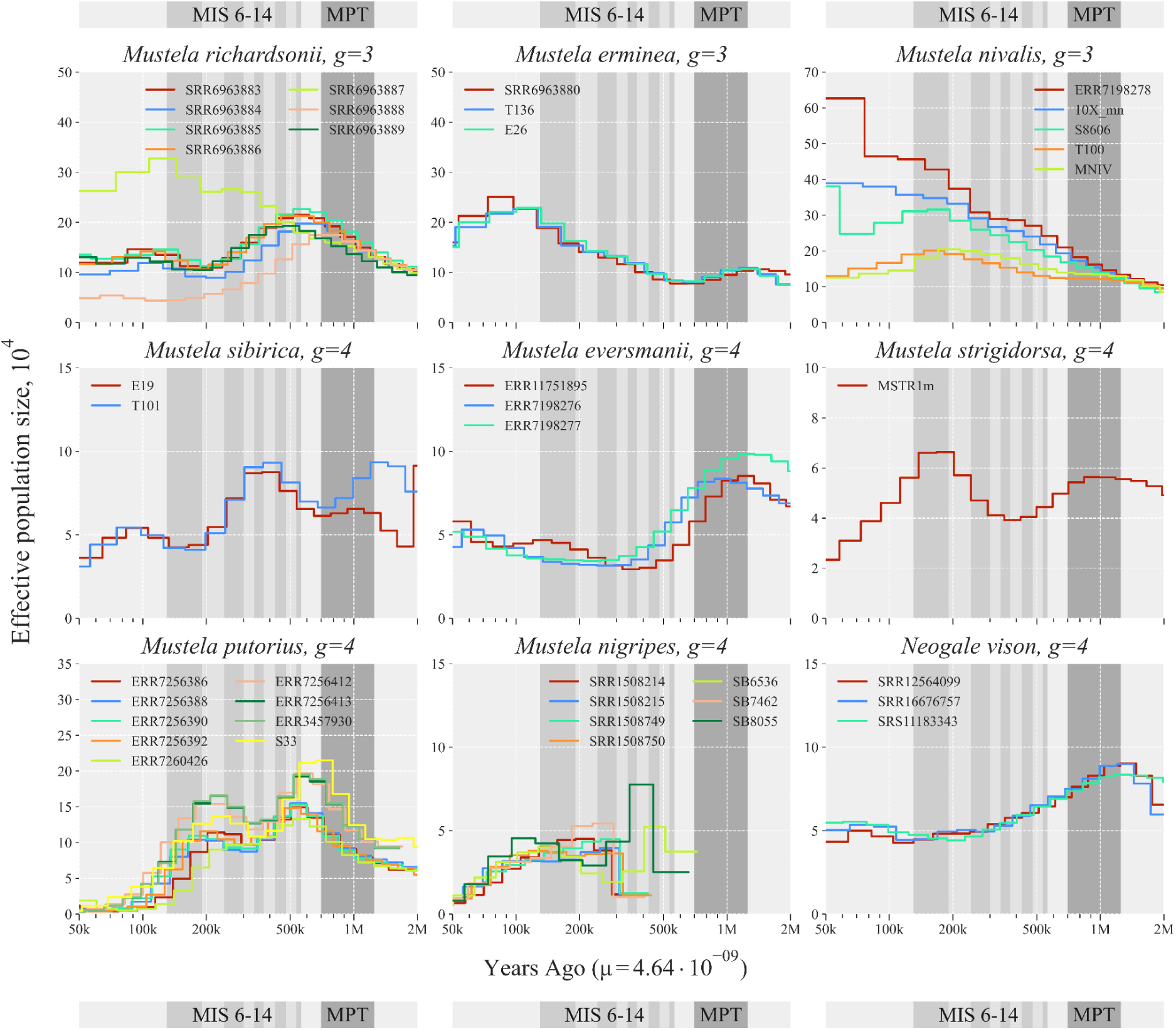
Demographic history of Mustelinae species. The X chromosome was excluded from the analysis. Parameters: *g* – generation time, *μ* – mutation rate. Abbreviations: MIS – Marine Isotope Stages: 6 (191-130 kya), 7 (243-191 kya), 8 (300-243 kya), 9 (337-300 kya), 10 (374-337 kya), 11 (424-374 kya), 12 (478-424 kya), 13 (533-478 kya), 14 (563-533 kya); MPT – Mid-Pleistocene Transition (1.25-0.7 Mya)

Among highly heterozygous species, *M. nivalis* samples from Europe showed a predominantly steady increase in N_e_ following divergence from a common ancestor shared with Asian samples around 1 Mya (MPT, confidence interval (CI): 629.6 kya – 1.6 Mya). The maximum N_e_ among European samples reached 62.6k (ERR7198278, France), exceeding values observed in both Asian samples (up to 20.4k, MNIV) and other *Mustela* species (Figure 9). Notably, the Berlin sample (S8606) exhibited a prolonged N_e_ decline beginning ∼150 kya (CI: 94.4 – 236.7 kya), followed by a sharp increase around ∼60 kya (CI: 37.8 – 94.7 kya). In contrast, the Asian samples did not show such recovery; their N_e_ declined to ∼13k and remained low. N_e_ dynamics were nearly identical among geographically proximate samples, although minor fluctuations were observed among European individuals. For *M. erminea*, represented by three samples from Asia, a steady increase in N_e_ was observed until ∼100 kya (CI: 63 – 157.8 kya), followed by a gradual decline (Figure 9). The trajectories were consistent across samples, with slight differences in maximum Ne, ranging from 22.7k (T136) to 25.1k (SRR6963880). As previously reported (Colella et al. 2018), divergence between *M. richardsonii* and *M. erminea* occurred around 2 Mya (CI: 1.3 – 3.2 Mya), followed by an increase in N_e_ reaching maxima around 570 kya (CI: 358.9 – 899.6 kya), ranging from 17.5k (SRR6963888) to 22.6k (SRR6963885). After a period of decline, the trajectory stabilized around ∼200 kya (CI: 125.9 – 315.6 kya). Remarkably, the hybrid sample SRR6963887 showed a continuous N_e_ increase (max 32.7k) until ∼150 kya (CI: 94.4 – 236.7 kya) following divergence from *M. erminea*.

Among species with lower heterozygosity levels, *M. sibirica* stands out with at least two major population bottlenecks (Figure 9). The first, ∼1.5 Mya (CI: 944.4 kya – 2.4 Mya), is likely an artifact, as sample E19 displayed anomalous fluctuations. The remaining bottlenecks (∼350 kya, CI: 220.4 – 552.4 kya, and ∼100 kya, CI: 63 – 157.8 kya) reduced N_e_ from 8.8k to 3.6k in E19, and from 9.3k to 4.1k in T101. A similar sharp N_e_ decline was observed in Asian samples of *M. eversmanii*, beginning ∼1 Mya (CI: 629.6 kya – 1.6 Mya), resulting in minimum N_e_ values: from 8.5k to 2.9k (ERR11751895), from 8.4k to 3.2k (ERR7198276), and from 9.8k to 3.4k (ERR7198277). The tropical species *M. strigidorsa* exhibited two distinct periods of N_e_ reduction. The first (∼850 kya, CI: 535.1 kya – 1.3 Mya, to 350 kya, CI: 220.4 – 552.4 kya) was followed by recovery to a maximum N_e_ of 6.6k, after which a second decline (∼150 kya, CI: 94.4 – 236.7 kya) decreased N_e_ to a minimum of 2.3k. *M. putorius* experienced the most pronounced N_e_ declines due to two sharp bottlenecks (Figure 9). Samples formed two population clusters with nearly parallel N_e_ trajectories: one group comprising samples from continental Europe, and the other from the United Kingdom. Interestingly, the Pskov sample (S33, Russia) clustered with the latter. Maximum N_e_ values differed between groups: 13.3k (ERR7260426) to 15.5k (ERR7256388) in the first, and 19.2k (ERR7256413) to 21.5k (S33) in the second. After reaching peak values, the first sharp decline occurred around ∼500 kya (CI: 314.8 – 789.1 kya), followed by a second around ∼200 kya (CI: 125.9 – 315.6 kya), reducing N_e_ to critically low levels in both groups: from 0.4k (ERR7256392) to 1.2k (S33). The demographic history of *M. nigripes*, a critically endangered species, could be reconstructed only up to ∼350 kya (CI: 220.4 – 552.4 kya), likely due to extremely low genetic diversity caused by a strong founder effect (Derežanin et al. 2025). Minimum N_e_ values ranged from 0.7k (SRR1508214) to 1.1k (SB6536). All *N. vison* samples exhibited a prolonged decline in N_e_ beginning ∼1 Mya (CI: 629.6 kya – 1.6 Mya), reaching relatively stable values around ∼300 kya (CI: 188.9 – 473.5 kya).

## Discussion

### Genome assemblies and rearrangements

In recent years, chromosome-level genome assemblies have become the gold standard in comparative and population genomics, providing highly detailed genomic structures and facilitating sophisticated genetic studies (Theissinger et al. 2023). In this work, we expanded the availability of such high-quality data for species within the Mustelinae, promoting a deeper understanding of genetic diversity within the Mustelidae family. Currently, only six chromosome-level genome assemblies and two scaffold-level assemblies are available for the Mustelinae. The quality of these assemblies varies due to the use of different sequencing technologies (Supplementary Table ST4). Notably, the genome assemblies of *M. erminea*, *M. lutreola* and *N. vison*, generated with PacBio long reads, stand out for their high scaffold N50 and completeness metrics (Supplementary Table ST4 and <u>ST6</u>). Other assemblies, including the new chromosome-level assemblies of *M. nivalis* (Supplementary Figure SF1A) and *M. strigidorsa* (Supplementary Figure SF1B), as well as the scaffold-level assembly of *M. sibirica*, are comparable in quality to the existing genomes of other *Mustela* species, with scaffold N50 115.57-138.37 Mbp (Supplementary Table ST4) and 94.6-96.1% complete BUSCOs (Supplementary Table ST6).

However, genome assemblies are still lacking for 11 species of the Mustelinae, even at the scaffold-level (Supplementary Table ST1). The main challenges are associated with the difficulty of obtaining samples, especially for tropical species such as *M. kathiah*, *M. nudipes*, and *M. lutreolina* and for insular species such as *M. haidarum* and *M. itatsi*. As part of our research, we present for the first time the genome assembly of *M. strigidorsa* (Supplementary Figure SF1B), a species found in the tropics of Southeast Asia, which is an important step in expanding the database of genomic resources.

We observed a number of chromosomal rearrangements among the genomes of the seven compared Mustelinae species (Figure 3; Supplementary Table ST12) and confirmed all known fusions and fissions from previous cytogenetic studies (Graphodatsky et al. 2002). Most of the inversions we identified previously eluded detection, as they were smaller than 10 Mbp and thus unresolvable using only cytogenetic data. However, some of the inversions we identified had been previously proposed (Graphodatsky et al. 2002), such as the pericentric inversion on NVIS 14. The improved resolution of genomic data also enabled the clarification of another predicted inversion, which actually consists of two small paracentromeric inversions that occur in MNIV 10 and not in MLUT and MNIV (Graphodatsky et al. 2002). Consequently, we show that both inversion events (MNIV 10 and MNIV 14) are specific for *N. vison*.

### Genome sizes and repeats content

According to previous cytogenetic studies, Mustelinae species are characterized by completely heterochromatic arms on originally acrocentric chromosomes (Graphodatsky et al. 1977; Graphodatsky et al. 1989). The amount of the additional heterochromatin varies from species to species, affecting their respective genome sizes (Graphodatsky et al. 1977). Heterochromatin blocks are known to consist of repetitive sequences, mainly tandem repeats, which are difficult to sequence due to high GC content and their representation in genome assemblies varies depending on library preparation methods and sequencing technologies (Biscotti et al. 2015). Moreover, during the genome assembly process, a significant portion of repeat-associated reads is often discarded (Sedlazeck et al. 2018). Our data appear to confirm that differences in heterochromatin content underlie the variability in genome size among Mustelinae species, as indicated by discrepancies between estimates derived from genome assemblies and 23-mer distribution of WGS reads (Figure 2; Supplementary Table ST5).

Genome size can be estimated using methods such as k-mer distribution analysis of WGS data (Kliver et al. 2017; Ranallo-Benavidez et al. 2020) or cytogenetic analysis (Hardie et al. 2002). Cytogenetic data suggests the euchromatic portion remains stable across Mustelidae species (2.4-2.9 Gbp, which corresponds to 2.5 to 3 pg, where 1 pg corresponds to 978 Mbp (Doležel et al. 2003)) (Graphodatsky 1989), with size differences attributed to heterochromatin variation (Graphodatsky et al. 1977). Our study revealed notable discrepancies between different measurement approaches – our genome assemblies (2.41-2.68 Gbp; Supplementary Table ST4) and 23-mer WGS-based estimates (Supplementary Table ST5) consistently yielded smaller sizes than cytogenetic measurements. Species with minimal heterochromatin content such as *M. sibirica* (13.4% ± 4.7% (Graphodatsky et al. 1977)) with an assembly size of 2.4 Gbp showed moderate differences in genome size (cytogenetic: 3.0-3.1 Gbp = 3.1 to 3.2 pg vs. WGS: ∼2.54-2.59 Gbp), while those with substantial heterochromatin like *M. nivalis* (34.5% ± 3.2% heterochromatin (Graphodatsky et al. 1977)) with an assembly size of 2.45 Gbp exhibited more dramatic disparities (cytogenetic: 4.4 Gbp = 4.5pg vs. WGS: 3.08-3.35 Gbp). In other Mustelinae species, such as *M. eversmanii*, *M. putorius*, *M. erminea* and *N. vison*, the heterochromatin content is 20.6% ± 2.2%, 22.7% ± 3.4%, 22.8% ± 3.5% and 25.4% ± 4.7%, respectively (Graphodatsky et al. 1977). These inconsistencies likely stem from incomplete assembly of heterochromatic regions in genome sequencing approaches, compounded by the high error margins (up to 15%) in heterochromatin quantification methods (Graphodatsky et al. 1983; Graphodatsky 1989).

### Heterozygosity and Runs of Homozygosity

Numerous studies highlight the critical importance of maintaining genome-wide genetic variation to enhance population viability (Lande and Shannon 1996; Abascal et al. 2016; Bozzuto et al. 2019), although some researchers have critiqued traditional conservation genetic approaches based on neutral variation by placing a greater emphasis on functionally significant genetic variation (Robinson et al. 2016; Robinson et al. 2019; Teixeira and Huber 2021). Despite ongoing debates, there is broad consensus that a reduction in genome-wide genetic diversity can have important consequences for the long-term viability of populations, though its impact may vary depending on genetic and ecological contexts. For instance, some studies suggest that the loss of heterozygosity in small populations is not inherently detrimental, as it may reduce the frequency of deleterious recessive alleles (Vidal et al. 2025). However, reduced genetic diversity can also limit adaptive potential and increase vulnerability to environmental changes. Mustelinae species, like others, are subject to varying climatic and ecological pressures, leading to significant fluctuations in their genetic diversity (Colella et al. 2018; Derežanin et al. 2022; de Ferran et al. 2022). These fluctuations influence their ability to adapt to environmental changes. In our study, we evaluated genetic diversity by assessing and comparing heterozygosity and homozygosity within and between species, as well as investigating their distribution within the genomes (Supplementary Table ST8, <u>ST9</u>). Although our data do not cover population-level genetic diversity, they provide an initial genome-wide overview of variability for both widespread and geographically restricted *Mustela* species.

We found considerable intraspecific and interspecific differences in mean heterozygosity between analyzed *Mustela* species: from 0.02 SNPs/kbp to 2.9 SNPs/kbp (Figure 4; Supplementary Table ST8). The most heterozygous species included *M. nivalis* (2.64 SNP/kbp), *M. erminea* (2.03 SNP/kbp) and *M. richardsonii* (2.22 SNP/kbp). These species have extensive ranges (Figure 1). For example, *M. nivalis* and *M. erminea* inhabit nearly the entire Holarctic region, whereas *M. richardsonii* is restricted to North America. The observed high genetic diversity in *M. nivalis* samples aligns with previous studies based on different molecular markers, including mitochondrial sequences (Lebarbenchon et al. 2010; Sato et al. 2020; Tissaoui et al. 2024). Despite the high mean heterozygosity of this species, we observed clear signs of recent inbreeding, indicated by the presence of ultra long RoH in samples such as T100 (Figures 4A, 5; Supplementary Table ST9) covering significant portions of their chromosomes. While mean genetic diversity is maintained at a high level, monitoring of *M. nivalis* shows declines in local populations (e.g. USA (Jachowski et al. 2021), UK (Sainsbury et al. 2019), Spain (Llorca et al. 2024) and Tunisia (Hayder et al. 2023)).

In contrast, we observed significantly lower heterozygosity and extensive RoH in the less widespread *Mustela* species. For example, the single sample of the tropical species *M. strigidorsa* from Vietnam has a mean heterozygosity of 0.62 SNP/kbp only (Figure 10, Supplementary Table ST8), and RoH cover 23.62% of the genome (Supplementary Figure SF5; Supplementary Table ST9). This study provides the first such assessment of this understudied species. There are concerns regarding habitat degradation and fragmentation for this species (Abramov et al. 2015), but their effect on *M. strigidorsa* population size remains unknown due to limited data (Abramov et al. 2008). We noted both long (106 or 10.21%) and ultra long (3 or 1.81%) RoH in this species’ genome. While *M. strigidorsa* exhibits relatively moderate homozygosity, exceptional cases of extreme homozygosity within *Mustela* are notably found in *M. nigripes*, a species that once was declared extinct (Fricke 2015), where RoH cover at least 87% of the genome (Figure 5; Supplementary Table ST9), and heterozygosity levels range from 0.02 to 0.04 SNP/kbp (Supplementary Table ST8) (Derežanin et al. 2025) – significantly lower than those of the domesticated *M. putorius furo* (0.13-0.25 SNP/kbp).

**Figure 10.**
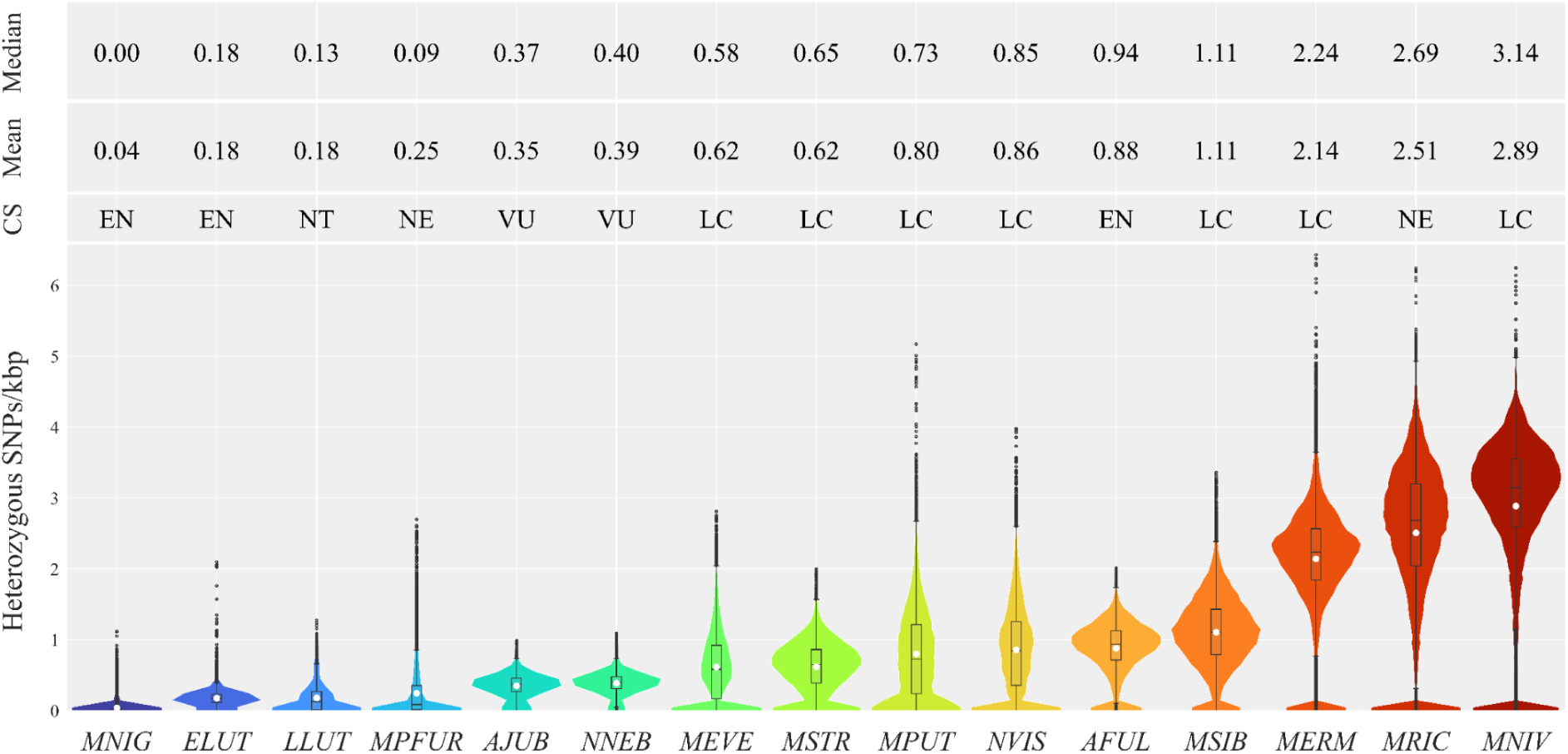
Distribution of the heterozygous SNP density. Heterozygous SNPs were counted in 1 Mbp windows and 100 kbp steps and scaled SNPs/kbp. Only samples with highest mean values are shown (one sample per species, Supplementary Table ST8). Median and mean heterozygosity values are indicated within boxplots by black line and white dots, respectively. Color gradient indicates the level of heterozygosity, ranging from lower (blue) to higher (red) values. Abbreviations: GS – Global conservation status according to IUCN Red List data: LC – Least Concern, NT – Near Threatened, VU – Vulnerable, EN – Endangered, NE – Not Evaluated; Species: MNIG – *Mustela nigripes*, ELUT – *Enhydra lutris*, LLUT – *Lutra lutra*, MPFUR – *Mustela putorius furo*, AJUB – *Acinonyx jubatus*, NNEB – *Neofelis nebulosa*, MEVE – *Mustela eversmanii*, MSTR – *Mustela strigidorsa*, MPUT – *Mustela putorius*, NVIS – *Neogale vison*, AFUL – *Ailurus fulgens*, MSIB – *Mustela sibirica*, MERM – *Mustela erminea*, MRIC – *Mustela richardsonii*, MNIV – *Mustela nivalis*

Another example is *M. eversmanii*, represented in our study by three samples from its Asian range, which exhibit relatively low mean heterozygosities (0.54, 0.58 and 0.62 SNPs/kbp; Supplementary Table ST8). These samples contain mostly ultra long RoH, covering large portions of the autosomes and encompassing 14.49-19.24% of the genome (Supplementary Figure SF5; Supplementary Table ST9). *M. eversmanii* is considered to be rapidly disappearing in several European countries (Šálek et al. 2013), facing severe threats including habitat loss and fragmentation, declining prey base, competition and rodenticide poisoning (Šálek et al. 2013; Szapu et al. 2024). Conversely, a recent study based on low-resolution data challenges the alarming rate of decline in Europe, suggesting either insufficient research or a rapid population recovery following a sharp decline (Szatmári et al. 2021). Assessments in its Asian range are scarce, but reports indicate a significant decline in Russia (FGU Centrokhotkontrol 2007; Minprirody of Russia 2019; Minprirody of Russia 2021; Kiseleva 2024; Minprirody of Russia 2024), where the combined population estimate of *M. eversmanii* and *M. putorius* (due to nearly indistinguishable tracks (FGU Centrokhotkontrol 2007)) dropped from 90.6 to 48.3 thousand individuals between 2003 and 2023 (Figure 11; Supplementary Table ST13) (FGU Centrokhotkontrol 2007; Minprirody of Russia 2019; Minprirody of Russia 2021; Minprirody of Russia 2024). *M. eversmanii* and *M. putorius* are believed to contribute approximately equally to these counts, with an assumed geographic division according to their ranges (Figure 1) (FGU Centrokhotkontrol 2007). The causes of *M. eversmanii* decline in Russia remain unclear, with habitat loss considered minor due to protected range areas (Kiseleva 2024) and illegal hunting limited by the low commercial value of its fur (Minprirody of Russia 2024). Competition with the invasive *N. vison*, known for its aggressiveness and infanticide, has been suggested as a contributing factor (Maran et al. 1998; Sidorovich and Macdonalds 2001; MaCdonald and Harrington 2003; Croose et al. 2018; Kiseleva 2024) with similar concerns in Central Asia, where *N. vison* is actively spreading into Kazakhstan (Maran et al. 2015). The diet of *M. eversmanii* in Asia is primarily composed of rodents, particularly the red-cheeked ground squirrel *Spermophilus erythrogenys* (Moskvitina et al. 2023), which is considered an agricultural pest (Shilova 2011). The two species are tightly bound not only by the predator-prey relationship, but also by the fact that steppe polecats habitually den in ground squirrel burrows. In Russia, efforts to control ground squirrels began as early as the 1920’s and involved a wide range of methods, including the use of selective strains of erysipeloidum and salmonellosis in the late 20th century (Shilova 2011; Vazhov et al. 2016). Introduction of new pesticides and intensive agricultural methods may have played a role as well (Wright et al. 2022). As a result, the population of *S. erythrogenys* has declined sharply, and in some regions, including Novosibirsk and Altai Krai, the species is extremely rare (Vazhov et al. 2016; Moskvitina et al. 2023). This inevitably affected the population of *M. eversmanii*, as a reduction in food availability leads to a decline. Considering the significant threats across its range, *M. eversmanii* populations will probably continue to decline in both European and Asian regions.

**Figure 11.**
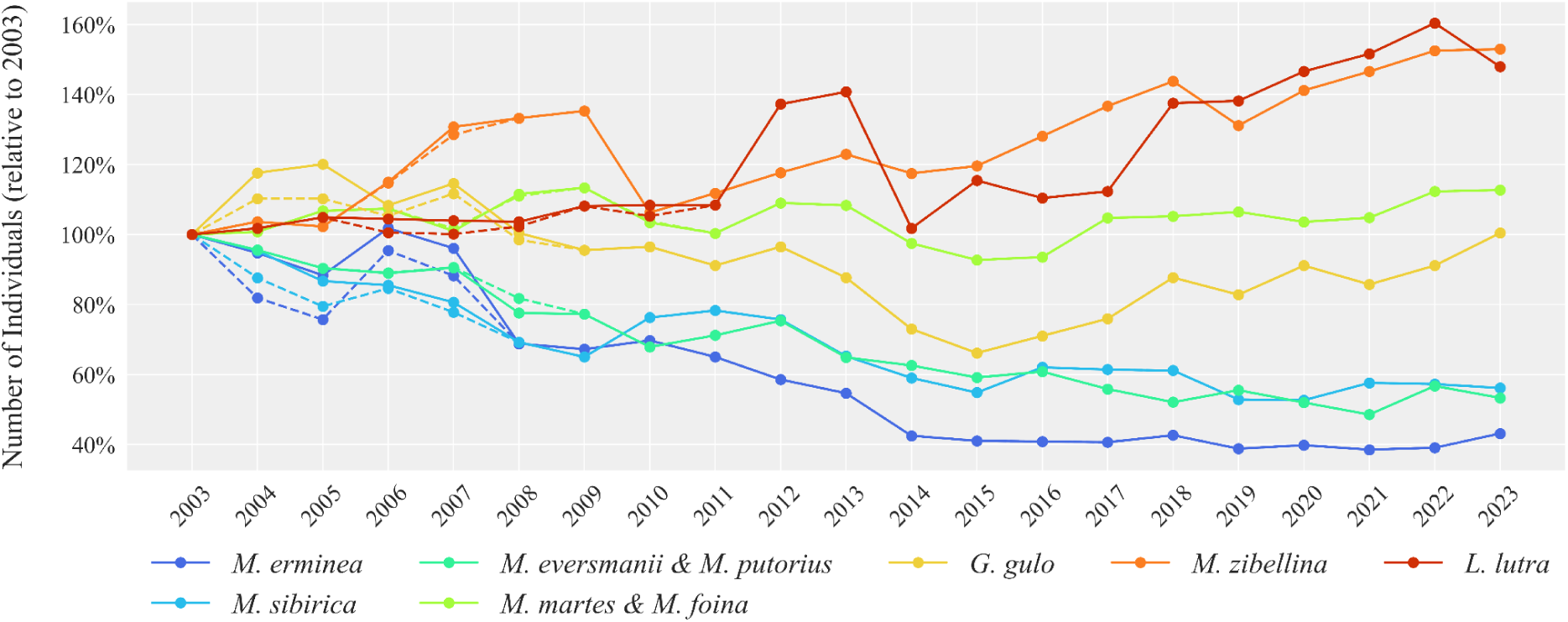
The main fur game animal resources in the Russian Federation for the years 2003-2023. Data are based on public reports of the Ministry of Natural Resources and Environment of the Russian Federation and the Federal Research Center for Hunting Development (FGU Centrokhotkontrol 2007; Minprirody of Russia 2019; Minprirody of Russia 2021; Minprirody of Russia 2024). Dotted lines indicate discrepancies in the datasets (Supplementary Table ST13)

*M. putorius* shows low genetic diversity (mean heterozygosity 0.43 SNPs/kbp) (Supplementary Table ST8) and high RoH (15.84%-59.28%) (Supplementary Table ST9). Populations are declining in Europe, with factors overlapping those affecting *M. eversmanii*, such as competition with invasive predators and hybridization with other *Mustela* species (Croose et al. 2018). While hybridization with *M. putorius furo* has contributed to population recovery in the UK (Shaw et al. 2024), increasing competition with *N. vison* remains a threat (Croose et al. 2018).

### Phylogeny, admixture, and ancestral rearrangements

Previously, the phylogeny of the Mustelinae was reconstructed on the basis of concatenated sets of nuclear and/or mitochondrial markers (Koepfli, Deere, et al. 2008; Sato et al. 2012; Law et al. 2018; Hassanin et al. 2021). We performed such a reconstruction using whole genome data for the first time, including new genome assembly and resequencing data of two Asian species - *M. strigidorsa* and *M. sibirica*. *M. strigidorsa*, which is distributed in the tropical regions of Southeast Asia, occupied a basal position to other *Mustela* species in the phylogenetic trees (Figure 6, 8; Supplementary Figures SF7). For three more tropical Asian species (*M. kathiah*, *M. nudipes* and *M. lutreolina*) whole genome data is still unavailable, but mitochondrial genome reconstructions suggest successive splitting of *M. nudipes* and *M. kathiah* from the stem (Hassanin et al. 2021), while concatenated nuclear + mitochondrial DNA datasets group *M. nudipes* with *M. strigidorsa* together as sister taxa (Koepfli, Deere, et al. 2008; Sato et al. 2012; Law et al. 2018). However, mtDNA trees might be unreliable in case of mustelids, due to active interspecies hybridization within this family, which was reported for multiple species: badgers, martens, ermines, ferrets and others (Rozhnov et al. 2013; Kinoshita et al. 2019; Colella et al. 2021; Szatmári et al. 2021; Etherington et al. 2022; Tomarovsky et al. 2025). Breeding in captivity has also revealed that distantly related species such as *M. putorius* and *M. sibirica* are capable of hybridizing (Graphodatsky et al. 1982; Graphodatsky et al. 1985). In our study we observed some discordance between nuclear (Figures 6) and mitochondrial (Figures 8) trees. For example, the *M. putorius* ERR7260426 (BK069848) sample was previously shown to have such a discordance (Etherington et al. 2022); inclusion of all available genome data from the Mustelinae clarified that this sample carries a *M. lutreola* mitochondrial genome.

Our species tree (Supplementary Figures SF7, <u>SF8</u>) supports the distinction between *N. vison* and the genus *Mustela*, with high main posterior probability support (pp1=1.0). While quartet support for this split was somewhat lower than for other interspecific nodes (q1=61.7%), it remains above the threshold typically considered indicative of primary topological signal. This moderate value may reflect inherent gene tree discordance due to incomplete lineage sorting or ancient gene flow at the early stages of Mustelinae diversification. Nonetheless, both concatenated and coalescent-based analyses consistently recover *N. vison* as a sister lineage to *Mustela*, supporting the current taxonomic treatment that recognizes *Neogale* as a distinct genus. While most interspecific nodes were characterized by high quartet support, a notable exception is node 9, uniting *M. sibirica* and *M. lutreola*, which showed markedly reduced support (q1=47.7%), likely reflecting conflicting gene tree signals in this part of the phylogeny. At the same time, the position of *M. strigidorsa* as the earliest branching lineage within *Mustela* is strongly supported, further reinforcing its placement within this genus.

Our analysis underscores the critical role of incorporating heterozygous positions in phylogenetic reconstruction, particularly when dealing with individuals that may have hybrid ancestry.

In our study, the placement of the *M. nivalis* reference individual (red background in Supplementary Figure SF10) shifted dramatically depending on whether a haploid assembly or a heterozygous SNP set derived from read alignments was used. This discordance suggests that the haploid consensus sequence may not adequately represent the genetic makeup of admixed individuals, potentially masking one of the parental signals. The 10X_mn genome may carry introgressed segments, and collapsing its heterozygous sites into a single haplotype can mislead phylogenetic inference by emphasizing one ancestral lineage over the other. While the effect was less pronounced for *M. sibirica* (green), *M. strigidorsa* (blue), and *M. eversmanii* (orange), branch lengths were still affected. Mustelidae are known to hybridize, and our findings highlight how such genetic complexity can confound phylogenetic inference when standard haploid assemblies are used.

*M. erminea sensu lato* (all stoats) and *M. nivalis* are the Mustelinae species with the largest ranges, encompassing nearly the whole Holarctic (Figure 1). The combination of such a wide distribution and limited dispersal due to small body sizes suggested to researchers that these two taxa might contain multiple species. Recently, *M. erminea sensu lato* was split into three species: *M. erminea (sensu stricto*, Eurasia and Alaska), *M. richardsonii* (continental America) and *M. haidarum* (Haida Gwaii and Alexander Archipelago of coastal North America) (Colella et al. 2021). However, this suggestion was based on mitochondrial sequences and morphological traits only. By combining samples from a previous dataset (Colella et al. 2018) and sequencing two more *M. erminea* individuals from Yakutia, Russia, we confirmed the hypothesis of multiple species (Supplementary <u>MR</u>). Due to the lack of samples, the status of European *M. erminea* remains unclear. For *M. nivalis* we found that despite clear division of Western European and Asian samples, pairwise distances between them are comparable to distances within *M. erminea* (Figure 7 and Supplementary Figure SF6), i.e., no signs of the speciation was observed. However, it is important to note that within-species divergence was assessed for Eurasian *M. nivalis* only, due to the lack of samples from North America.

The order Carnivora was suggested to have an ancestral karyotype with 2n=38 (Nash et al. 2008; Perelman et al. 2012; Beklemisheva et al. 2016), which is observed across a wide range of terrestrial and semiaquatic species (Graphodatsky et al. 2020). Within Carnivora, the Mustelidae also exhibits a predominantly syntenic conservatism with 2n=38 in most species (Graphodatsky et al. 2020). However, several lineages within Mustelidae show notable variation in chromosome number, reflecting independent chromosomal rearrangements. One such example is the Mustelinae, which displays considerable karyotypic variability (Figures 3, 6). Within this group, the diploid number ranges from 2n=30 in *N. vison* to 2n=44 in species such as *M. strigidorsa*, *M. kathiah* (Abramov et al. 2013), *M. erminea* and *M. altaica* (Graphodatsky et al. 2020). This variability likely reflects a history of recurrent chromosomal fissions and fusions, which have contributed significantly to the evolutionary diversification of the subfamily.

The results of our study allow us to clarify the timing of the three latest chromosomal fissions in Mustelinae (Figure 6). All three rearrangements occurred after the divergence of Mustelinae from other Mustelidae lineages and at early stages of the subfamily’s radiation (Supplementary Figure SF9). These events were previously identified (Graphodatsky et al. 2002), but two of them are now confirmed to have taken place before the diversification of extant Mustelinae, while the timing of the third fission has been revised based on our data. Earlier hypotheses suggested that the fission fis(MSTR2, MSTR21) occurred after the divergence of the *Neogale* lineage but before the separation of *M. erminea* (Graphodatsky et al. 2002). The inclusion of *M. strigidorsa* in our phylogenetic analysis provides important new evidence. This species occupies a basal position within the *Mustela* clade (Figures 6, 8; Supplementary Figure SF7), which shifts the previously proposed timing of the fission event. Additional support for the ancestral Mustelinae rearrangements comes from data on other tropical *Mustela* species. For example,

*M. kathiah* has been shown in previous studies (Sato et al. 2012; Law et al. 2018), as well as in our mitochondrial genome analysis (Figure 8), to have diverged after *M. strigidorsa*. Furthermore, *M. nudipes* is phylogenetically close to *M. strigidorsa* (Koepfli, Deere, et al. 2008; Sato et al. 2012; Law et al. 2018), and *M. lutreolina* is another tropical species potentially related to this group. However, the phylogenetic position of *M. lutreolina* remains uncertain, as no cytogenetic or genomic data are currently available for this species. These observations support the inference that the ancestral *Mustela* karyotype most likely had a diploid number of 2n=42 (Figure 12).

**Figure 12.**
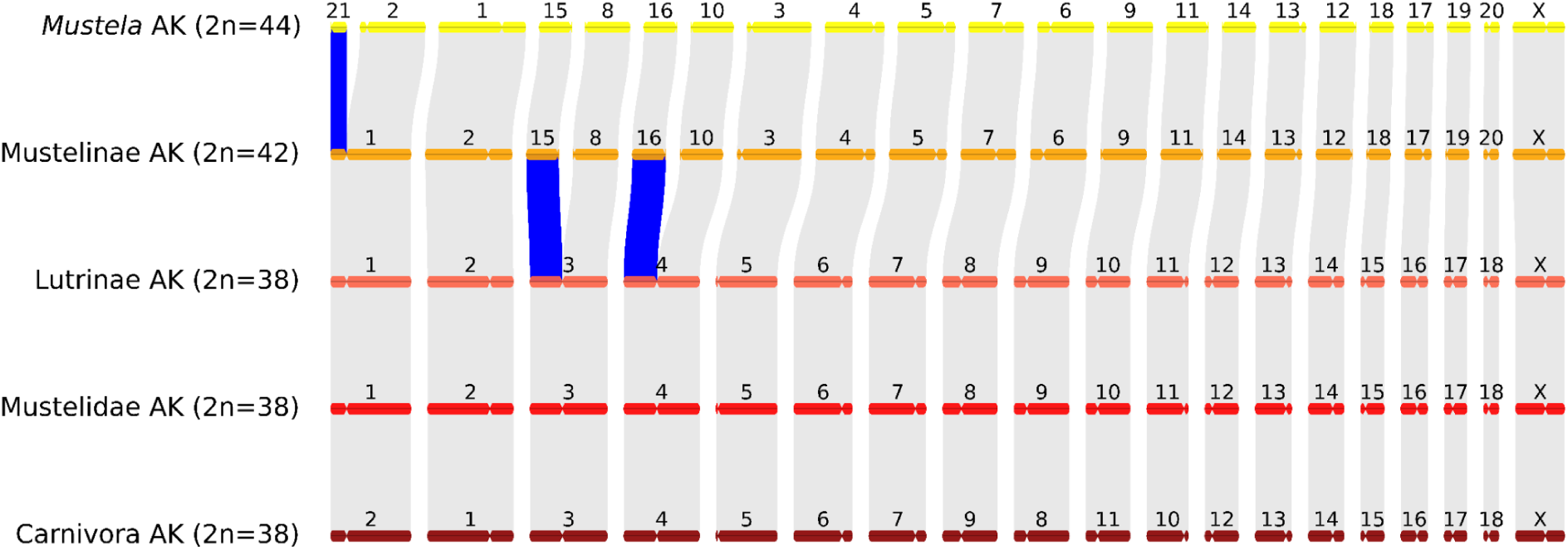
The chromosomal synteny (scheme) between ancestral karyotypes (AK) of Carnivora, Mustelidae, Lutrinae, Mustelinae and *Mustela*. Fissions are highlighted in blue. Chromosome numbering for Mustelidae AK and Lutrinae AK is based on that of *Martes foina*, for Carnivora AK we followed the previously proposed nomenclature (Beklemisheva et al. 2016), for Mustelinae AK we used *M. strigidorsa* nomenclature with modification for chr 1 and 2, and for *Mustela* AK it completely corresponds to *Mustela strigidorsa*

The ancestral karyotype of the genus *Neogale* remains uncertain. In *N. vison*, several chromosomal fusions have been identified, resulting in a reduced chromosome number of 2n=30 (Figure 3). However, it is not yet clear whether these chromosomal rearrangements are specific to this species or represent a broader pattern shared across the genus. Further studies, including data from other *Neogale* species are necessary to address this question.

### Demographic history

Сomparative analysis of demographic history revealed significant variation in effective population size among *Mustela* species (Figure 9), likely reflecting both biogeographic history and species-specific responses to Pleistocene climate fluctuations. Although each species exhibits its own pattern of N_e_ change, several common trends indicate the influence of the same large-scale environmental factors.

*M. nivalis* and *M. erminea* have a relatively high heterozygosity (Figure 4B) and broad ecological tolerance (Sommer and Crees 2022), likely promoting demographic stability during climatic oscillations. Both show either continuous growth or consistently elevated N_e_ during MIS (Marine Isotope Stage) 14-6 a period of strong climatic instability (Figure 9). These patterns are consistent with their persistence across glacial–interglacial cycles and broad geographic ranges (Sommer and Benecke 2004; Sommer and Crees 2022). Fossil evidence further supports this (Sommer and Benecke 2004; Marciszak and Socha 2014; Krajcarz et al. 2015; Kosintsev et al. 2016; Crégut-Bonnoure et al. 2018; Giustini et al. 2024): *M. nivalis* (including its ancestral form *M. praenivalis*) and *M. erminea* are represented by 34 and 17 Pleistocene–early Holocene records, mostly from Europe and North America (PBDB 2025). *M. nivalis* shows a structured demographic history, with divergence between western (European) and eastern (Asian) lineages around 1 Mya (Figure 9), during the MPT (Mid-Pleistocene Transition). Western populations display sustained N_e_ growth, while eastern ones remain stable. Elevated N_e_ in Europe may indicate historical gene flow, though this requires testing via expanded sampling and formal modeling. A similar pattern was observed in the hybrid *M. richardsonii* × *M. erminea* individual (SRR6963887), likely due to recurrent gene flow (Colella et al. 2018). In the case of SRR6963887, introgressive hybridization is well documented (Colella et al. 2018), whereas for European *M. nivalis*, the elevated N_e_ may reflect high intraspecific diversity, a hypothesis that remains to be investigated using a larger sample set. Similar patterns are also seen in other Mustelidae hybrids like *M. zibellina* × *M. martes* (Tomarovsky et al. 2025). Although PSMC does not explicitly infer admixture, such signatures offer indirect evidence of complex histories. The demographic stability in eastern *M. nivalis* is consistent with a possible Asian origin and later westward expansion. *M. erminea* shows a broadly similar trajectory, with N_e_ increase during MIS 14-6 followed by decline near the end of MIS 6. However, due to sampling from eastern populations only, spatial inferences are limited. Nevertheless, the concordant demographic patterns observed in *M. nivalis* and *M. erminea* support the hypothesis that their persistence was shaped by shared ecological mechanisms, particularly their high adaptability to cold climates during periods of Quaternary climatic instability.

In contrast, species with lower heterozygosity such as *M. sibirica*, *M. eversmanii*, *M. strigidorsa*, *M. putorius*, and *M. nigripes* show evidence of recurrent or prolonged bottlenecks (Figure 9). *M. putorius* widespread in Europe (Figure 1), experienced two sharp N_e_ declines during MIS 12 and MIS 6. Despite this, it is considered an early post-LGM (Last Glacial Maximum) recolonizer of Central Europe, supported by fossil data (Sommer and Benecke 2004; Sommer and Crees 2022). Other species, sampled from Asia, show bottlenecks around the onset of the 100-kyr glacial cycles (Figure 9). Although Asia is thought to have been less impacted by Quaternary climate shifts (Fu and Wen 2023), our results suggest substantial demographic changes even in steppe and tropical-adapted species. Fossil data for these taxa remain limited (Krajcarz et al. 2015; Kosintsev et al. 2016; Giustini et al. 2024; Fourvel et al. 2025): *M. eversmanii* is known from seven European Pleistocene sites (including *M. stromeri*, its common ancestor with *M. putorius* and *M. nigripes*); *M. sibirica* from two Japanese finds; and *M. strigidorsa* lacks confirmed Pleistocene records (PBDB 2025). Southern Polish (Krajcarz et al. 2015) and Ural (Kosintsev et al. 2016) fossils suggest a broader range for *M. eversmanii* during the Pleistocene, later restricted to Eastern European steppes. The North American species, *M. nigripes*, exhibits a truncated N_e_ trajectory (Derežanin et al. 2025), consistent with extreme founder effects and its well-documented population collapse (Fricke 2015).

Overall, *Mustela* demographic histories highlight how long-term population dynamics reflect the interplay of environmental change and climatic history. Future studies incorporating broader genomic sampling, paleodistribution models, and molecular dating are crucial to provide a more comprehensive overview.

### Conclusions and perspectives

This study represents the first comprehensive analysis of the Mustelinae at the whole genome level, in which we thoroughly investigated phylogenetic relationships, assessed levels of genetic diversity, reconstructed demographic histories and identified structural rearrangements using chromosome-scale scaffolds. One of the key findings of this study is the identification of significant variation in heterozygosity and homozygosity, both at the interspecific and intraspecific levels. Our data highlight the relevance of genetic diversity measures in conservation assessments. Specifically, *M. eversmanii* exhibits low genome-wide genetic diversity in its Asian range, alongside population declines in its European range and in Russia, where the majority of its global population resides. Similarly, *M. putorius* shows signs of population decline, increased inbreeding, and low genetic diversity despite its current classification as a species of Least Concern (Maran et al. 2016). While these findings alone do not directly imply a need for Red List reclassification, they suggest that a more detailed evaluation of the genetic and demographic trends in these species could be valuable in refining conservation priorities. Although some species have persisted with low genetic diversity over long evolutionary timescales, such cases are typically associated with historically small populations that have reached equilibrium (Robinson et al. 2018; Westbury et al. 2018; Robinson et al. 2022; Yang et al. 2025). In contrast, populations experiencing rapid declines from previously high levels of heterozygosity may follow a different trajectory, as reduced efficacy of purifying selection could lead to the accumulation of deleterious alleles (Robinson et al. 2023). To better understand the potential implications for long-term viability, further studies incorporating analyses of runs of homozygosity, effective population size dynamics, and functional consequences of genetic variation are needed.

Our study presents the first phylogenomic reconstruction of the genus *Mustela* based on whole-genome data, allowing for a more precise understanding of evolutionary relationships among species. Nuclear and mitochondrial genome data revealed largely congruent topologies, yet some discrepancies were observed, likely resulting from interspecific hybridization, which was reported not only for Mustelinae (Colella et al. 2018; Szatmári et al. 2021; Etherington et al. 2022), but for other Mustelidae lineages, including badgers (Melinae), otters (Lutrinae), and martens (Guloninae) (Rozhnov et al. 2013; Moretti et al. 2017; Kinoshita et al. 2019; Tomarovsky et al. 2025). Our results support the previously suggested split of *M. richardsonii* and *M. erminea* but do not provide evidence of speciation within *M. nivalis*, despite the observed geographic differentiation of its populations. Demographic history analysis revealed diverse trajectories of effective population size changes among *Mustela* species, reflecting their unique evolutionary histories. Major shifts in N_e_ correlate with key climatic events such as glacial and interglacial periods during the Pleistocene, which likely contributed to range expansions and population radiations in certain species, including *M. nivalis* and *M. erminea*. At the same time, species with more restricted distributions, such as *M. sibirica* and *M. strigidorsa*, experienced significant population declines.

These findings highlight the importance of an integrative approach to studying Mustelinae evolution, combining nuclear and mitochondrial data while accounting for factors influencing demographic processes and phylogenetic structure. Further research on intraspecific structure, genetic diversity, and conservation strategies remains a crucial direction for future studies in this field.

## Materials and methods

### Samples, DNA extraction and sequencing

To generate data for the chromosome-level genome assembly, we used a primary fibroblast cell line from male *M. strigidorsa* (MSTR1m, AVA 19-021, origin: Quang Nam, Vietnam). The cell lines were provided by the large-scale research facilities “Cryobank of cell cultures” Institute of Molecular and Cellular Biology, Siberian Branch of the Russian Academy of Sciences (IMCB SB RAS). We used frozen muscle tissue from a male *M. sibirica* (MSIB1m, origin: Republic of Sakha, Russia) to generate a scaffold-level genome assembly. For whole genome resequencing of *M. nivalis*, *M. erminea*, *M. sibirica* and *M. putorius*, we used both muscle tissue and primary fibroblast cell lines. The resulting dataset, combined with publicly available data, consisted of 50 whole-genome samples representing 10 species from the Mustelinae. A detailed description of the resequencing data is provided in Supplementary Table ST14 and includes both previously published and new data generated within our study.

DNA extraction was performed using the standard phenol-chloroform protocol (Sambrook and Russell 2006). The extracted DNA was fragmented using a Covaris instrument to reach the desired insert size and libraries were constructed with the TruSeq DNA PCR-Free kit (Illumina, Inc., San Diego, CA, USA). For the chromosome-level genome assemblies, we generated Hi-C libraries following the original protocol (Rao et al. 2014). All prepared libraries were sequenced with paired-end 150 bp reads on the Illumina NovaSeq 6000 platform. An Oxford Nanopore Technologies library was also prepared for the *M. strigidorsa* sample using the SQK-ULK114 kit and sequenced on a 10.4.1 PromethION flow cell with triple loading.

#### Assembly of the *Mustela strigidorsa* genome

We assembled the *M. strigidorsa* genome using Oxford Nanopore long reads (SRR30615031), generated as part of this study, and Hi-C (SRR34091397) and Illumina libraries (SRR34068022), generated by the DNA Zoo consortium (Dudchenko et al. 2017). We removed adapters from the Oxford Nanopore data using Porechop v0.5.0 with *ab initio* adapter detection turned off, as in the test runs with the “--ab_initio” option we detected trimming of mustelid-specific sequences present in the genome assembled using different technologies (Wick et al. 2017). Next, we filtered the data by quality (only reads with mean quality >7 were retained) and trimmed 30 bp from each end using the Chopper v0.8.0 tool (De Coster and Rademakers 2023).

We assembled the genome of the male *M. strigidorsa* using Illumina, Oxford Nanopore and Hi-C data. First, an initial draft assembly was generated from a single paired PCR-free Illumina library using w2rap-contigger (Clavijo et al. 2017), see (Dudchenko et al. 2018) for details. On the second step, the generated assembly was scaffolded using the SAMBA (script *samba.sh* with “-m 2500” option) tool (Zimin and Salzberg 2022) from MaSuRCA package v 4.1.0 (Zimin et al. 2013) and the filtered Nanopore data. Errors introduced by the long reads were corrected using the POLCA (from the same version of MaSuRCA) polisher (Zimin and Salzberg 2020) and the Illumina data. Next, we aligned the Hi-C data to the corrected intermediate assembly using Juicer v1.6 (Durand et al. 2016), followed by scaffolding to chromosome level using 3D-DNA pipeline v210623 (Dudchenko et al. 2017) with subsequent manual curation in Juicebox v2.16.00 (Dudchenko et al. 2018). Finally, we closed gaps using the SAMBA script *close_scaffold_gaps.sh* (with option “-m 2500”) and the Oxford Nanopore reads and performed a second and third round of polishing.

#### Assembly of the *Mustela nivalis* genome

We upgraded the publicly available draft genome assembly of *M. nivalis* (GCA_019141155.1) to chromosome-level using the Hi-C sequencing data (SRR34082581), generated by DNA Zoo Consortium. Hi-C data were aligned to the draft genome assembly using Juicer v1.6. Before alignment, the restriction sites for *Csp6I*+*MseI* were generated using *generate_site_positions.py* script in Juicer. Haplotype duplications in the alignments were identified using purge_dups v1.2.6 (Guan et al. 2020), based on sequence similarity and coverage. Regions where more than 90% of the sequence was involved in haplotype duplications were removed. Each deduplicated alignment was then processed using the 3D-DNA pipeline v210623 with the default parameters. A manual curation was performed using Juicebox v2.16.00.

#### Assembly of the *Mustela sibirica* genome

To generate a draft genome assembly for *M. sibirica*, we used w2rap-contigger with the default parameters to assemble paired-end Illumina reads from sample E19 (SRR30226579). Subsequently, this assembly was scaffolded using the *M erminea* genome assembly (GCF_009829155.1) as a reference. For construction of such a pseudochromosome assembly we used RagTag v2.1.0 (Alonge et al. 2022) with default parameters.

#### Pairwise whole-genome alignments and connection between assembly and karyotype

To determine the nomenclature of chromosomal scaffolds in the *M. nivalis* genome assembly, pairwise whole-genome alignments were performed between the obtained assembly and publicly available chromosomal-level genome assemblies of closely related species: *N. vison* (GCF_020171115.1), *M. lutreola* (GCF_030435805.1) and *M. erminea* (GCF_009829155.1). To determine the nomenclature of chromosomal scaffolds in the *M. strigidorsa* genome assembly, pairwise genome-wide alignment was performed only between the resulting assembly and the *M. erminea* assembly. Initially, tandemly organized and dispersed repeats in the genome assemblies were identified using Tandem Repeats Finder v4.09.1 (Benson 1999) with parameters “2 7 7 80 10 50 2000-l 10”, WindowMasker v1.0.0 (Morgulis et al. 2006) with default parameters, and RepeatMasker v4.1.6 (Tarailo-Graovac and Chen 2009) with the parameter “-species carnivora”. Tandem Repeats Finder and RepeatMasker were run using the Dfam TETools v1.88.5 container (https://github.com/Dfam-consortium/TETools). Subsequently, BEDTools v2.31.0 (Quinlan and Hall 2010) with the “-soft” parameter was employed to mask the genome assemblies using the identified repeat elements. The masked genome assemblies were then used for whole-genome alignments. Indexed masked genome assemblies were created with lastdb from the LAST v1519 (Kiełbasa et al. 2011) program using parameters “-c-u YASS-R11.” Alignments were performed with lastal from the LAST package using the parameters “-R11-f MAF.” The alignment results were visualized with the *dotplot_from_last_tab.py* script from the MAVR v0.97 software package (https://github.com/mahajrod/MAVR). We assigned chromosome numbers to the chromosomal scaffolds of the genome assemblies for *M. nivalis* according to previously established nomenclature (Graphodatsky et al. 2020), and the *M. strigidorsa* assembly based on one-to-one synteny with *M. erminea*. For this we combined our pairwise whole-genome alignments with published Zoo-FISH data of (*N. vison* probes vs *M. nivalis* and *M. lutreola* chromosomes (Graphodatsky et al. 2002)), as well as G-banding karyotypes of these species (Graphodatsky et al. 2020). Finally, all chromosomal scaffolds were renamed using the prefix “chr.”

#### Multiple whole-genome alignment, detection of synteny blocks and localization of centromeres

The chromosomal-level genome assemblies were used for synteny block analysis (Supplementary Table ST12). Initially, BUSCO v5.6.1 (Manni et al. 2021) was run on each genome assembly using the mammalia_odb10.2024-01-08 database from OrthoDB v.10.1 (Kriventseva et al. 2019). A phylogenetic tree was reconstructed from the BUSCO results using the maximum likelihood method implemented in IQ-TREE v.2.2.0 (Nguyen et al. 2015), with parameters “-m TESTNEW” for model selection and “-bb 1000” to generate 1000 ultra-fast bootstrap replicates. The BuscoClade v1.7 pipeline (https://github.com/tomarovsky/BuscoClade) was used to reconstruct the phylogenetic tree. Multiple whole-genome alignment of all masked genome assemblies (see section “*Pairwise whole-genome alignments and connection between assembly and karyotype*”) was performed using Progressive Cactus v2.8.0 (Armstrong et al. 2020), with the obtained phylogenetic tree and default parameters.

Synteny blocks were extracted from the resulting multiple alignment using halSynteny v2.2 (Krasheninnikova et al. 2020) with parameters “--minBlockSize 50000--maxAnchorDistance 50000”. The results were visualized with the script *draw_macrosynteny.py* from the MACE v1.1.32 software package (https://github.com/mahajrod/MACE), using the parameter “--min_len_threshold 1000000” to highlight only inversions and translocations of 1 Mbp or longer. Centromeres in the analyzed genome assemblies were localized on chromosomal scaffolds by comparison of the corresponding FISH maps and the synteny blocks. The *Canis familiaris* (UU_Cfam_GSD_1.0, GCF_011100685.1) (Hoeppner et al. 2014) and *Homo sapiens* (GRCh38.p14, GCF_000001405.40) (Schneider et al. 2017) genome assemblies were used as references. The detailed algorithm for identification of centromere coordinates is described in (Kliver et al. 2025).

#### Ancestral reconstruction of chromosomal rearrangements

For translocations, we created a two-state, presence (Y) or absence (N), trait matrix with species IDs as columns, and rearrangement IDs as rows. Ancestral reconstruction for the internal nodes of the inferred maximum likelihood tree from the trait matrices was performed using the continuous time Markov model implemented in the ape v5.8 package (function rerootingMethod with model=”SYM”) (Paradis and Schliep 2019). Depending on the predicted probabilities (PN) for “N”, we classified states at internal nodes as “N” (PN >= 0.7), “Y” (PN <= 0.3) and “U” (unclear, 0.3 < PN < 0.7). Our coordinate system was based on the genome assembly of *Mustela strigidorsa* (MSTR), which was chosen because this species occupies a basal position in the phylogenetic tree relative to all other analyzed *Mustela* species. Moreover, it provides the most convenient coordinate framework, as its chromosomes are involved only in fusion events but not in fissions relative to other species. Therefore, for each trait, we checked if the ancestral reconstruction showed “Y” for the root node. In such cases, we inverted the state of the trait (from “Y” to “N” and vice versa) for both internal nodes and leaves.

#### Raw data quality control, filtration and genome size estimation

Barcodes were removed from 10X Genomics linked reads (*M. nivalis* sample 10X_mn) using EMA v0.6.2 (Shajii et al. 2018). Quality control checking of raw and filtered reads was performed using FastQC v0.12.0 [31], KrATER v2.5 (Kliver et al. 2017) and GenomeScope v2.0 (Ranallo-Benavidez et al. 2020). Distributions of 23-mers before and after read filtration were counted using Jellyfish v2.3.0 (Marçais and Kingsford 2011) with the parameters “-m 23-s 30G” for *jellyfish count* and “-l 1-h 100000000” for *jellyfish histo*; and then visualized using KrATER with parameters “-m 23-u 1” to check for possible anomalies and signs of contamination (Supplementary <u>MR</u>). Genome size estimation of the studied species and coverage based on paired-end reads was performed using KrATER, incorporating the GenomeScope-based assessment in diploid mode with a k-mer length of 23. The contamination check of the data (Supplementary <u>MR</u>) was conducted using Kraken v2.1.3 (Wood and Salzberg 2014) and the custom database, which included archaeal, bacterial, viral, human, fungal, plant and protozoan sequences, downloaded from the NCBI RefSeq database on May 10, 2023. In addition to the sequences listed above, the corresponding genome assembly from NCBI RefSeq was added for each species. If such an assembly was missing, an assembly of a closely related species was added (Supplementary <u>MR</u>). The adapter trimming and quality filtering of whole-genome reads were conducted in two stages: an initial k-mer based trimming of large adapter fragments using Cookiecutter (Starostina et al. 2015), followed by an additional trimming of small fragments and quality filtering using Trimmomatic v0.39 (Bolger et al. 2014) with the parameters “ILLUMINACLIP:TruSeq2-PE.fa:2:30:10:1 SLIDINGWINDOW:8:20 MINLEN:50”.

#### Read downsampling and alignments

The coverage of the resequencing data generated for the study varied significantly. To avoid coverage-related bias, we downsampled all the data to ∼12x (desired) coverage, as this coverage was detected for multiple samples (Supplementary <u>MR</u>). The downsampling fraction was calculated as the ratio of the desired coverage to the initial coverage of the sample. If this fraction was >=0.8, no downsampling was performed for that sample. The *reformat.sh* script from the BBmap v38.96 (Bushnell 2014) software package was used to perform random subsampling, specifying the downsampling fraction with the “samplerate=” parameter. The reads were aligned to the corresponding genome assemblies using BWA v0.7.17-r1188 (Li and Durbin 2009) with the default parameters. The Samtools package v1.18 (Li et al. 2009) was used to pair reads, sort them, filter by quality, mark duplicates and index the alignments. The genome coverage in the resulting alignment files was calculated using Mosdepth v0.3.3 (Pedersen and Quinlan 2018) with default parameters.

#### Pseudoautosomal region identification, variant calling and runs of homozygosity

To set the correct ploidy of X chromosome segments during variant calling, we identified coordinates of the pseudoautosomal region (PAR) in samples obtained from males (Supplementary <u>MR</u>) using the *pseudoautosomal_region.py* script from the Biocrutch software package (https://github.com/tomarovsky/Biocrutch). The coverage-based algorithm used for PAR determination is detailed in (Tomarovsky et al. 2025).

SNPs and short indels were identified using Bcftools v1.18 (Li 2011) with the following parameters: “-d 250-q 30-Q 30--adjust-MQ 50-a AD,INFO/AD,ADF,INFO/ADF,ADR,INFO/ADR,DP,SP,SCR,INFO/SCR-O u” for the *bcftools mpileup* and “--ploidy-file--samples-file--group-samples--m-O u-v-f GQ,GP” for the *bcftools call*. The PAR coordinates were specified for variant calling via the “--ploidy-file” parameter. Low-quality genetic variants were filtered out using the *bcftools filter* with the parameters “-S.-O z--exclude’QUAL < 20.0 || (FORMAT/SP > 60.0 | FORMAT/DP < 5.0 | FORMAT/GQ < 20.0)’”. Based on the coverage assessments obtained from the Mosdepth analyses, individual masks for each sample were created using the *generate_mask_from_coverage_bed.py* script from the MAVR v0.97 package with the parameters “-x 2.5-n 0.33”. These masks were used to remove genetic variants if coverage exceeded 250% or was less than 33% of the median genome coverage. The genetic variants were masked using the BEDTools v2.31.0 with the default parameters. The filtered and masked genetic variants for each sample were divided into heterozygous and homozygous single nucleotide polymorphisms (SNPs) and insertions/deletions (indels) using the *bcftools filter* with the parameters “-i’TYPE=“snp”’”, “-i’TYPE=“indel”’”, “-i’FMT/GT=“het”’” and “-i’FMT/GT=“hom”’”. Heterozygous SNPs were counted in 1 Mbp windows and 100 kbp steps.

Runs of homozygosity (RoH) were identified based on the previously calculated heterozygous SNP density in 100 kbp windows with 10 kbp steps. The algorithm is detailed in (Tomarovsky et al. 2025). RoH coordinates for each sample were determined across all autosomes. The visualization of RoH on the chromosomes was performed using the *draw_features.py* script from MACE v1.1.32.

#### Demographic history

Demographic history was reconstructed using the Pairwise Sequentially Markovian Coalescent v0.6.5 (Li and Durbin 2011) software package with the parameters “-N25-t15-r5-b-p ‘4+25*2+4+6’” applied to data sets both including and excluding the sex chromosomes. Genetic variants were identified using an earlier version of Samtools v0.1.19 (compatible with PSMC), with the alignment quality parameter “-C 50” for *samtools mpileup* and the variant calling parameter “-c” for *bcftools view*. Diploid consensus sequences were generated using *vcfutils.pl vcf2fq* from Samtools, specifying the minimum (“-d”) and maximum (“-D”) coverage values, calculated for each sample as follows: the median genome coverage divided by 3 for the “-d” parameter and the median genome coverage multiplied by 2.5 for the “-D” parameter. Coverage values outside these ranges were filtered out. Fasta-like sequences were created using *fq2psmcfa* with a minimum nucleotide quality parameter of “-q20”. Initial bootstrapping, using *splitfa*, divided the sequences into shorter segments, with 100 rounds of bootstrapping specified by the “-b” parameter for each sample. The files were prepared for visualization using *psmc_plot.pl* with “-R” parameter. Starting from values reported for the musteline species included in the IUCN Red List of Threatened Species (Supplementary Table ST1), generation times (“-g”) were slightly adjusted: 3 years were set for the relatively smaller species (*M. nivalis*, *M. erminea*, and *M. richardsonii*) and 4 years for the larger ones (*M. putorius*, *M. sibirica*, *M. strigidorsa*, *M. nigripes*, *M. eversmanii*, and *N. vison*). A mutation rate (“-u”) of 4.64×10⁻⁹ substitutions per generation was applied, with lower (2.94×10⁻⁹) and upper (7.37×10⁻⁹) confidence interval derived from previous estimates of the germline mutation rate in *N. vison* (Bergeron et al. 2023).

#### Phylogenetic analyses

We reconstructed genome assemblies for all the resequenced samples based on the reference genome assemblies of the studied species and SNPs. Indels were not taken into account for this analysis. We used the *FastaAlternateReferenceMaker* tool from the GATK package v4.4.0.0 (Van der Auwera et al. 2013) with the parameter “--use-iupac-sample” to account for heterozygous SNPs. The obtained assemblies and the reference genome assemblies were used to identify single-copy orthologs (6,599) using BUSCO v5.6.1 with the mammalia_odb10.2024-01-08 OrthoDB v.10.1 database. To construct phylogenetic trees, we prepared two datasets: (1) a complete dataset of all resequenced samples and genome assemblies; (2) a dataset excluding all identified hybrids (see “*Admixture analysis of stoat samples*” in Supplementary <u>MR</u>), genome assemblies, and sex chromosomes. Multiple sequence alignment was performed using MAFFT v7.490 (Kuraku et al. 2013), followed by alignment correction by the filtering of hypervariable and poorly aligned regions using Gblocks v0.91b (Castresana 2000). The multiple sequence alignment length after filtering was 11,067,417 bp. Phylogenetic trees were constructed using the Maximum Likelihood method implemented in RAxML-NG v1.2.2 (Kozlov et al. 2024) using the GTGTR4 model and 1000 bootstrap replicates. An alternative phylogenetic reconstruction was performed using ASTRAL-III v5.7.1 (Zhang et al. 2018), based on a set of separately reconstructed gene trees using RAxML-NG. To reduce noise, we applied a filtering step by collapsing nodes with bootstrap support below 70% prior to species tree inference. The resulting phylogenetic trees were visualized using the ETE Toolkit v3.1.2 (Huerta-Cepas et al. 2016). All trees were rooted using the *Martes foina* genome assembly (mfoi.min_150.pseudohap2.1_HiC, DNAZoo). For reproducibility of the analysis, we employed the BuscoClade v1.7 pipeline. The genetic distance matrix was obtained using the Neighbor Joining (NJ) method implemented in RapidNJ v2.3.3 (Simonsen et al. 2008).

#### Mitochondrial genome assemblies and analysis

To reconstruct the mitochondrial DNA phylogenetic tree, we generated mitochondrial genomes for all resequenced samples involved in this study, except for *M. putorius furo*, *N. vison* and *M. nigripes* for which mitochondrial genome were downloaded from NCBI’s Genbank (Supplementary Table ST11). Initially, read subsampling was conducted for each studied sample to 20 Mbp using the Seqtk v1.4 (r122) toolkit (https://github.com/lh3/seqtk). Next, the whole-genome reads of each sample with 20 Mbp were processed using MitoZ v3.6 (Meng et al. 2019) to extract mitochondrial DNA reads, assemble the mitochondrial genomes and annotate the resulting assemblies. MitoZ was run in the “all” mode with the following parameters: “--clade Chordata--genetic_code 2--assembler megahit--requiring_taxa Chordata”. Following the assembly of mitochondrial genomes, all publicly available assemblies from the NCBI database were included in the analysis with the query ““Mustelinae”[Orgn] AND (mitochondrial OR mitochondrion OR mitochondrion complete genome) AND 10000:20000[SLEN]”. These data were manually checked for correctness. Sequence orientation for each sample was checked and corrected using Unipro UGENE v50.0 (Okonechnikov et al. 2012). Multiple sequence alignment was performed using MAFFT v7.490, followed by the filtering of hypervariable and poorly aligned regions using trimAl v1.4.1 (Capella-Gutiérrez et al. 2009) with the parameters “-automated1-nogaps”. The multiple sequence alignment length prior to and after filtering was 18,050 bp and 15,793 bp, respectively. A phylogenetic tree was reconstructed using the maximum likelihood method with IQ-TREE v.2.2.0 with automatic selection of the best-fitting substitution model using ModelFinder (Kalyaanamoorthy et al. 2017). The stability of the tree was evaluated using 1000 ultra-fast bootstrap replicates. To root the tree we used *Martes foina* (NC_020643.1) as an outgroup. The resulting phylogenetic tree was visualized using the ETE Toolkit v3.1.2.

## Supporting information

Supplementary Methods and Results

## Acknowledgements

The unpublished genome assembly of the domestic ferret (*Mustela putorius furo*), Hi-C (SRR34091397) and Illumina libraries (SRR34068022) of *M. strigidorsa*, as well as Hi-C (SRR34082581) library of *M. nivalis*, were used by permission from the DNA Zoo Consortium (https://www.dnazoo.org). We acknowledge Valeriy A. Zelepukhin for helping with sample gathering. The research was completed using equipment (materials) of the large-scale research facilities “Cryobank of cell cultures” Institute of Molecular and Cellular Biology SB RAS (Novosibirsk, Russia). This research was supported in part through computational resources of HPC facilities at the collaborative center «Bioinformatics» ICG SB RAS, as well as the computing cluster of ITMO University.

## Data availability

Genome assemblies and resequencing data are available in BioProject PRJNA1146985. Mitochondrial data obtained during the study are available at the following accession numbers: PQ246107-PQ246113, PQ821906. Mitochondrial data derived from publicly available resequencing data are available in the GenBank Third Party Annotation database under the following accession numbers: BK068807, BK068808, BK069841-BK069851.

## Supplementary

Supplementary Methods and Results: <u>link</u>

Supplementary Figures: <u>link</u>

Supplementary Tables: <u>link</u>

Supplementary File: <u>link</u>

