## Supplementary Methods and Results for "Comparative genomics and phylogenomics of the Mustelinae lineage (Mustelidae, Carnivora)"

[11] Departamento de Biologia, Faculdade de Ciências da Universidade do Porto, Rua Campo Alegre s/n, 4169-007 Porto, Portugal.

[12] BIOPOLIS Program in Genomics, Biodiversity and Land Planning, CIBIO, 4485-661 Vairão, Portugal.

[18] Centre for Computational Biology, Peter the Great Saint Petersburg Polytechnic University, 29 Polytechnicheskaya str., St. Petersburg, 195251, Russia. (https://orcid.org/0000-0003-1416-0200).

[21] Laboratory of Amyloid Biology, St. Petersburg State University, 199034 St. Petersburg, Russia.

[23] Center for Evolutionary Hologenomics, The Globe Institute, The University of Copenhagen, 5A Oester Farimagsgade, Copenhagen, 1353, Denmark. (https://orcid.org/0000-0002-2965-3617).

* corresponding author

### Supplementary Methods

#### *Admixture analysis of stoat samples*

For the analysis of population structure and admixture in *M. erminea* and *M. richardsonii* samples, we used the filtered and masked SNP data located on autosomes and in the PAR. A final mask was created based on previously obtained individual coverage masks for each sample. This mask was developed through a two-step procedure: first, using BEDOPS v2.4.40 (Neph et al. 2012), we intersected individual masks for each pair of samples and then combined all intersections. The resulting masking track contained sites with excessively high (>250%) or excessively low (<33%) coverage in at least two samples. Filtered and masked SNP data were further filtered using PLINK v1.9 (Purcell et al. 2007), excluding variants with a 100% genotyping rate across all samples ("--geno 0"). We also pruned the SNPs with high pairwise linkage disequilibrium (LD) in sliding windows of 50 SNPs with a step size of 10 SNPs, using an $r^2$ threshold of 0.7 ("--indep-pairwise 50 10 0.7").

Principal component analysis (PCA) was performed using PLINK v1.9 (option "--pca"). Genome-wide admixture (global admixture) was estimated with ADMIXTURE v1.3.0 (Alexander et al. 2009) for K (number of populations) values ranging from 2 to 5, with each K value analyzed in three replicates. The results were visualized using pong v1.5 (Behr et al. 2016). For K=2, we performed local ancestry analysis (local admixture) to identify admixed ancestry regions. The *M. erminea* reference genome was split into sliding windows of 1 Mbp with a step size of 100 kbp and admixture analysis was performed independently for each window.

### Supplementary Results

*Coverage and contamination check*

Quality assessment of the whole-genome reads based on 23-mer distribution revealed significant variation in coverage across the samples, ranging from 9.82x to 117.82x (Figure 1).

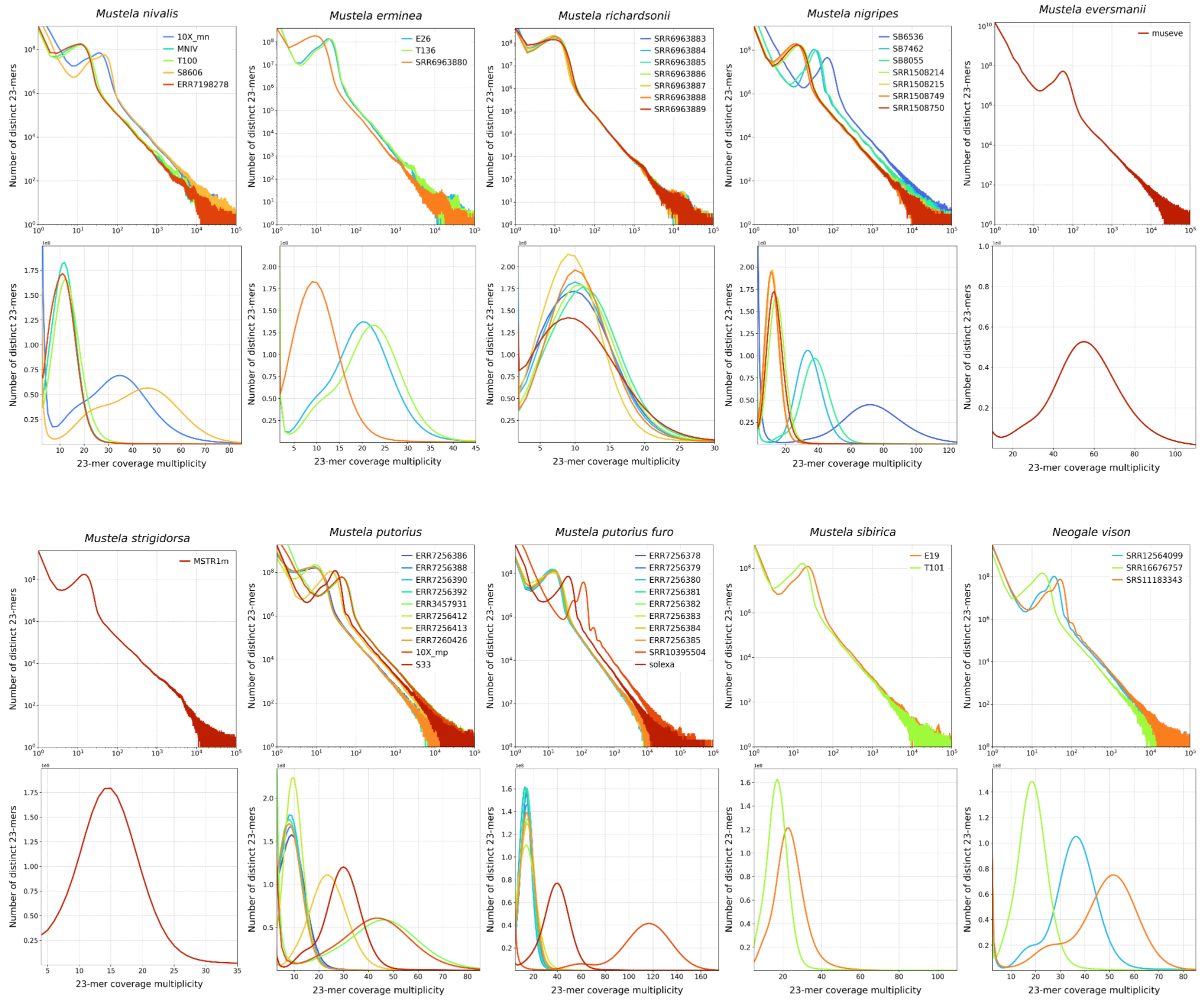


**Figure 1.** Distribution of 23-mers for samples of each studied species in logarithmic (top graph) and linear (bottom graph) scales.

Due to this coverage variation, all samples were downsampled to approximately 12x coverage (Table 1).

**Table 1.** Whole genome coverage statistics.

| Species | Sample | Whole genome coverage statistics | | | |
|---|---|---|---|---|---|
| | | median | mean | max | min |
| *M. nivalis* | MNIV | 14.0 | 17.59 | 139235.0 | 0.0 |
| | T100 | 12.0 | 17.14 | 134598.0 | 0.0 |
| | 10X_mn | 13.0 | 17.07 | 148809.0 | 0.0 |
| | ERR7198278 | 15.0 | 18.52 | 85657.0 | 0.0 |
| | S8606 | 11.0 | 15.31 | 94073.0 | 0.0 |
| *M. strigidorsa* | MSTR1m | 14.0 | 18.93 | 126028.0 | 0.0 |
| *M. sibirica* | E19 | 13.0 | 13.89 | 78173.0 | 0.0 |
| | T101 | 13.0 | 13.76 | 97072.0 | 0.0 |
| *M. erminea* | T136 | 13.0 | 15.16 | 131335.0 | 0.0 |
| | E26 | 13.0 | 15.11 | 111787.0 | 0.0 |
| | SRR6963880 | 13.0 | 14.27 | 48319.0 | 0.0 |
| *M. richardsonii* | SRR6963883 | 14.0 | 15.94 | 46069.0 | 0.0 |
| | SRR6963884 | 14.0 | 15.95 | 42918.0 | 0.0 |
| | SRR6963885 | 15.0 | 17.15 | 54270.0 | 0.0 |
| | SRR6963886 | 14.0 | 15.82 | 44997.0 | 0.0 |
| | SRR6963887 | 13.0 | 14.64 | 62647.0 | 0.0 |
| | SRR6963888 | 13.0 | 14.75 | 63303.0 | 0.0 |
| | SRR6963889 | 11.0 | 12.49 | 43735.0 | 0.0 |
| *M. nigripes* | SRR1508214 | 13.0 | 14.89 | 40896.0 | 0.0 |
| | SRR1508215 | 14.0 | 16.07 | 48044.0 | 0.0 |

| | | | | | |
|---|---|---|---|---|---|
| | SRR1508750 | 15.0 | 17.0 | 35137.0 | 0.0 |
| | SRR1508749 | 14.0 | 15.13 | 38700.0 | 0.0 |
| | SB7462 | 13.0 | 15.24 | 62969.0 | 0.0 |
| | SB8055 | 13.0 | 14.53 | 48176.0 | 0.0 |
| | SB6536 | 13.0 | 15.4 | 76727.0 | 0.0 |
| *M. putorius* | ERR3457930 | 12.0 | 13.32 | 33825.0 | 0.0 |
| | ERR7256386 | 8.0 | 9.3 | 136014.0 | 0.0 |
| | ERR7256388 | 9.0 | 9.45 | 41122.0 | 0.0 |
| | ERR7256390 | 9.0 | 9.37 | 30000.0 | 0.0 |
| | ERR7256392 | 8.0 | 9.14 | 77004.0 | 0.0 |
| | ERR7256412 | 12.0 | 12.6 | 70638.0 | 0.0 |
| | ERR7256413 | 12.0 | 13.06 | 34538.0 | 0.0 |
| | ERR7260426 | 8.0 | 8.9 | 34270.0 | 0.0 |
| | S33 | 13.0 | 14.3 | 41972.0 | 0.0 |
| *M. putorius furo* | Solexa | 11.0 | 11.76 | 65490.0 | 0.0 |
| | SRR10395504 | 13.0 | 14.32 | 219675.0 | 0.0 |
| | ERR7256378 | 13.0 | 14.3 | 201239.0 | 0.0 |
| | ERR7256379 | 13.0 | 13.74 | 216036.0 | 0.0 |
| | ERR7256380 | 16.0 | 16.91 | 242836.0 | 0.0 |
| | ERR7256381 | 13.0 | 14.22 | 196452.0 | 0.0 |
| | ERR7256382 | 12.0 | 12.85 | 168507.0 | 0.0 |
| | ERR7256383 | 11.0 | 11.81 | 162446.0 | 0.0 |

| | | | | | |
|---|---|---|---|---|---|
| | ERR7256384 | 12.0 | 12.88 | 188682.0 | 0.0 |
| | ERR7256385 | 12.0 | 12.63 | 152067.0 | 0.0 |
| *M. eversmanni* | ERR11751895 | 11.0 | 12.61 | 449035.0 | 0.0 |
| | ERR7198276 | 8.0 | 09.07 | 26430.0 | 0.0 |
| | ERR7198277 | 9.0 | 10.49 | 861008.0 | 0.0 |
| *N. vison* | SRR12564099 | 13.0 | 13.28 | 428894.0 | 0.0 |
| | SRR16676757 | 12.0 | 12.72 | 176772.0 | 0.0 |
| | SRS11183343 | 14.0 | 13.84 | 28599.0 | 0.0 |

Additionally, several samples showed extra peaks at the beginning of the 23-mer distribution (Figure 1). As these peaks could be indicative of both high levels of heterozygosity and potential contamination, we classified the reads for each sample (Figure 2).

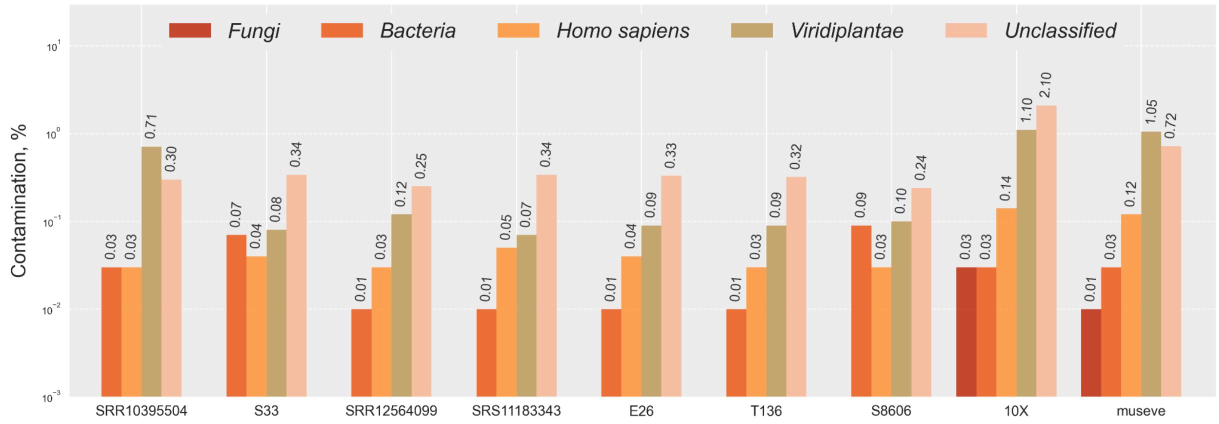


**Figure 2.** Checking the data for contamination. The samples showing additional peaks on the 23-mer distribution are presented.

Contamination analysis did not reveal any significant deviations, with the percentage of target species content (Table 2) across all samples ranging from 95.31% to 99.68% (Table 3).

**Table 2.** Genome assemblies used in the classification of reads.

| Target species | Reference species | Reference assembly name/ID |
|---|---|---|
| *M. nivalis* | *M. nivalis* | GCA_019141155.1 |
| *M. strigidorsa* | *M. erminea* | GCF_009829155.1 |
| *M. erminea* | *M. erminea* | GCF_009829155.1 |
| *M. sibirica* | *M. erminea* | GCF_009829155.1 |
| *M. eversmanii* | *M. eversmanii* | GCA_963422785.1 |
| *M. nigripes* | *M. nigripes* | GCF_022355385.1 |
| *M. putorius* | *M. putorius* | GCA_902207235.1 |
| *M. putorius furo* | *M. putorius furo* | GCA_011764305.2 |
| *N. vison* | *N. vison* | GCF_020171115.1 |

**Table 3.** Classification of reads.

| Species | Sample ID | Number of read pairs | Target species | Unclassified | *Homo sapiens* | Bacteria | Viridiplantae | Fungi | SAR | Viruses | Archaea |
|---|---|---|---|---|---|---|---|---|---|---|---|
| *M. nivalis* | 10X_mn | 618349704 | 96.35 | 2.1 | 0.14 | 0.03 | 1.1 | 0.03 | 0 | 0 | 0 |
| | ERR7198278 | 193408869 | 99.13 | 0.26 | 0.3 | 0.2 | 0.07 | 0 | 0 | 0 | 0 |
| | MNIV | 162669637 | 99.58 | 0.22 | 0.04 | 0.01 | 0.11 | 0 | 0 | 0 | 0 |
| | S8606 | 651649526 | 99.5 | 0.24 | 0.03 | 0.09 | 0.1 | 0 | 0 | 0 | 0 |
| | T100 | 190641593 | 99.58 | 0.24 | 0.04 | 0.01 | 0.1 | 0 | 0 | 0 | 0 |
| *M. strigidorsa* | MSTR1m | 214749346 | 95.31 | 4.00 | 0.08 | 0.11 | 0.36 | 0 | 0 | 0 | 0 |
| *M. sibirica* | E19 | 251146712 | 98.84 | 0.74 | 0.11 | 0.04 | 0.18 | 0 | 0 | 0 | 0 |
| | T101 | 183857602 | 99.01 | 0.66 | 0.09 | 0.02 | 0.16 | 0 | 0 | 0 | 0 |
| *M. erminea* | E26 | 238732014 | 99.49 | 0.33 | 0.04 | 0.01 | 0.09 | 0 | 0 | 0 | 0 |
| | T136 | 264805617 | 99.5 | 0.32 | 0.03 | 0.01 | 0.09 | 0 | 0 | 0 | 0 |
| | SRR6963880 | 201919606 | 99.29 | 0.56 | 0.02 | 0.01 | 0.08 | 0 | 0 | 0 | 0 |

| | | | | | | | | | | | |
|---|---|---|---|---|---|---|---|---|---|---|---|
| *M. richardsonii* | SRR6963883 | 215858519 | 99.19 | 0.61 | 0.03 | 0.01 | 0.11 | 0 | 0 | 0 | 0 |
| | SRR6963884 | 216618365 | 99.27 | 0.53 | 0.03 | 0.01 | 0.11 | 0 | 0 | 0 | 0 |
| | SRR6963885 | 236185815 | 99.27 | 0.48 | 0.05 | 0.03 | 0.13 | 0 | 0 | 0 | 0 |
| | SRR6963886 | 221551197 | 99.34 | 0.49 | 0.03 | 0.01 | 0.09 | 0 | 0 | 0 | 0 |
| | SRR6963887 | 194715867 | 99.31 | 0.5 | 0.03 | 0.01 | 0.11 | 0 | 0 | 0 | 0 |
| | SRR6963888 | 205827781 | 99.24 | 0.57 | 0.03 | 0.01 | 0.1 | 0 | 0 | 0 | 0 |
| | SRR6963889 | 204272335 | 99.41 | 0.47 | 0.03 | 0 | 0.05 | 0 | 0 | 0 | 0 |
| *M. nigripes* | SB6536 | 1052588227 | 98.51 | 0.75 | 0.07 | 0.02 | 0.5 | 0.01 | 0 | 0 | 0 |
| | SB7462 | 401310398 | 99.51 | 0.31 | 0.04 | 0.01 | 0.09 | 0 | 0 | 0 | 0 |
| | SB8055 | 421914722 | 99.5 | 0.29 | 0.05 | 0.01 | 0.11 | 0 | 0 | 0 | 0 |
| | SRR1508214 | 270590776 | 99.12 | 0.5 | 0.05 | 0.02 | 0.19 | 0 | 0 | 0 | 0 |
| | SRR1508215 | 225975899 | 99.08 | 0.5 | 0.05 | 0.02 | 0.18 | 0 | 0 | 0 | 0 |
| | SRR1508749 | 214227048 | 99.11 | 0.49 | 0.05 | 0.02 | 0.18 | 0 | 0 | 0 | 0 |
| | SRR1508750 | 244374194 | 99.13 | 0.49 | 0.05 | 0.02 | 0.19 | 0 | 0 | 0 | 0 |

| | | | | | | | | | | | |
|---|---|---|---|---|---|---|---|---|---|---|---|
| *M. putorius* | S33 | 326257265 | 99.44 | 0.34 | 0.04 | 0.07 | 0.08 | 0 | 0 | 0 | 0 |
| | ERR3457930 | 316464462 | 99.34 | 0.42 | 0.08 | 0.03 | 0.1 | 0 | 0 | 0 | 0 |
| | ERR7256386 | 64944975 | 97.74 | 0.43 | 0.03 | 1.38 | 0.08 | 0 | 0 | 0.01 | 0 |
| | ERR7256388 | 61723609 | 99.31 | 0.22 | 0.04 | 0.05 | 0.05 | 0 | 0 | 0 | 0 |
| | ERR7256390 | 61081647 | 99.16 | 0.31 | 0.04 | 0.08 | 0.06 | 0 | 0 | 0 | 0 |
| | ERR7256392 | 62695929 | 99.17 | 0.23 | 0.04 | 0.1 | 0.05 | 0 | 0 | 0 | 0 |
| | ERR7256412 | 135051845 | 98.67 | 0.83 | 0.03 | 0.06 | 0.15 | 0 | 0.01 | 0 | 0 |
| | ERR7256413 | 152916463 | 99.53 | 0.29 | 0.07 | 0.02 | 0.07 | 0 | 0 | 0 | 0 |
| | ERR7260426 | 60404590 | 99.25 | 0.23 | 0.04 | 0.06 | 0.05 | 0 | 0 | 0 | 0 |
| *M. putorius furo* | ERR7256378 | 200811196 | 98.27 | 0.28 | 0.03 | 0.12 | 0.05 | 0 | 0 | 0 | 0 |
| | ERR7256379 | 198793435 | 98.31 | 0.28 | 0.03 | 0.11 | 0.05 | 0 | 0 | 0 | 0 |
| | ERR7256380 | 181557812 | 98.07 | 0.28 | 0.03 | 0.13 | 0.05 | 0 | 0 | 0 | 0 |
| | ERR7256381 | 196895149 | 98.31 | 0.28 | 0.03 | 0.11 | 0.06 | 0 | 0 | 0 | 0 |
| | ERR7256382 | 205707310 | 98.06 | 0.35 | 0.03 | 0.2 | 0.08 | 0 | 0 | 0 | 0 |

| | | | | | | | | | | | |
|---|---|---|---|---|---|---|---|---|---|---|---|
| | ERR7256383 | 207090454 | 98.39 | 0.31 | 0.03 | 0.12 | 0.07 | 0 | 0 | 0 | 0 |
| | ERR7256384 | 204561178 | 98.21 | 0.33 | 0.03 | 0.16 | 0.08 | 0 | 0 | 0 | 0 |
| | ERR7256385 | 204201493 | 98.14 | 0.31 | 0.03 | 0.14 | 0.09 | 0 | 0 | 0 | 0 |
| | SRR10395504 | 1268568171 | 98.41 | 0.3 | 0.03 | 0.03 | 0.71 | 0 | 0 | 0 | 0 |
| | Solexa | 704126807 | 98.02 | 01.04 | 0.04 | 0.01 | 0.2 | 0 | 0 | 0 | 0 |
| *M. eversmanii* | ERR11751895 | 443701651 | 97.98 | 0.72 | 0.12 | 0.03 | 01.05 | 0.01 | 0 | 0 | 0 |
| | ERR7198276 | 61722595 | 99.03 | 0.34 | 0.04 | 0.07 | 0.08 | 0.01 | 0 | 0 | 0 |
| | ERR7198277 | 121381979 | 97.78 | 1.56 | 0.05 | 0.12 | 0.22 | 0.01 | 0 | 0 | 0 |
| *N. vison* | SRR12564099 | 395644985 | 99.55 | 0.25 | 0.03 | 0.01 | 0.12 | 0 | 0 | 0 | 0 |
| | SRR16676757 | 203489410 | 99.68 | 0.2 | 0.01 | 0.01 | 0.06 | 0 | 0 | 0 | 0 |
| | SRS11183343 | 857282157 | 99.48 | 0.34 | 0.05 | 0.01 | 0.07 | 0 | 0 | 0 | 0 |

*Pseudoautosomal region*

We have identified the coordinates of the PAR (PseudoAutosomal Region) on the sex chromosomes of males. The maximum length of the PAR varied between the species from 6.1 for *M. eversmanii* to 6.6 Mbp for *M. erminea*, *M. putorius* and *N. vison*. The rest of the species had intermediate values: 6.2 Mbp (*M. richardsonii*), 6.4 Mbp (*M. nivalis*, *M. strigidorsa* and *M. sibirica*) and 6.5 Mbp (*M. putorius furo* and *M. nigripes)* (Table 4).

**Table 4**. PAR coordinates of male sex chromosomes.

| **Species** | **Sample** | **PAR coordinates** | | | **Sex X-chromosome ID** | **Sex X-chromosome length, bp.** |
|---|---|---|---|---|---|---|
| | | **start** | **stop** | **length** | | |
| *M. nivalis* | 10X_mn | 118960000 | 125400000 | 6440000 | HiC_scaffold_21 | 125416964 |
| | MNIV | 118960000 | 125400000 | 6440000 | | |
| | S8606 | 118960000 | 125400000 | 6440000 | | |
| | ERR7198278 | 119160000 | 124480000 | 5320000 | | |
| *M. strigidorsa* | MSTR1m | 0 | 6390000 | 6390000 | HiC_scaffold_23 | 73834643 |
| *M. erminea* | SRR6963880 | 980000 | 6060000 | 5080000 | NC_045635.1 | 130149454 |
| | E26 | 0 | 6600000 | 6600000 | | |
| | T136 | 0 | 6600000 | 6600000 | | |
| *M. richardsonii* | SRR6963883 | 690000 | 6540000 | 5850000 | NC_045635.1 | 130149454 |
| | SRR6963885 | 430000 | 6590000 | 6160000 | | |

| Species | Sample | PAR coordinates | | | Sex X-chromosome ID | Sex X-chromosome length, bp. |
|---|---|---|---|---|---|---|
| | | start | stop | length | | |
| *M. putorius* | ERR3457930 | 180000 | 6570000 | 6390000 | HiC_scaffold_10_RagTag | 126329207 |
| | ERR7256386 | 1000000 | 6500000 | 5500000 | | |
| | ERR7256388 | 1000000 | 6020000 | 5020000 | | |
| | ERR7256392 | 980000 | 6510000 | 5530000 | | |
| | ERR7256413 | 220000 | 6570000 | 6350000 | | |
| | ERR7260426 | 1030000 | 6020000 | 4990000 | | |
| | S33 | 0 | 6570000 | 6570000 | | |
| *M. putorius furo* | ERR7256379 | 670000 | 6510000 | 5840000 | HiC_scaffold_10 | 123993920 |
| | ERR7256382 | 1000000 | 6280000 | 5280000 | | |
| | ERR7256383 | 1000000 | 5710000 | 4710000 | | |
| | SRR10395504 | 0 | 6510000 | 6510000 | | |
| *M. nigripes* | SB6536 | 117770000 | 124120000 | 6350000 | NC_081575.1 | 124252308 |
| | SB8055 | 117770000 | 124240000 | 6470000 | | |
| | SRR1508214 | 117770000 | 124130000 | 6360000 | | |
| | SRR1508215 | 117770000 | 124240000 | 6470000 | | |
| | SRR1508749 | 117770000 | 124200000 | 6430000 | | |
| *M. eversmanii* | ERR11751895 | 123990000 | 130080000 | 6090000 | HiC_scaffold_10_Rag | 130183722 |
| | ERR7198276 | 124210000 | 129230000 | 5020000 | | |
| | ERR7198277 | 123990000 | 130080000 | 6090000 | | |

| Species | Sample | PAR coordinates | | | Sex X-chromosome ID | Sex X-chromosome length, bp. |
|---|---|---|---|---|---|---|
| | | start | stop | length | | |
| | | | | | Tag | |
| *M. sibirica* | E19 | 0 | 6380000 | 6380000 | NC_045635.1_RagTag | 112012753 |
| | T101 | 0 | 6380000 | 6380000 | | |
| *N. vison* | SRR12564099 | 125120000 | 131670000 | 6550000 | NC_058105.1 | 131682864 |
| | SRR16676757 | 125120000 | 131670000 | 6550000 | | |
| | SRS11183343 | 125120000 | 131560000 | 6440000 | | |

The correct localization of the pseudoautosomal region (PAR) of sex chromosomes is crucial for the accurate detection of genetic variants in heterogametic individuals. However, precise identification of the PAR borders from a low coverage WGS data (our case) is challenging. For our samples we observed variations in its size both within the same species and across different species (Table 4). However, these variations may not necessarily reflect biological differences but may be technical artifacts. This is due to several factors: (1) PAR is located at the ends of sex chromosomes, making it difficult to accurately define its borders since repeats at the ends of chromosomes can reduce coverage; (2) fluctuations of coverage in a low coverage sample may bias a result; (3) a X chromosome in an assembly might be of lower quality than autosomes if it was generated from a male; and (4) structural rearrangements, such as segmental duplications, within chrX may bias coverage estimates via both assembly and alignment artifacts. Together, these factors result in an uneven coverage, complicating the precise determination of PAR borders using coverage-based algorithms.

Given these limitations, we chose to use the largest PAR size among all samples as a reference for each species and applied this information for identifying genetic variants. Despite these assumptions, our results align with previously estimated PAR lengths for members of the Mustelidae family and other lineages within the Carnivorans, obtained from high-coverage whole-genome data (Totikov et al. 2021).

*Comparison of M. nivalis genome assemblies*

During manuscript preparation for submission, one more chromosome-level genome assembly of the *M. nivalis* (mMusNiv2.hap1.1, GCA_964662115.1) was released to the scientific community. Compared with our assembly, it is longer (3.4 Gbp vs 2.45 Gbp), but its N50 scaffold is slightly lower (115.39 Mbp vs 138.37 Mbp). In terms of completeness, mMusNiv2.hap1.1 contains 8808 (95.5%) complete BUSCOs (93.7% single-copy and 1.8% duplicated), which is lower than in our assembly (96%). The fraction of fragmented (1.4%) and missing (3.1%) BUSCOs is also higher compared to our assembly (1.1% fragmented and 2.9% missing).

*Principal component analysis and admixture of Mustela erminea and Mustela richardsonii*

The analysis of heterozygosity levels in *M. richardsonii* samples revealed relatively high mean values, ranging from 1.28 SNPs/kbp in sample SRR6963888 to 2.94 SNPs/kbp in sample SRR6963887 (Figure 3A). The high heterozygosity level in sample SRR6963887 may indicate admixture, as noted in a previous study (Colella et al. 2018).

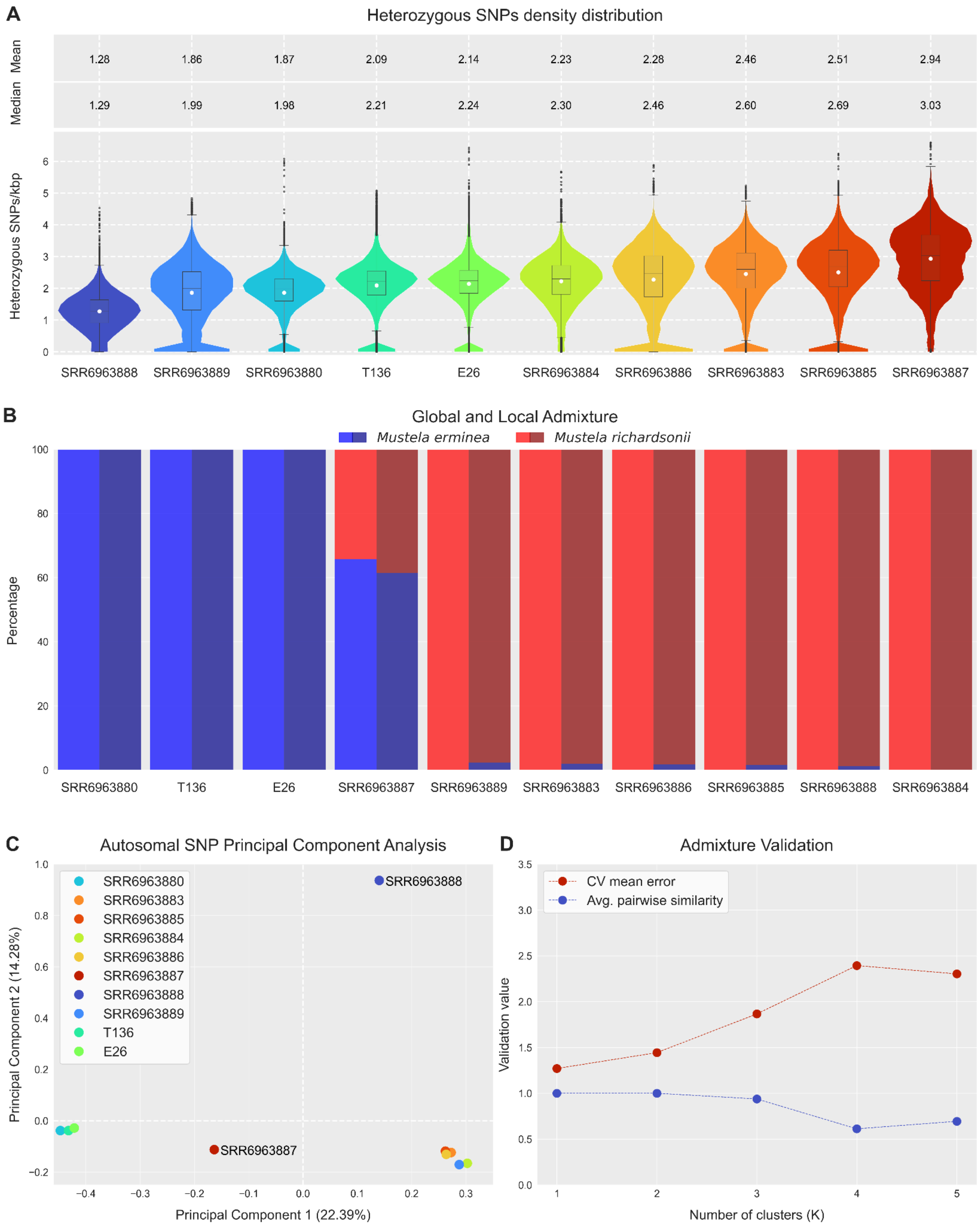


**Figure 3**. Global and Local Admixture, Heterozygosity, PCA and Admixture Validation of the *Mustela erminea* and *Mustela richardsonii* samples.

(A) Distribution of the *Mustela erminea* and *Mustela richardsonii* heterozygous SNP density counted in 1 Mbp sliding windows and 100 kbp steps and converted to heterozygous SNPs/kbp; (B) Global Admixture (left half – blue, red) and Local Admixture (right half – dark blue, dark red) of 1 Mbp sliding windows with 100 kbp step based on *M. erminea* genome assembly. (C) Principal component analysis (PCA) based on heterozygous SNPs from autosomes and pseudoautosomal region (PAR) of *M. erminea* genome assembly; (D) Plot of mean values of cross-validation error (red) and values of average pairwise similarity (blue) based on Global Admixture of *M. erminea* genome assembly.

Compared with that study we incorporated two newly sequenced Asian *M. erminea* samples, and performed both global and local ADMIXTURE analysis (Figure 3B, Table 5). Cross-validation error estimates for the global analysis support two clusters (K=2) (Figure 3D), indicating admixture in sample SRR6963887 (Canada, Southern Yukon Territory). The level of admixture is 34.3%, while no signs of admixture were detected in the other samples according to the global Admixture results. Local Admixture analysis identified admixture of more than 1% in five additional samples: SRR6963889 (2.31%), SRR6963883 (1.95%), SRR6963886 (1.77%), SRR6963885 (1.57%) and SRR6963888 (1.16%) (Table 5). Principal Component Analysis (PCA) revealed two distinct clusters as well (Figure 3C): *M. erminea* (left) and *M. richardsonii* (right), with samples SRR6963887 and SRR6963888 positioned between them. Sample SRR6963887 is closer to the *M. erminea* cluster along the horizontal axis PC1 (22.39%), while sample SRR6963888 is closer to the *M. richardsonii* cluster along the vertical axis PC2 (14.28%).

**Table 5**. Global and Local Admixture (K=2).

| **Sample** | **Global Admixture** | | **Local Admixture** | |
|---|---|---|---|---|
| | ***M. erminea*, %** | ***M. richardsonii*, %** | ***M. erminea*, %** | ***M. richardsonii*, %** |
| E26 | 100 | 0 | 99.94 | 0.060 |
| T136 | 100 | 0 | 99.98 | 0.017 |
| SRR6963880 | 100 | 0 | 99.94 | 0.057 |
| SRR6963883 | 0 | 100 | 1.955 | 98.04 |

| Sample | Global Admixture | | Local Admixture | |
|---|---|---|---|---|
| | ***M. erminea*, %** | ***M. richardsonii*, %** | ***M. erminea*, %** | ***M. richardsonii*, %** |
| SRR6963884 | 0 | 100 | 0.062 | 99.94 |
| SRR6963885 | 0 | 100 | 1.566 | 98.43 |
| SRR6963886 | 0 | 100 | 1.769 | 98.23 |
| SRR6963887 | 65.7 | 34.3 | 61.38 | 38.62 |
| SRR6963888 | 0 | 100 | 1.156 | 98.84 |
| SRR6963889 | 0 | 100 | 2.308 | 97.69 |
