## Supplementary Figures for "Comparative genomics and phylogenomics of the Mustelinae lineage (Mustelidae, Carnivora)"

[11] Departamento de Biologia, Faculdade de Ciências da Universidade do Porto, Rua Campo Alegre s/n, 4169-007 Porto, Portugal.

[12] BIOPOLIS Program in Genomics, Biodiversity and Land Planning, CIBIO, 4485-661 Vairão, Portugal.

[18] Centre for Computational Biology, Peter the Great Saint Petersburg Polytechnic University, 29 Polytechnicheskaya str., St. Petersburg, 195251, Russia. (https://orcid.org/0000-0003-1416-0200).

[21] Laboratory of Amyloid Biology, St. Petersburg State University, 199034 St. Petersburg, Russia.

[23] Center for Evolutionary Hologenomics, The Globe Institute, The University of Copenhagen, 5A Oester Farimagsgade, Copenhagen, 1353, Denmark. (https://orcid.org/0000-0002-2965-3617).

* corresponding author

**SF1 (Supplementary Figure 1).** Hi-C contact maps of newly generated genome assemblies of two *Mustela* species and their synteny.

(A) Hi-C density map of the *M. nivalis* genome assembly with 21 chromosomes. (B) Hi-C density map of the *M. strigidorsa* genome assembly with 22 chromosomes. (C) Macro-level synteny (gray lines) among assemblies of *M. strigidorsa*, *M. erminea* and *M. nivalis*. Inversions (larger than 1 Mbp) and fusion/fission events are highlighted in red and blue, respectively. Chromosomes labeled by primes (‘) were reverse complemented to follow the orientation in the *M. erminea* assembly. Numbering of the *M. nivalis* chromosomes was done according to the published cytogenetic data (Graphodatsky et al. 2020). For *M. strigidorsa*, chromosome numbering was transferred from *M. erminea* due to one-to-one synteny. Centromeric positions are tentative, approximately drawn based on the analysis of comparative chromosome painting maps and G-banded karyotypes.

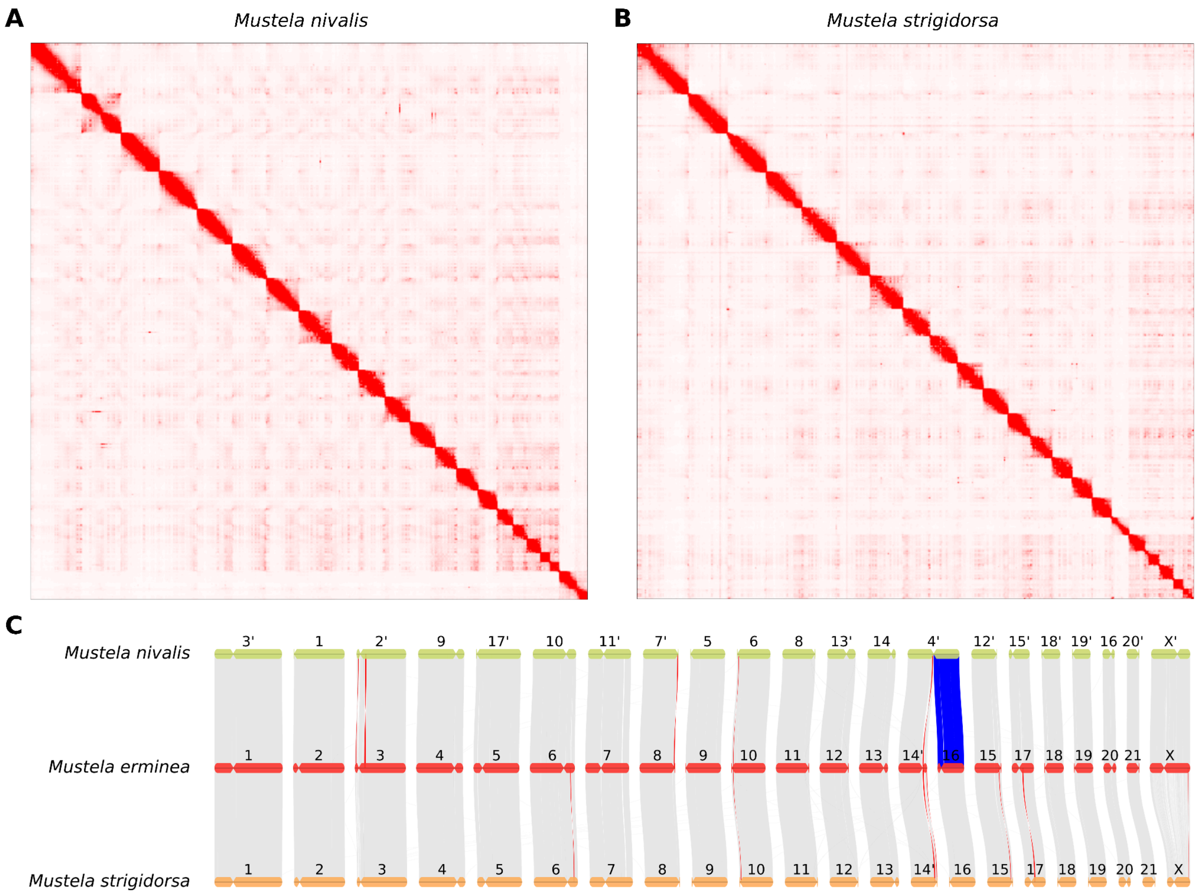


**SF2 (Supplementary Figure 2).** Quality assessment of genome assemblies by conserved orthologous groups (BUSCOs) from three OrthoDB v10.1 datasets: Mammalia (A), Laurasiatheria (B), and Carnivora (C). Abbreviations: Chr – chromosome-level genome assembly, pChr – pseudochromosome-level genome assembly.

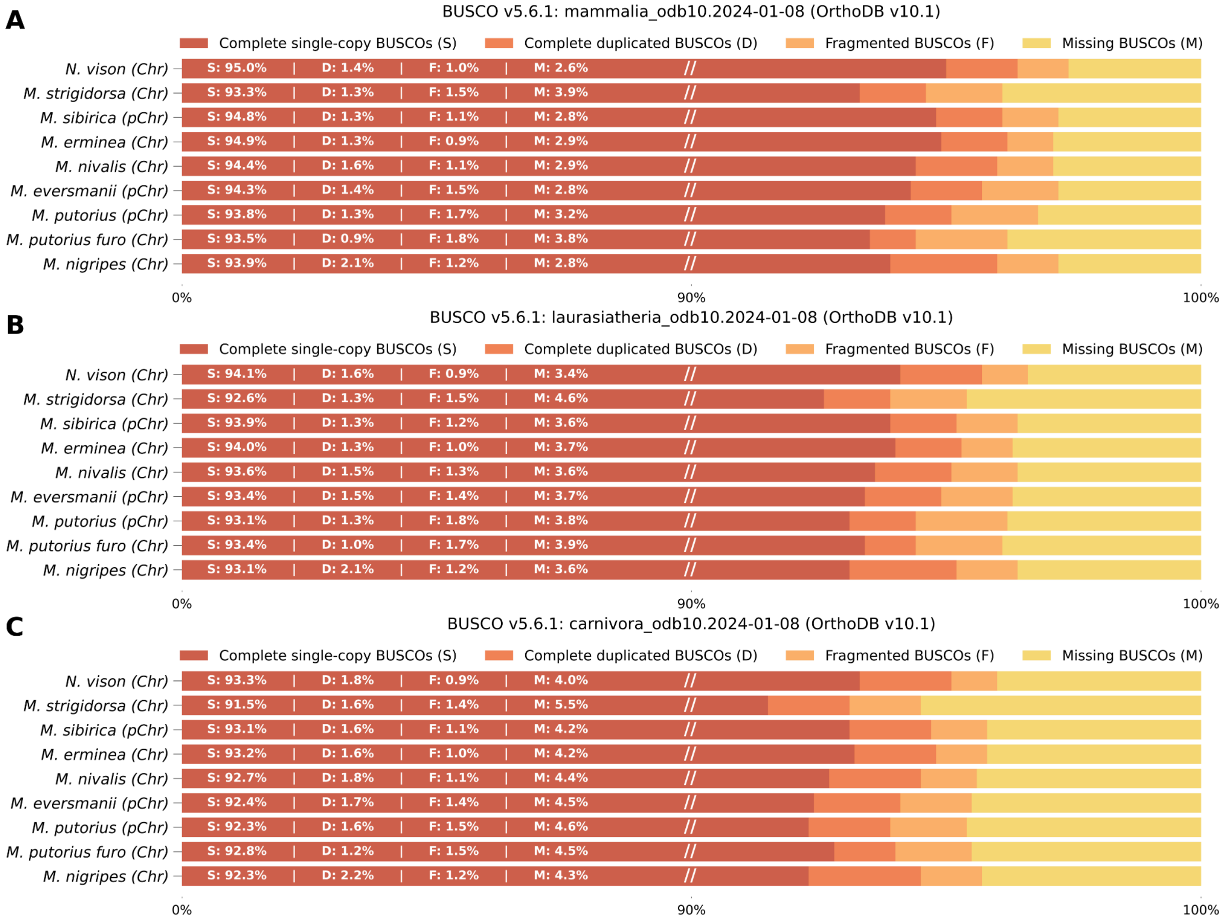


**SF3 (Supplementary Figure 3).** Kimura distance-based copy divergence analyses of Carnivora transposable elements in genome assemblies.

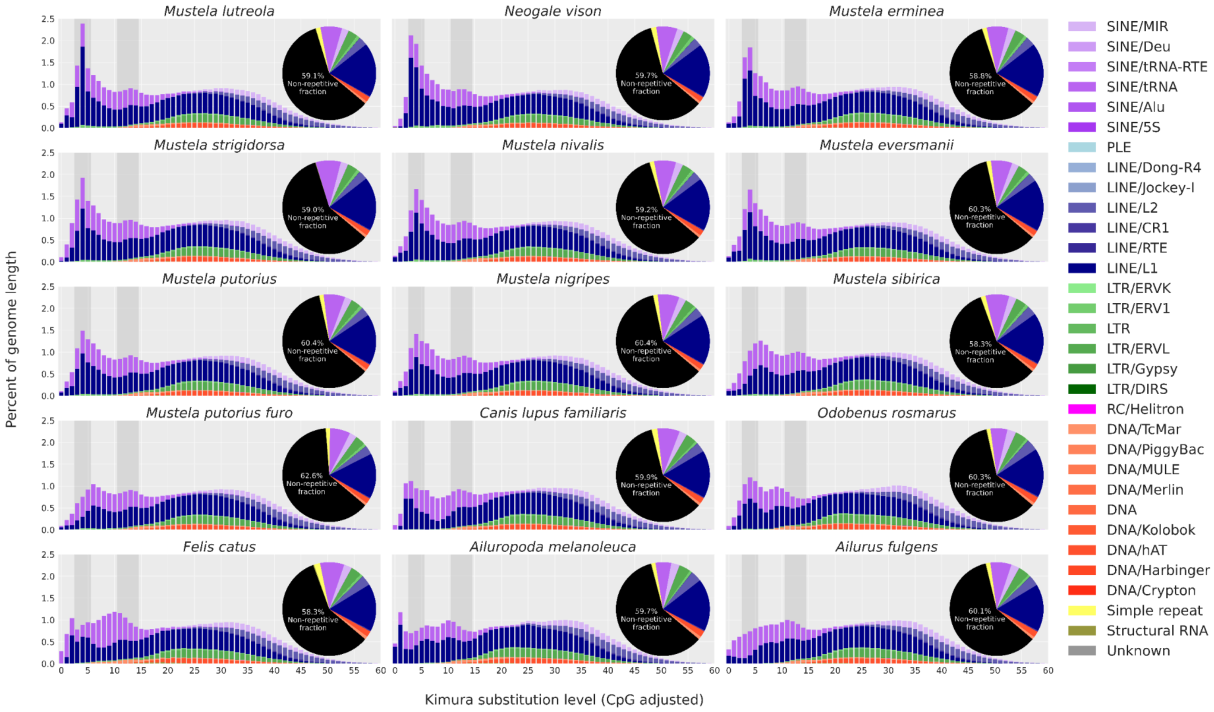


**SF4 (Supplementary Figure 4).** Centromere positions shift within the genus *Mustela*.

(A) Different centromere positions on MERM 12 and homologous chromosomes. (B) Different centromere positions on MERM 5 and homologous chromosomes. Chromosomes labeled by ‘ were reverse complemented. The dot on the G-bands indicates the position of the centromere. Original images of G-banded chromosomes from: (Graphodatsky and Radzhabli 1988; Cavagna et al. 2000; Graphodatsky et al. 2002; Nie et al. 2002; Graphodatsky et al. 2020; Kliver et al. 2023). Abbreviation: MFOI – *Martes foina*, ELUT – *Enhydra lutris*, NVIS – *Neogale vison*, MSTR – *Mustela strigidorsa*, MERM – *Mustela erminea*, MNIV – *Mustela nivalis*, MLUT – *Mustela lutreola*, MPFUR – *Mustela putorius furo*, MNIG – *Mustela nigripes*. Macro-level synteny is shown by gray lines, inversions (larger than 1 Mbp) are highlighted in red. Centromeric positions on chromosome scaffolds are tentative, and approximately drawn based on the analysis of comparative chromosome painting maps and G-banded karyotypes. Lutrinae (ELUT) and Gulioninae (MFOI) are used here as Mutelinae outgroups to demonstrate the ancestral centromeric position for the Mustelidae family. All Lutrinae, including ELUT, with reported karyotypes have 2n=38 and conserved centromeric positions based on conventional staining. For this figure in place of ELUT chromosomes, we used the only G-banded karyotype of Lutrinae (*Lutra lutra*) that has been published (Graphodatsky and Radzhabli 1988; Graphodatsky et al. 2020). Since the MSTR karyotype is not available and one-to-one synteny to MERM chromosome was observed (2n=44), the G-banded chromosomes of MERM are used to represent MSTR on this figure.

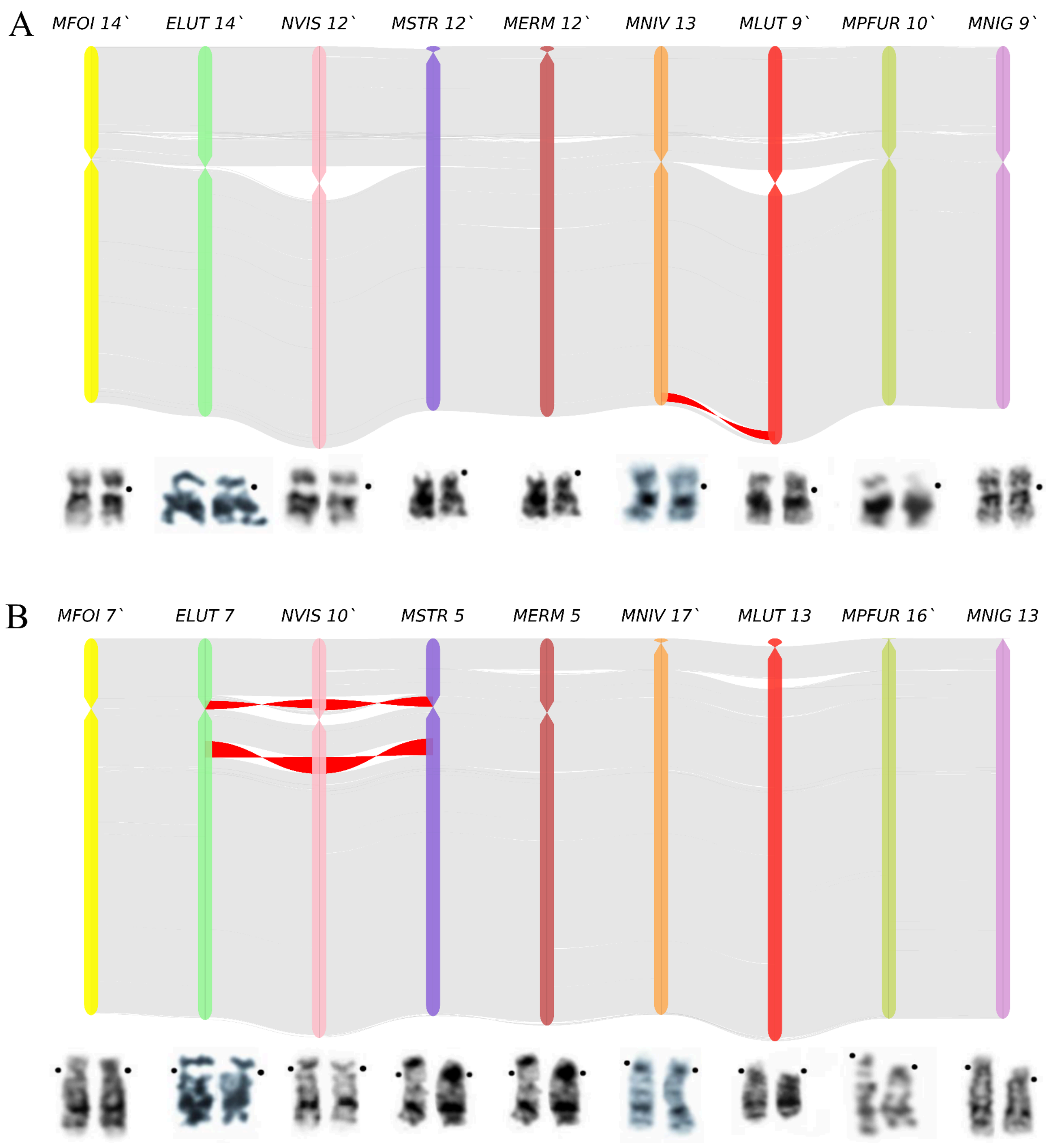

**SF5 (Supplementary Figure 5).** The Runs of Homozygosity (RoH) content in the samples. The fraction of different size categories of RoH fraction within the genome: Short RoH – <1 Mbp, Long RoH – >=1 Mbp, Ultra Long RoH – >=10 Mbp. Non-RoH indicates the fraction of the genome not covered by any detected RoH segments.

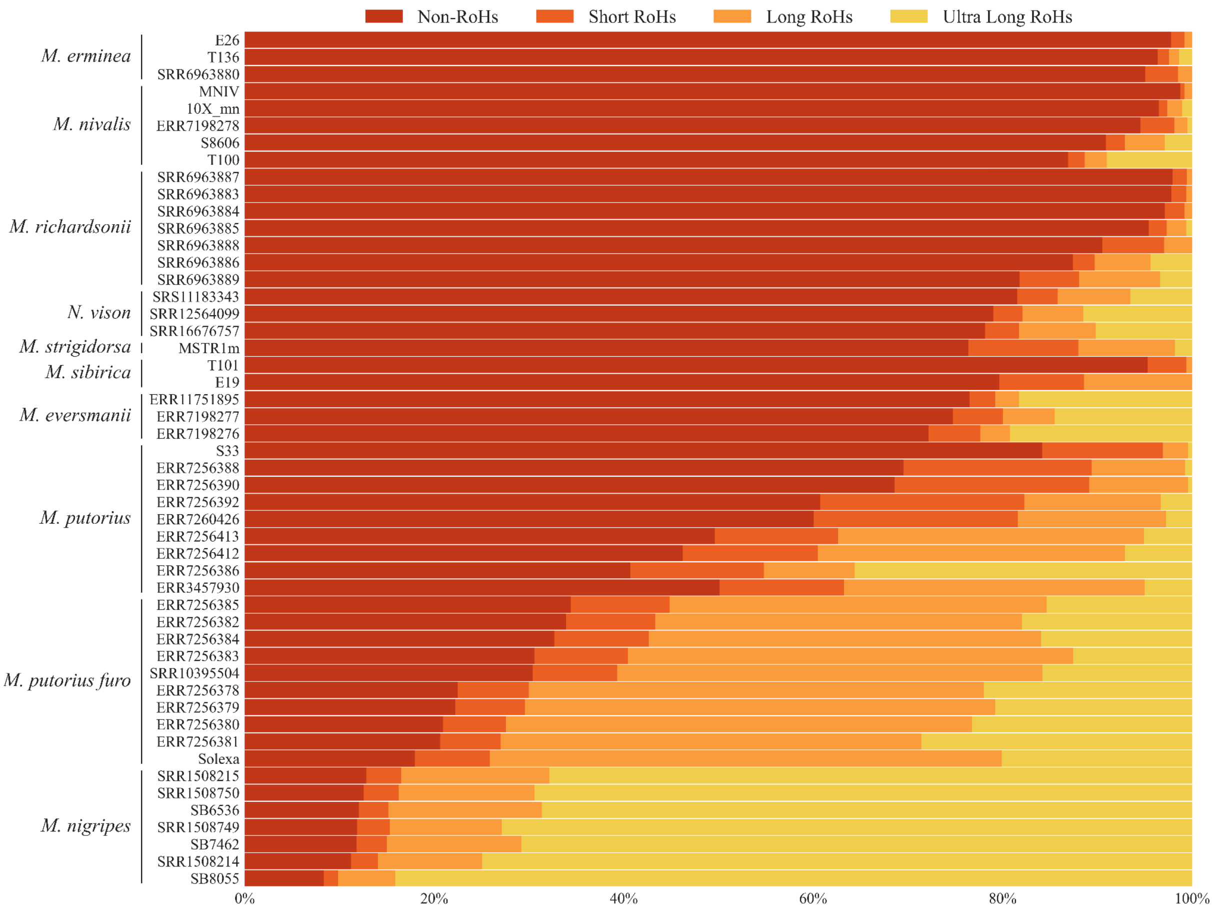


**SF6 (Supplementary Figure 6).** Pairwise distances between and within sister species in Mustelinae lineage.

A – *Mustela nivalis, B – Mustela erminea* and *Mustela richardsonii*; C – *Mustela eversmanii* and *Mustela nigripes*.

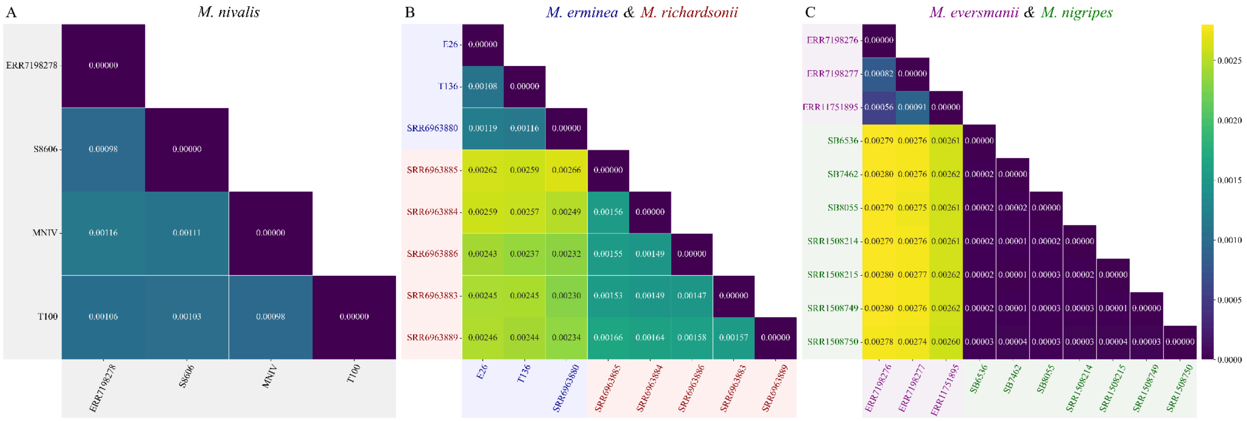

A
M. nivalis
ERR7198278
S8606
MNIV
T100
0.00000
0.00098 0.00000
0.00116 0.00111 0.00000
0.00106 0.00103 0.00098 0.00000
B
M. erminea & M. richardsonii
E26
T136
SRR6963880
SRR6963885
SRR6963884
SRR6963886
SRR6963883
SRR6963889
0.00000
0.00108 0.00000
0.00119 0.00116 0.00000
0.00262 0.00259 0.00266 0.00000
0.00259 0.00257 0.00249 0.00156 0.00000
0.00243 0.00237 0.00232 0.00155 0.00149 0.00000
0.00245 0.00245 0.00230 0.00153 0.00149 0.00147 0.00000
0.00246 0.00244 0.00234 0.00166 0.00164 0.00158 0.00157 0.00000
C
M. eversmanii & M. nigripes
ERR7198276
ERR7198277
ERR11751895
SB6536
SB7462
SB8055
SRR1508214
SRR1508215
SRR1508749
SRR1508750
0.00000
0.00082 0.00000
0.00056 0.00091 0.00000
0.00279 0.00276 0.00261 0.00000
0.00280 0.00276 0.00262 0.00002 0.00000
0.00279 0.00275 0.00261 0.00002 0.00002 0.00000
0.00279 0.00276 0.00261 0.00002 0.00001 0.00002 0.00000
0.00280 0.00277 0.00262 0.00002 0.00001 0.00003 0.00002 0.00000
0.00280 0.00276 0.00262 0.00002 0.00001 0.00003 0.00003 0.00001 0.00000
0.00278 0.00274 0.00260 0.00003 0.00004 0.00003 0.00003 0.00004 0.00003 0.00000
0.0025
0.0020
0.0015
0.0010
0.0005
0.0000

**SF7 (Supplementary Figure 7).** ASTRAL-III phylogenetic tree was reconstructed based on 6,599 single-copy BUSCO gene alignments. Individual gene trees were inferred using RAxML-NG v1.2.2 (1000 bootstraps, model GTR+G4). Nodes with bootstrap support values below 70 were excluded from the input trees. Numbers displayed at each node represent unique node identifiers, for which detailed statistical values are provided in Supplementary Table ST10. Colored dots indicate the main local posterior probabilities (pp1), expressed as percentages.

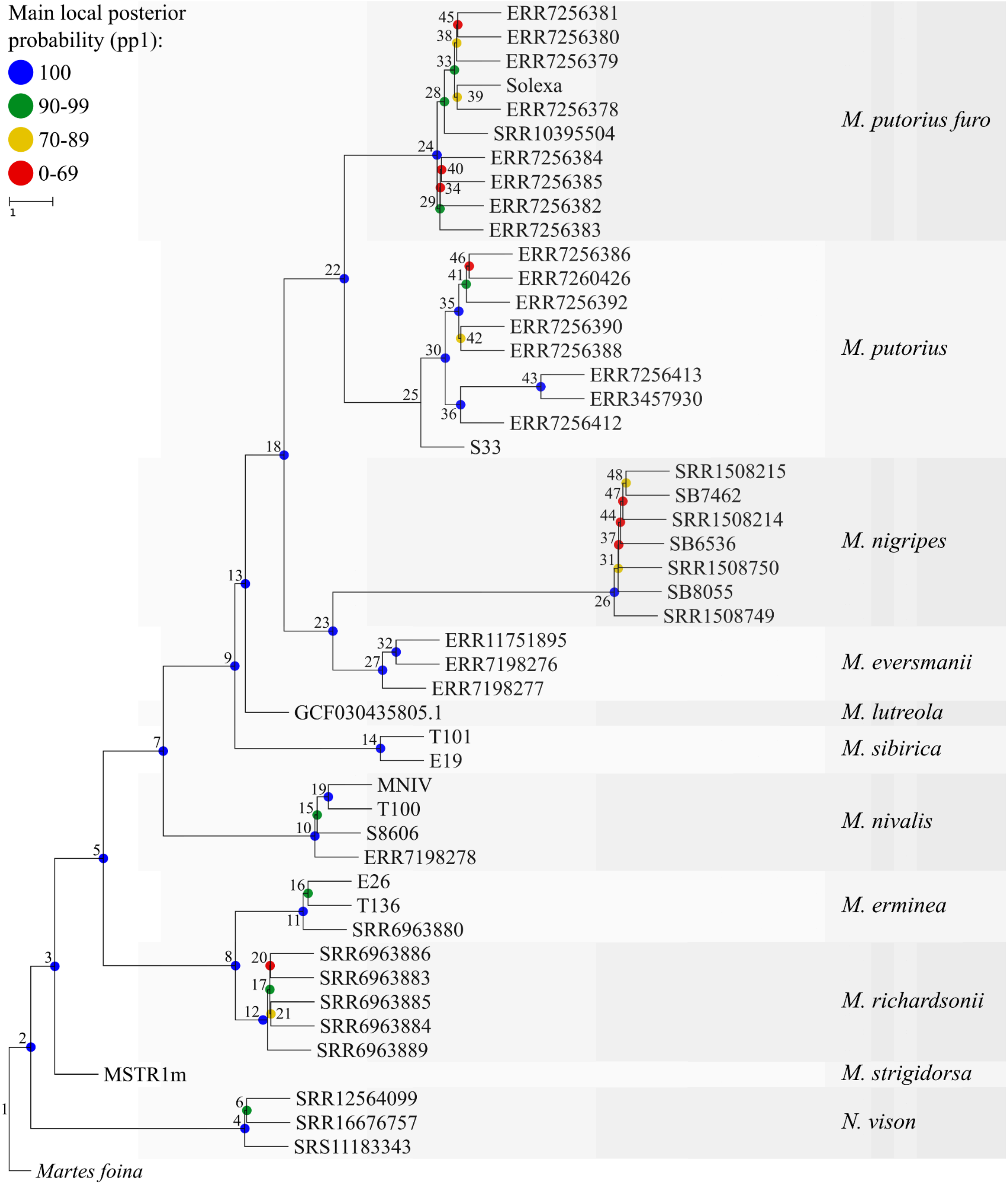

**SF8 (Supplementary Figure 8).** ASTRAL-III phylogenetic tree was reconstructed based on 6,599 single-copy BUSCO gene alignments. Individual gene trees were inferred using RAxML-NG v1.2.2 (1000 bootstraps, model GTR+G4). Nodes with bootstrap support values below 70 were excluded from the input trees. Detailed statistical values are provided in Supplementary Table ST10. Piecharts indicate the quartet supports.

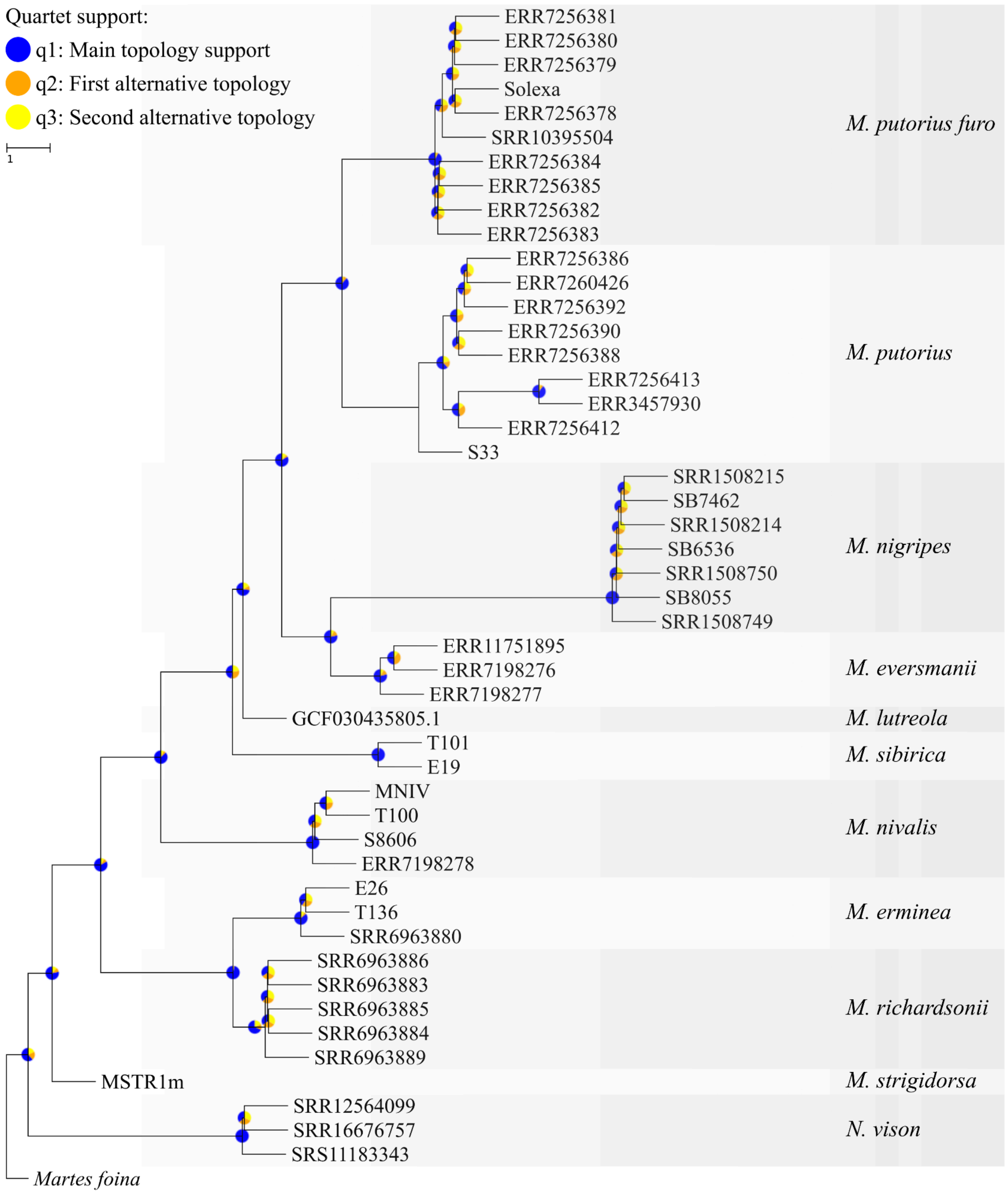

**SF9 (Supplementary Figure 9).** Chromosomal fusions and fission in the Mustelinae lineage.

Each row corresponds to a particular translocation: fission (dark gray) or fusion (light gray) of chromosomes.Yellow color indicates the presence of the translocation, and dark violet its absence. For example, consider a case of homoplasy involving fus(MSTR2; MSTR9). This fusion is present in MSIB, MLUT, MNIG, and MEVE, but absent in MPUT and MPFUR. According to the phylogeny, MLUT is the basal lineage to the two sister clades: MPUT-MPFUR and MNIG-MEVE. The most likely explanation is that the fusion appeared in their common ancestor and subsequently separated independently in MPUT and MPFUR. This is an example of homoplasy caused by secondary loss, where the same ancestral state was reversed in two different lineages. Mustelinae species: NVIS – *Neogale vison*, MSTR – *Mustela strigidorsa*, MERM – *Mustela erminea*, MNIV – *Mustela nivalis*, MSIB – *Mustela sibirica*, MLUT – *Mustela lutreola*, MPUT – *Mustela putorius*, MPFUR – *Mustela putorius furo*, MEVE – *Mustela eversmanii*, MNIG – *Mustela nigripes*. Outgroup: MFOI – *Martes foina*, PBRA – *Pteronura brasiliensis*, ELUT – *Enhydra lutris*, LCAN – *Lutra canadensis*, LLUT – *Lutra lutra*, ACIN – *Aonyx cinerea*. For additional details see caption for Figure 6 and methods.

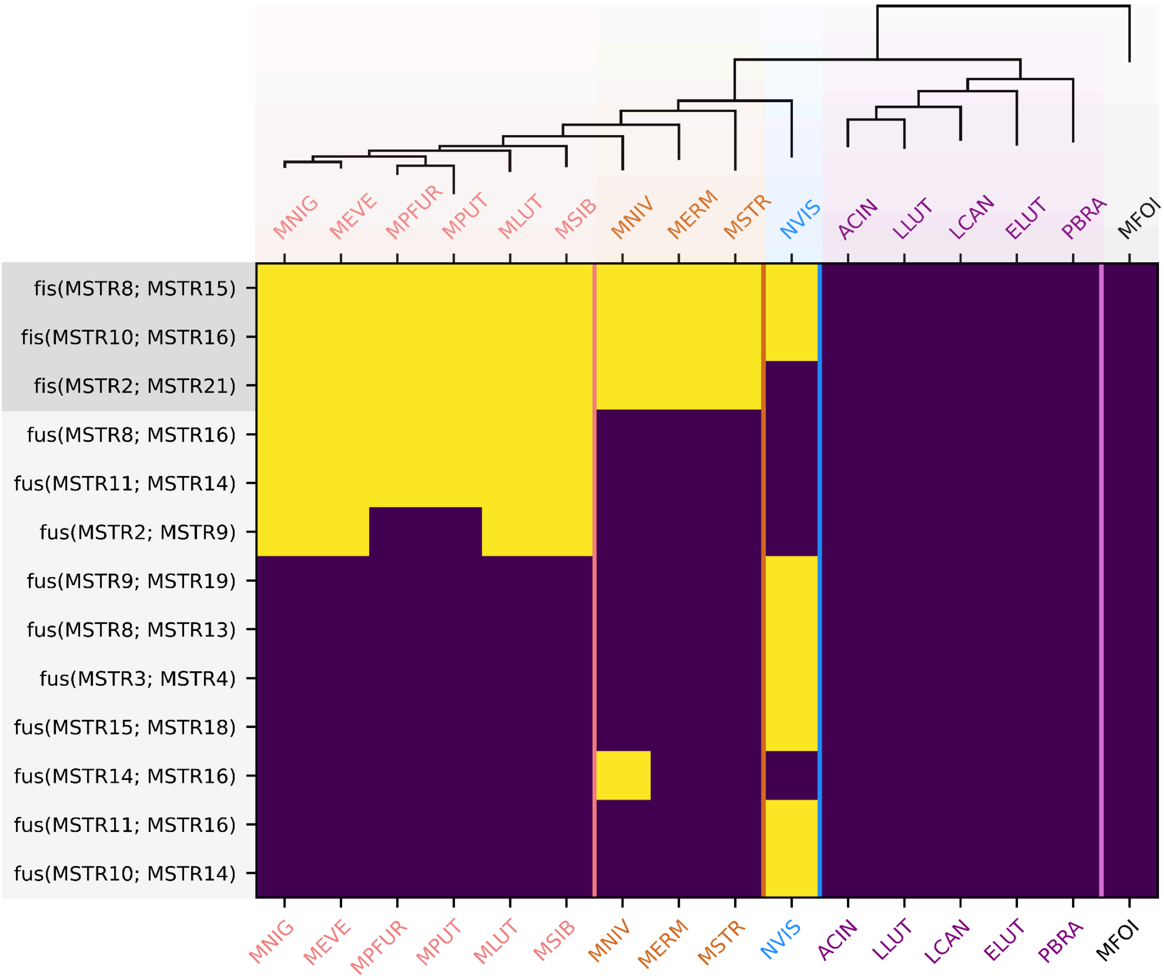

**SF10 (Supplementary Figure 10).** Maximum likelihood nuclear phylogeny of analyzed Mustelinae species with genome assemblies.

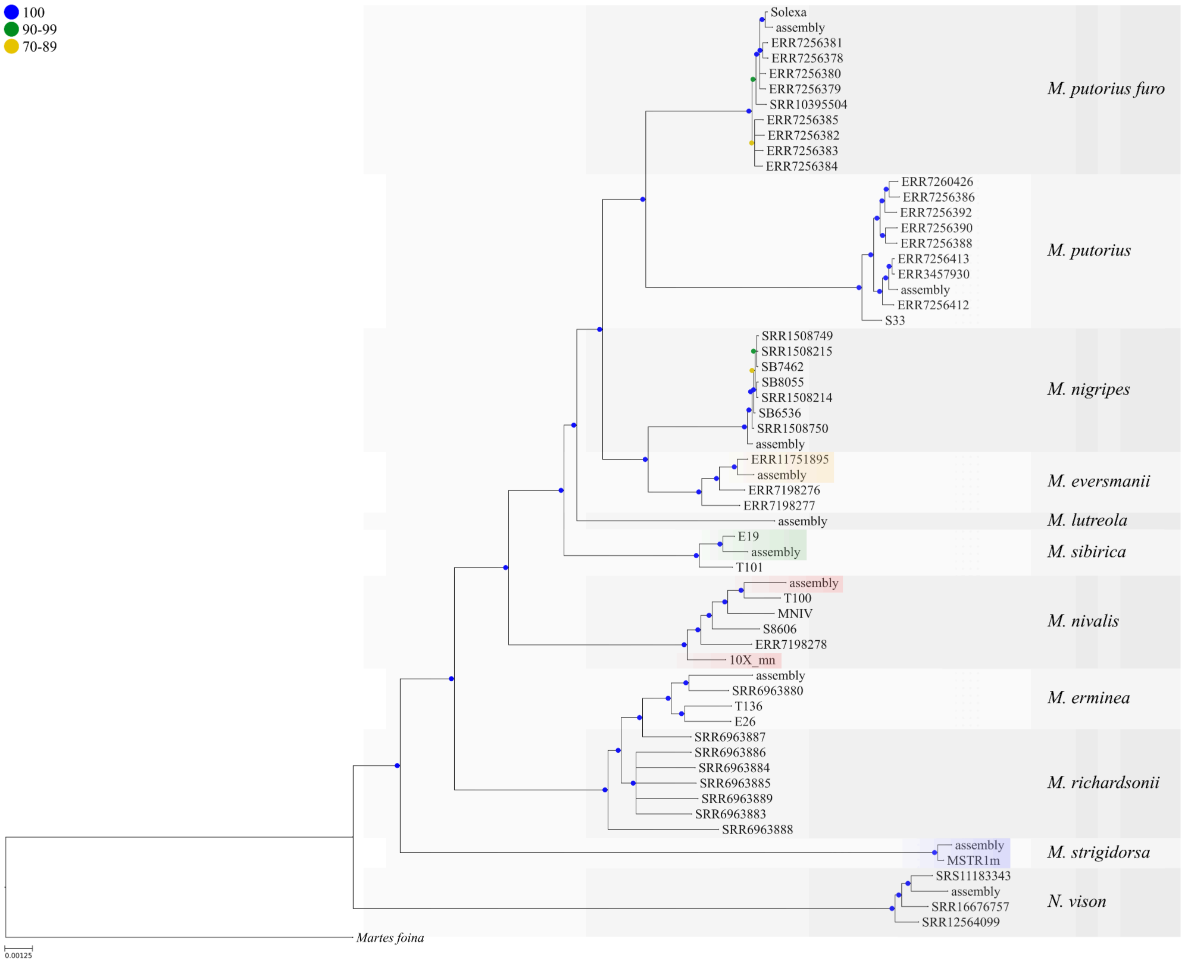
