## Supplementary Tables for "Comparative genomics and phylogenomics of the Mustelinae lineage (Mustelidae, Carnivora)"

**ST1 (Supplementary Table 1). Available genetic data for Mustelinae species.**

| Species | Abbreviation | Assembly | Resequencing data | Mitochondrial genome | STR | Nuclear or mitochondrial markers | Global conservation status | Generation time | Area | References |
|---|---|---|---|---|---|---|---|---|---|---|
| *Mustela erminea* | MERM | + | +* | +* | + | + | LC | 3 | Holarctic | (Sato et al. 2004; Kranz et al. 2015; Colella et al. 2018) |
| *Mustela lutreola* | MLUT | + | + | + | + | + | CR | 2 | Northwest Palearctic | (Cabria et al. 2007; Maran, Skumatov, Gomez, et al. 2015; Skorupski 2022; Skorupski et al. 2023) |
| *Mustela eversmanii* | MEVE | + | + | + | + | + | LC | 3.3 | Palearctic | (Maran, Skumatov, Kranz, et al. 2015; Szatmári et al. 2021; Etherington et al. 2022) |
| *Mustela nigripes* | MNIG | + | + | + | + | + | EN | 4 | Nearctic | (Belant et al. 2015; Zhao et al. 2016; Kliver et al. 2023) |
| *Mustela nivalis* | MNIV | +* | +* | +* | + | + | LC | 3.3 | Holarctic | (Kryštufek et al. 2015; Miranda et al. 2021) |
| *Mustela putorius* | MPUT | + | +* | +* | + | + | LC | 4.5 | Western Palearctic | (Maran et al. 2016; Szatmári et al. 2021; Etherington et al. 2022) |
| *Mustela sibirica* | MSIB | –** | –** | +* | + | + | LC | 4 | Eastern Palearctic | (Abramov, Timmins, Duckworth, Choudhury, Chan, Ghimirey, et al. 2015; Shalabi et al. 2017) |
| *Mustela strigidorsa* | MSTR | –** | –** | –** | – | + | LC | 6.4 | Indomalayan | (Koepfli et al. 2008; Abramov, Timmins, Duckworth, Choudhury, Dinets, et al. 2015) |
| *Mustela haidarum* | MHAI | – | + | + | – | – | NE | – | Western Nearctic | (Colella et al. 2018; Colella et al. 2021) |
| *Mustela richardsonii* | MRIC | – | + | + | – | – | NE | – | Nearctic | (Colella et al. 2018; Colella et al. 2021) |
| *Mustela itatsi* | MITA | – | – | + | + | + | NT | 5.3 | Eastern Palearctic | (Abramov, Kaneko, and Masuda 2015; Shalabi et al. 2017) |
| *Mustela altaica* | MALT | – | – | + | – | + | NT | 5 | Southeastern Palearctic | (Sato et al. 2004; Huang et al. 2014; Abramov 2015) |
| *Mustela aistoodonnivalis* | MAIS | – | – | – | – | + | – | – | Indomalayan | (Liu et al. 2023) |
| *Mustela kathiah* | MKAT | – | – | + | – | + | LC | 3.4 | Indomalayan | 'u et al. 2011; Abramov et al. 2013; Abramov, Timmins, Duckworth, Choudhury, Chan, Lau, et al. 201 |
| *Mustela nudipes* | MNUD | – | – | +^ | – | + | LC | 6.6 | Indomalayan | (Koepfli et al. 2008; Duckworth et al. 2015; Hassanin et al. 2021) |
| *Mustela lutreolina* | MLIN | – | – | – | – | – | LC | 4.8 | Indomalayan | (Abramov, Duckworth, Meijaard, et al. 2015) |
| *Neogale felipei* | NFEL | – | – | – | – | + | VU | 5.3 | Neotropical | (Harding and Smith 2009; González-Maya et al. 2016) |
| *Neogale frenata* | NFRE | – | – | + | – | + | LC | – | Nearctic & Neotropical | (Koepfli et al. 2008; Yu et al. 2011; Helgen and Reid 2015) |
| *Neogale africana* | NAFR | – | – | – | – | + | LC | – | Neotropical | (Harding and Smith 2009; Helgen and Emmons 2015) |
| *Neogale vison* | NVIS | + | + | + | + | + | LC | – | Holarctic | (Reid et al. 2015; Karimi et al. 2022; Lukashkova et al. 2023) |

* Data complemented by us;

** Data obtained by us for the first time;

^ Mitochondrial genome of *M. nudipes* probably was misidentified as a different species (MH464792) (Hassanin et al. 2021).

**ST2 (Supplementary Table 2). Assignments of the chromosome IDs in the *Mustela nivalis* genome assembly.**

| *M. nivalis* assembly | | Homologous *N. vison* chromosomes (WGA *) | Homologous *N. vison* chromosomes (cytogenetic data **) | Homologous *M. lutreola* chromosomes (WGA *) | Homologous *M. lutreola* chromosomes (cytogenetic data **) |
|---|---|---|---|---|---|
| **C-scaffold ID** | **Chromosome** | | | | |
| HiC_scaffold_1 | chr3 | 5 | 5 | 2 | 2 |
| HiC_scaffold_2 | chr4 | 2, 3 | 2, 3 | 3, 4 | 3, 4 |
| HiC_scaffold_3 | chr1 | 6 | 6 | 1 | 1 |
| HiC_scaffold_4 | chr2 | 1 | 1 | 12 | 12 |
| HiC_scaffold_5 | chr9 | 1 | 1 | 5 | 5 |
| HiC_scaffold_6 | chr17 | 10 | 10 | 13 | 13 |
| HiC_scaffold_7 | chr10 | 9 | 9 | 6 | 6 |
| HiC_scaffold_8 | chr11 | 11 | 11 | 7 | 7 |
| HiC_scaffold_9 | chr7 | 4 | 4 | 3 | 3 |
| HiC_scaffold_10 | chr5 | 7 | 7 | 1 | 1 |
| HiC_scaffold_11 | chr6 | 2 | 2 | 14 | 14 |
| HiC_scaffold_12 | chr8 | 3 | 3 | 4 | 4 |
| HiC_scaffold_13 | chr13 | 12 | 12 | 9 | 9 |
| HiC_scaffold_14 | chr14 | 4 | 4 | 8 | 8 |
| HiC_scaffold_15 | chr12 | 8 | 8 | 15 | 15 |
| HiC_scaffold_16 | chr15 | 13 | 13 | 10 | 10 |
| HiC_scaffold_17 | chr18 | 8 | 8 | 16 | 16 |
| HiC_scaffold_18 | chr19 | 7 | 7 | 17 | 17 |
| HiC_scaffold_19 | chr20 | 6 | 6 | 18 | 18 |
| HiC_scaffold_20 | chr16 | 14 | 14 | 11 | 11 |
| HiC_scaffold_21 | chrX | X | X | X | X |

* WGA — based on pairwise whole-genome alignment.

** Cytogenetic data — summary of hybridization patterns of American mink (N. vison) probes onto G-banded mustelid chromosomes (Figure 3a and 3b by (Graphodatsky et al. 2002)).

**ST3 (Supplementary Table 3). Assignments of the chromosome IDs in the *Mustela strigidorsa* genome assembly.**

| *M. strigidorsa* assembly | | Homologous *M. erminea* chromosomes (WGA *) |
|---|---|---|
| **C-scaffold ID** | **Chromosome** | |
| HiC_scaffold_1 | chr1 | 1 |
| HiC_scaffold_2 | chr19 | 19 |
| HiC_scaffold_3 | chr18 | 18 |
| HiC_scaffold_4 | chr10 | 10 |
| HiC_scaffold_5 | chr9 | 9 |
| HiC_scaffold_6 | chr2 | 2 |
| HiC_scaffold_7 | chr14 | 14 |
| HiC_scaffold_8 | chr13 | 13 |
| HiC_scaffold_9 | chr5 | 5 |
| HiC_scaffold_10 | chr7 | 7 |
| HiC_scaffold_11 | chr16 | 16 |
| HiC_scaffold_12 | chr6 | 6 |
| HiC_scaffold_13 | chr17 | 17 |
| HiC_scaffold_14 | chr20 | 20 |
| HiC_scaffold_15 | chr4 | 4 |
| HiC_scaffold_16 | chr3 | 3 |
| HiC_scaffold_17 | chr21 | 21 |
| HiC_scaffold_18 | chr8 | 8 |
| HiC_scaffold_19 | chr12 | 12 |
| HiC_scaffold_20 | chr11 | 11 |
| HiC_scaffold_21 | chr15 | 15 |
| HiC_scaffold_22 | chrY | Y |
| HiC_scaffold_23 | chrX | X |

* WGA — based on pairwise whole-genome alignment.

**ST4 (Supplementary Table 4). Quality metrics for genome assemblies (Scaffolds length >= 1000 bp).**

| Species | 2n | Assembly name/ID | Assembly type * | Sequencing technology | Number of scaffolds | Genome length, Gbp | Ns, Mbp | N50, Mbp | L50 |
|---|---|---|---|---|---|---|---|---|---|
| *M. nivalis* | 42 | MNIV1m | Chr | HiC | 16872 | 2.45 | 22.01 | 138.37 | 8 |
| *M. strigidorsa* | 44 | MSTR1m | Chr | Illumina + Nanopore + Hi-C | 29284 | 2.42 | 2.4 | 115.1 | 8 |
| *M. erminea* | 44 | GCF_009829155.1 | Chr | PacBio + llumina + Hi-C + Bionano | 94 | 2.45 | 21.48 | 130.15 | 8 |
| *M. sibirica* | 38 | MSIB1m + GCF_009829155.1 | pChr | llumina + PacBio + Hi-C + Bionano | 13639 | 2.41 | 26.69 | 113.57 | 8 |
| *M. eversmanii* | 38 | GCA_963422785.1 + M. nigripes (DNAZoo) | pChr | 10X Genomics + Bionano | 25219 | 2.55 | 64.12 | 146.97 | 7 |
| *M. nigripes* | 38 | DNAZoo | Chr | 10X Genomics + Bionano + Hi-C | 20510 | 2.5 | 82.35 | 145.43 | 7 |
| *M. putorius* | 40 | M. putorius (GCA_902207235.1 ) + M. putorius furo (DNAZoo) | pChr | 10X Genomics + Illumina + Bionano + Hi-C | 11913 | 2.5 | 23.15 | 150.61 | 7 |
| *M. putorius furo* | 40 | DNAZoo | Chr | Illumina + Hi-C | 7417 | 2.41 | 131.95 | 145.37 | 7 |
| *M. lutreola* | 38 | GCF_030435805.1 | Chr | PacBio + OmniC | 25 | 2.59 | 0.007800 | 154.08 | 7 |
| *N. vison* | 30 | GCF_020171115.1 | Chr | PacBio + Illumina | 181 | 2.68 | 0 | 220.35 | 6 |

* Assembly type: Chr – chromosome-level genome assembly; pChr – pseudochromosome-level genome assembly.

**ST5 (Supplementary Table 5). Samples quality. Number of reads, k-mers coverage, genome sizes and downsampling fraction of sequenced individuals.**

| Species | Sample | Read length | Number of reads, Mbp | | Kmer multiplicity at first maximum | Estimated haplome coverage | | Coverage | Genome size, Gbp | | Downsampling fraction * |
|---|---|---|---|---|---|---|---|---|---|---|---|
| | | | before filtering | after filtering | | | | | | | |
| *M. nivalis* | MNIV | 150+150 | 167.7 | 162.7 | 12 | 6.56 | ±0.01 | 13.12 | 03.08 | ±0.00 | 0.91 |
| | T100 | 150+150 | 193.8 | 190.6 | 12 | 7.12 | ±0.05 | 14.24 | 3.35 | ±0.02 | 0.84 |
| | 10X_mn | 128+150 | 805.2 | 618.3 | 35 | 19.13 | ±0.03 | 38.26 | 3.21 | ±0.00 | 0.31 |
| | ERR7198278 | 126+126 | 228.7 | 193.4 | 11 | 6.64 | ±0.01 | 13.28 | 2.84 | ±0.00 | 0.90 |
| | S8606 | 150+150 | 667.3 | 651.6 | 46 | 24.65 | ±0.02 | 49.3 | 3.22 | ±0.00 | 0.24 |
| *M. strigidorsa* | MSTR1m | 150+150 | 225.7 | 214.7 | 15 | 7.73 | ±0.03 | 15.46 | 3.34 | ±0.01 | 0.78 |
| *M. sibirica* | E19 | 150+150 | 256.3 | 251.1 | 23 | 11.90 | ±0.03 | 23.8 | 2.59 | ±0.00 | 0.50 |
| | T101 | 150+150 | 187.9 | 183.8 | 17 | 8.97 | ±0.01 | 17.94 | 2.54 | ±0.00 | 0.67 |
| *M. erminea* | T136 | 150+150 | 270.4 | 264.8 | 22 | 11.71 | ±0.01 | 23.42 | 2.81 | ±0.01 | 0.51 |
| | E26 | 150+150 | 243.9 | 238.7 | 20 | 10.70 | ±0.01 | 21.4 | 2.76 | ±0.01 | 0.56 |
| | SRR6963880 | 100+100 | 220.4 | 201.9 | 9 | 5.75 | ±0.01 | 11.5 | 2.64 | ±0.01 | 01.04 |
| *M. richardsonii* | SRR6963883 | 100+100 | 250.0 | 215.8 | 10 | 6.23 | ±0.01 | 12.46 | 2.58 | ±0.00 | 0.96 |
| | SRR6963884 | 100+100 | 261.9 | 216.6 | 10 | 06.01 | ±0.01 | 12.02 | 2.67 | ±0.00 | 1.00 |
| | SRR6963885 | 100+100 | 250.0 | 236.2 | 11 | 6.46 | ±0.04 | 12.92 | 2.79 | ±0.02 | 0.93 |
| | SRR6963886 | 100+100 | 242.5 | 221.5 | 11 | 6.24 | ±0.01 | 12.48 | 2.66 | ±0.00 | 0.96 |
| | SRR6963887 | 100+100 | 216.7 | 194.7 | 9 | 5.37 | ±0.01 | 10.74 | 2.74 | ±0.00 | 1.12 |
| | SRR6963888 | 100+100 | 225.7 | 205.8 | 10 | 5.91 | ±0.07 | 11.82 | 2.62 | ±0.03 | 01.02 |
| | SRR6963889 | 100+100 | 225.4 | 204.3 | 9 | 6.64 | ±0.02 | 13.28 | 2.29 | ±0.01 | 0.90 |
| *M. nigripes* | SRR1508214 | 100+100 | 281 | 270.5 | 14 | 7.68 | ±0.01 | 15.36 | 2.70 | ±0.00 | 0.78 |
| | SRR1508215 | 100+100 | 233.3 | 225.9 | 12 | 6.30 | ±0.01 | 12.6 | 2.71 | ±0.00 | 0.95 |
| | SRR1508750 | 100+100 | 253.1 | 244.3 | 13 | 07.01 | ±0.01 | 14.02 | 2.64 | ±0.00 | 0.86 |
| | SRR1508749 | 100+100 | 221.9 | 214.2 | 11 | 6.15 | ±0.01 | 12.3 | 2.64 | ±0.00 | 0.98 |
| | SB7462 | 150+150 | 422.6 | 401.3 | 34 | 17.35 | ±0.01 | 34.7 | 2.75 | ±0.00 | 0.35 |
| | SB8055 | 150+150 | 437.9 | 421.9 | 38 | 19.51 | ±0.01 | 39.02 | 2.59 | ±0.00 | 0.31 |
| | SB6536 | 128+150 | 1.16 Gbp | 1.05 Gbp | 72 | 37.73 | ±0.02 | 75.46 | 2.79 | ±0.00 | 0.16 |
| | ERR3457930 | 239+225 | 374.04 | 61.1 | 48 | 25.28 | ±0.01 | 50.56 | 2.54 | ±0.00 | 0.24 |
| | ERR7256386 | 236+215 | 73.7 | 64.9 | 9 | 5.75 | ±0.02 | 11.5 | 2.21 | ±0.01 | 01.04 |
| | ERR7256388 | 236+215 | 65.7 | 61.7 | 9 | 5.49 | ±0.02 | 10.98 | 2.21 | ±0.01 | 01.09 |
| | ERR7256390 | 236+215 | 64.7 | 61.1 | 8 | 5.22 | ±0.02 | 10.44 | 2.3 | ±0.01 | 1.15 |

| | | | | | | | | | | | |
|---|---|---|---|---|---|---|---|---|---|---|---|
| *M. putorius* | ERR7256392 | 230+205 | 67.6 | 62.7 | 8 | 5.27 | ±0.02 | 10.54 | 2.23 | ±0.01 | 1.14 |
| | ERR7256412 | 126+126 | 144.2 | 135 | 9 | 5.21 | ±0.01 | 10.42 | 2.58 | ±0.00 | 1.15 |
| | ERR7256413 | 250+250 | 175.1 | 152.9 | 23 | 12.56 | ±0.01 | 25.12 | 2.53 | ±0.00 | 0.48 |
| | ERR7260426 | 225+225 | 64.9 | 60.4 | 8 | 5.25 | ±0.02 | 10.5 | 2.16 | ±0.01 | 1.14 |
| | S33 | 150+150 | 333.6 | 326.2 | 30 | 15.35 | ±0.01 | 30.7 | 2.63 | ±0.00 | 0.39 |
| | Solexa | 100+100 | 710.2 | 704.1 | 40 | 20.71 | ±0.01 | 41.42 | 2.47 | ±0.00 | 0.29 |
| | SRR10395504 | 150+150 | 1.32 Gbp | 1.27 Gbp | 60 | 58.91 | ±0.02 | 117.82 | 2.59 | ±0.00 | 0.10 |
| | ERR7256378 | 123+123 | 204.8 | 200.8 | 15 | 7.92 | ±0.01 | 15.84 | 2.53 | ±0.00 | 0.76 |
| | ERR7256379 | 123+123 | 203.2 | 198.8 | 15 | 8.10 | ±0.01 | 16.2 | 2.46 | ±0.00 | 0.74 |
| | ERR7256380 | 123+123 | 185.2 | 181.5 | 13 | 7.27 | ±0.01 | 14.54 | 2.48 | ±0.00 | 0.83 |
| *M. putorius furo* | ERR7256381 | 123+123 | 201.2 | 196.9 | 14 | 7.79 | ±0.01 | 15.58 | 2.52 | ±0.00 | 0.77 |
| | ERR7256382 | 123+123 | 210.5 | 205.7 | 15 | 8.53 | ±0.01 | 17.06 | 2.4 | ±0.00 | 0.70 |
| | ERR7256383 | 123+123 | 212.6 | 207.1 | 14 | 9.10 | ±0.03 | 18.2 | 2.26 | ±0.01 | 0.66 |
| | ERR7256384 | 123+123 | 209.5 | 204.5 | 15 | 8.54 | ±0.01 | 17.08 | 2.38 | ±0.00 | 0.70 |
| | ERR7256385 | 123+123 | 208.4 | 204.2 | 14 | 8.19 | ±0.01 | 16.38 | 2.5 | ±0.00 | 0.73 |
| | ERR11751895 | 228+250 | 452.1 | 443.7 | 55 | 29.39 | ±0.02 | 58.78 | 2.83 | ±0.00 | 0.20 |
| *M. eversmanni* | ERR7198276 | 225+199 | 65.2 | 61.7 | 7 | 4.99 | ±0.02 | 9.98 | 2.28 | ±0.01 | 1.20 |
| | ERR7198277 | 123+122 | 131.3 | 121.4 | 7 | 4.91 | ±0.02 | 9.82 | 2.45 | ±0.01 | 1.22 |
| | SRR12564099 | 150+150 | 407.6 | 395.7 | 36 | 18.70 | ±0.01 | 37.4 | 2.61 | ±0.00 | 0.32 |
| *N. vison* | SRR16676757 | 150+150 | 207.7 | 203.4 | 18 | 9.77 | ±0.01 | 19.54 | 2.58 | ±0.00 | 0.61 |
| | SRS11183343 | 100+100 | 898.7 | 857.2 | 52 | 26.13 | ±0.01 | 52.26 | 2.47 | ±0.00 | 0.23 |

* Downsampling was not performed for samples with a fraction value greater than or equal to 0.8.

**ST6 (Supplementary Table 6). Completeness of genome assemblies assessed using BUSCO v5.6.1.**

| OrthoDB v10.1 dataset (2024-01-08) | Total BUSCOs in dataset | Species * | Complete BUSCOs | | Complete single-copy BUSCOs | | Complete duplicated BUSCOs | | Fragmented BUSCOs | | Missing BUSCOs | |
|---|---|---|---|---|---|---|---|---|---|---|---|---|
| | | | abs. | % | abs. | % | abs. | % | abs. | % | abs. | % |
| Mammalia | 9226 | *M. nigripes (Chr)* | 8857 | 96 | 8662 | 93.9 | 195 | 2.1 | 114 | 1.2 | 255 | 2.8 |
| | | *M. putorius furo (Chr)* | 8714 | 94.4 | 8627 | 93.5 | 87 | 0.9 | 170 | 1.8 | 342 | 3.8 |
| | | *M. putorius (pChr)* | 7860 | 85.2 | 7760 | 84.1 | 100 | 1.1 | 524 | 5.7 | 842 | 9.1 |
| | | *M. eversmanii (pChr)* | 8842 | 95.8 | 8701 | 94.3 | 141 | 1.5 | 109 | 1.2 | 275 | 3.0 |
| | | *M. nivalis (Chr)* | 8853 | 96 | 8709 | 94.4 | 144 | 1.6 | 105 | 1.1 | 268 | 2.9 |
| | | *M. strigidorsa (Chr)* | 8728 | 94.6 | 8611 | 93.3 | 117 | 1.3 | 136 | 1.5 | 362 | 3.9 |
| | | *M. erminea (Chr)* | 8874 | 96.2 | 8758 | 94.9 | 116 | 1.3 | 85 | 0.9 | 267 | 2.9 |
| | | *M. sibirica (pChr)* | 8865 | 96.1 | 8749 | 94.8 | 116 | 1.3 | 102 | 1.1 | 259 | 2.8 |
| | | *N. vison (Chr)* | 8893 | 96.4 | 8762 | 95 | 131 | 1.4 | 93 | 1 | 240 | 2.6 |
| Laurasiatheria | 12234 | *M. nigripes (Chr)* | 11643 | 95.2 | 11391 | 93.1 | 252 | 2.1 | 152 | 1.2 | 439 | 3.6 |
| | | *M. putorius furo (Chr)* | 11553 | 94.4 | 11431 | 93.4 | 122 | 1 | 206 | 1.7 | 475 | 3.9 |
| | | *M. putorius (pChr)* | 10622 | 86.8 | 10490 | 85.7 | 132 | 1.1 | 652 | 5.3 | 960 | 7.9 |
| | | *M. eversmanii (pChr)* | 11606 | 94.9 | 11399 | 93.2 | 207 | 1.7 | 178 | 1.5 | 450 | 3.6 |
| | | *M. nivalis (Chr)* | 11643 | 95.1 | 11455 | 93.6 | 188 | 1.5 | 153 | 1.3 | 438 | 3.6 |
| | | *M. strigidorsa (Chr)* | 11487 | 93.9 | 11328 | 92.6 | 159 | 1.3 | 189 | 1.5 | 558 | 4.6 |
| | | *M. erminea (Chr)* | 11666 | 95.3 | 11506 | 94 | 160 | 1.3 | 126 | 1 | 442 | 3.7 |
| | | *M. sibirica (pChr)* | 11646 | 95.2 | 11492 | 93.9 | 154 | 1.3 | 149 | 1.2 | 439 | 3.6 |
| | | *N. vison (Chr)* | 11706 | 95.7 | 11510 | 94.1 | 196 | 1.6 | 113 | 0.9 | 415 | 3.4 |
| Carnivora | 14502 | *M. nigripes (Chr)* | 13715 | 94.5 | 13391 | 92.3 | 324 | 2.2 | 179 | 1.2 | 608 | 4.3 |
| | | *M. putorius furo (Chr)* | 13635 | 94 | 13457 | 92.8 | 178 | 1.2 | 212 | 1.5 | 655 | 4.5 |
| | | *M. putorius (pChr)* | 12309 | 84.9 | 12133 | 83.7 | 176 | 1.2 | 638 | 4.4 | 1555 | 10.7 |
| | | *M. eversmanii (pChr)* | 13649 | 94.2 | 13351 | 92.1 | 298 | 2.1 | 200 | 1.4 | 653 | 4.4 |
| | | *M. nivalis (Chr)* | 13707 | 94.5 | 13448 | 92.7 | 259 | 1.8 | 166 | 1.1 | 629 | 4.4 |
| | | *M. strigidorsa (Chr)* | 13490 | 93.1 | 13264 | 91.5 | 226 | 1.6 | 205 | 1.4 | 807 | 5.5 |
| | | *M. erminea (Chr)* | 13760 | 94.8 | 13522 | 93.2 | 238 | 1.6 | 148 | 1 | 594 | 4.2 |
| | | *M. sibirica (pChr)* | 13720 | 94.7 | 13495 | 93.1 | 225 | 1.6 | 157 | 1.1 | 625 | 4.2 |
| | | *N. vison (Chr)* | 13800 | 95.1 | 13532 | 93.3 | 268 | 1.8 | 132 | 0.9 | 570 | 4 |

* Assembly type: Chr – chromosome-level genome assembly; pChr – pseudochromosome-level genome assembly.

**ST7 (Supplementary Table 7). Transposable elements in genome assemblies.**

| Transposable elements | *M. nivalis* | | *M. strigidorsa* | | *M. erminea* | | *M. nigripes* | | *M. sibirica* | | *M. eversmanii* | | *M. putorius* | | *M. putorius furo* | | *N. vison* | |
|---|---|---|---|---|---|---|---|---|---|---|---|---|---|---|---|---|---|---|
| | **Length, bp** | **Length, %** | **Length, bp** | **Length, %** | **Length, bp** | **Length, %** | **Length, bp** | **Length, %** | **Length, bp** | **Length, %** | **Length, bp** | **Length, %** | **Length, bp** | **Length, %** | **Length, bp** | **Length, %** | **Length, bp** | **Length, %** |
| Retroelements | 894084071 | 36.45 | 918586772 | 36.89 | 899953705 | 36.80 | 880332663 | 35.23 | 892302471 | 35.24 | 903233886 | 35.45 | 873770992 | 35.32 | 798488117 | 33.13 | 962176199 | 35.89 |
| SINEs | 247914698 | 10.11 | 273308052 | 10.98 | 246456942 | 10.08 | 249508254 | 9.99 | 262152139 | 10.35 | 250121976 | 9.82 | 245621060 | 9.93 | 231675859 | 9.61 | 257947145 | 9.62 |
| Penelope | 57589 | 0.00 | 58275 | 0.00 | 57262 | 0.00 | 57626 | 0.00 | 57516 | 0.00 | 56700 | 0.00 | 56511 | 0.00 | 56850 | 0.00 | 57287 | 0.00 |
| LINEs | 533790057 | 21.76 | 532182702 | 21.37 | 539806520 | 22.08 | 518551456 | 20.75 | 513303624 | 20.27 | 538686843 | 21.14 | 517014330 | 20.9 | 458419554 | 19.02 | 584189646 | 21.79 |
| CRE/SLACS | 0 | 0.00 | 0 | 0.00 | 0 | 0.00 | 0 | 0.00 | 0 | 0.00 | 0 | 0.00 | 0 | 0.00 | 0 | 0.00 | 0 | 0.00 |
| L2/CR1/Rex | 82192496 | 3.35 | 82518422 | 3.31 | 83614387 | 3.42 | 82865339 | 3.32 | 84909443 | 3.35 | 83182117 | 3.26 | 81534271 | 3.3 | 81213027 | 3.37 | 84010161 | 3.13 |
| R1/LOA/Jockey | 0 | 0.00 | 0 | 0.00 | 0 | 0.00 | 0 | 0.00 | 0 | 0.00 | 0 | 0.00 | 0 | 0.00 | 0 | 0.00 | 0 | 0.00 |
| R2/R4/NeSL | 78334 | 0.00 | 79828 | 0.00 | 78913 | 0.00 | 79558 | 0.00 | 83287 | 0.00 | 79626 | 0.00 | 77005 | 0.00 | 78253 | 0.00 | 79405 | 0.00 |
| RTE/Bov-B | 2603526 | 0.11 | 2624672 | 0.11 | 2669016 | 0.11 | 2622782 | 0.10 | 2692701 | 0.11 | 2638241 | 0.10 | 2585528 | 0.1 | 2594494 | 0.11 | 2665482 | 0.10 |
| L1/CIN4 | 448871804 | 18.30 | 446917791 | 17.95 | 453400007 | 18.54 | 432939776 | 17.33 | 425573537 | 16.81 | 452743041 | 17.77 | 432773785 | 17.49 | 374491811 | 15.54 | 497390192 | 18.55 |
| LTR elements | 112379316 | 4.58 | 113096018 | 4.54 | 113690243 | 4.65 | 112272953 | 4.49 | 116846708 | 4.61 | 114425067 | 4.49 | 111135602 | 4.49 | 108392704 | 4.50 | 120039408 | 4.48 |
| BEL/Pao | 0 | 0.00 | 0 | 0.00 | 0 | 0.00 | 0 | 0.00 | 0 | 0.00 | 0 | 0.00 | 0 | 0.00 | 0 | 0.00 | 0 | 0.00 |
| Ty1/Copia | 0 | 0.00 | 0 | 0.00 | 0 | 0.00 | 0 | 0.00 | 0 | 0.00 | 0 | 0.00 | 0 | 0.00 | 0 | 0.00 | 0 | 0.00 |
| Gypsy/DIRS1 | 3097794 | 0.13 | 3095699 | 0.12 | 3171926 | 0.13 | 3113243 | 0.12 | 3214000 | 0.13 | 3114315 | 0.12 | 3073639 | 0.12 | 3076605 | 0.13 | 3236726 | 0.12 |
| Retroviral | 107292770 | 4.37 | 108007898 | 4.34 | 108502410 | 4.44 | 107156966 | 4.29 | 111563718 | 4.41 | 109288357 | 4.29 | 106091086 | 4.29 | 103353576 | 4.29 | 114750047 | 4.28 |
| DNA transposons | 65999064 | 2.69 | 66789636 | 2.68 | 66665240 | 2.73 | 66509749 | 2.66 | 69276007 | 2.74 | 66788759 | 2.62 | 65776870 | 2.66 | 64963944 | 2.70 | 66968712 | 2.50 |
| hobo-Activator | 45971432 | 1.87 | 46326066 | 1.86 | 46525219 | 1.90 | 46318265 | 1.85 | 48045163 | 1.90 | 46424586 | 1.82 | 45760606 | 1.85 | 45240517 | 1.88 | 46679477 | 1.74 |
| Tc1-IS630-Pogo | 18698766 | 0.76 | 19126895 | 0.77 | 18796855 | 0.77 | 18857337 | 0.75 | 19866123 | 0.78 | 19025887 | 0.75 | 18695699 | 0.76 | 18403917 | 0.76 | 18937280 | 0.71 |
| En-Spm | 0 | 0.00 | 0 | 0.00 | 0 | 0.00 | 0 | 0.00 | 0 | 0.00 | 0 | 0.00 | 0 | 0.00 | 0 | 0.00 | 0 | 0.00 |
| MULE-MuDR | 57809 | 0.00 | 58170 | 0.00 | 57315 | 0.00 | 57952 | 0.00 | 58292 | 0.00 | 58411 | 0.00 | 59467 | 0.00 | 59262 | 0.00 | 57323 | 0.00 |
| PiggyBac | 194154 | 0.01 | 194028 | 0.01 | 194630 | 0.01 | 196029 | 0.01 | 197634 | 0.01 | 194607 | 0.01 | 194962 | 0.01 | 190214 | 0.01 | 197415 | 0.01 |
| Tourist/Harbinger | 62004 | 0.00 | 62001 | 0.00 | 60455 | 0.00 | 59913 | 0.00 | 62371 | 0.00 | 60525 | 0.00 | 61102 | 0.00 | 60881 | 0.00 | 59316 | 0.00 |
| Other | 0 | 0.00 | 0 | 0.00 | 0 | 0.00 | 0 | 0.00 | 0 | 0.00 | 0 | 0.00 | 0 | 0.00 | 0 | 0.00 | 0 | 0.00 |
| Rolling-circles | 314385 | 0.01 | 316354 | 0.01 | 324062 | 0.01 | 320354 | 0.01 | 323406 | 0.01 | 318931 | 0.01 | 313188 | 0.01 | 308550 | 0.01 | 323233 | 0.01 |
| Unclassified | 509788 | 0.02 | 508240 | 0.02 | 509159 | 0.02 | 512180 | 0.02 | 522125 | 0.02 | 537780 | 0.02 | 505854 | 0.02 | 501985 | 0.02 | 509736 | 0.02 |
| Total interspersed repeats | 960650512 | 39.17 | 985942923 | 39.60 | 967185366 | 39.55 | 947412218 | 37.92 | 962158119 | 38.00 | 970617125 | 38.09 | 940110227 | 38.00 | 864010896 | 35.85 | 1029711934 | 38.40 |
| Small RNA | 189439366 | 7.72 | 214526694 | 8.62 | 187176545 | 7.65 | 190443040 | 7.62 | 201480416 | 7.96 | 191193296 | 7.50 | 187385664 | 7.57 | 173640949 | 7.21 | 198897395 | 7.42 |
| Satellites | 0 | 0.00 | 0 | 0.00 | 1025 | 0.00 | 0 | 0.00 | 0 | 0.00 | 0 | 0.00 | 0 | 0.00 | 0 | 0.00 | 0 | 0.00 |
| Simple repeats | 31363798 | 1.28 | 0 | 0.00 | 31454968 | 1.29 | 31415803 | 1.26 | 32053320 | 1.27 | 31839810 | 1.25 | 80540452 | 3.26 | 27974595 | 1.16 | 37258342 | 1.39 |
| Low complexity | 5677226 | 0.23 | 0 | 0.00 | 5612716 | 0.23 | 5801632 | 0.23 | 5946577 | 0.23 | 6076698 | 0.24 | 5651127 | 0.23 | 5221885 | 0.22 | 10165995 | 0.38 |

**ST8 (Supplementary Table 8). Total number of mean and median heterozygous SNP.**

| Species | Median heterozygosity for species | Mean heterozygosity for species | Sample | Number of heterozygous SNP, abs | Median value, SNP/kbp | Mean value, SNP/kbp |
|---|---|---|---|---|---|---|
| *M. nivalis* | 2.8528 | 2.6433 | 10X_mn | 64980179 | 2.9850 | 2.7659 |
| | | | ERR7198278 | 67839830 | 3.1450 | 2.8877 |
| | | | S8606 | 62845099 | 2.9630 | 2.6751 |
| | | | MNIV | 62592879 | 2.7750 | 2.6643 |
| | | | T100 | 52240846 | 2.3960 | 2.2237 |
| *M. strigidorsa* | 0.6490 | 0.6209 | MSTR1m | 14379184 | 0.6490 | 0.6209 |
| *M. sibirica* | 0.9995 | 0.9783 | T101 | 25972898 | 1.1090 | 1.1074 |
| | | | E19 | 19918151 | 0.8900 | 0.8492 |
| *M. erminea* | 2.1453 | 2.0344 | SRR6963880 | 45058854 | 1.9840 | 1.8656 |
| | | | E26 | 51760030 | 2.2370 | 2.1430 |
| | | | T136 | 50591316 | 2.2150 | 2.0946 |
| *M. richardsonii* | 2.3366 | 2.2215 | SRR6963883 | 59376754 | 2.6010 | 2.4584 |
| | | | SRR6963884 | 53814613 | 2.3000 | 2.2281 |
| | | | SRR6963885 | 60608591 | 2.6850 | 2.5094 |
| | | | SRR6963886 | 55023878 | 2.4620 | 2.2781 |
| | | | SRR6963887 | 70941448 | 3.0290 | 2.9372 |
| | | | SRR6963888 | 30869525 | 1.2890 | 1.2781 |
| | | | SRR6963889 | 44957478 | 1.9900 | 1.8614 |
| *M. nigripes* | 0.0039 | 0.0329 | SB6536 | 804209 | 0.0080 | 0.0339 |
| | | | SB7462 | 830528 | 0.0030 | 0.0350 |
| | | | SB8055 | 482212 | 0.0020 | 0.0203 |
| | | | SRR1508214 | 724873 | 0.0030 | 0.0306 |
| | | | SRR1508215 | 867958 | 0.0030 | 0.0366 |
| | | | SRR1508749 | 796328 | 0.0040 | 0.0336 |
| | | | SRR1508750 | 948903 | 0.0040 | 0.0400 |
| *M. putorius* | 0.3023 | 0.4333 | S33 | 19179667 | 0.7290 | 0.8015 |
| | | | ERR3457930 | 7687401 | 0.8015 | 0.3212 |
| | | | ERR7256386 | 6226441 | 0.0810 | 0.2602 |
| | | | ERR7256388 | 12645518 | 0.4480 | 0.5284 |
| | | | ERR7256390 | 13027247 | 0.4690 | 0.5444 |
| | | | ERR7256392 | 9719128 | 0.3040 | 0.4061 |
| | | | ERR7256412 | 7814090 | 0.1340 | 0.3265 |
| | | | ERR7256413 | 7646523 | 0.1260 | 0.3195 |

| | | | | | | |
|---|---|---|---|---|---|---|
| | | | ERR7260426 | 9369504 | 0.3040 | 0.3915 |
| *M. putorius furo* | 0.0575 | 0.1934 | ERR7256378 | 3889066 | 0.0450 | 0.1645 |
| | | | ERR7256379 | 3939550 | 0.0380 | 0.1667 |
| | | | ERR7256380 | 3964542 | 0.0350 | 0.1677 |
| | | | ERR7256381 | 3721125 | 0.0330 | 0.1574 |
| | | | ERR7256382 | 5504838 | 0.0780 | 0.2329 |
| | | | ERR7256383 | 5016253 | 0.0680 | 0.2122 |
| | | | ERR7256384 | 5373606 | 0.0790 | 0.2273 |
| | | | ERR7256385 | 5830466 | 0.0870 | 0.2466 |
| | | | SRR10395504 | 5398236 | 0.0680 | 0.2284 |
| | | | Solexa | 3074658 | 0.0440 | 0.1301 |
| *M. eversmanii* | 0.5763 | 0.5758 | ERR11751895 | 13800229 | 0.6040 | 0.5755 |
| | | | ERR7198276 | 12851776 | 0.5400 | 0.5359 |
| | | | ERR7198277 | 14772369 | 0.5850 | 0.6160 |
| *N. vison* | 0.8433 | 0.8548 | SRR12564099 | 22307660 | 0.8350 | 0.8480 |
| | | | SRR16676757 | 22551113 | 0.8500 | 0.8573 |
| | | | SRS11183343 | 22600472 | 0.8450 | 0.8592 |

**ST9 (Supplementary Table 9). RoHs content.**

| Species | Genome assembly length, bp. | Sex X chromosome length, bp. | Sample | Sex | Number of RoH * | | | | RoH length * | | | | RoH length, % of all RoH length * | | | RoH fraction in genome, % ** | | | |
|---|---|---|---|---|---|---|---|---|---|---|---|---|---|---|---|---|---|---|---|
| | | | | | all | short | long | ultra long | all | short | long | ultra long | short | long | ultra long | all | short | long | ultra long |
| *M. sibirica* | 2532161059 | 112012753 | E19 | Male | 816 | 696 | 120 | 0 | 492830000 | 216940000 | 275890000 | 0 | 44.02 | 55.98 | 0 | 20.36 | 8.96 | 11.4 | 0 |
| | | | T101 | Male | 458 | 449 | 9 | 0 | 113020000 | 98830000 | 14190000 | 0 | 87.44 | 12.56 | 0 | 4.67 | 04.08 | 0.59 | 0 |
| *M. nivalis* | 2452646332 | 125416964 | 10X_mn | Male | 106 | 90 | 15 | 1 | 81450000 | 20680000 | 36710000 | 24060000 | 25.39 | 45.07 | 29.54 | 3.5 | 0.89 | 1.58 | 01.03 |
| | | | ERR7198278 | Male | 361 | 344 | 16 | 1 | 127000000 | 83590000 | 32580000 | 10830000 | 65.82 | 25.65 | 8.53 | 5.46 | 3.59 | 1.4 | 0.47 |
| | | | S8606 | Male | 205 | 166 | 36 | 3 | 131000000 | 46770000 | 97840000 | 67080000 | 35.7 | 74.69 | 51.21 | 5.63 | 02.01 | 4.2 | 2.88 |
| | | | MNIV | Male | 55 | 51 | 4 | 0 | 29280000 | 11030000 | 18250000 | 0 | 37.67 | 62.33 | 0 | 1.26 | 0.47 | 0.78 | 0 |
| | | | T100 | Female | 174 | 156 | 11 | 7 | 269860000 | 36110000 | 42710000 | 191040000 | 13.38 | 15.83 | 70.79 | 11.6 | 1.55 | 1.84 | 8.21 |
| *M. strigidorsa* | 2489756210 | 73834643 | MSTR1m | Male | 1062 | 953 | 106 | 3 | 570560000 | 280130000 | 246690000 | 43740000 | 49.1 | 43.24 | 7.67 | 23.62 | 11.6 | 10.21 | 1.81 |
| *M. erminea* | 2445217270 | 130149454 | SRR6963880 | Male | 364 | 348 | 16 | 0 | 113990000 | 80000000 | 33990000 | 0 | 70.18 | 29.82 | 0 | 4.92 | 3.46 | 1.47 | 0 |
| | | | E26 | Male | 150 | 143 | 7 | 0 | 51370000 | 33090000 | 18280000 | 0 | 64.42 | 35.58 | 0 | 2.22 | 1.43 | 0.79 | 0 |
| | | | T136 | Male | 148 | 134 | 13 | 1 | 84490000 | 28110000 | 24890000 | 31490000 | 33.27 | 29.46 | 37.27 | 3.65 | 1.21 | 01.08 | 1.36 |
| *M. richardsonii* | 2445217270 | 130149454 | SRR6963883 | Male | 166 | 158 | 8 | 0 | 51180000 | 36960000 | 14220000 | 0 | 72.22 | 27.78 | 0 | 2.21 | 1.6 | 0.61 | 0 |
| | | | SRR6963884 | Female | 212 | 203 | 9 | 0 | 64110000 | 45460000 | 18650000 | 0 | 70.91 | 29.09 | 0 | 2.77 | 1.96 | 0.81 | 0 |
| | | | SRR6963885 | Male | 192 | 173 | 18 | 1 | 106020000 | 43940000 | 47930000 | 14150000 | 41.45 | 45.21 | 13.35 | 4.58 | 1.9 | 02.07 | 0.61 |
| | | | SRR6963886 | Female | 250 | 203 | 40 | 7 | 277990000 | 50890000 | 129740000 | 97360000 | 18.31 | 46.67 | 35.02 | 12.01 | 2.2 | 5.6 | 4.21 |
| | | | SRR6963887 | Female | 143 | 140 | 3 | 0 | 40000000 | 33290000 | 6710000 | 0 | 83.23 | 16.78 | 0 | 1.73 | 1.44 | 0.29 | 0 |
| | | | SRR6963888 | Female | 656 | 626 | 30 | 0 | 214460000 | 149290000 | 65170000 | 0 | 69.61 | 30.39 | 0 | 9.26 | 6.45 | 2.82 | 0 |
| | | | SRR6963889 | Female | 544 | 453 | 87 | 4 | 403850000 | 143220000 | 188090000 | 72540000 | 35.46 | 46.57 | 17.96 | 17.44 | 6.19 | 8.12 | 3.13 |
| *M. nigripes* | 2498707582 | 124252308 | SB6536 | Male | 355 | 191 | 119 | 45 | 2089080000 | 74330000 | 384920000 | 1629830000 | 3.56 | 18.43 | 78.02 | 87.98 | 3.13 | 16.21 | 68.64 |
| | | | SB7462 | Female | 367 | 202 | 110 | 55 | 2082080000 | 80050000 | 350310000 | 1651720000 | 3.84 | 16.83 | 79.33 | 87.69 | 3.37 | 14.75 | 69.56 |
| | | | SB8055 | Male | 199 | 99 | 51 | 49 | 2176460000 | 35590000 | 142710000 | 1998160000 | 1.64 | 6.56 | 91.81 | 91.66 | 1.5 | 06.01 | 84.15 |
| | | | SRR1508214 | Male | 313 | 176 | 85 | 52 | 2108950000 | 67420000 | 261760000 | 1779770000 | 3.2 | 12.41 | 84.39 | 88.82 | 2.84 | 11.02 | 74.95 |
| | | | SRR1508215 | Male | 409 | 227 | 131 | 51 | 2069990000 | 87170000 | 371750000 | 1611070000 | 4.21 | 17.96 | 77.83 | 87.18 | 3.67 | 15.66 | 67.85 |
| | | | SRR1508749 | Male | 357 | 217 | 100 | 40 | 2092720000 | 81900000 | 279650000 | 1731170000 | 3.91 | 13.36 | 82.72 | 88.13 | 3.45 | 11.78 | 72.91 |
| | | | SRR1508750 | Female | 414 | 235 | 132 | 47 | 2061830000 | 91340000 | 356820000 | 1613670000 | 4.43 | 17.31 | 78.26 | 86.83 | 3.85 | 15.03 | 67.97 |
| *M. putorius* | 2473963644 | 126329207 | S33 | Male | 1315 | 1276 | 38 | 1 | 371750000 | 299030000 | 62340000 | 10380000 | 80.44 | 16.77 | 2.79 | 15.84 | 12.74 | 2.66 | 0.44 |
| | | | ERR3457930 | Male | 1304 | 993 | 304 | 7 | 1170720000 | 307330000 | 745490000 | 117900000 | 26.25 | 63.68 | 10.06 | 49.87 | 13.09 | 31.75 | 05.02 |
| | | | ERR7256386 | Male | 1192 | 1054 | 121 | 17 | 1391740000 | 330770000 | 224590000 | 836380000 | 23.77 | 16.14 | 60.1 | 59.28 | 14.09 | 9.57 | 35.63 |
| | | | ERR7256388 | Male | 1745 | 1620 | 124 | 1 | 714730000 | 465370000 | 231770000 | 17590000 | 65.11 | 32.43 | 2.46 | 30.44 | 19.82 | 9.87 | 0.75 |
| | | | ERR7256390 | Female | 1727 | 1599 | 127 | 1 | 717340000 | 476160000 | 230240000 | 10940000 | 66.38 | 32.1 | 10.53 | 30.56 | 20.28 | 9.81 | 0.47 |
| | | | ERR7256392 | Male | 1814 | 1624 | 186 | 4 | 920970000 | 506120000 | 336890000 | 77960000 | 54.96 | 36.58 | 8.46 | 39.23 | 21.56 | 14.39 | 3.32 |
| | | | ERR7256412 | Female | 1384 | 1065 | 312 | 7 | 1238470000 | 336520000 | 773380000 | 128570000 | 27.17 | 62.45 | 10.38 | 52.75 | 14.33 | 32.94 | 5.48 |
| | | | ERR7256413 | Male | 1304 | 977 | 320 | 7 | 1182670000 | 304810000 | 758750000 | 119110000 | 25.77 | 64.16 | 10.07 | 50.38 | 12.98 | 32.32 | 05.07 |
| | | | ERR7260426 | Male | 1792 | 1582 | 208 | 2 | 938200000 | 506090000 | 367560000 | 64550000 | 53.94 | 39.18 | 6.88 | 39.96 | 21.56 | 15.66 | 2.75 |
| *M. putorius furo* | 2409982543 | 123993920 | ERR7256378 | Female | 865 | 489 | 351 | 25 | 1756730000 | 174960000 | 1131380000 | 450390000 | 9.96 | 64.4 | 25.64 | 76.85 | 7.65 | 49.49 | 19.7 |
| | | | ERR7256379 | Male | 833 | 450 | 356 | 28 | 1778290000 | 168450000 | 1135010000 | 474830000 | 9.47 | 63.83 | 26.7 | 77.79 | 7.37 | 49.65 | 20.77 |
| | | | ERR7256380 | Female | 773 | 389 | 358 | 26 | 1792790000 | 156850000 | 1139610000 | 496330000 | 8.75 | 63.57 | 27.68 | 78.43 | 6.86 | 49.85 | 21.71 |
| | | | ERR7256381 | Female | 797 | 429 | 337 | 31 | 1804020000 | 147730000 | 1034980000 | 621310000 | 8.19 | 57.37 | 34.44 | 78.92 | 6.46 | 45.27 | 27.18 |
| | | | ERR7256382 | Male | 1037 | 680 | 333 | 25 | 1511500000 | 215450000 | 885910000 | 410140000 | 14.25 | 58.61 | 27.13 | 66.12 | 9.42 | 38.75 | 17.94 |
| | | | ERR7256383 | Male | 1018 | 656 | 344 | 19 | 1587140000 | 225450000 | 1074630000 | 287060000 | 14.2 | 67.71 | 18.09 | 69.43 | 9.86 | 47.01 | 12.56 |
| | | | ERR7256384 | Female | 1029 | 690 | 319 | 20 | 1524230000 | 222540000 | 945530000 | 356160000 | 14.6 | 62.03 | 23.37 | 66.68 | 9.73 | 41.36 | 15.58 |
| | | | ERR7256385 | Female | 1057 | 724 | 318 | 15 | 1483670000 | 238550000 | 924410000 | 320710000 | 16.08 | 62.31 | 21.62 | 64.9 | 10.44 | 40.44 | 14.03 |
| | | | SRR10395504 | Male | 944 | 584 | 338 | 23 | 1591520000 | 204070000 | 1026350000 | 361100000 | 12.82 | 64.49 | 22.69 | 69.62 | 8.93 | 44.9 | 15.8 |
| | | | Solexa | Male | 855 | 472 | 354 | 30 | 1875760000 | 180460000 | 1236410000 | 458890000 | 9.62 | 65.92 | 24.46 | 82.05 | 7.89 | 54.09 | 20.07 |
| *M. eversmanii* | 2547922575 | 108794627 | ERR11751895 | Male | 335 | 301 | 23 | 11 | 572570000 | 65990000 | 60560000 | 446020000 | 11.53 | 10.58 | 77.9 | 23.47 | 2.71 | 2.48 | 18.29 |
| | | | ERR7198276 | Male | 631 | 584 | 34 | 13 | 679130000 | 134000000 | 75880000 | 469250000 | 19.73 | 11.17 | 69.1 | 27.84 | 5.49 | 3.11 | 19.24 |
| | | | ERR7198277 | Male | 591 | 542 | 40 | 9 | 616290000 | 128640000 | 134230000 | 353420000 | 20.87 | 21.78 | 57.35 | 25.27 | 5.27 | 5.5 | 14.49 |
| | | | SRR12564099 | Male | 373 | 306 | 54 | 13 | 534810000 | 77650000 | 164690000 | 292470000 | 14.52 | 30.79 | 54.69 | 20.98 | 03.05 | 6.46 | 11.47 |

| | | | | | | | | | | | | | | | | | | |
|---|---|---|---|---|---|---|---|---|---|---|---|---|---|---|---|---|---|---|
| *N. vison* | 2681215271 | 131682864 | SRR16676757 | Male | 389 | 320 | 61 | 8 | 557530000 | 91360000 | 207280000 | 258890000 | 16.39 | 37.18 | 46.44 | 21.87 | 3.58 | 8.13 | 10.15 |
| | | | SRS11183343 | Male | 462 | 384 | 70 | 8 | 471220000 | 108630000 | 196240000 | 166350000 | 23.05 | 41.65 | 35.3 | 18.48 | 4.26 | 7.7 | 6.52 |

* Short RoH (< 1 Mbp), long RoH (>=1 Mbp < 10 Mbp) and ultra long RoH (>= 10 Mbp).

** The proportion of RoH in the genome was calculated as the percentage of the total length of all RoH segments relative to the genome assembly length used (excluding the length of the X chromosome).

**ST10 (Supplementary Table 10). Summary of support metrics for each internal branch in the ASTRAL species tree.**

The table reports quartet support values (q1, q2, q3), representing the proportion of gene tree quartets supporting the main topology and two alternatives; counts of supporting quartet trees across all gene trees (f1, f2, f3); local posterior probabilities for each topology (pp1, pp2, pp3); total number of informative quartets (QC); and the effective number of genes contributing to each branch (EN).

| Node | q1 | q2 | q3 | f1 | f2 | f3 | pp1 | pp2 | pp3 | QC | EN |
|---|---|---|---|---|---|---|---|---|---|---|---|
| 1 | - | - | - | - | - | - | - | - | - | - | - |
| 2 | 0,6172965154 | 0,1990426448 | 0,1836608398 | 2508,833333 | 808,9545455 | 746,4393939 | 1 | 0 | 0 | 132 | 4064,227273 |
| 3 | 0,7833379007 | 0,1023194875 | 0,1143426118 | 3161,202257 | 412,9157986 | 461,4357639 | 1 | 0 | 0 | 1152 | 4035,553819 |
| 4 | 0,995435386 | 0,00243111551 | 0,002133498522 | 6392,066667 | 15,61111111 | 13,7 | 1 | 0 | 0 | 90 | 6421,377778 |
| 5 | 0,8338170287 | 0,08522076488 | 0,08096220645 | 3316,442578 | 338,9589844 | 322,0208984 | 1 | 0 | 0 | 5120 | 3977,422461 |
| 6 | 0,3631639197 | 0,3171464623 | 0,3196896179 | 1213,804348 | 1060 | 1068,5 | 0,9995170594 | 0,0002266176964 | 0,0002563229334 | 46 | 3342,304348 |
| 7 | 0,8730294312 | 0,05895889459 | 0,0680116742 | 3248,065705 | 219,3538462 | 253,0342949 | 1 | 0 | 0 | 3120 | 3720,453846 |
| 8 | 0,9688150665 | 0,01682027539 | 0,01436465816 | 5004,395926 | 86,88481481 | 74,20037037 | 1 | 0 | 0 | 2700 | 5165,481111 |
| 9 | 0,4774700217 | 0,3033953406 | 0,2191346377 | 1084,25355 | 688,959432 | 497,6176471 | 1 | 2,39E-45 | 0 | 986 | 2270,830629 |
| 10 | 0,9802119204 | 0,01008131611 | 0,009706763443 | 5491,967949 | 56,48397436 | 54,38541667 | 1 | 0 | 0 | 1248 | 5602,83734 |
| 11 | 0,8612563329 | 0,05046481261 | 0,08827885446 | 2157,285366 | 126,404878 | 221,1219512 | 1 | 0 | 0 | 410 | 2504,812195 |
| 12 | 0,683912005 | 0,1361463234 | 0,1799416716 | 1416,556911 | 281,9939024 | 372,7052846 | 1 | 0 | 0 | 492 | 2071,256098 |
| 13 | 0,7278843628 | 0,1008363464 | 0,1712792908 | 1892,801385 | 262,2163435 | 445,3972299 | 1 | 0 | 0 | 3610 | 2600,414958 |
| 14 | 0,9772006868 | 0,01024710572 | 0,01255220744 | 4529,396078 | 47,49607843 | 58,18039216 | 1 | 0 | 0 | 510 | 4635,072549 |
| 15 | 0,3677525167 | 0,3292829289 | 0,3029645543 | 956,7 | 856,6222222 | 788,1555556 | 0,9994906621 | 0,000389456794 | 0,0001198810869 | 90 | 2601,477778 |
| 16 | 0,4054516412 | 0,301499459 | 0,2930488999 | 855,2826087 | 636 | 618,173913 | 1 | 4,63E-12 | 3,76E-12 | 46 | 2109,456522 |
| 17 | 0,3674589057 | 0,3157983214 | 0,316742773 | 683,0852273 | 587,0511364 | 588,8068182 | 0,9967402194 | 0,001599535472 | 0,001660245097 | 176 | 1858,943182 |
| 18 | 0,8348605274 | 0,04974456692 | 0,1153949057 | 2742,851333 | 163,4308333 | 379,1185 | 1 | 0 | 0 | 18000 | 3285,400667 |
| 19 | 0,4853740443 | 0,2787431644 | 0,2358827914 | 1259,978261 | 723,5869565 | 612,326087 | 1 | 2,23E-57 | 0 | 46 | 2595,891304 |
| 20 | 0,341512605 | 0,3338463807 | 0,3246410143 | 643,4666667 | 629,0222222 | 611,6777778 | 0,5590409629 | 0,2825034994 | 0,1584555377 | 90 | 1884,166667 |
| 21 | 0,3494840815 | 0,3141252903 | 0,3363906282 | 660,4666667 | 593,6444444 | 635,7222222 | 0,7743272067 | 0,05046761777 | 0,1752051755 | 90 | 1889,833333 |
| 22 | 0,8867970971 | 0,08261496985 | 0,03058793303 | 2982,741667 | 277,8754167 | 102,8825 | 1 | 0 | 0 | 2400 | 3363,499583 |
| 23 | 0,7850795415 | 0,1308734149 | 0,08404704357 | 2081,334211 | 346,9601504 | 222,8181704 | 1 | 0 | 0 | 7980 | 2651,112531 |
| 24 | 0,922828795 | 0,04107679526 | 0,03609440974 | 1340,602932 | 59,67268519 | 52,43472222 | 1 | 0 | 0 | 6480 | 1452,71034 |
| 25 | - | - | - | - | - | - | - | - | - | - | - |
| 26 | 0,9993037566 | 0,0002640630559 | 0,0004321803712 | 3353,074074 | 0,886039886 | 1,45014245 | 1 | 0 | 0 | 702 | 3355,410256 |
| 27 | 0,788650315 | 0,1276427977 | 0,08370688726 | 1500,272894 | 242,8186813 | 159,2380952 | 1 | 0 | 0 | 546 | 1902,32967 |
| 28 | 0,4373179772 | 0,3231665343 | 0,2395154885 | 211,7628205 | 156,4871795 | 115,9807692 | 0,9999946889 | 4,27E-06 | 1,04E-06 | 780 | 484,2307692 |
| 29 | 0,3760208631 | 0,2940974574 | 0,3298816795 | 174,7891738 | 136,7079772 | 153,3418803 | 0,9109221299 | 0,02586187902 | 0,06321599103 | 702 | 464,8390313 |
| 30 | 0,6211275321 | 0,1643847756 | 0,2144876923 | 514,0016667 | 136,0333333 | 177,495 | 1 | 0 | 5,38E-64 | 600 | 827,53 |
| 31 | 0,3931996982 | 0,3381291087 | 0,268671193 | 69,4952381 | 59,76190476 | 47,48571429 | 0,8393994788 | 0,1216980197 | 0,03890250158 | 210 | 176,7428571 |
| 32 | 0,5131460415 | 0,3826283703 | 0,1042255882 | 1064,956522 | 794,0869565 | 216,3043478 | 1 | 1,60E-57 | 0 | 46 | 2075,347826 |
| 33 | 0,4742509064 | 0,2582398013 | 0,2675092923 | 201,2790698 | 109,6007752 | 113,5348837 | 0,9999999965 | 1,66E-09 | 1,86E-09 | 258 | 424,4147287 |

| | | | | | | | | | | | |
|---|---|---|---|---|---|---|---|---|---|---|---|
| 34 | 0,3390059308 | 0,3224086041 | 0,3385854652 | 170,2111111 | 161,8777778 | 170 | 0,3951421774 | 0,2167581886 | 0,388099634 | 90 | 502,0888889 |
| 35 | 0,5146079196 | 0,284159742 | 0,2012323384 | 313,9254743 | 173,3455285 | 122,7574526 | 1 | 5,25E-20 | 2,14E-20 | 738 | 610,0284553 |
| 36 | 0,5340653434 | 0,3257387571 | 0,1401958994 | 378,0804878 | 230,6 | 99,24878049 | 1 | 2,92E-27 | 0 | 410 | 707,9292683 |
| 37 | 0,3424995654 | 0,3249782722 | 0,3325221624 | 57,27906977 | 54,34883721 | 55,61046512 | 0,4043012117 | 0,2745897391 | 0,3211090492 | 172 | 167,2383721 |
| 38 | 0,3661458077 | 0,3274839388 | 0,3063702535 | 126,6136364 | 113,2443182 | 105,9431818 | 0,7546886428 | 0,1531916724 | 0,09211968473 | 176 | 345,8011364 |
| 39 | 0,3733525713 | 0,3232836592 | 0,3033637695 | 131,3409091 | 113,7272727 | 106,719697 | 0,8441078278 | 0,09540600113 | 0,06048617103 | 132 | 351,7878788 |
| 40 | 0,3516047602 | 0,3282445807 | 0,3201506591 | 190,7608696 | 178,0869565 | 173,6956522 | 0,6262531269 | 0,2113182863 | 0,1624285869 | 46 | 542,5434783 |
| 41 | 0,4414893617 | 0,3064058568 | 0,2521047815 | 219,2897727 | 152,1931818 | 125,2215909 | 0,9999988505 | 8,06E-07 | 3,44E-07 | 176 | 496,7045455 |
| 42 | 0,3647890724 | 0,3035909523 | 0,3316199753 | 194,6287879 | 161,9772727 | 176,9318182 | 0,8179421693 | 0,05617859495 | 0,1258792357 | 132 | 533,5378788 |
| 43 | 0,8971670346 | 0,0620729528 | 0,04076001261 | 742,1521739 | 51,34782609 | 33,7173913 | 1 | 0 | 0 | 46 | 827,2173913 |
| 44 | 0,3637207425 | 0,3212671509 | 0,3150121065 | 68,28030303 | 60,31060606 | 59,13636364 | 0,6212703086 | 0,2007238163 | 0,1780058751 | 132 | 187,7272727 |
| 45 | 0,350090854 | 0,3412477286 | 0,3086614173 | 125,6521739 | 122,4782609 | 110,7826087 | 0,5054651935 | 0,3535787936 | 0,1409560129 | 46 | 358,9130435 |
| 46 | 0,3753669725 | 0,2570642202 | 0,3675688073 | 177,8913043 | 121,826087 | 174,1956522 | 0,6506244263 | 0,01093578213 | 0,3384397915 | 46 | 473,9130435 |
| 47 | 0,3719411648 | 0,3414682409 | 0,2865905943 | 62,65555556 | 57,52222222 | 48,27777778 | 0,6431553864 | 0,2597227801 | 0,09712183355 | 90 | 168,4555556 |
| 48 | 0,3819610978 | 0,2796429523 | 0,3383959499 | 62,32608696 | 45,63043478 | 55,2173913 | 0,7346065868 | 0,07209603873 | 0,1932973745 | 46 | 163,173913 |

**ST11 (Supplementary Table 11). List of mitochondrial genome assembly IDs used in the phylogenetic analyses.**

| Species | Accession ID | Assembly publication |
|---|---|---|
| *M. eversmanii* | NC_028013 | - |
| | PQ821906 (ERR11751895) | - |
| | BK068807 (ERR7198276) | (Etherington et al. 2022) |
| | BK068808 (ERR7198277) | |
| *M. nudipes* | MH464792 | - |
| *M. nigripes* | NC_024942 | (Zhao et al. 2016) |
| *M. altaica* | NC_021751 | (Huang et al. 2014) |
| *M. strigidorsa* | CM107193.1 (MSTR1m) | This study |
| *M. putorius* | BK069841 (ERR3457930) | (Etherington et al. 2020) |
| | BK069847 (ERR7256413) | (Etherington et al. 2022) |
| | BK069845 (ERR7256392) | |
| | BK069844 (ERR7256390) | |
| | BK069846 (ERR7256412) | |
| | BK069848 (ERR7260426) | |
| | BK069842 (ERR7256386) | |
| | BK069843 (ERR7256388) | |
| | PQ246112 (S33) | This study |
| | NC_020638 | (Yu et al. 2011) |
| *M. kathiah* | NC_023210 | |
| *N. frenata* | NC_020640 | |
| *N. vison* | NC_020641 | |
| | KM488625 | (Sun et al. 2016) |
| | KU146454 | (Hua and Xu 2016) |
| | MT410953 | - |
| *M. putorius furo* | KT693383 | (Emami-Khoyi et al. 2016) |
| | KT693382 | |
| | CM032670 (10X_mn) | (Miranda et al. 2021) |

| | | |
|---|---|---|
| *M. nivalis* | PQ246109 (MNIV) | This study |
| | MW257229 | (Hassanin et al. 2021) |
| | KT901457 | - |
| | MF459691 | (Lim et al. 2017) |
| | BK069849 (ERR7198278) | (Etherington et al. 2022) |
| | PQ246110 (S8606) | This study |
| | PQ246111 (T100) | This study |
| | NC_020639 | (Yu et al. 2011) |
| *M. lutreola* | CM059646 | (Skorupski 2022) |
| | NC_056132.1 | |
| | MW197425 | |
| | MW197426 | |
| | MT304869 | |
| | MW197423 | |
| | MW197424 | |
| *M. itatsi* | NC_034330 | |
| | AP017400 | |
| | AP017401 | |
| | AP017388 | |
| | AP017402 | |
| | AP017403 | |
| | AP017404 | |
| | AP017405 | |
| | AP017389 | |
| | AP017390 | |
| | AP017406 | |
| | AP017407 | |
| | AP017391 | |
| | AP017408 | |
| | AP017409 | |

| | | |
|---|---|---|
| | AP017410 | (Shalabi et al. 2017) |
| | AP017411 | |
| | AP017392 | |
| | AP017412 | |
| *M. sibirica* | AP017393 | |
| | AP017413 | |
| | AP017414 | |
| | AP017415 | |
| | AP017416 | |
| | AP017394 | |
| | AP017417 | |
| | AP017395 | |
| | AP017418 | |
| | AP017396 | |
| | AP017397 | |
| | AP017419 | |
| | AP017420 | |
| | AP017421 | |
| | MN206976 | (Gao et al. 2020) |
| | MW625812 | - |
| | (CM107191.1) E19 | This study |
| | MN264435 | (Yu et al. 2019) |
| | MH818224 | - |
| | PQ246113 (T101) | This study |
| | NC_020637 | (Yu et al. 2011) |
| | MW257230 | (Hassanin et al. 2021) |
| | MT584107 | - |
| | CM020617 | - |
| | NC_025516 | - |
| | PQ246107 (E26) | This study |

| | | |
|---|---|---|
| *M. erminea* | PQ246108 (T136) | This study |
| | MK603870 | |
| | MK603871 | |
| | MK603873 | |
| | MK603904 (SRR6963880) | |
| | MK603896 | |
| | MK603875 | |
| | MK603876 | |
| | MK603892 | |
| *M. haidarum* | MK603899 | |
| | MK603898 | |
| | MK603004 | |
| | MK603005 | |
| | MK603006 | |
| | MK603007 | |
| | MK603008 | |
| | MK603009 | |
| | MK603010 | |
| | MK603011 | |
| | MK603012 | |
| | MK603014 | |
| | MK603003 | |
| | MK603013 | |
| | MK603900 (SRR6963882) | |
| | MK603879 | |
| | MK603880 | |
| | MK603881 | |
| | MK603882 | |
| | MK603908 | |
| | MK603909 | (Colella et al. 2021) |

| | | |
|---|---|---|
| | MK603910 | (Colella et al. 2021) |
| | MK603913 | |
| | MK603897 | |
| | MK603894 | |
| | MK603906 | |
| | MK603907 | |
| | MK603886 | |
| | MK603887 | |
| | MK603911 | |
| | MK603912 | |
| | MK603895 (SRR6963885, SRR6978696) | |
| | MK603890 | |
| *M. richardsonii* | MK603915 | |
| | MK603891 | |
| | MK603893 | |
| | MK603905 | |
| | MK603902 | |
| | MK603903 | |
| | MK603883 (SRR6963886) | |
| | MK603884 (SRR6963889) | |
| | MK603874 | |
| | MK603877 | |
| | MK603878 | |
| | MK603885 (SRR6963883) | |
| | MK603889 | |
| | MK603869 (SRR6963884) | |
| | MK603872 | |
| | MK603888 | |
| | MK603914 | |
| | BK069851 (SRR6963887) | (Colella et al. 2018) |

| | | |
|---|---|---|
| | BK069850 (SRR6963888) | (Colella et al. 2018) |
| *Martes foina* | NC_020643 | (Yu et al. 2011) |

**ST12 (Supplementary Table 12). List of species used in the synteny analysis.**

| Latin name | Common name | Family | 2n | Assembly name | Source | Assembly publication |
|---|---|---|---|---|---|---|
| *Homo sapiens* | Human | Hominidae | 46 | GRCh38.p14 | NCBI | (Schneider et al. 2017) |
| *Canis familiaris* | Dog | Canidae | 78 | UU_Cfam_GSD_1.0 (CanFam4) | NCBI | (Hoeppner et al. 2014) |
| *Enhydra lutris* | Sea otter | Mustelidae | 38 | ASM228890v2_HiC | DNAZoo | (Peng et al. 2014; Dudchenko et al. 2017; Dudchenko et al. 2018) |
| *Martes foina* | Stone marten | Mustelidae | 38 | mfoi.min_150.pseudohap2.1_HiC | DNAZoo | (Dudchenko et al. 2017; Dudchenko et al. 2018) |
| *Neogale vison* | American mink | Mustelidae | 30 | ASM_NN_V1 | NCBI | (Karimi et al. 2022) |
| *Mustela nigripes* | Black-footed ferret | Mustelidae | 38 | musNig1_HiC | DNAZoo | (Kliver et al. 2023) |
| *Mustela nivalis* | Least weasel | Mustelidae | 42 | MusNiv_Pri1.0.no_dups.0.9_HiC | This study | This study |
| *Mustela strigidorsa* | Black-striped weasel | Mustelidae | 44 | MSTR1m | This study | This study |
| *Mustela erminea* | Ermine/Stoat | Mustelidae | 44 | mMusErm1.Pri | NCBI | - |
| *Mustela lutreola* | European mink | Mustelidae | 38 | mMusLut2.pri | NCBI | (Skorupski et al. 2023) |
| *Mustela putorius furo* | Domestic ferret | Mustelidae | 40 | MusPutFur1.0_HiC | DNAZoo | (Peng et al. 2014; Dudchenko et al. 2017; Dudchenko et al. 2018) |

**ST13 (Supplementary Table 13). The main fur game animal resources in the Russian Federation for the years 2003-2023.**

Data are based on public reports of the Ministry of Natural Resources and Environment of the Russian Federation and the Federal Research Center for Hunting Development (FGU Centrokhotkontrol 2007; Minprirody of Russia 2019; Minprirody of Russia 2021; Minprirody of Russia 2024). Differences indicate discrepancies in the datasets.

| **Species** | **Number for the year, thousands** | | | | | | | | | | | | | | | | | | | | |
|---|---|---|---|---|---|---|---|---|---|---|---|---|---|---|---|---|---|---|---|---|---|
| | **2003** | **2004** | **2005** | **2006** | **2007** | **2008** | **2009** | **2010** | **2011** | **2012** | **2013** | **2014** | **2015** | **2016** | **2017** | **2018** | **2019** | **2020** | **2021** | **2022** | **2023** |
| *Mustela erminea* | 997.4 | 945.0 or 816.9 | 881.4 or 755.2 | 1015.5 or 951.7 | 958.4 or 880.1 | 686.4 | 670.7 or 670.8 | 695.5 | 648.6 | 584.1 | 545.2 | 423.8 | 409.4 | 407.3 | 405.5 | 425.3 | 387.1 | 397.0 | 384.2 | 389.7 | 430.4 |
| *Mustela sibirica* | 197.7 | 188.5 or 173.3 | 171.4 or 157.1 | 169.1 or 167.3 | 159.4 or 153.8 | 136.9 or 136.8 | 128.5 or 128.6 | 150.8 | 154.8 | 149.7 | 129.0 | 116.7 | 108.4 | 122.7 | 121.4 | 120.9 | 104.5 | 104.1 | 113.9 | 113.2 | 111.0 |
| *Mustela eversmanii & Mustela putorius* | 90.6 | 86.6 | 81.9 | 80.6 or 80.7 | 82.0 or 82.1 | 70.3 or 74.1 | 70.0 | 61.5 | 64.5 | 68.3 | 58.8 | 56.7 | 53.6 | 55.1 | 50.6 | 47.2 | 50.3 | 47.1 | 44.0 | 51.4 | 48.3 |
| *Martes martes & Martes foina* | 218.6 | 220.3 | 233.4 | 234.8 | 220.8 or 222.3 | 243.9 or 242.7 | 247.9 | 226.1 or 226.8 | 219.4 | 238.3 | 236.9 | 213.1 | 202.7 | 204.5 | 229.0 | 230.0 | 232.8 | 226.5 | 229.1 | 245.5 | 246.4 |
| *Gulo gulo* | 20.41 | 24.01 or 22.5 | 24.51 or 22.5 | 22.11 or 21.5 | 23.39 or 22.8 | 20.5 or 20.1 | 19.5 | 19.7 | 18.6 | 19.7 | 17.9 | 14.9 | 13.5 | 14.5 | 15.5 | 17.9 | 16.9 | 18.6 | 17.5 | 18.6 | 20.5 |
| *Martes zibellina* | 1095.1 | 1135.4 | 1120.1 | 1259.3 | 1432.0 or 1407.5 | 1459.5 | 1481.9 | 1163.8 | 1224.5 | 1288.9 | 1346.3 | 1286.7 | 1309.7 | 1402.7 | 1497.1 | 1574.8 | 1436.4 | 1546.0 | 1605.2 | 1670.4 | 1675.4 |
| *Lutra lutra* | 73.8 | 75.12 | 77.43 | 77.09 or 74.2 | 76.74 or 73.9 | 76.5 or 75.5 | 79.8 | 80.0 or 77.7 | 80.0 | 101.3 | 103.9 | 75.1 | 85.2 | 81.5 | 82.9 | 101.5 | 102.0 | 108.2 | 111.9 | 118.4 | 109.2 |

**ST14 (Supplementary Table 14). Sample information.**

| Species | Data type* | Sample ID | SRA ID | Sex | Origin |
|---|---|---|---|---|---|
| *M. nivalis* | PE | MNIV | SRR30238154 | male | Russia, Novosibirsk |
| | PE | T100 | SRR30238153 | female | Russia, Far East |
| | 10X | 10X_mn | SRR13788917<br>SRR13788918 | male | Poland, Grzymały-Sierzputy Stare, along the main road Ostrołęka—Łomża (road-killed) |
| | PE | ERR7198278 | ERR7198278 | male | France, La Couvertoirade, Averyron |
| | PE | S8606 | SRR30238155 | male | Germany, Berlin, Lichtenberg, Karlshorst, Beerfelder Str. |
| *M. strigidorsa* | PE | MSTR1m | SRR34068022 | male | Vietnam, Quang Nam |
| *M. sibirica* | PE | E19 | SRR30226579 | male | Russia, Republic of Sakha |
| | PE | T101 | SRR30238152 | male | Russia, Far East, Primorsky krai, Sikhote-Alin Nature Reserve, river Western Kema |
| *M. erminea* | PE | T136 | SRR30238151 | male | Russia, Republic of Sakha, Tomponskii region, village Khandyga |
| | PE | E26 | SRR30238150 | male | Russia, Republic of Sakha |
| | PE | SRR6963880 | SRR6963880 | male | Mongolia, Bayan-Ölgii Province |
| *M. richardsonii* | PE | SRR6963883 | SRR6963883 | male | Canada, Southern British Columbia |
| | PE | SRR6963884 | SRR6963884 | female | USA, Vermont |
| | PE | SRR6963885<br>SRR6978696 | SRR6963885<br>SRR6978696 | male | USA, Alaska, Revillagigedo Island |
| | PE | SRR6963886 | SRR6963886 | female | USA, Alaska, Annette Island |
| | PE | SRR6963887 | SRR6963887 | female | Canada, Southern Yukon Territory |
| | PE | SRR6963888 | SRR6963888 | female | USA, New Mexico |
| | PE | SRR6963889 | SRR6963889 | female | USA, Alaska, Kupreanof Island |
| *M. nigripes* | PE | SB6573 | SRR1508214 | male | USA, Wyoming |
| | PE | SB6815 | SRR1508215 | male | USA, Wyoming |
| | PE | SB10 | SRR1508750 | female | USA, Wyoming |
| | PE | W2094(SB2) | SRR1508749 | male | USA, Wyoming |
| | PE | SB7462 | SRR11941224 | female | Captive population |
| | PE | SB8055 | SRR11941223 | male | Captive population |

| | | | | | |
|---|---|---|---|---|---|
| | PE | SB6536 | SRR12036677<br>SRR12036676<br>SRR11940030 | male | Captive population |
| *M. putorius* | PE | ERR3457930 | ERR3457930 | male | United Kingdom, Community of Magor with Undy |
| | PE | ERR7256386 | ERR7256386 | male | Spain, Vera de Bidasoa, Navarre |
| | PE | ERR7256388 | ERR7256388 | male | Austria, Feldbach, Styria |
| | PE | ERR7256390 | ERR7256390 | female | Austria, Wolfsberg, Styria |
| | PE | ERR7256392 | ERR7256392 | male | France, Audon |
| | PE | ERR7256412 | ERR7256412 | female | United Kingdom, Bartestree |
| | PE | ERR7256413 | ERR7256413 | male | United Kingdom, Undy |
| | PE | ERR7260426 | ERR7260426 | male | France, Saint-Jean-de-Lier |
| | PE | S33 | SRR30238149 | male | Russia, Pskov |
| *M. putorius furo* | PE | Solexa | SRX034840<br>SRX034844<br>SRX034846 | female | USA |
| | PE | SRR10395504 | SRR10395504 | male | USA |
| | PE | ERR7256378 | ERR7256378 | female | USA |
| | PE | ERR7256379 | ERR7256379 | male | USA |
| | PE | ERR7256380 | ERR7256380 | female | USA |
| | PE | ERR7256381 | ERR7256381 | female | USA |
| | PE | ERR7256382 | ERR7256382 | male | China |
| | PE | ERR7256383 | ERR7256383 | male | China |
| | PE | ERR7256384 | ERR7256384 | female | China |
| | PE | ERR7256385 | ERR7256385 | female | China |
| *M. eversmanni* | PE | ERR11751895 | ERR11751895 | male | Russia, Altai Krai |
| | PE | ERR7198276 | ERR7198276 | male | Mongolia |
| | PE | ERR7198277 | ERR7198277 | male | Mongolia |
| | PE | SRR12564099 | SRR12564099 | male | Experimental Fur Farm, IC&G SB RAS (Novosibirsk mink population) |
| | PE | SRR16676757 | SRR16676757 | male | Experimental Fur Farm, IC&G SB RAS (Novosibirsk mink population) |

| | | | | | |
|---|---|---|---|---|---|
| *N. vison* | PE | SRS11183343 | SRR17072712<br>SRR17072713<br>SRR17072714<br>SRR17072715 | male | China, Institute of Special animal and plant sciences |

* Data type: PE – paired-end Illumina reads; 10X – 10X Genomics linked reads.

Abramov AV. 2015. IUCN Red List of Threatened Species: Mustela altaica. IUCN Red List Threat. Species.

Abramov AV, Duckworth W, Meijaard E, Eaton J, Holden J, Long B. 2015. IUCN Red List of Threatened Species: Mustela lutreolina. IUCN Red List Threat. Species.

Abramov AV, Kaneko Y, Masuda R. 2015. IUCN Red List of Threatened Species: Mustela itatsi. IUCN Red List Threat. Species.

Abramov AV, Meschersky IG, Aniskin VM, Rozhnov VV. 2013. The mountain weasel Mustela kathiah (Carnivora: Mustelidae): Molecular and Karyological data. Biol. Bull. 40:52–60.

Abramov AV, Timmins R, Duckworth W, Choudhury A, Chan B, Ghimirey Y, Dinets V, Chutipong W. 2015. IUCN Red List of Threatened Species: Mustela sibirica. IUCN Red List Threat. Species.

Abramov AV, Timmins R, Duckworth W, Choudhury A, Chan B, Lau M, Chutipong W, Roberton S, Willcox D. 2015. IUCN Red List of Threatened Species: Mustela kathiah. IUCN Red List Threat. Species.

Abramov AV, Timmins R, Duckworth W, Choudhury A, Dinets V, Chutipong W, Roberton S, Willcox D. 2015. IUCN Red List of Threatened Species: Mustela strigidorsa. IUCN Red List Threat. Species.

Belant J, Biggins D, Griebel R, Hughes J, Garelle D. 2015. IUCN Red List of Threatened Species: Mustela nigripes. IUCN Red List Threat. Species.

Cabria MT, González EG, Gómez-Moliner BJ, Zardoya R. 2007. Microsatellite markers for the endangered European mink (Mustela lutreola) and closely related mustelids. Molecular Ecology Notes:1185–1188.

Colella JP, Frederick LM, Talbot SL, Cook JA. 2021. Extrinsically reinforced hybrid speciation within Holarctic ermine (Mustela spp.) produces an insular endemic. Divers. Distrib. 27:747–762.

Colella JP, Lan T, Schuster SC, Talbot SL, Cook JA, Lindqvist C. 2018. Whole-genome analysis of Mustela erminea finds that pulsed hybridization impacts evolution at high latitudes. Commun. Biol. 1:1–10.

Duckworth W, Hearn A, Ross J, Chutipong W. 2015. IUCN Red List of Threatened Species: Mustela nudipes. IUCN Red List Threat. Species.

Dudchenko O, Batra SS, Omer AD, Nyquist SK, Hoeger M, Durand NC, Shamim MS, Machol I, Lander ES, Aiden AP, et al. 2017. De novo assembly of the Aedes aegypti genome using Hi-C yields chromosome-length scaffolds. Science 356: 92–95.

Dudchenko O, Shamim MS, Batra SS, Durand NC, Musial NT, Mostofa R, Pham M, Glenn St Hilaire B, Yao W, Stamenova E, et al. 2018. The Juicebox Assembly Tools module facilitates de novo assembly of mammalian genomes with chromosome-length scaffolds for under $1000. Genomics Available from: http://biorxiv.org/lookup/doi/10.1101/254797

Emami-Khoyi A, Hartley DA, Ross JG, Murphy EC, Paterson AM, Cruickshank RH, Else T-A. 2016. Complete mitochondrial genome of the stoat (Mustela erminea) and New Zealand fur seal (Arctocephalus forsteri) and their significance for mammalian phylogeny. Mitochondrial DNA Part A 27:4597–4599.

Etherington GJ, Ciezarek A, Shaw R, Michaux J, Croose E, Haerty W, Di Palma F. 2022. Extensive genome introgression between domestic ferret and European polecat during population recovery in Great Britain. J. Hered. 113:500–515.

Etherington GJ, Heavens D, Baker D, Lister A, McNelly R, Garcia G, Clavijo B, Macaulay I, Haerty W, Di Palma F. 2020. Sequencing smart: De novo sequencing and assembly approaches for a non-model mammal. GigaScience 9:giaa045.

FGU Centrokhotkontrol. 2007. Status of resources game animals in Russian Federation 2003-2007. Moscow, FGU Centrokhotkontrol

Gao W, Lu Z, Liang Y, Ren Z-M. 2020. Complete mitochondrial genome of Mustela sibirica (Carnivora: Mustelidae), a protected and endangered species in China. Mitochondrial DNA Part B 5:1081–1083.

González-Maya J, Helgen K, Emmons L, Arias-Alzate A. 2016. IUCN Red List of Threatened Species: Mustela felipei. IUCN Red List Threat. Species.

Graphodatsky AS, Yang F, Perelman PL, O'Brien PCM, Serdukova NA, Milne BS, Biltueva LS, Fu B, Vorobieva NV, Kawada S-I, et al. 2002. Comparative molecular cytogenetic studies in the order Carnivora: mapping chromosomal rearrangements onto the phylogenetic tree. Cytogenet. Cell Genet. 96:137–145.

Harding LE, Smith FA. 2009. Mustela or Vison? Evidence for the taxonomic status of the American mink and a distinct biogeographic radiation of American weasels. Mol. Phylogenet. Evol. 52:632–642.

Hassanin A, Veron G, Ropiquet A, Vuuren BJ van, Lécu A, Goodman SM, Haider J, Nguyen TT. 2021. Evolutionary history of Carnivora (Mammalia, Laurasiatheria) inferred from mitochondrial genomes. PLOS ONE 16:e0240770.

Helgen K, Emmons L. 2015. IUCN Red List of Threatened Species: Mustela africana. IUCN Red List Threat. Species.

Helgen K, Reid F. 2015. IUCN Red List of Threatened Species: Mustela frenata. IUCN Red List Threat. Species.

Hoeppner MP, Lundquist A, Pirun M, Meadows JRS, Zamani N, Johnson J, Sundström G, Cook A, FitzGerald MG, Swofford R, et al. 2014. An Improved Canine Genome and a Comprehensive Catalogue of Coding Genes and Non-Coding Transcripts. PLOS ONE 9:e91172.

Hua Y, Xu Y. 2016. Evolutionary status of the invasive American mink Neovison vison revealed by complete mitochondrial genome. Mitochondrial DNA Part B 1:6–7.

Huang J, Yang B, Yan C, Yang C, Tu F, Zhang X, Yue B. 2014. Phylogenetic analysis of the Mustela altaica (Carnivora: Mustelidae) based on complete mitochondrial genome. Mitochondrial DNA 25:255–256.

Karimi K, Do DN, Wang J, Easley J, Borzouie S, Sargolzaei M, Plastow G, Wang Z, Miar Y. 2022. A chromosome-level genome assembly reveals genomic characteristics of the American mink (Neogale vison). Commun. Biol. 5:1–11.

Kliver S, Houck ML, Perelman PL, Totikov A, Tomarovsky A, Dudchenko O, Omer AD, Colaric Z, Weisz D, Aiden EL, et al. 2023. Chromosome-length genome assembly and karyotype of the endangered black-footed ferret (Mustela nigripes). J. Hered. 114:539–548.

Koepfli K-P, Deere KA, Slater GJ, Begg C, Begg K, Grassman L, Lucherini M, Veron G, Wayne RK. 2008. Multigene phylogeny of the Mustelidae: Resolving relationships, tempo and biogeographic history of a mammalian adaptive radiation. BMC Biol. 6:10.

Kranz A, Reid F, Helgen K. 2015. IUCN Red List of Threatened Species: Mustela erminea. IUCN Red List Threat. Species.

Kryštufek B, Stubbe M, Tikhonov A, Herrero J, McDonald R, Cavallini P, Reid F, Abramov AV, Kranz A, Maran T. 2015. IUCN Red List of Threatened Species: Mustela nivalis. IUCN Red List Threat. Species [Internet]. Available from: https://www.iucnredlist.org/en

Lim SJ, Kim HR, Cho JY, Park YC. 2017. Complete mitochondrial genome of the least weasel Mustela nivalis (Mustelidae) in Korea. Mitochondrial DNA Part B 2:740–741.

Liu Y, Pu Y, Chen S, Wang Xuming, Murphy RW, Wang Xin, Liao R, Tang K, Yue B, Liu S. 2023. Revalidation and expanded description of Mustela aistoodonnivalis (Mustelidae: Carnivora) based on a multigene phylogeny and morphology. Ecol. Evol. 13:e9944.

Lukashkova VM, Spivak AA, Kotova SA. 2023. Informative Relevance of 11 Microsatellite Loci for Forensic DNA Identification of Wild and Farm American Mink (Mustela vison) in Belarus. Russ. J. Genet. 59:396–407.

Maran T, Skumatov D, Gomez A, Põdra M, Abramov AV, Dinets V. 2015. IUCN Red List of Threatened Species: Mustela lutreola. IUCN Red List Threat. Species.

Maran T, Skumatov D, Kranz A, Abramov AV. 2015. IUCN Red List of Threatened Species: Mustela eversmanii. IUCN Red List Threat. Species.

Maran T, Skumatov D, Kranz A, Saveljev A, Abramov AV, Sándor AD, Kitchener A, Herrero J, Savour-Soubelet A, Zuberogoitia I, et al. 2016. IUCN Red List of Threatened Species: Mustela putorius. IUCN Red List Threat. Species.

Minprirody of Russia. 2019. State report “On the State and Environmental Protection of the Russian Federation in 2018.” Ministry of Natural Resources and Environment of the Russian Federation Available from: https://gosdoklad-ecology.ru/2018/

Minprirody of Russia. 2021. Number of main species of hunting resources in the Russian Federation (thousands of species). Ministry of Natural Resources and Environment of the Russian Federation. Available from: https://www.mnr.gov.ru/opendata/7710256289-hunting-resources

Minprirody of Russia. 2024. State report “On the State and Environmental Protection of the Russian Federation in 2023.” Ministry of Natural Resources and Environment of the Russian Federation Available from: https://2023.ecology-gosdoklad.ru/

Miranda I, Giska I, Farelo L, Pimenta J, Zimova M, Bryk J, Dalén L, Mills LS, Zub K, Melo-Ferreira J. 2021. Museomics Dissects the Genetic Basis for Adaptive Seasonal Coloration in the Least Weasel. Mol. Biol. Evol. 38:4388–4402.

Peng X, Alföldi J, Gori K, Eisfeld AJ, Tyler SR, Tisoncik-Go J, Brawand D, Law GL, Skunca N, Hatta M, et al. 2014. The draft genome sequence of the ferret (Mustela putorius furo) facilitates study of human respiratory disease. Nat. Biotechnol. 32:1250–1255.

Reid F, Schipper J, Schiaffini M. 2015. IUCN Red List of Threatened Species: Neovison vison. IUCN Red List Threat. Species.

Sato JJ, Hosoda T, Wolsan M, Suzuki H. 2004. Molecular phylogeny of arctoids (Mammalia: Carnivora) with emphasis on phylogenetic and taxonomic positions of the ferret-badgers and skunks. Zoolog. Sci. 21:111–118.

Schneider VA, Graves-Lindsay T, Howe K, Bouk N, Chen H-C, Kitts PA, Murphy TD, Pruitt KD, Thibaud-Nissen F, Albracht D, et al. 2017. Evaluation of GRCh38 and de novo haploid genome assemblies demonstrates the enduring quality of the reference assembly. Genome Res. 27:849–864.

Shalabi MA, Abramov AV, Kosintsev PA, Lin L-K, Han S-H, Watanabe S, Yamazaki K, Kaneko Y, Masuda R. 2017. Comparative phylogeography of the endemic Japanese weasel (Mustela itatsi) and the continental Siberian weasel (Mustela sibirica) revealed by complete mitochondrial genome sequences. Biol. J. Linn. Soc. 120:333–348.

Skorupski J. 2022. Characterisation of the Complete Mitochondrial Genome of Critically Endangered Mustela lutreola (Carnivora: Mustelidae) and Its Phylogenetic and Conservation Implications. Genes 13:125.

Skorupski J, Brandes F, Seebass C, Festl W, Śmietana P, Balacco J, Jain N, Tilley T, Abueg L, Wood J, et al. 2023. Prioritizing Endangered Species in Genome Sequencing: Conservation Genomics in Action with the First Platinum-Standard Reference-Quality Genome of the Critically Endangered European Mink Mustela lutreola L., 1761. Int. J. Mol. Sci. 24:14816.

Sun W, Wang S, Wang Z, Liu H, Zhong W, Yang Y, Li G. 2016. The complete mitochondrial genome sequence of Neovison vison (Carnivora: Mustelidae). Mitochondrial DNA Part A 27:1840–1841.

Szatmári L, Cserkész T, Laczkó L, Lanszki J, Pertoldi C, Abramov AV, Elmeros M, Ottlecz B, Hegyeli Z, Sramkó G. 2021. A comparison of microsatellites and genome-wide SNPs for the detection of admixture brings the first molecular evidence for hybridization between Mustela eversmanii and M. putorius (Mustelidae, Carnivora). Evol. Appl. 14:2286–2304.

Yu L, Peng D, Liu J, Luan P, Liang L, Lee H, Lee M, Ryder OA, Zhang Y. 2011. On the phylogeny of Mustelidae subfamilies: analysis of seventeen nuclear non-coding loci and mitochondrial complete genomes. BMC Evol. Biol. 11:92.

Yu M, Xu H, Li D, Wu J, Wen A, Xie M, Wang Q, Zhu G, Ni Q, Zhang M, et al. 2019. The complete mitochondrial genome sequence and phylogenetic analysis of yellow weasel (Mustela sibirica). Mitochondrial DNA Part B 4:3698–3699.

Zhao R-B, Zhou C-Y, Lu Z-X, Hu P, Liu J-Q, Tan W-W, Yang T-H. 2016. The complete mitochondrial genome of black-footed ferret, Mustela nigripes (Mustela, Mustelinae). Mitochondrial DNA Part A [Internet]. Available from: https://www.tandfonline.com/doi/abs/10.3109/19401736.2014.958685
