## Supplementary File for "Comparative genomics and phylogenomics of the Mustelinae lineage (Mustelidae, Carnivora)"

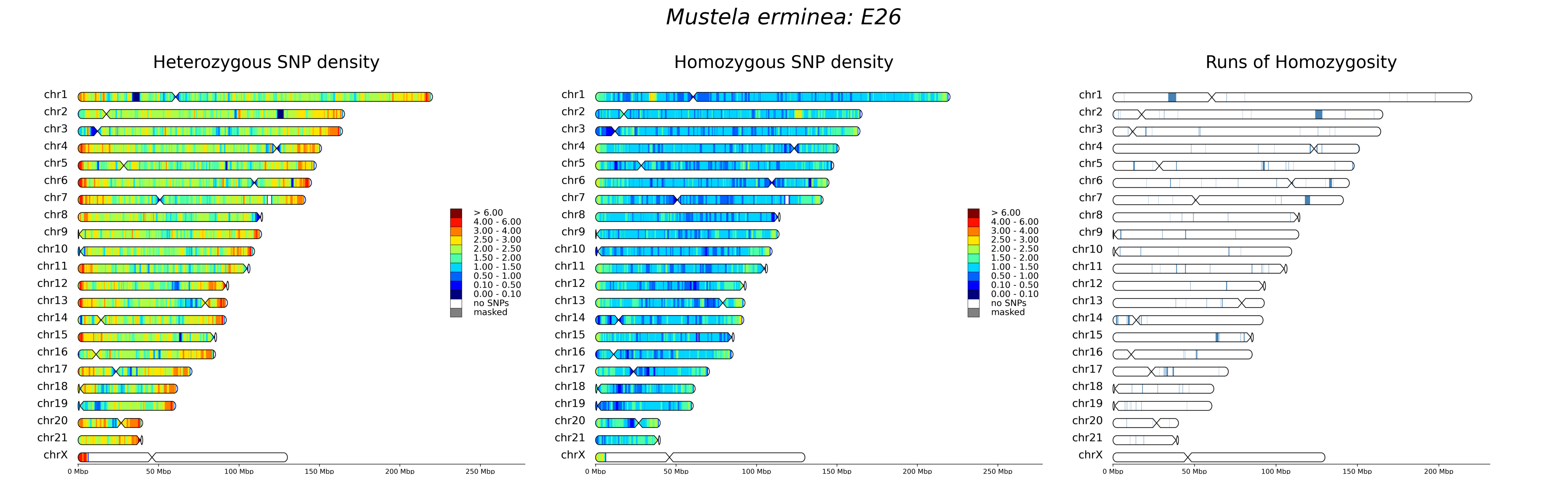

Mustela erminea: E26
Heterozygous SNP density
Homozygous SNP density
Runs of Homozygosity
chr1
chr2
chr3
chr4
chr5
chr6
chr7
chr8
chr9
chr10
chr11
chr12
chr13
chr14
chr15
chr16
chr17
chr18
chr19
chr20
chr21
chrX
> 6.00
4.00 - 6.00
3.00 - 4.00
2.50 - 3.00
2.00 - 2.50
1.50 - 2.00
1.00 - 1.50
0.50 - 1.00
0.10 - 0.50
0.00 - 0.10
no SNPs
masked
0 Mbp
50 Mbp
100 Mbp
150 Mbp
200 Mbp
250 Mbp

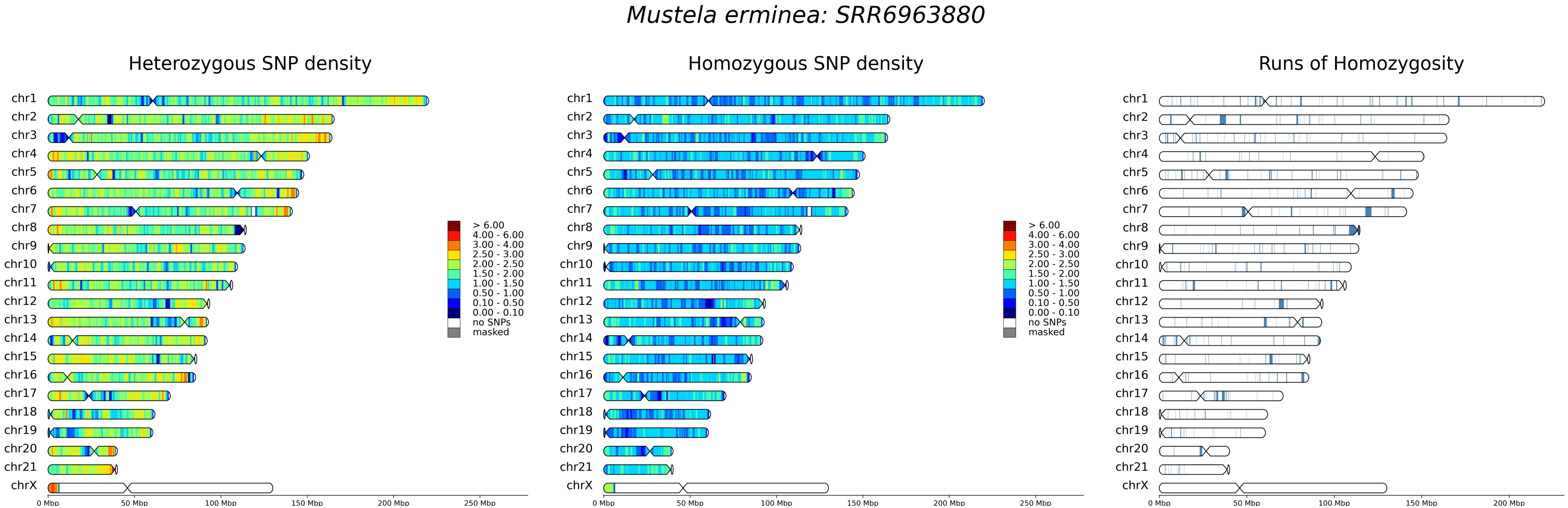
Mustela erminea: SRR6963880
Heterozygous SNP density
Homozygous SNP density
Runs of Homozygosity
chr1
chr2
chr3
chr4
chr5
chr6
chr7
chr8
chr9
chr10
chr11
chr12
chr13
chr14
chr15
chr16
chr17
chr18
chr19
chr20
chr21
chrX
> 6.00
4.00 - 6.00
3.00 - 4.00
2.50 - 3.00
2.00 - 2.50
1.50 - 2.00
1.00 - 1.50
0.50 - 1.00
0.10 - 0.50
0.00 - 0.10
no SNPs
masked
0 Mbp
50 Mbp
100 Mbp
150 Mbp
200 Mbp
250 Mbp

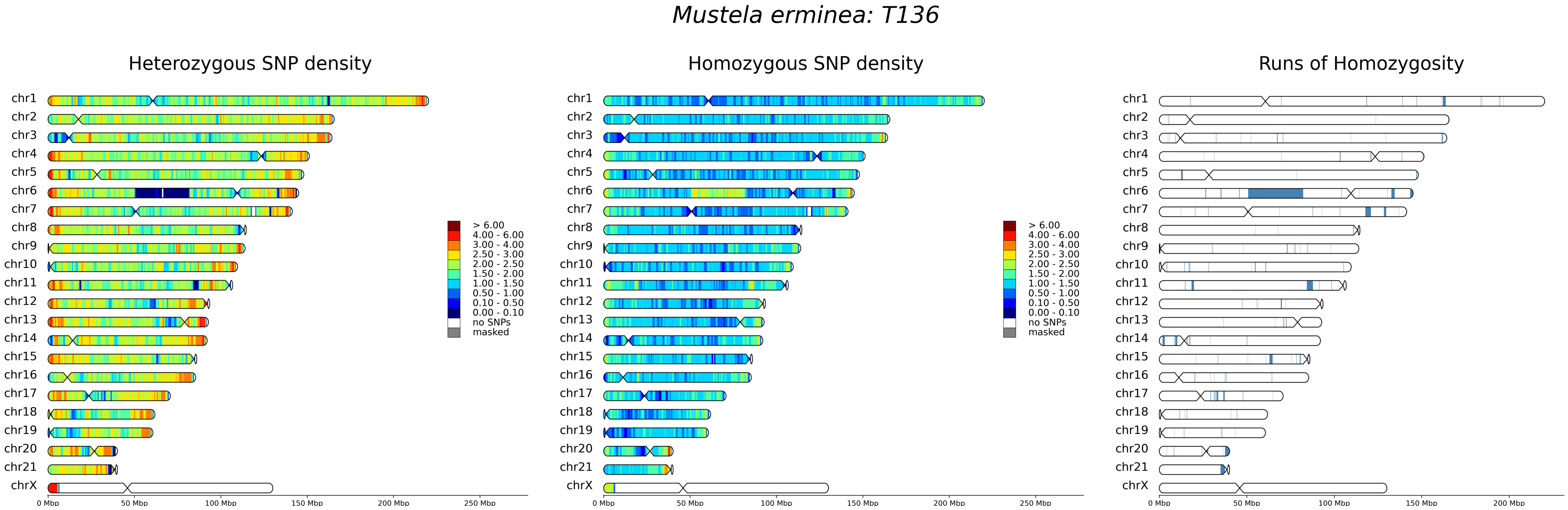
Mustela erminea: T136
Heterozygous SNP density
Homozygous SNP density
Runs of Homozygosity
chr1
chr2
chr3
chr4
chr5
chr6
chr7
chr8
chr9
chr10
chr11
chr12
chr13
chr14
chr15
chr16
chr17
chr18
chr19
chr20
chr21
chrX
> 6.00
4.00 - 6.00
3.00 - 4.00
2.50 - 3.00
2.00 - 2.50
1.50 - 2.00
1.00 - 1.50
0.50 - 1.00
0.10 - 0.50
0.00 - 0.10
no SNPs
masked
0 Mbp
50 Mbp
100 Mbp
150 Mbp
200 Mbp
250 Mbp

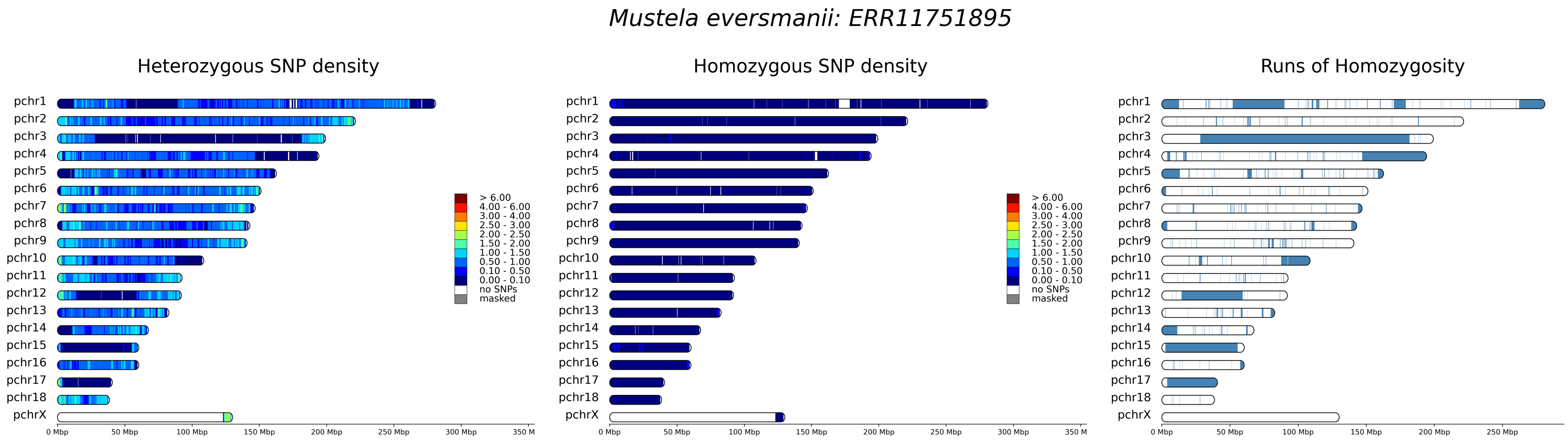
Mustela eversmanii: ERR11751895
Heterozygous SNP density
Homozygous SNP density
Runs of Homozygosity
pchr1
pchr2
pchr3
pchr4
pchr5
pchr6
pchr7
pchr8
pchr9
pchr10
pchr11
pchr12
pchr13
pchr14
pchr15
pchr16
pchr17
pchr18
pchrX
> 6.00
4.00 - 6.00
3.00 - 4.00
2.50 - 3.00
2.00 - 2.50
1.50 - 2.00
1.00 - 1.50
0.50 - 1.00
0.10 - 0.50
0.00 - 0.10
no SNPs
masked
0 Mbp
50 Mbp
100 Mbp
150 Mbp
200 Mbp
250 Mbp
300 Mbp
350 M

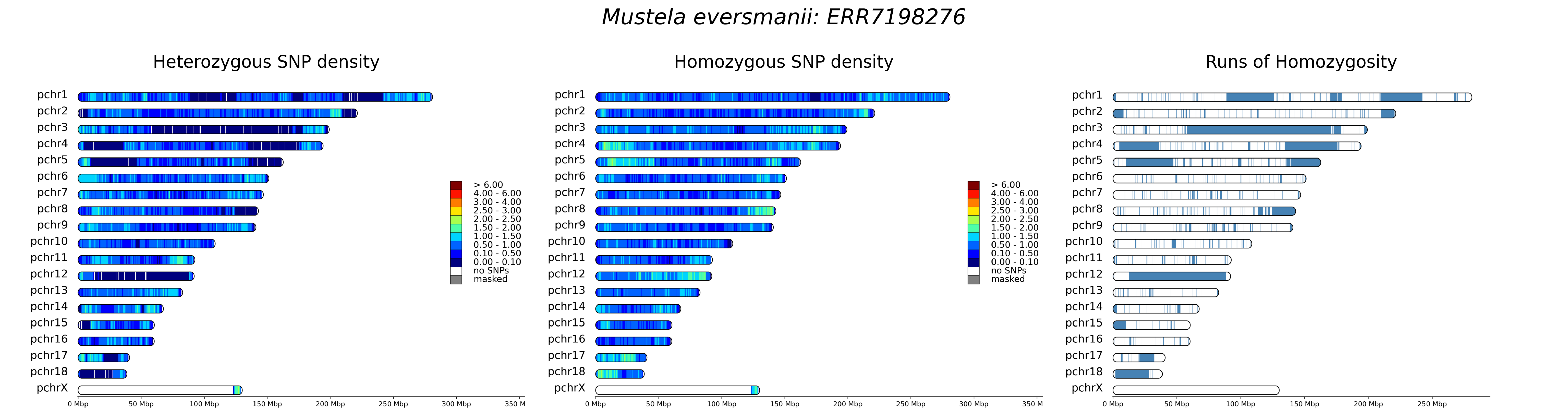

Mustela eversmanii: ERR7198276
Heterozygous SNP density
Homozygous SNP density
Runs of Homozygosity
pchr1
pchr2
pchr3
pchr4
pchr5
pchr6
pchr7
pchr8
pchr9
pchr10
pchr11
pchr12
pchr13
pchr14
pchr15
pchr16
pchr17
pchr18
pchrX
> 6.00
4.00 - 6.00
3.00 - 4.00
2.50 - 3.00
2.00 - 2.50
1.50 - 2.00
1.00 - 1.50
0.50 - 1.00
0.10 - 0.50
0.00 - 0.10
no SNPs
masked
0 Mbp
50 Mbp
100 Mbp
150 Mbp
200 Mbp
250 Mbp
300 Mbp
350 M

### *Mustela eversmanii: ERR7198277*

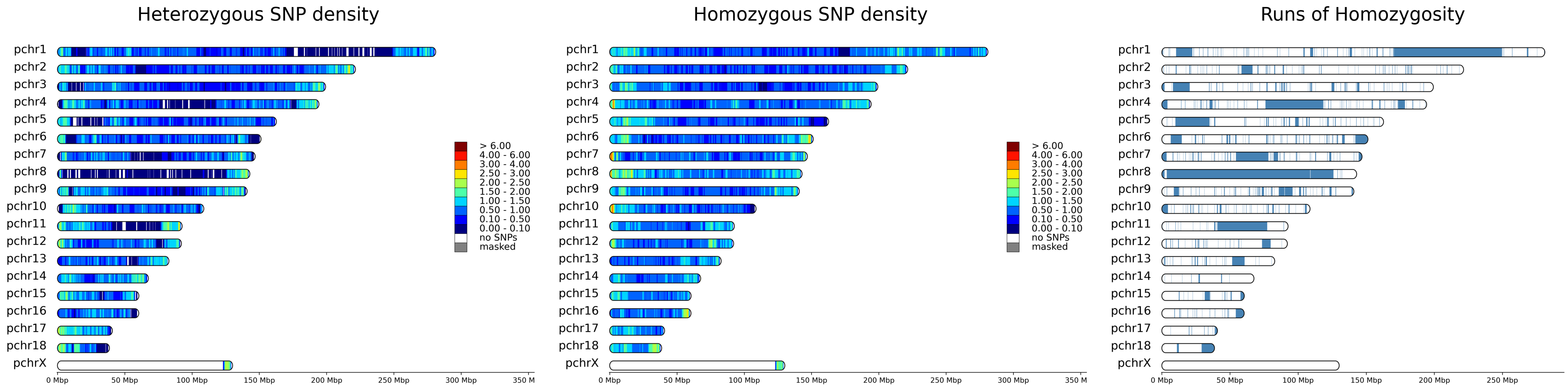

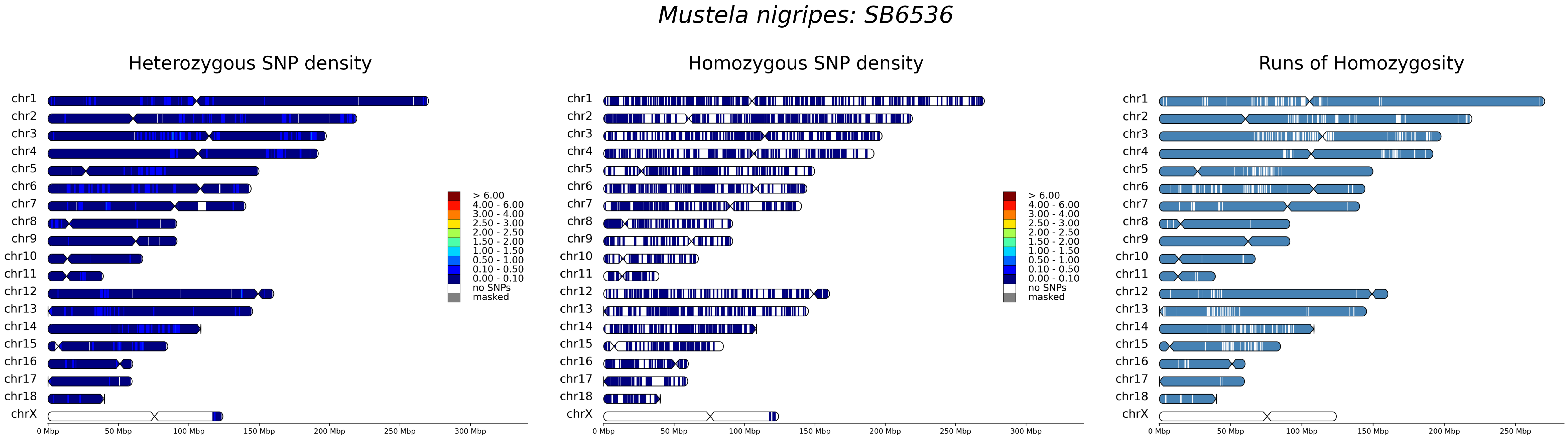

Mustela nigripes: SB6536
Heterozygous SNP density
Homozygous SNP density
Runs of Homozygosity
chr1
chr2
chr3
chr4
chr5
chr6
chr7
chr8
chr9
chr10
chr11
chr12
chr13
chr14
chr15
chr16
chr17
chr18
chrX
> 6.00
4.00 - 6.00
3.00 - 4.00
2.50 - 3.00
2.00 - 2.50
1.50 - 2.00
1.00 - 1.50
0.50 - 1.00
0.10 - 0.50
0.00 - 0.10
no SNPs
masked
0 Mbp
50 Mbp
100 Mbp
150 Mbp
200 Mbp
250 Mbp
300 Mbp

### *Mustela nigripes: SB7462*

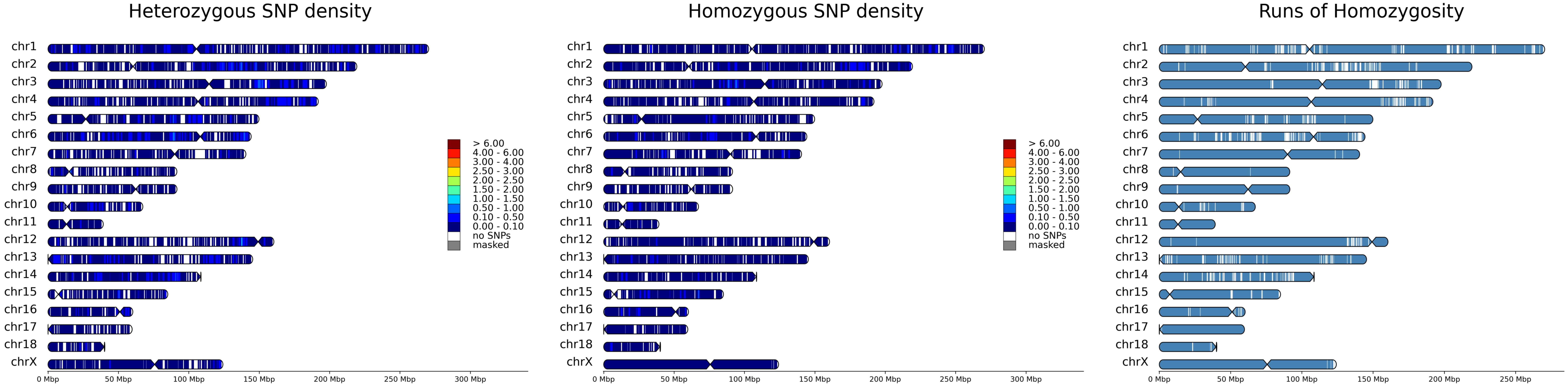

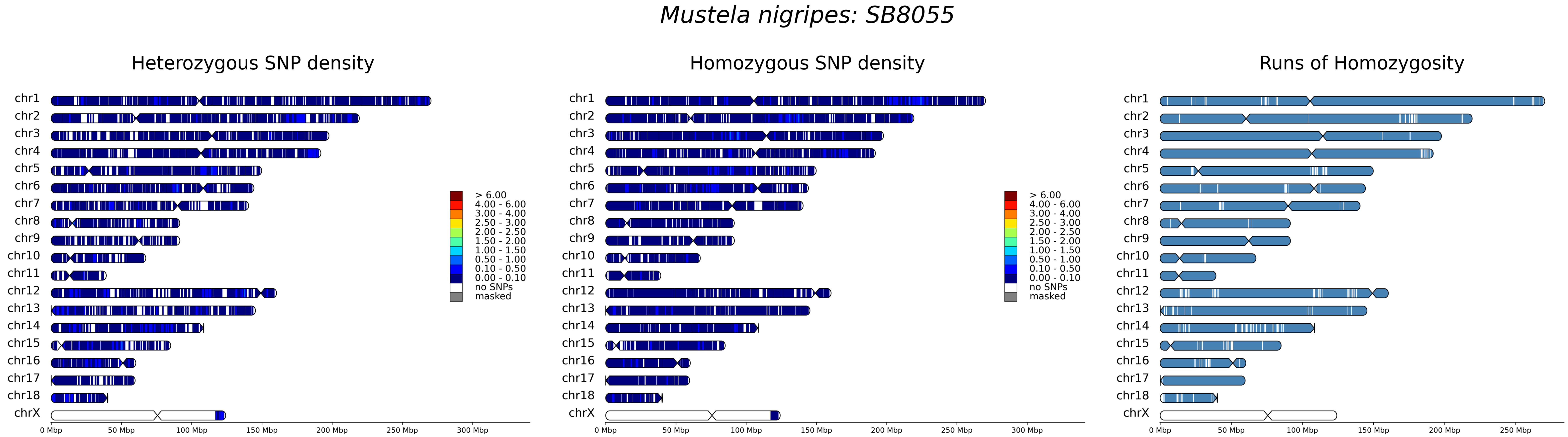
Mustela nigripes: SB8055
Heterozygous SNP density
Homozygous SNP density
Runs of Homozygosity
chr1
chr2
chr3
chr4
chr5
chr6
chr7
chr8
chr9
chr10
chr11
chr12
chr13
chr14
chr15
chr16
chr17
chr18
chrX
> 6.00
4.00 - 6.00
3.00 - 4.00
2.50 - 3.00
2.00 - 2.50
1.50 - 2.00
1.00 - 1.50
0.50 - 1.00
0.10 - 0.50
0.00 - 0.10
no SNPs
masked
0 Mbp
50 Mbp
100 Mbp
150 Mbp
200 Mbp
250 Mbp
300 Mbp

### *Mustela nigripes: SRR1508214*

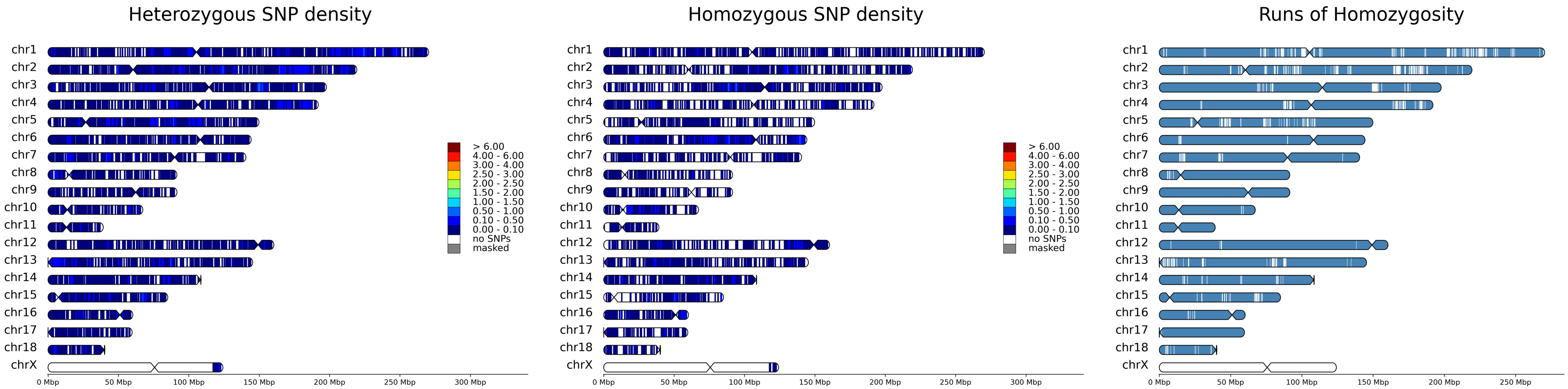

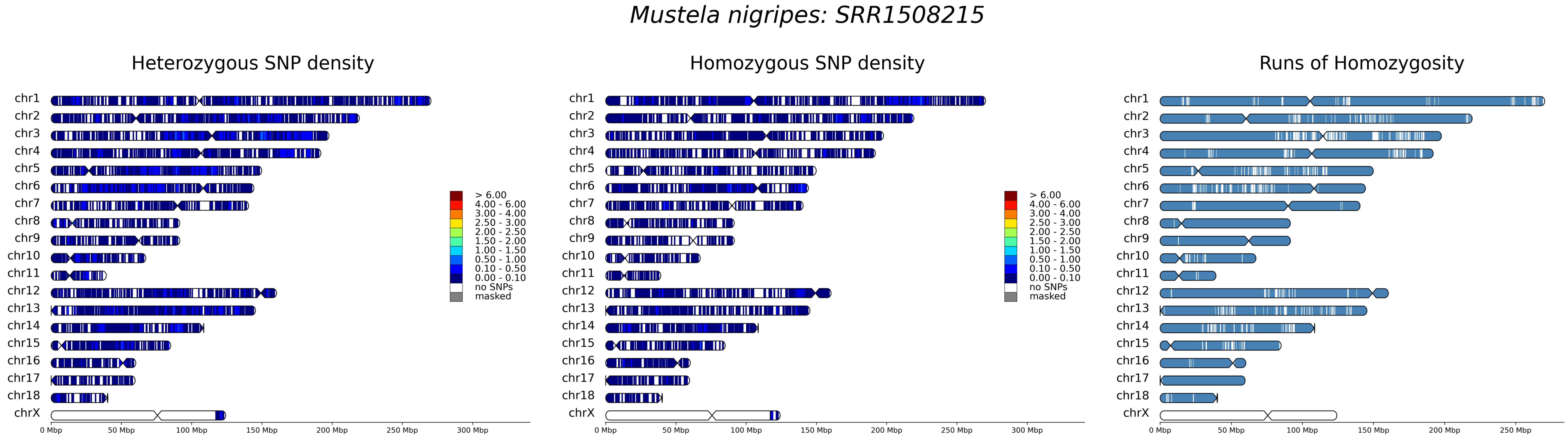

Mustela nigripes: SRR1508215
Heterozygous SNP density
Homozygous SNP density
Runs of Homozygosity
chr1
chr2
chr3
chr4
chr5
chr6
chr7
chr8
chr9
chr10
chr11
chr12
chr13
chr14
chr15
chr16
chr17
chr18
chrX
> 6.00
4.00 - 6.00
3.00 - 4.00
2.50 - 3.00
2.00 - 2.50
1.50 - 2.00
1.00 - 1.50
0.50 - 1.00
0.10 - 0.50
0.00 - 0.10
no SNPs
masked
0 Mbp
50 Mbp
100 Mbp
150 Mbp
200 Mbp
250 Mbp
300 Mbp

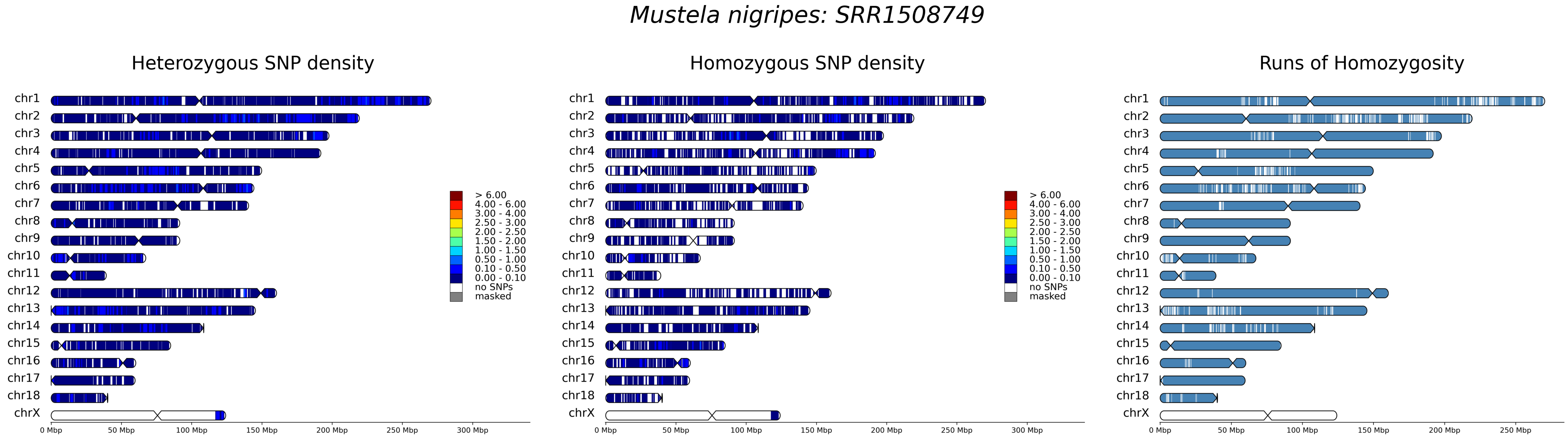
Mustela nigripes: SRR1508749
Heterozygous SNP density
Homozygous SNP density
Runs of Homozygosity
chr1
chr2
chr3
chr4
chr5
chr6
chr7
chr8
chr9
chr10
chr11
chr12
chr13
chr14
chr15
chr16
chr17
chr18
chrX
> 6.00
4.00 - 6.00
3.00 - 4.00
2.50 - 3.00
2.00 - 2.50
1.50 - 2.00
1.00 - 1.50
0.50 - 1.00
0.10 - 0.50
0.00 - 0.10
no SNPs
masked
0 Mbp
50 Mbp
100 Mbp
150 Mbp
200 Mbp
250 Mbp
300 Mbp

### *Mustela nigripes: SRR1508750*

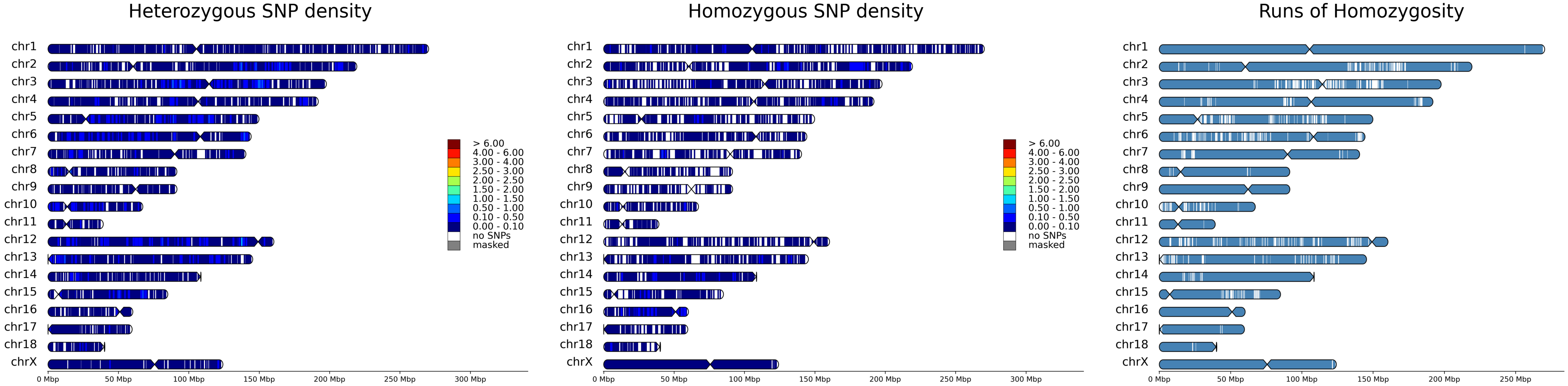

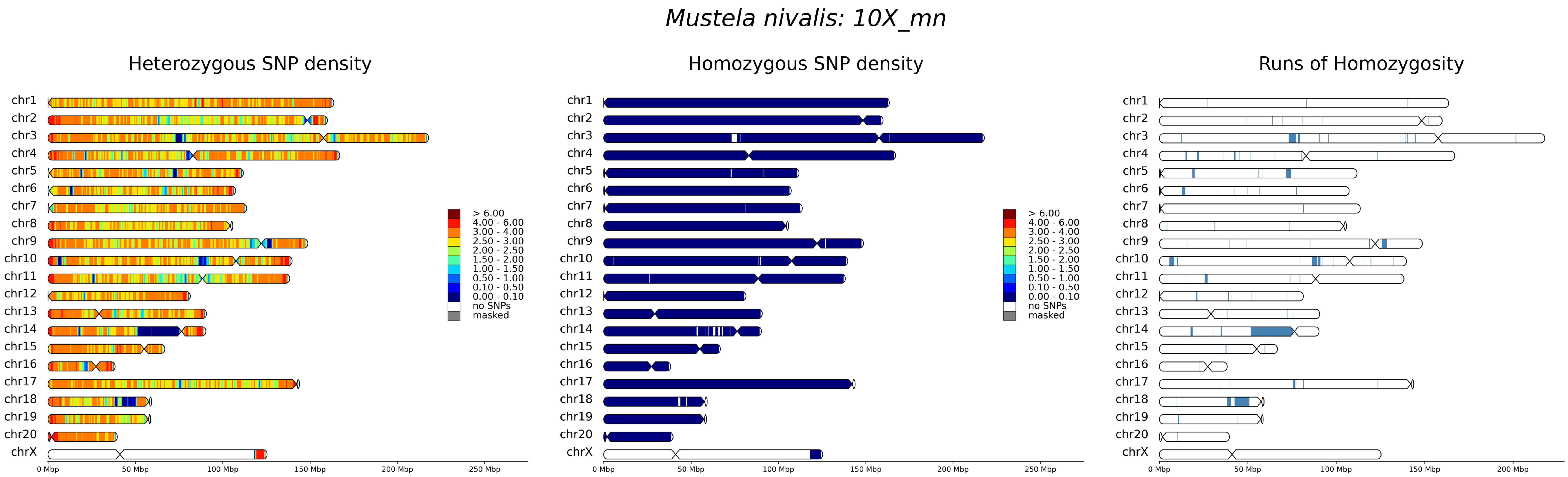

Mustela nivalis: 10X_mn
Heterozygous SNP density
Homozygous SNP density
Runs of Homozygosity
chr1
chr2
chr3
chr4
chr5
chr6
chr7
chr8
chr9
chr10
chr11
chr12
chr13
chr14
chr15
chr16
chr17
chr18
chr19
chr20
chrX
> 6.00
4.00 - 6.00
3.00 - 4.00
2.50 - 3.00
2.00 - 2.50
1.50 - 2.00
1.00 - 1.50
0.50 - 1.00
0.10 - 0.50
0.00 - 0.10
no SNPs
masked
0 Mbp
50 Mbp
100 Mbp
150 Mbp
200 Mbp
250 Mbp

### *Mustela nivalis: ERR7198278*

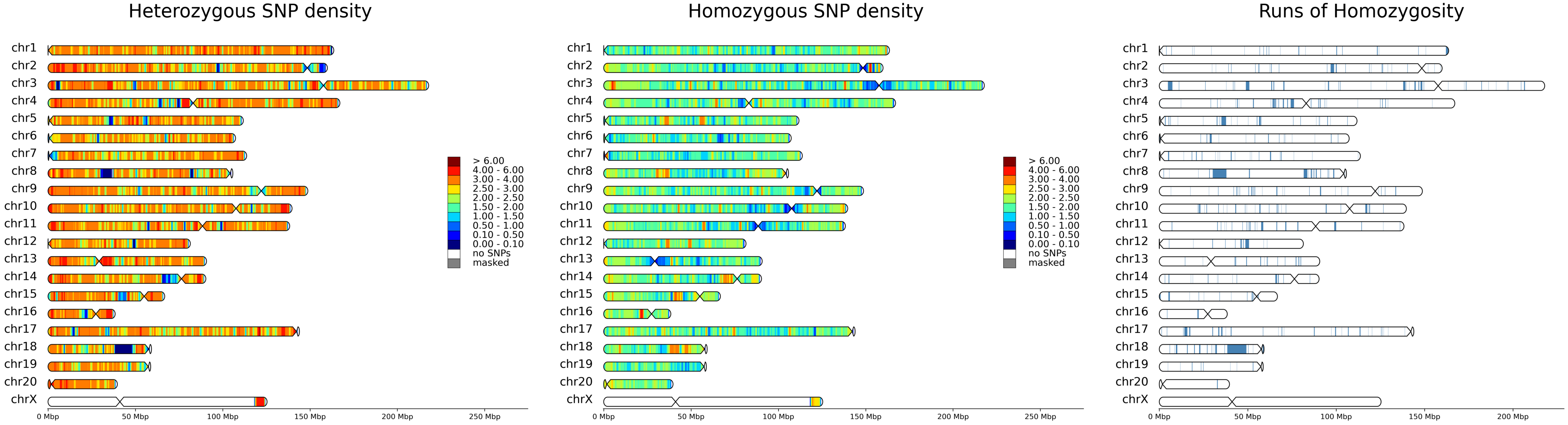

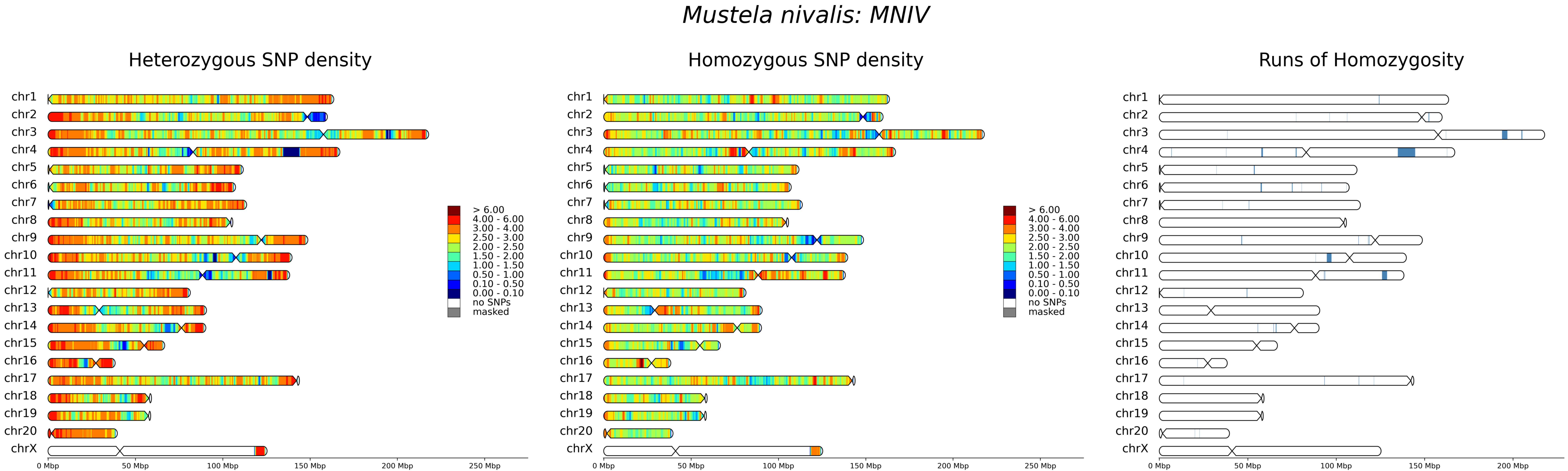

Mustela nivalis: MNIV
Heterozygous SNP density
Homozygous SNP density
Runs of Homozygosity
chr1
chr2
chr3
chr4
chr5
chr6
chr7
chr8
chr9
chr10
chr11
chr12
chr13
chr14
chr15
chr16
chr17
chr18
chr19
chr20
chrX
> 6.00
4.00 - 6.00
3.00 - 4.00
2.50 - 3.00
2.00 - 2.50
1.50 - 2.00
1.00 - 1.50
0.50 - 1.00
0.10 - 0.50
0.00 - 0.10
no SNPs
masked
0 Mbp
50 Mbp
100 Mbp
150 Mbp
200 Mbp
250 Mbp

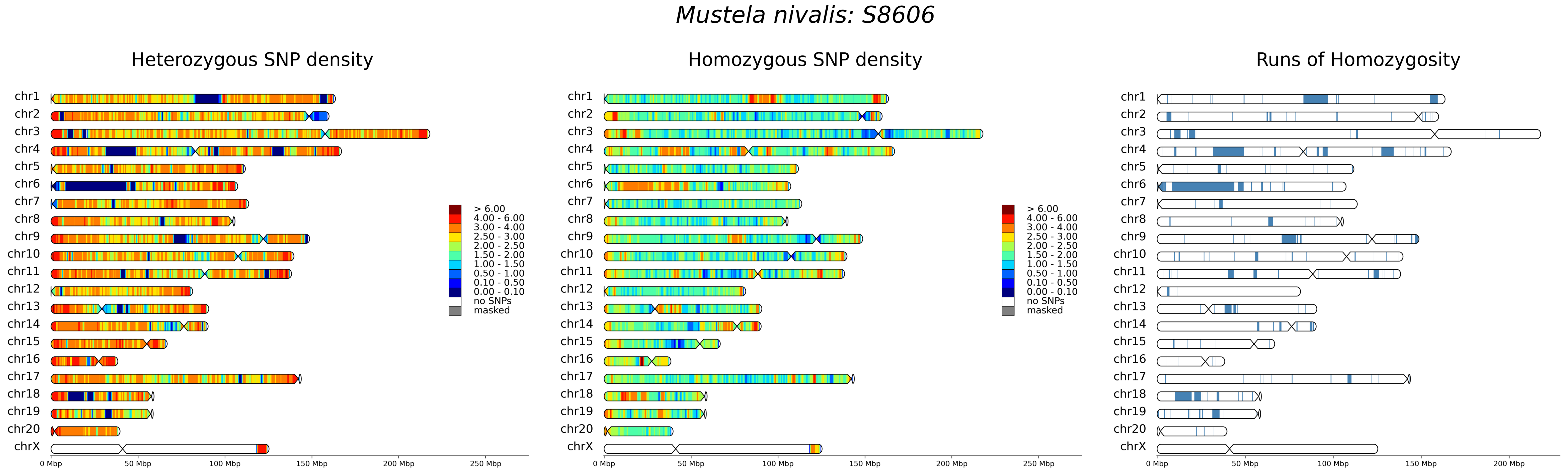

Mustela nivalis: S8606
Heterozygous SNP density
Homozygous SNP density
Runs of Homozygosity
chr1
chr2
chr3
chr4
chr5
chr6
chr7
chr8
chr9
chr10
chr11
chr12
chr13
chr14
chr15
chr16
chr17
chr18
chr19
chr20
chrX
> 6.00
4.00 - 6.00
3.00 - 4.00
2.50 - 3.00
2.00 - 2.50
1.50 - 2.00
1.00 - 1.50
0.50 - 1.00
0.10 - 0.50
0.00 - 0.10
no SNPs
masked
0 Mbp
50 Mbp
100 Mbp
150 Mbp
200 Mbp
250 Mbp

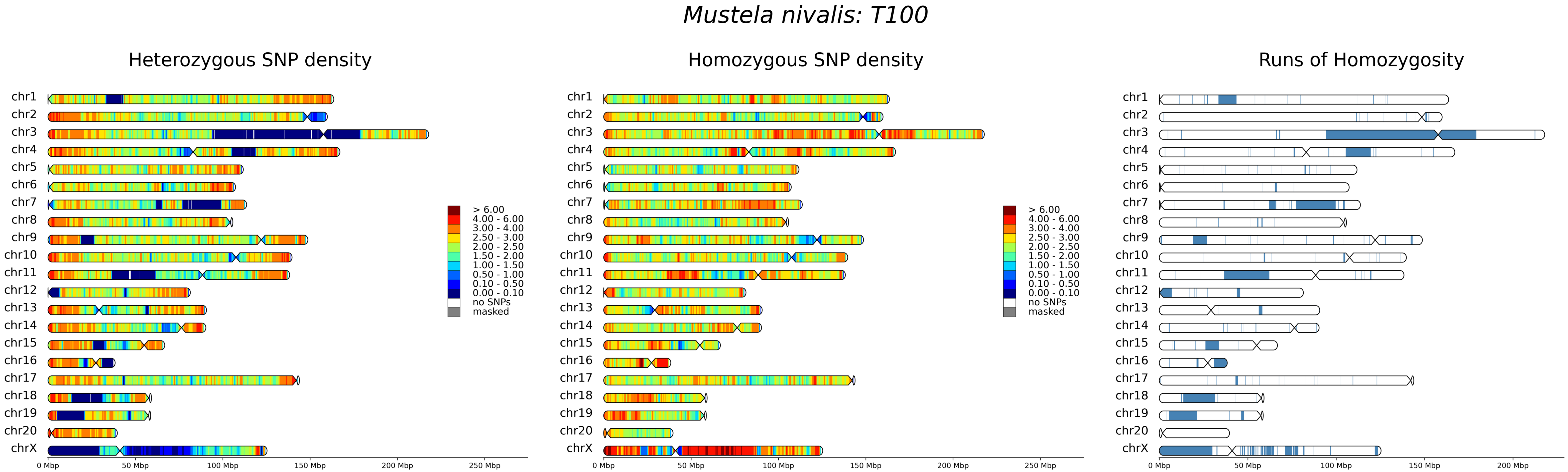

Mustela nivalis: T100
Heterozygous SNP density
Homozygous SNP density
Runs of Homozygosity
chr1
chr2
chr3
chr4
chr5
chr6
chr7
chr8
chr9
chr10
chr11
chr12
chr13
chr14
chr15
chr16
chr17
chr18
chr19
chr20
chrX
> 6.00
4.00 - 6.00
3.00 - 4.00
2.50 - 3.00
2.00 - 2.50
1.50 - 2.00
1.00 - 1.50
0.50 - 1.00
0.10 - 0.50
0.00 - 0.10
no SNPs
masked
0 Mbp
50 Mbp
100 Mbp
150 Mbp
200 Mbp
250 Mbp

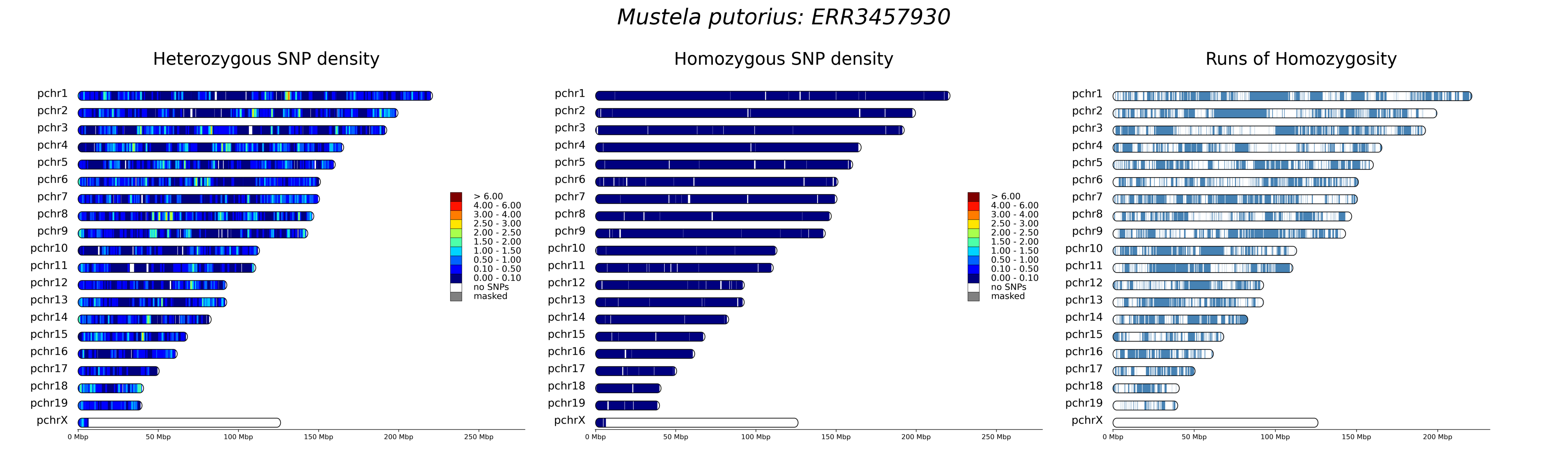
Mustela putorius: ERR3457930
Heterozygous SNP density
Homozygous SNP density
Runs of Homozygosity
pchr1
pchr2
pchr3
pchr4
pchr5
pchr6
pchr7
pchr8
pchr9
pchr10
pchr11
pchr12
pchr13
pchr14
pchr15
pchr16
pchr17
pchr18
pchr19
pchrX
> 6.00
4.00 - 6.00
3.00 - 4.00
2.50 - 3.00
2.00 - 2.50
1.50 - 2.00
1.00 - 1.50
0.50 - 1.00
0.10 - 0.50
0.00 - 0.10
no SNPs
masked
0 Mbp
50 Mbp
100 Mbp
150 Mbp
200 Mbp
250 Mbp

### *Mustela putorius: ERR7256386*

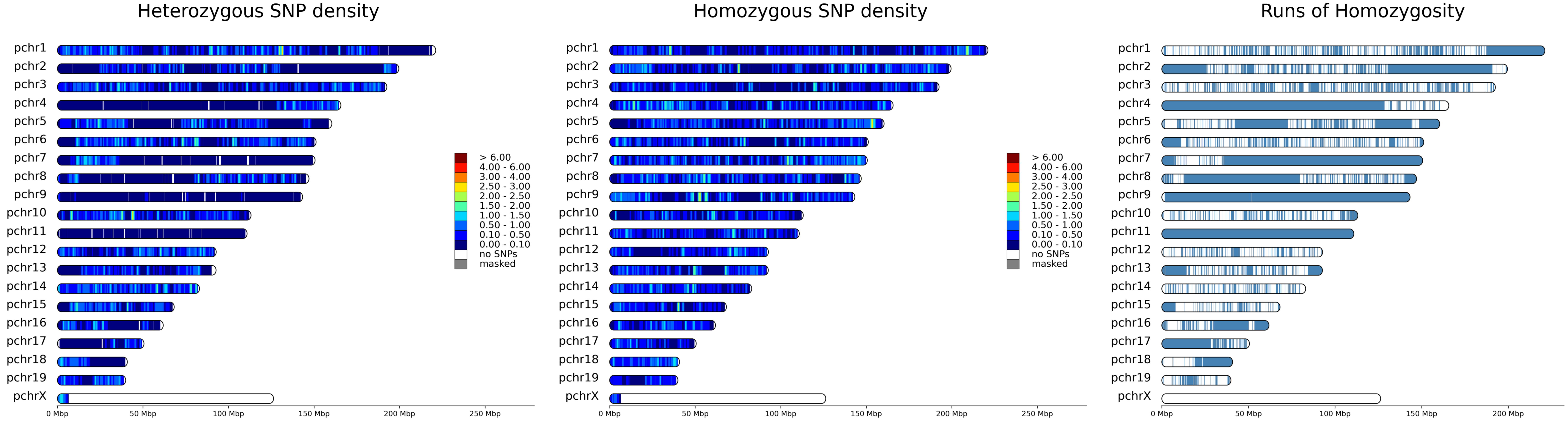

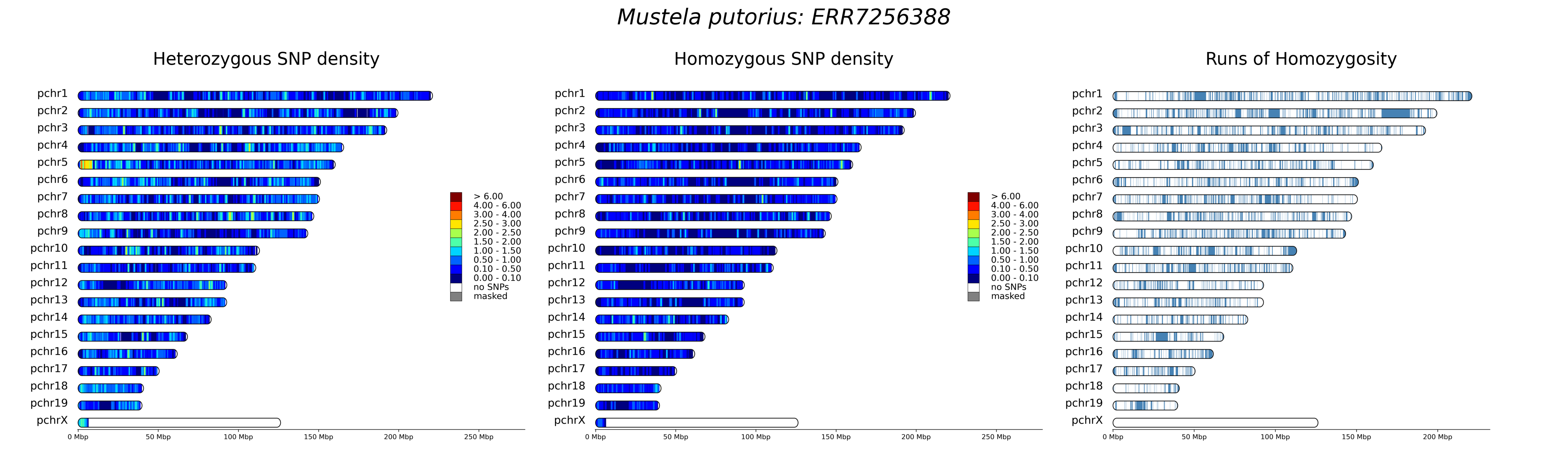
Mustela putorius: ERR7256388
Heterozygous SNP density
Homozygous SNP density
Runs of Homozygosity
pchr1
pchr2
pchr3
pchr4
pchr5
pchr6
pchr7
pchr8
pchr9
pchr10
pchr11
pchr12
pchr13
pchr14
pchr15
pchr16
pchr17
pchr18
pchr19
pchrX
> 6.00
4.00 - 6.00
3.00 - 4.00
2.50 - 3.00
2.00 - 2.50
1.50 - 2.00
1.00 - 1.50
0.50 - 1.00
0.10 - 0.50
0.00 - 0.10
no SNPs
masked
0 Mbp
50 Mbp
100 Mbp
150 Mbp
200 Mbp
250 Mbp

### *Mustela putorius: ERR7256390*

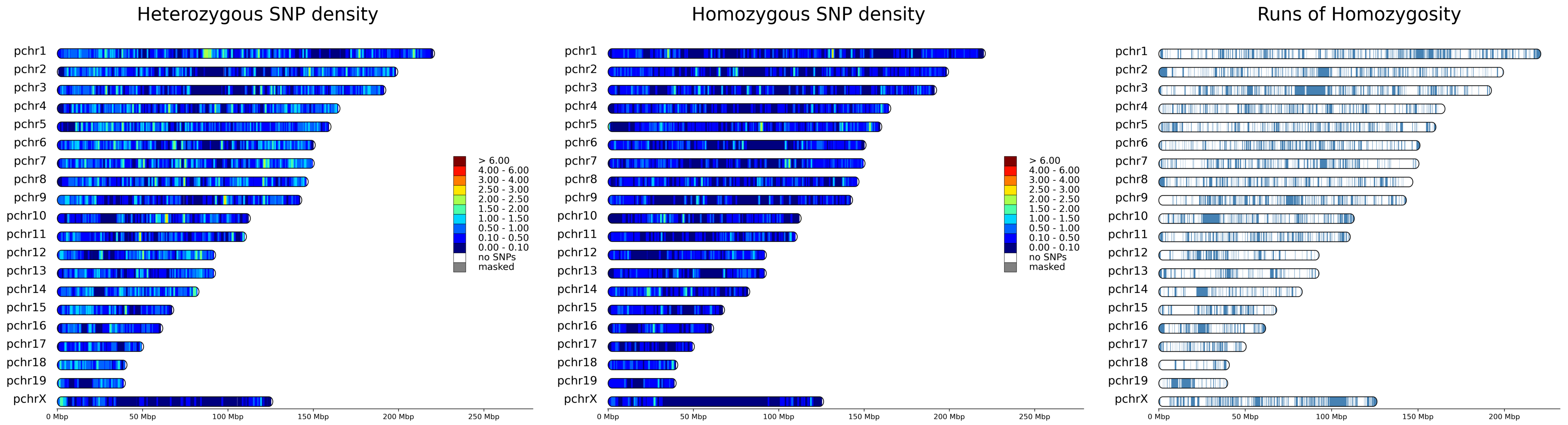

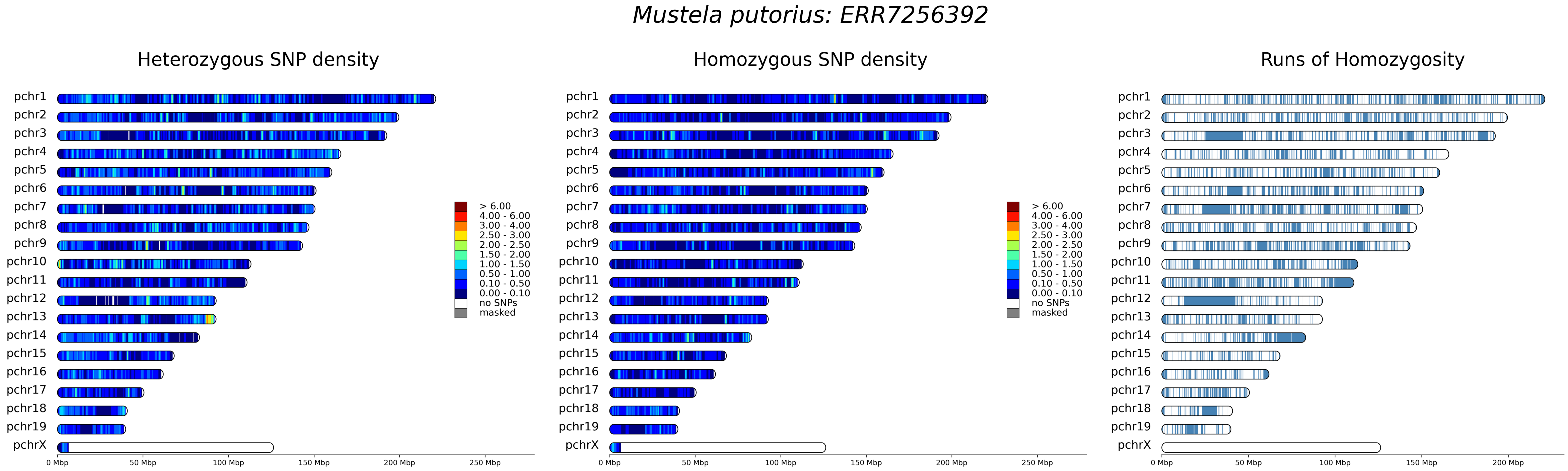

Mustela putorius: ERR7256392
Heterozygous SNP density
Homozygous SNP density
Runs of Homozygosity
pchr1
pchr2
pchr3
pchr4
pchr5
pchr6
pchr7
pchr8
pchr9
pchr10
pchr11
pchr12
pchr13
pchr14
pchr15
pchr16
pchr17
pchr18
pchr19
pchrX
> 6.00
4.00 - 6.00
3.00 - 4.00
2.50 - 3.00
2.00 - 2.50
1.50 - 2.00
1.00 - 1.50
0.50 - 1.00
0.10 - 0.50
0.00 - 0.10
no SNPs
masked
0 Mbp
50 Mbp
100 Mbp
150 Mbp
200 Mbp
250 Mbp

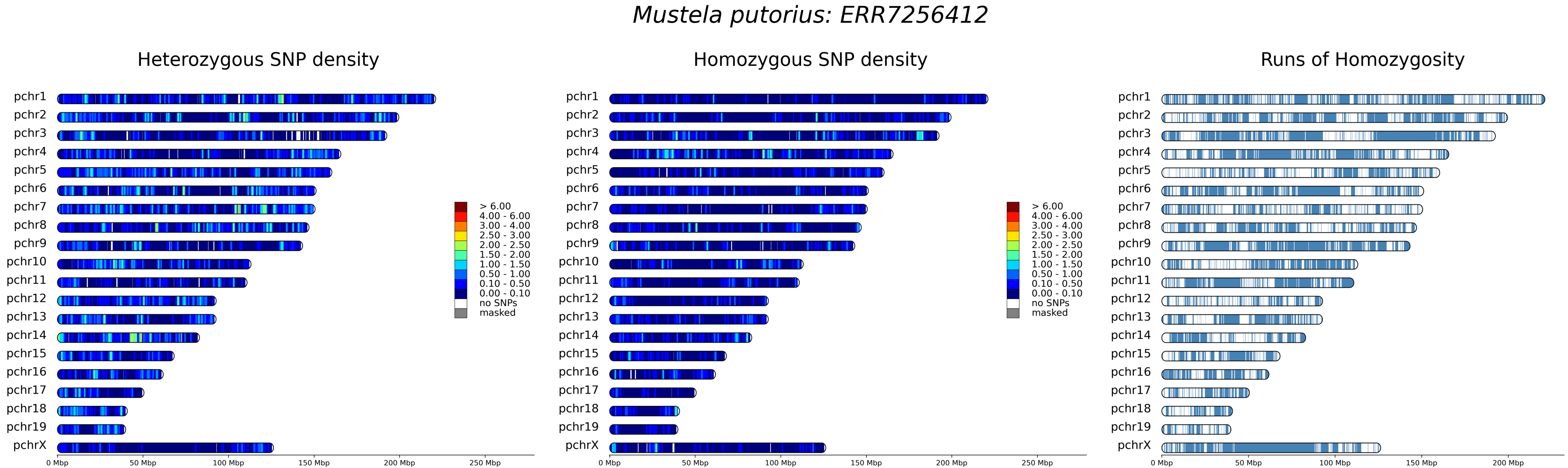

Mustela putorius: ERR7256412
Heterozygous SNP density
Homozygous SNP density
Runs of Homozygosity
pchr1
pchr2
pchr3
pchr4
pchr5
pchr6
pchr7
pchr8
pchr9
pchr10
pchr11
pchr12
pchr13
pchr14
pchr15
pchr16
pchr17
pchr18
pchr19
pchrX
> 6.00
4.00 - 6.00
3.00 - 4.00
2.50 - 3.00
2.00 - 2.50
1.50 - 2.00
1.00 - 1.50
0.50 - 1.00
0.10 - 0.50
0.00 - 0.10
no SNPs
masked
0 Mbp
50 Mbp
100 Mbp
150 Mbp
200 Mbp
250 Mbp

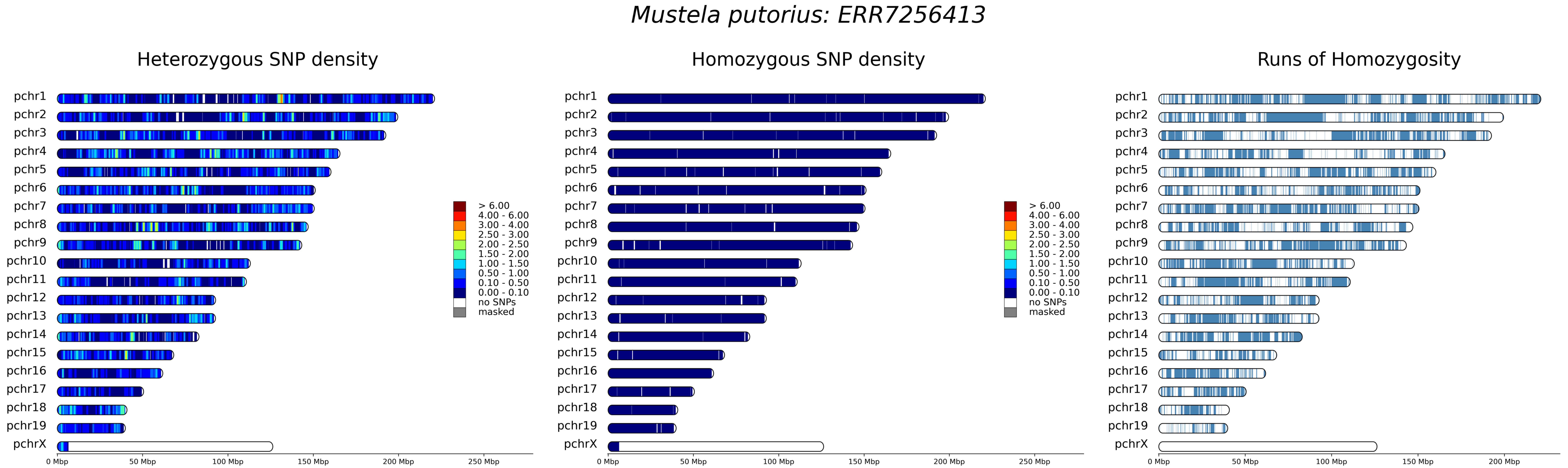
Mustela putorius: ERR7256413
Heterozygous SNP density
Homozygous SNP density
Runs of Homozygosity
pchr1
pchr2
pchr3
pchr4
pchr5
pchr6
pchr7
pchr8
pchr9
pchr10
pchr11
pchr12
pchr13
pchr14
pchr15
pchr16
pchr17
pchr18
pchr19
pchrX
> 6.00
4.00 - 6.00
3.00 - 4.00
2.50 - 3.00
2.00 - 2.50
1.50 - 2.00
1.00 - 1.50
0.50 - 1.00
0.10 - 0.50
0.00 - 0.10
no SNPs
masked
0 Mbp
50 Mbp
100 Mbp
150 Mbp
200 Mbp
250 Mbp

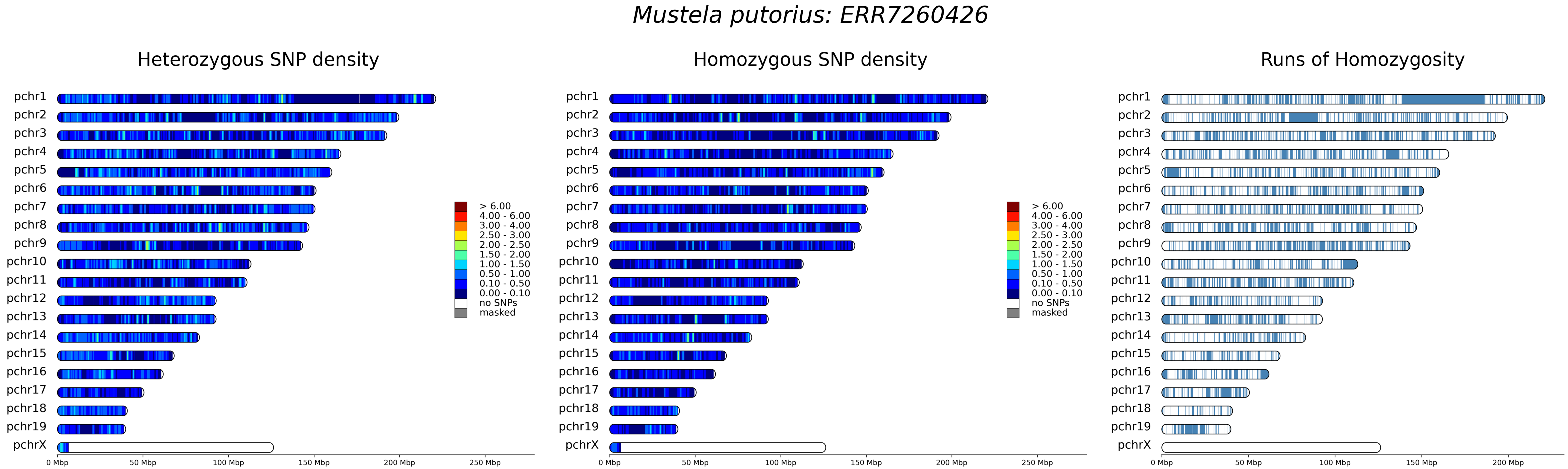
Mustela putorius: ERR7260426
Heterozygous SNP density
Homozygous SNP density
Runs of Homozygosity
pchr1
pchr2
pchr3
pchr4
pchr5
pchr6
pchr7
pchr8
pchr9
pchr10
pchr11
pchr12
pchr13
pchr14
pchr15
pchr16
pchr17
pchr18
pchr19
pchrX
> 6.00
4.00 - 6.00
3.00 - 4.00
2.50 - 3.00
2.00 - 2.50
1.50 - 2.00
1.00 - 1.50
0.50 - 1.00
0.10 - 0.50
0.00 - 0.10
no SNPs
masked
0 Mbp
50 Mbp
100 Mbp
150 Mbp
200 Mbp
250 Mbp

*Mustela putorius: S33*

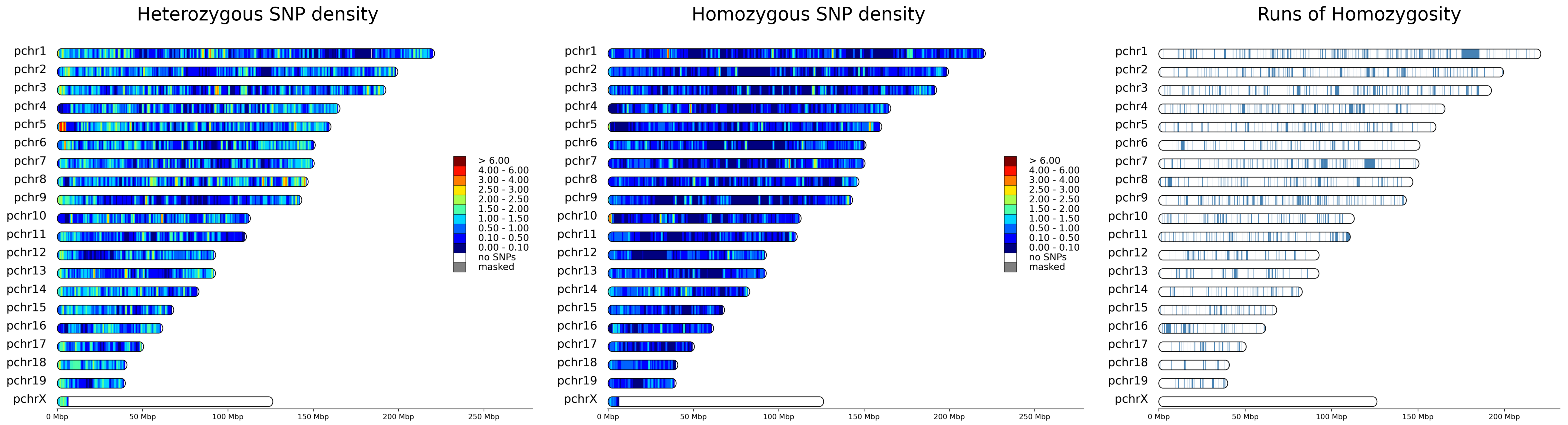

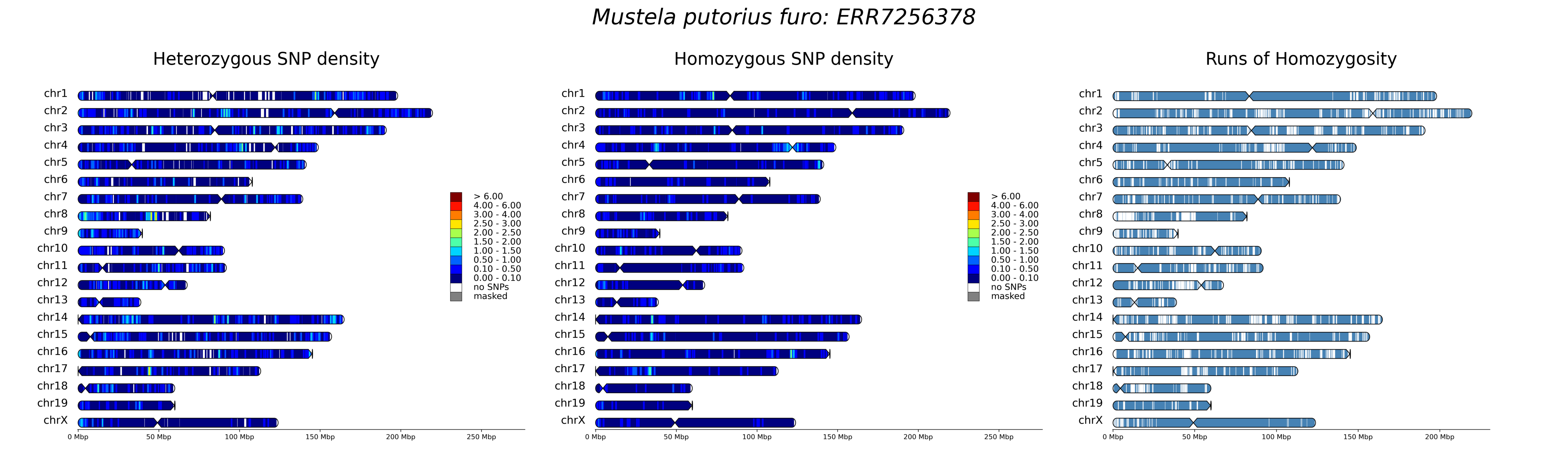

Mustela putorius furo: ERR7256378
Heterozygous SNP density
Homozygous SNP density
Runs of Homozygosity
chr1
chr2
chr3
chr4
chr5
chr6
chr7
chr8
chr9
chr10
chr11
chr12
chr13
chr14
chr15
chr16
chr17
chr18
chr19
chrX
> 6.00
4.00 - 6.00
3.00 - 4.00
2.50 - 3.00
2.00 - 2.50
1.50 - 2.00
1.00 - 1.50
0.50 - 1.00
0.10 - 0.50
0.00 - 0.10
no SNPs
masked
0 Mbp
50 Mbp
100 Mbp
150 Mbp
200 Mbp
250 Mbp

*Mustela putorius furo: ERR7256379*

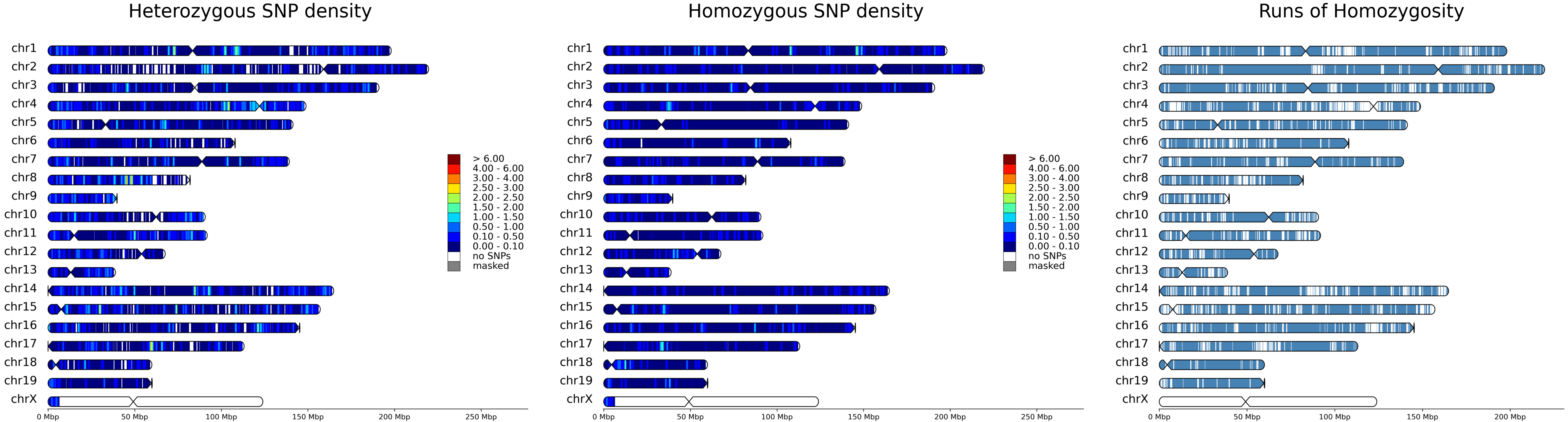

### *Mustela putorius furo: ERR7256380*

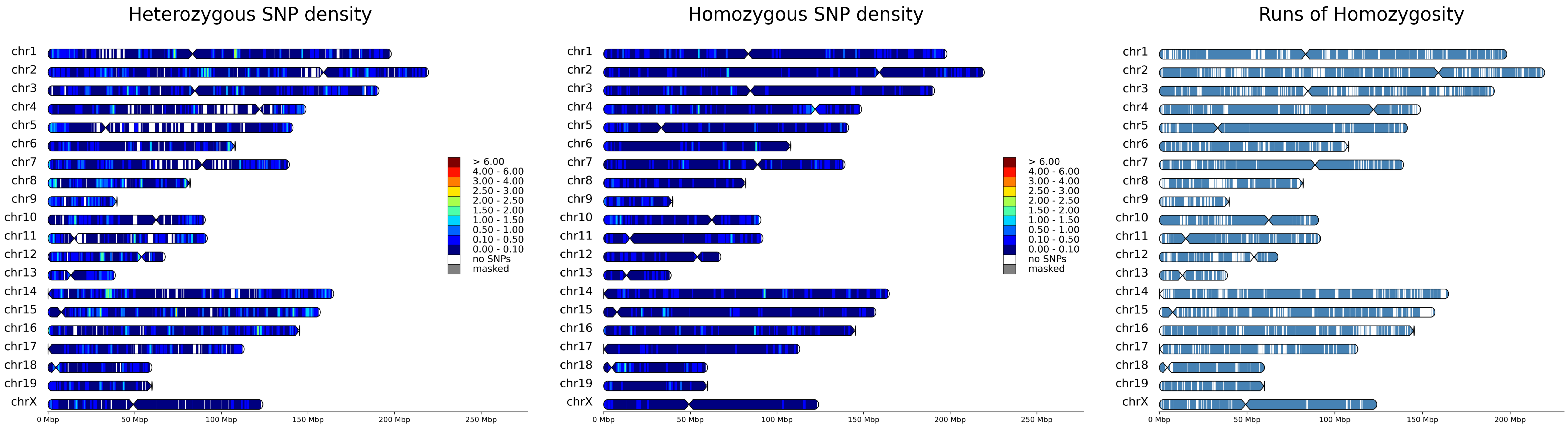

#### *Mustela putorius furo: ERR7256381*

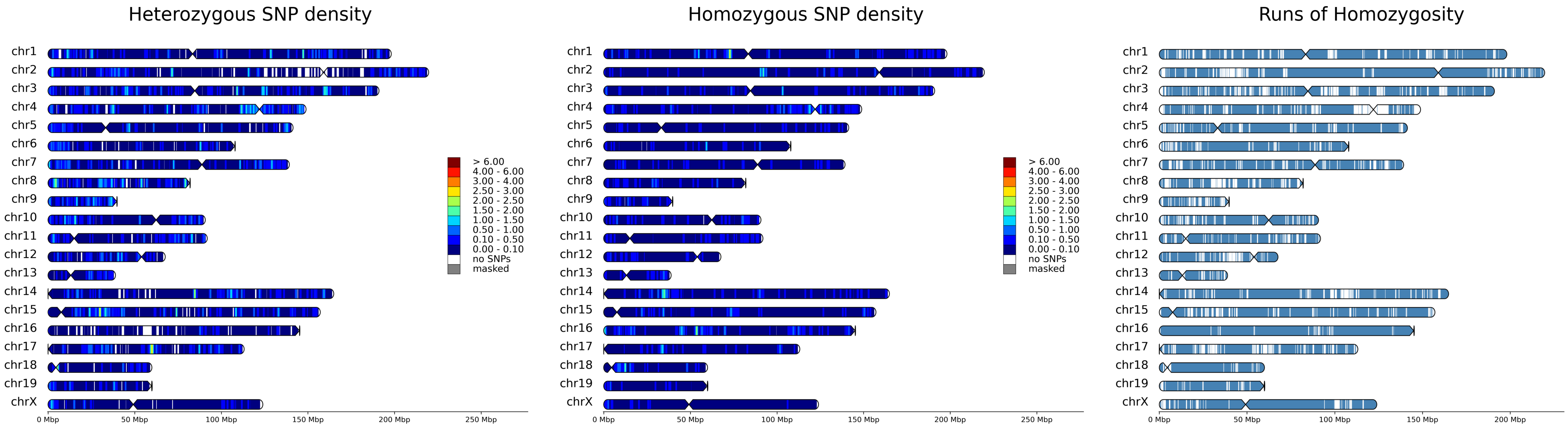

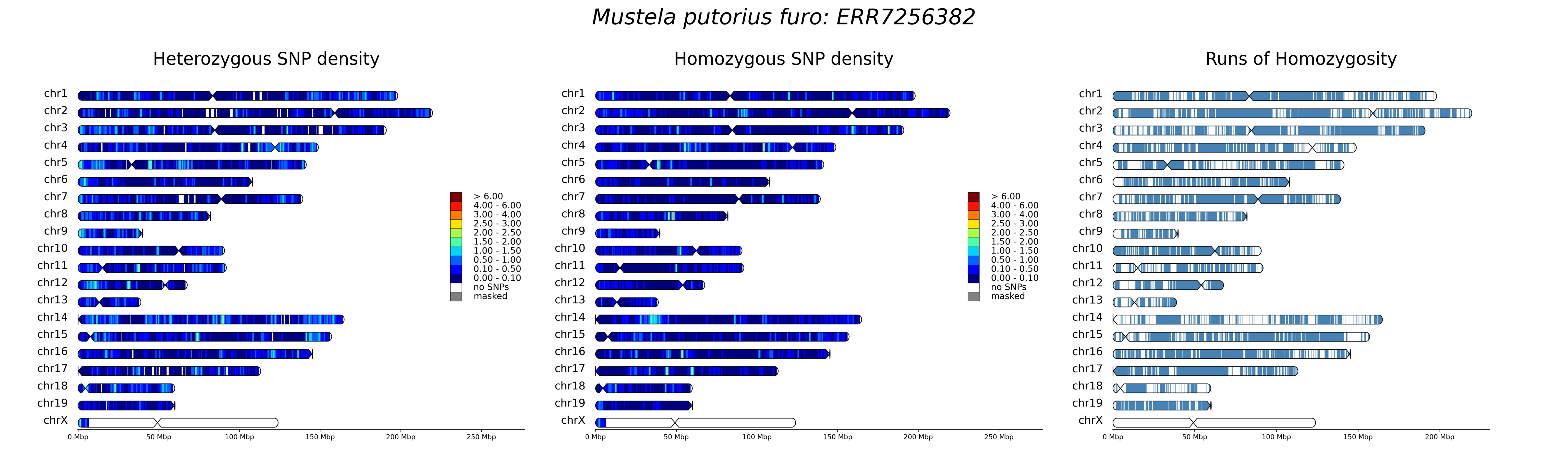

Mustela putorius furo: ERR7256382
Heterozygous SNP density
Homozygous SNP density
Runs of Homozygosity
chr1
chr2
chr3
chr4
chr5
chr6
chr7
chr8
chr9
chr10
chr11
chr12
chr13
chr14
chr15
chr16
chr17
chr18
chr19
chrX
> 6.00
4.00 - 6.00
3.00 - 4.00
2.50 - 3.00
2.00 - 2.50
1.50 - 2.00
1.00 - 1.50
0.50 - 1.00
0.10 - 0.50
0.00 - 0.10
no SNPs
masked
0 Mbp
50 Mbp
100 Mbp
150 Mbp
200 Mbp
250 Mbp

*Mustela putorius furo: ERR7256383*

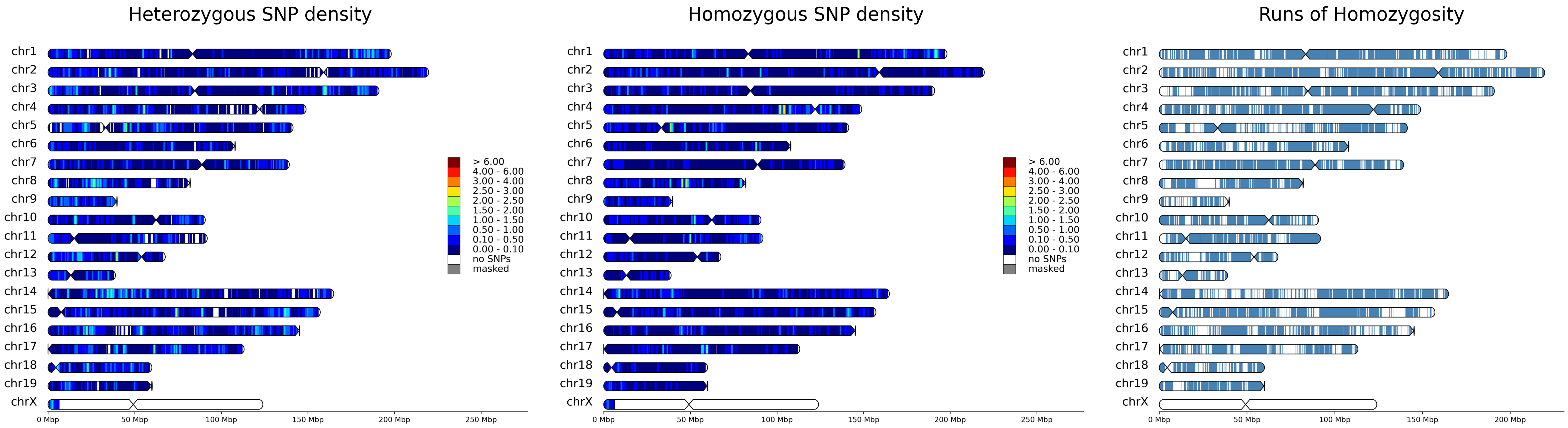

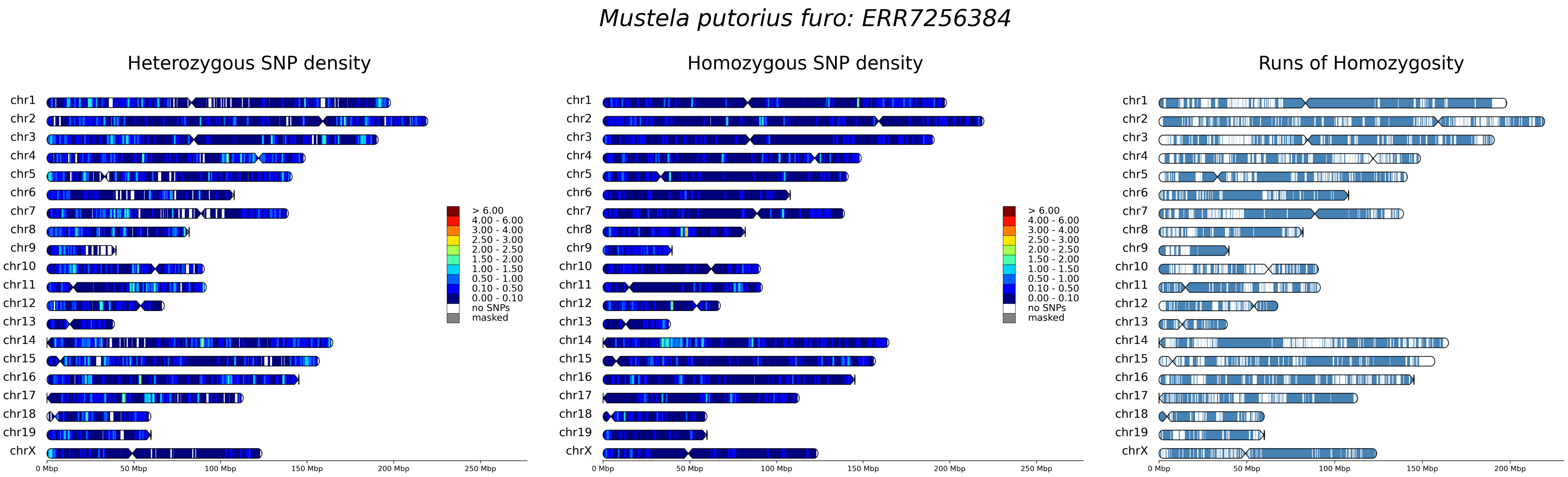
Mustela putorius furo: ERR7256384
Heterozygous SNP density
Homozygous SNP density
Runs of Homozygosity
chr1
chr2
chr3
chr4
chr5
chr6
chr7
chr8
chr9
chr10
chr11
chr12
chr13
chr14
chr15
chr16
chr17
chr18
chr19
chrX
> 6.00
4.00 - 6.00
3.00 - 4.00
2.50 - 3.00
2.00 - 2.50
1.50 - 2.00
1.00 - 1.50
0.50 - 1.00
0.10 - 0.50
0.00 - 0.10
no SNPs
masked
0 Mbp
50 Mbp
100 Mbp
150 Mbp
200 Mbp
250 Mbp

*Mustela putorius furo: ERR7256385*

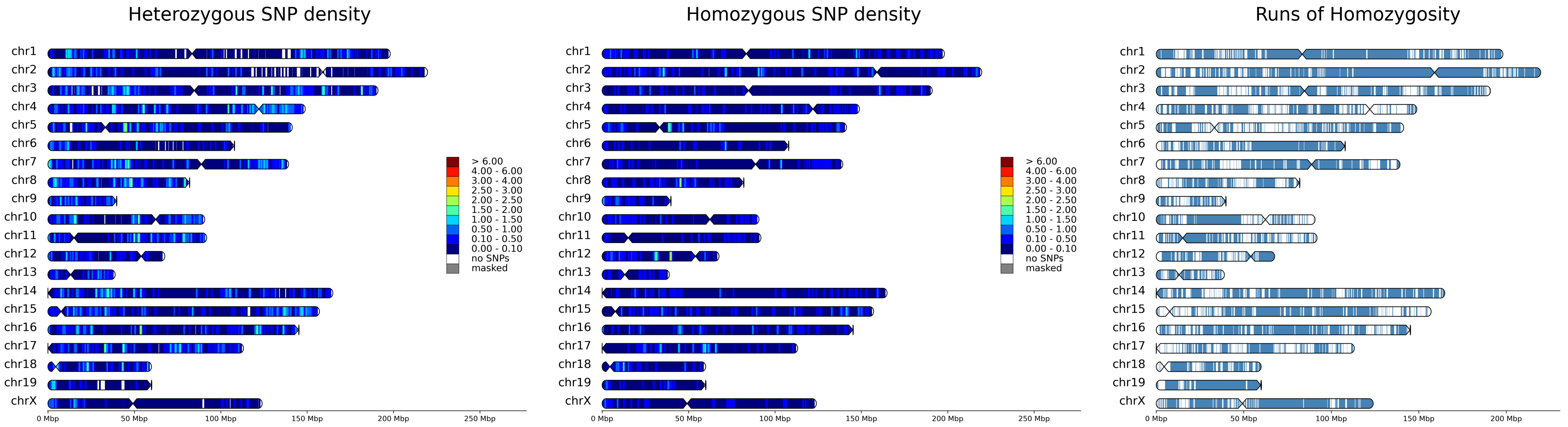

### *Mustela putorius furo: SRR10395504*

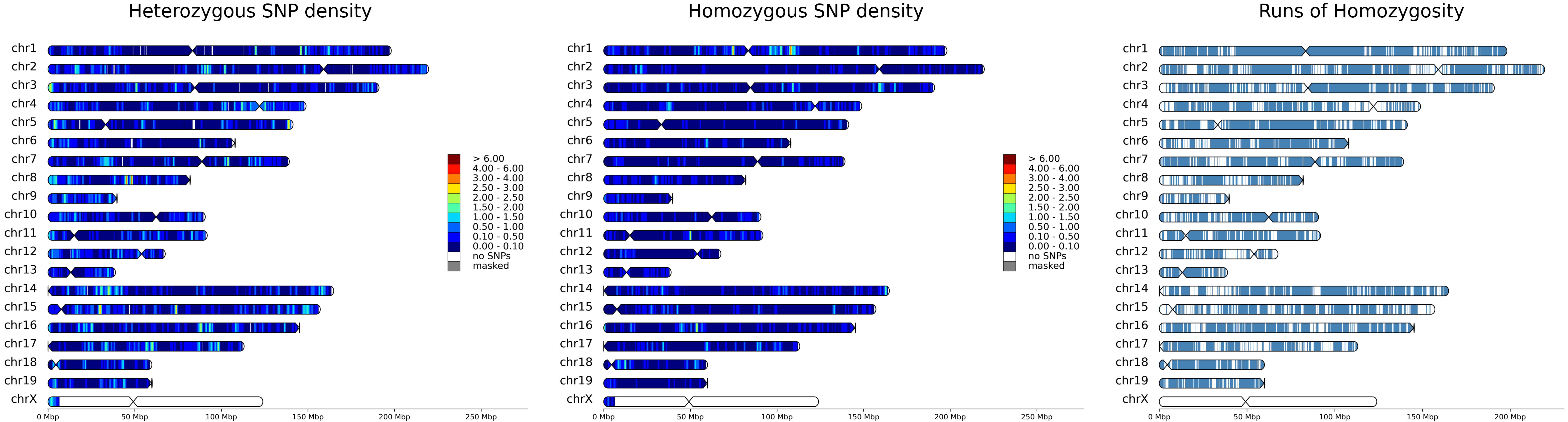

### *Mustela putorius furo: Solexa*

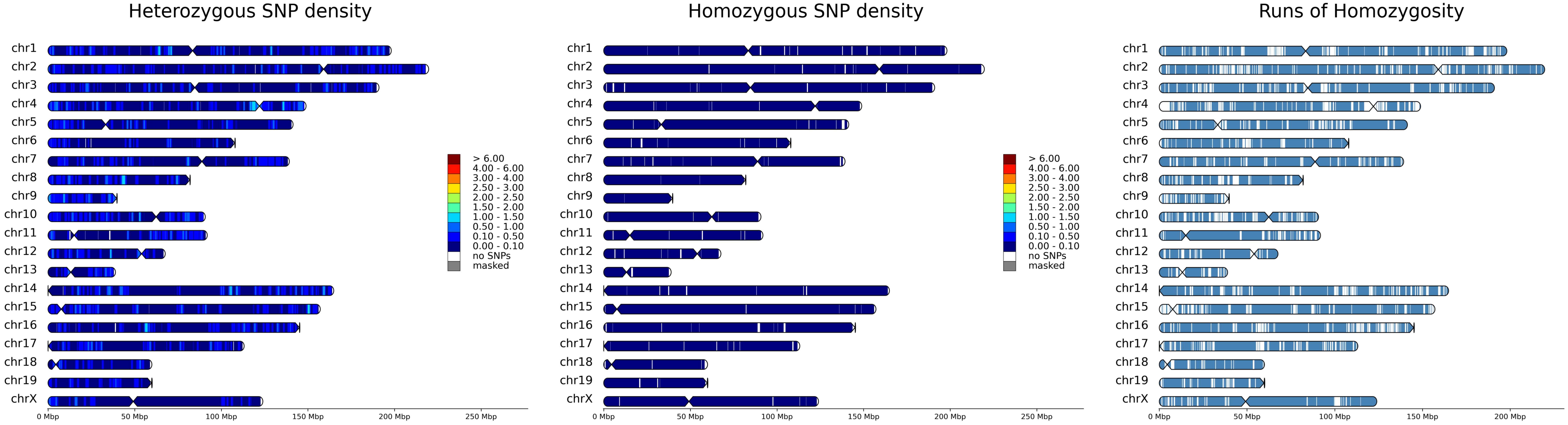

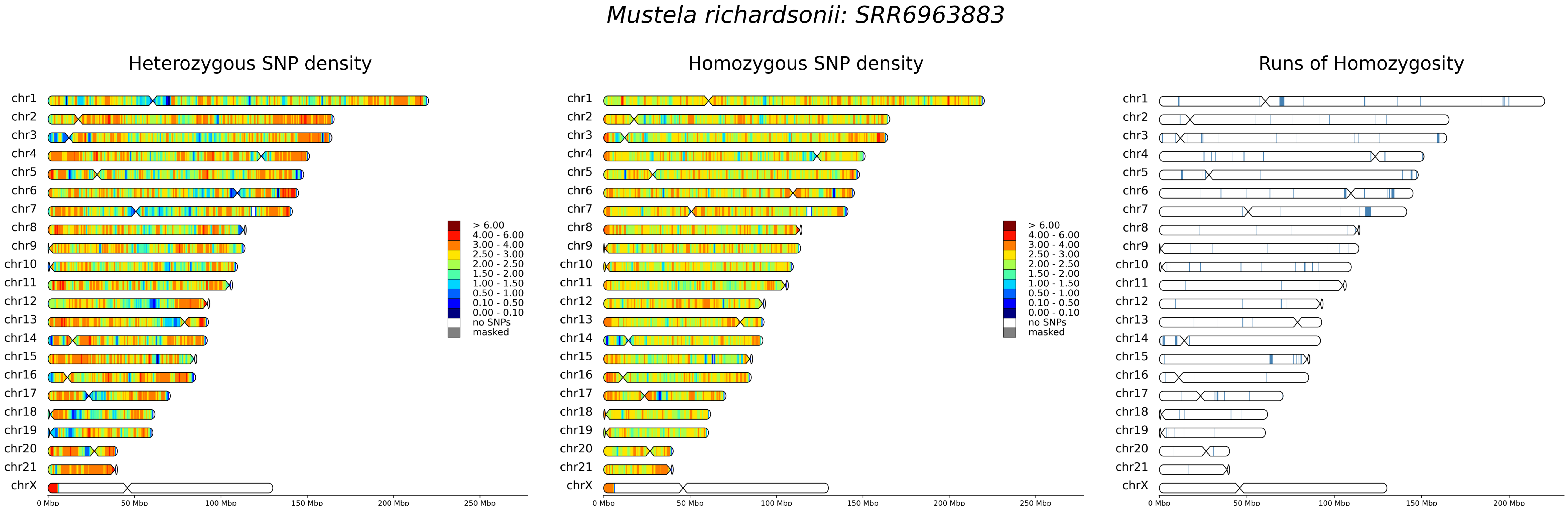
Mustela richardsonii: SRR6963883
Heterozygous SNP density
Homozygous SNP density
Runs of Homozygosity
chr1
chr2
chr3
chr4
chr5
chr6
chr7
chr8
chr9
chr10
chr11
chr12
chr13
chr14
chr15
chr16
chr17
chr18
chr19
chr20
chr21
chrX
> 6.00
4.00 - 6.00
3.00 - 4.00
2.50 - 3.00
2.00 - 2.50
1.50 - 2.00
1.00 - 1.50
0.50 - 1.00
0.10 - 0.50
0.00 - 0.10
no SNPs
masked
0 Mbp
50 Mbp
100 Mbp
150 Mbp
200 Mbp
250 Mbp

*Mustela richardsonii: SRR6963884*

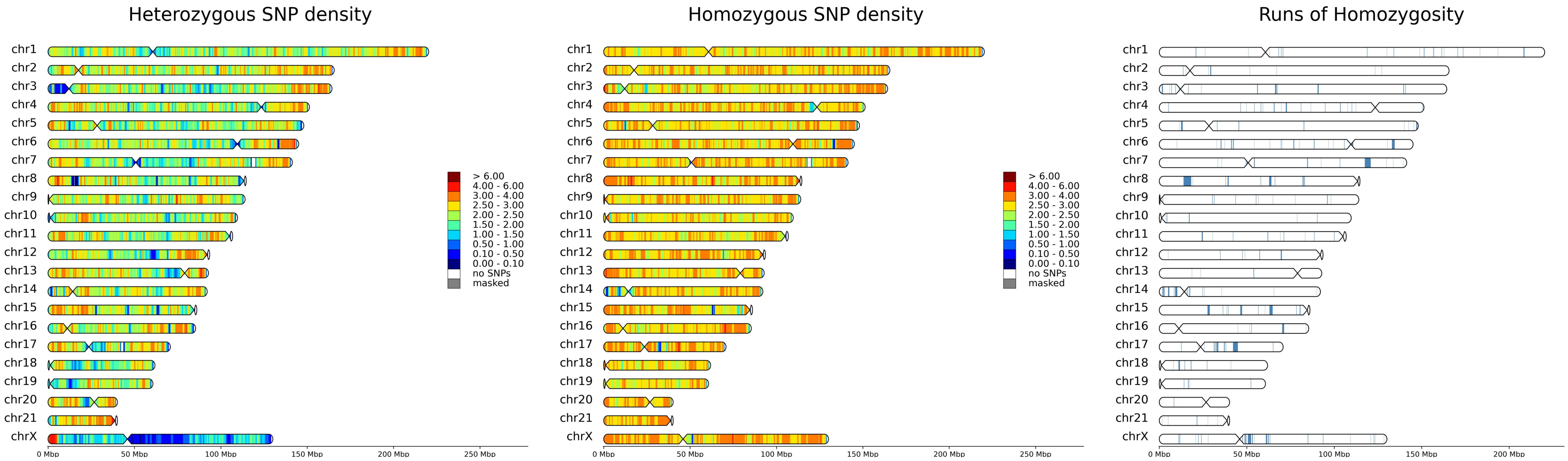

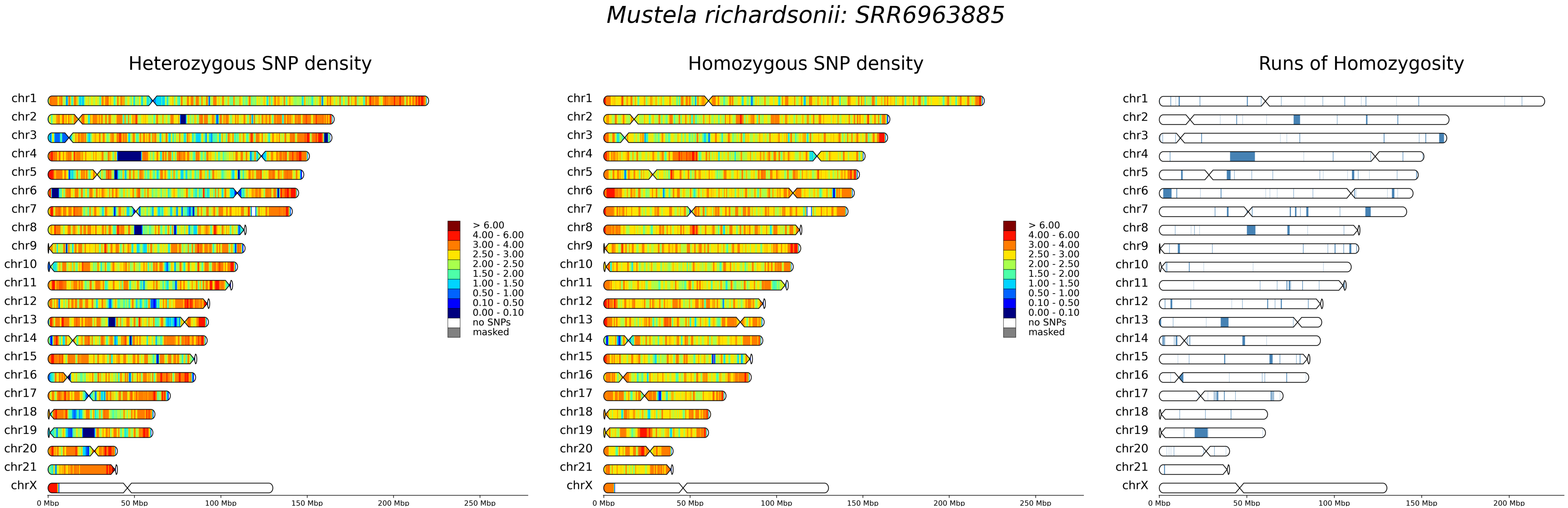
Mustela richardsonii: SRR6963885
Heterozygous SNP density
Homozygous SNP density
Runs of Homozygosity
chr1
chr2
chr3
chr4
chr5
chr6
chr7
chr8
chr9
chr10
chr11
chr12
chr13
chr14
chr15
chr16
chr17
chr18
chr19
chr20
chr21
chrX
> 6.00
4.00 - 6.00
3.00 - 4.00
2.50 - 3.00
2.00 - 2.50
1.50 - 2.00
1.00 - 1.50
0.50 - 1.00
0.10 - 0.50
0.00 - 0.10
no SNPs
masked
0 Mbp
50 Mbp
100 Mbp
150 Mbp
200 Mbp
250 Mbp

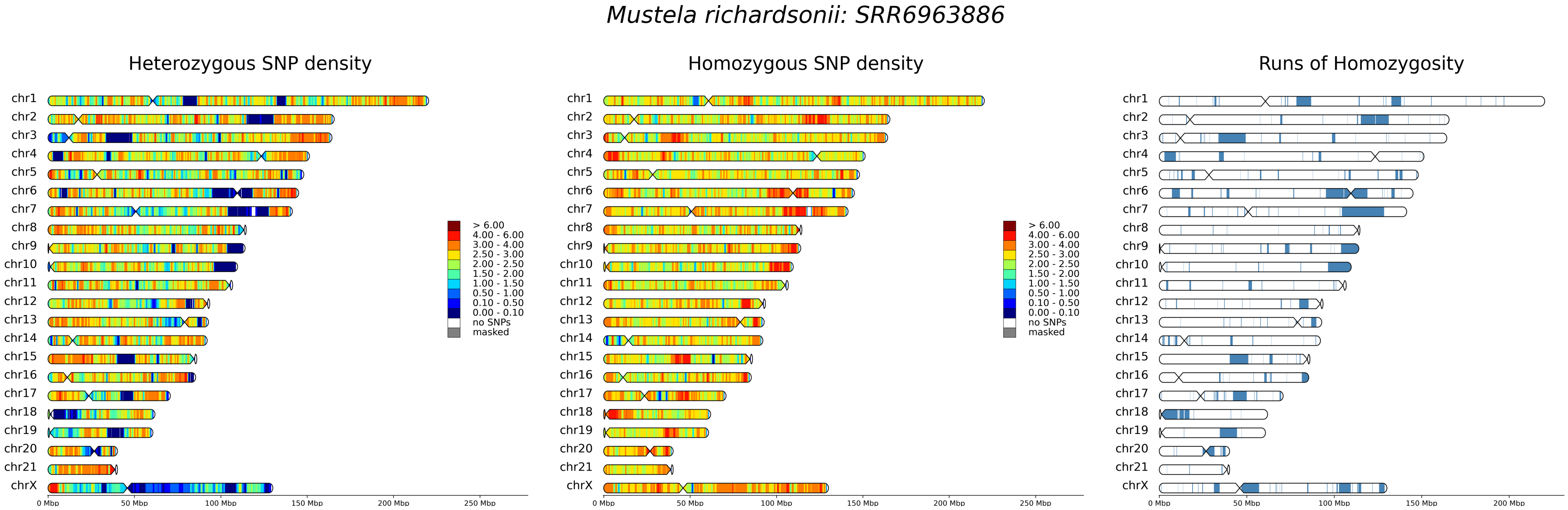

Mustela richardsonii: SRR6963886
Heterozygous SNP density
Homozygous SNP density
Runs of Homozygosity
chr1
chr2
chr3
chr4
chr5
chr6
chr7
chr8
chr9
chr10
chr11
chr12
chr13
chr14
chr15
chr16
chr17
chr18
chr19
chr20
chr21
chrX
> 6.00
4.00 - 6.00
3.00 - 4.00
2.50 - 3.00
2.00 - 2.50
1.50 - 2.00
1.00 - 1.50
0.50 - 1.00
0.10 - 0.50
0.00 - 0.10
no SNPs
masked
0 Mbp
50 Mbp
100 Mbp
150 Mbp
200 Mbp
250 Mbp

#### *Mustela richardsonii: SRR6963887*

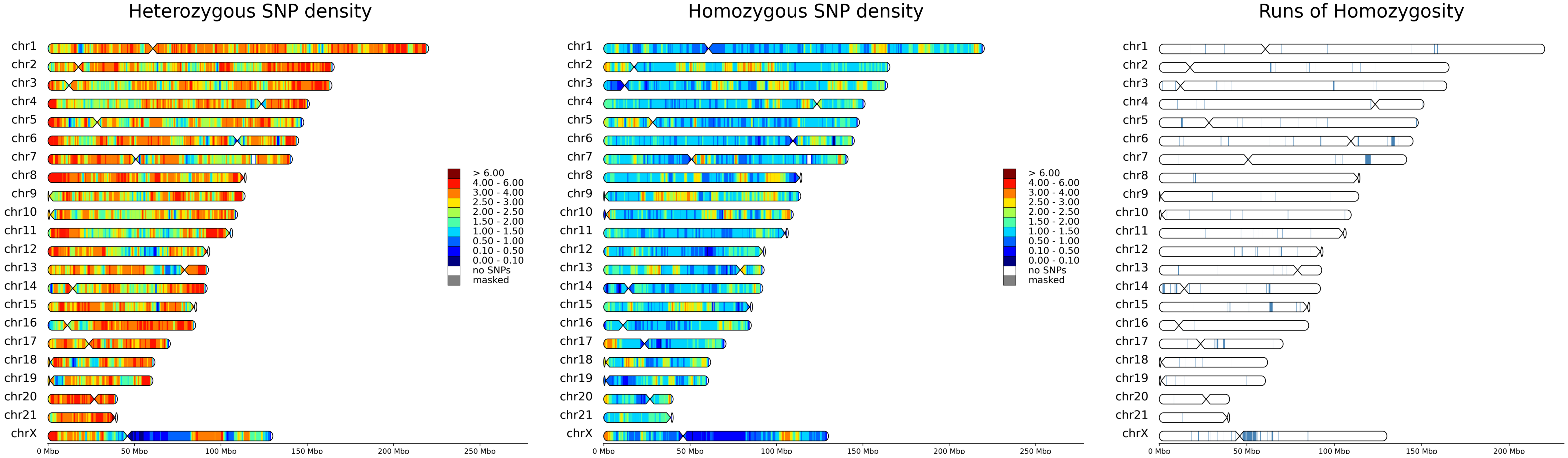

### *Mustela richardsonii: SRR6963888*

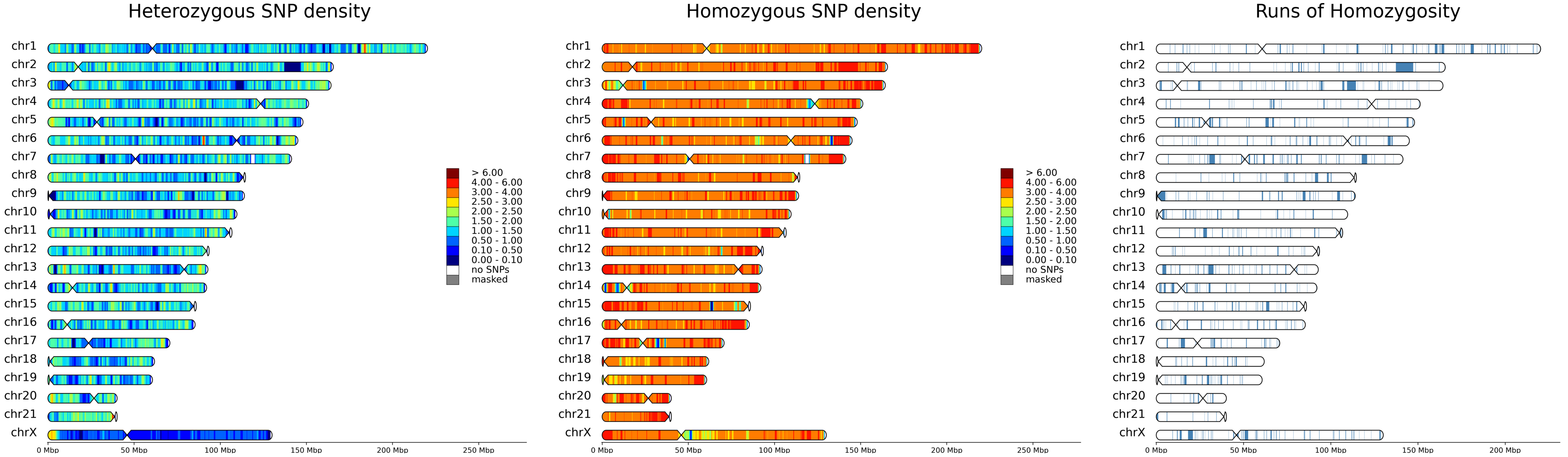

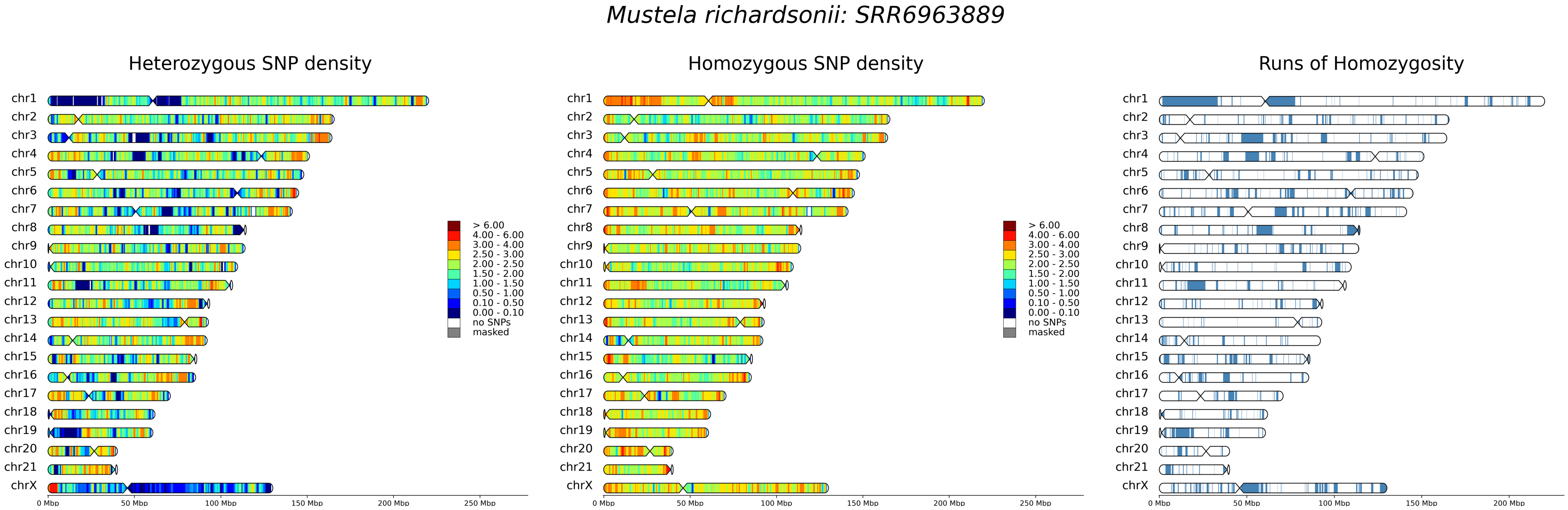

*Mustela richardsonii: SRR6963889*
Heterozygous SNP density
Homozygous SNP density
Runs of Homozygosity
chr1
chr2
chr3
chr4
chr5
chr6
chr7
chr8
chr9
chr10
chr11
chr12
chr13
chr14
chr15
chr16
chr17
chr18
chr19
chr20
chr21
chrX
> 6.00
4.00 - 6.00
3.00 - 4.00
2.50 - 3.00
2.00 - 2.50
1.50 - 2.00
1.00 - 1.50
0.50 - 1.00
0.10 - 0.50
0.00 - 0.10
no SNPs
masked
0 Mbp
50 Mbp
100 Mbp
150 Mbp
200 Mbp
250 Mbp

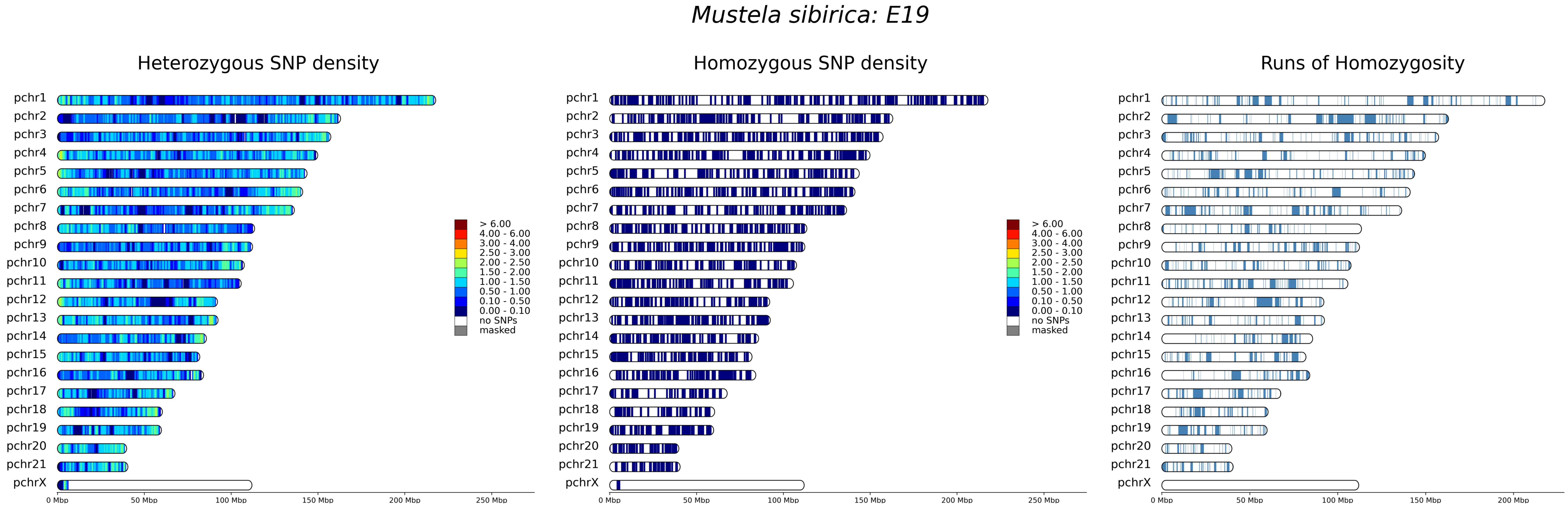
Mustela sibirica: E19
Heterozygous SNP density
Homozygous SNP density
Runs of Homozygosity
pchr1
pchr2
pchr3
pchr4
pchr5
pchr6
pchr7
pchr8
pchr9
pchr10
pchr11
pchr12
pchr13
pchr14
pchr15
pchr16
pchr17
pchr18
pchr19
pchr20
pchr21
pchrX
> 6.00
4.00 - 6.00
3.00 - 4.00
2.50 - 3.00
2.00 - 2.50
1.50 - 2.00
1.00 - 1.50
0.50 - 1.00
0.10 - 0.50
0.00 - 0.10
no SNPs
masked
0 Mbp
50 Mbp
100 Mbp
150 Mbp
200 Mbp
250 Mbp

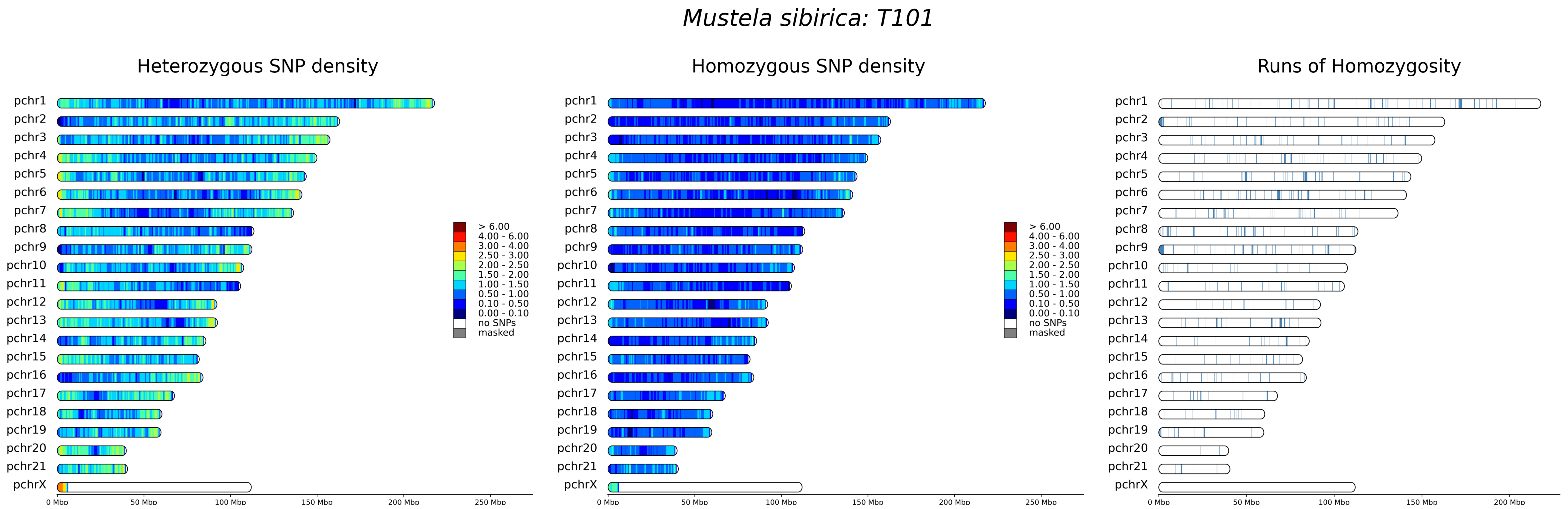

Mustela sibirica: T101
Heterozygous SNP density
Homozygous SNP density
Runs of Homozygosity
pchr1
pchr2
pchr3
pchr4
pchr5
pchr6
pchr7
pchr8
pchr9
pchr10
pchr11
pchr12
pchr13
pchr14
pchr15
pchr16
pchr17
pchr18
pchr19
pchr20
pchr21
pchrX
> 6.00
4.00 - 6.00
3.00 - 4.00
2.50 - 3.00
2.00 - 2.50
1.50 - 2.00
1.00 - 1.50
0.50 - 1.00
0.10 - 0.50
0.00 - 0.10
no SNPs
masked
0 Mbp
50 Mbp
100 Mbp
150 Mbp
200 Mbp
250 Mbp

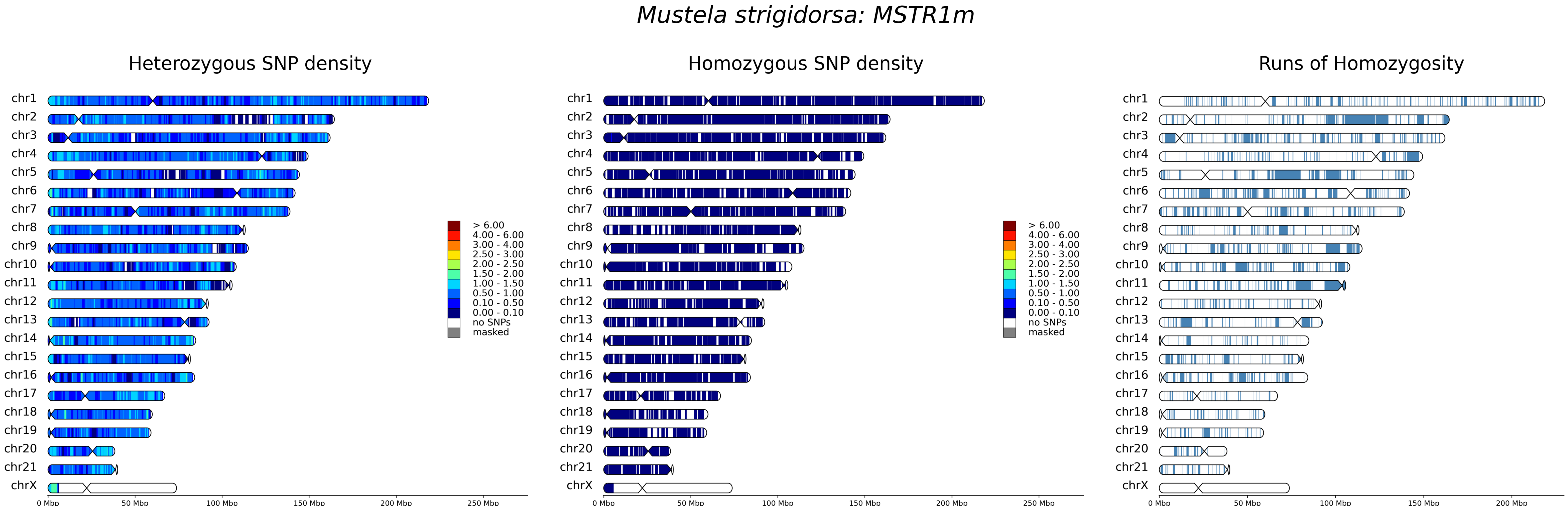
Mustela strigidorsa: MSTR1m
Heterozygous SNP density
Homozygous SNP density
Runs of Homozygosity
chr1
chr2
chr3
chr4
chr5
chr6
chr7
chr8
chr9
chr10
chr11
chr12
chr13
chr14
chr15
chr16
chr17
chr18
chr19
chr20
chr21
chrX
> 6.00
4.00 - 6.00
3.00 - 4.00
2.50 - 3.00
2.00 - 2.50
1.50 - 2.00
1.00 - 1.50
0.50 - 1.00
0.10 - 0.50
0.00 - 0.10
no SNPs
masked
0 Mbp
50 Mbp
100 Mbp
150 Mbp
200 Mbp
250 Mbp

### *Neogale vison: SRR12564099*

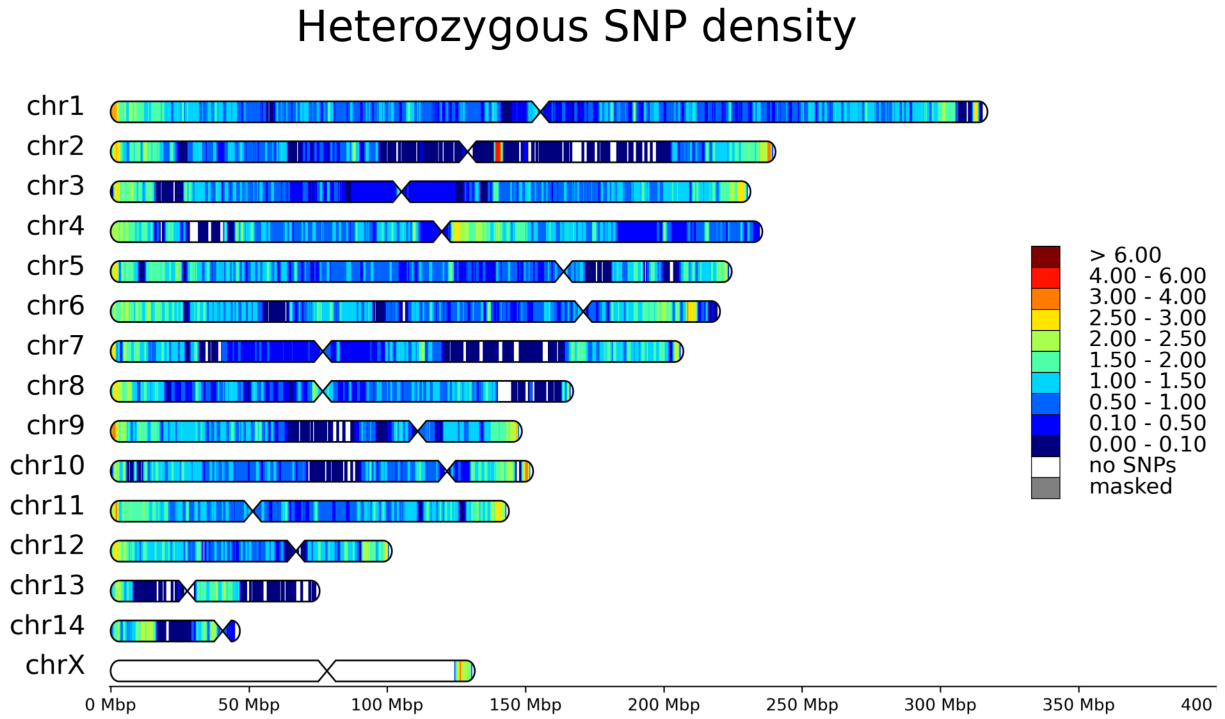


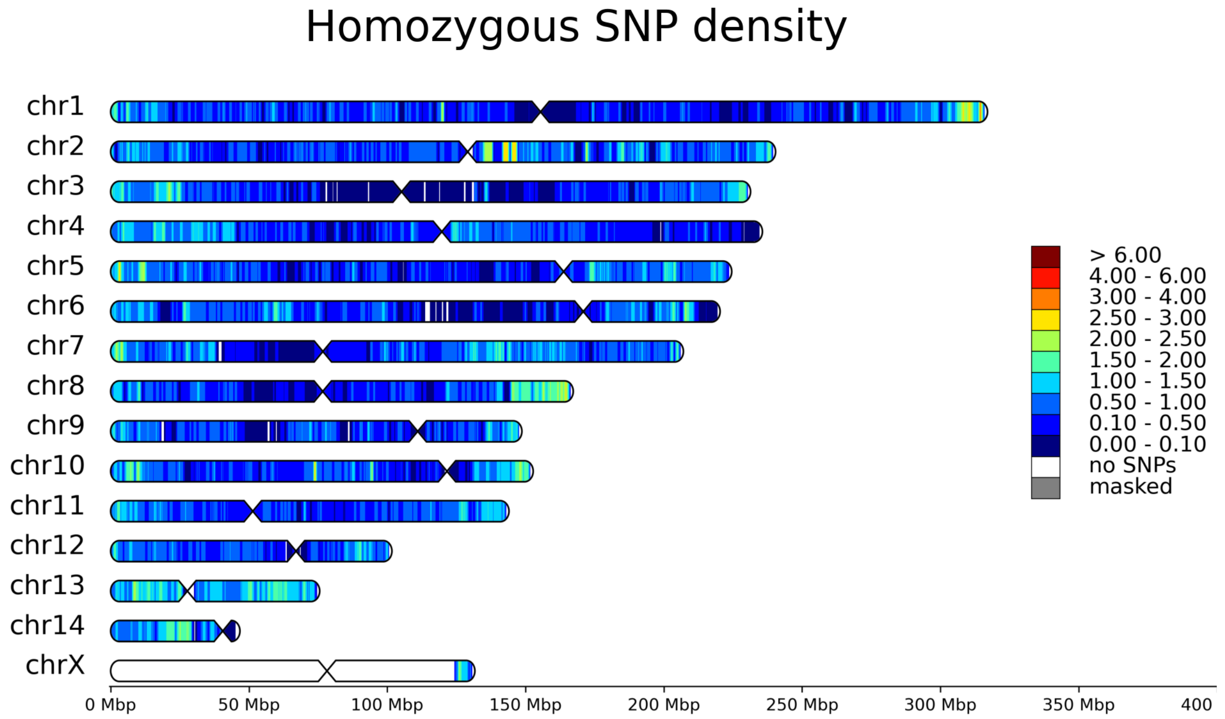


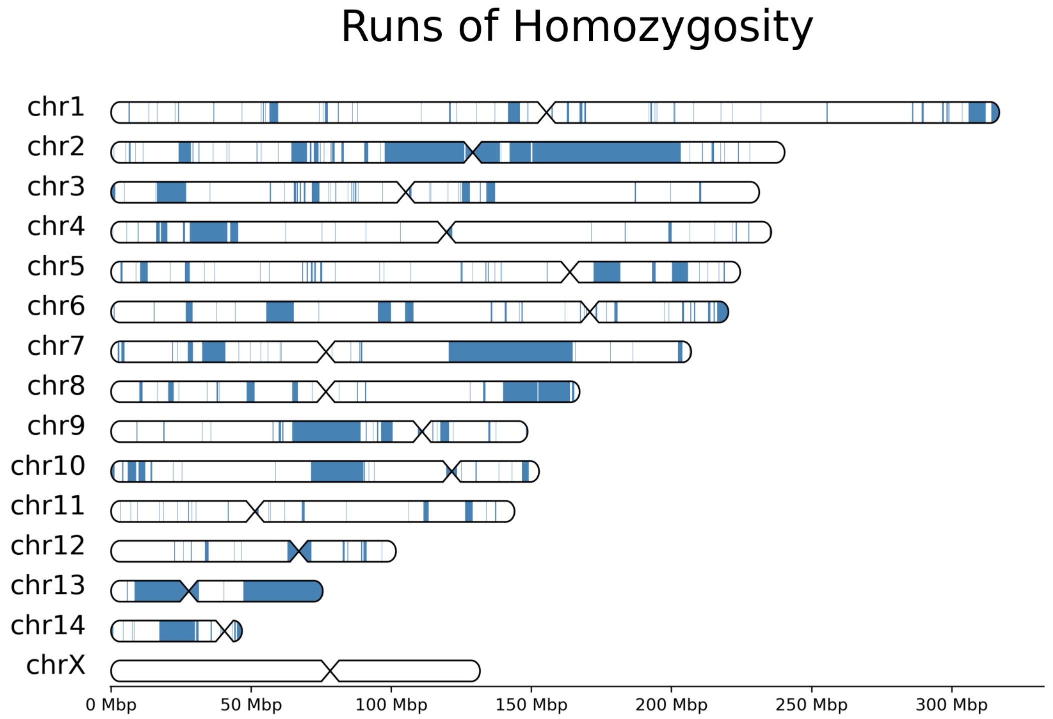

### Neogale vison: SRR16676757

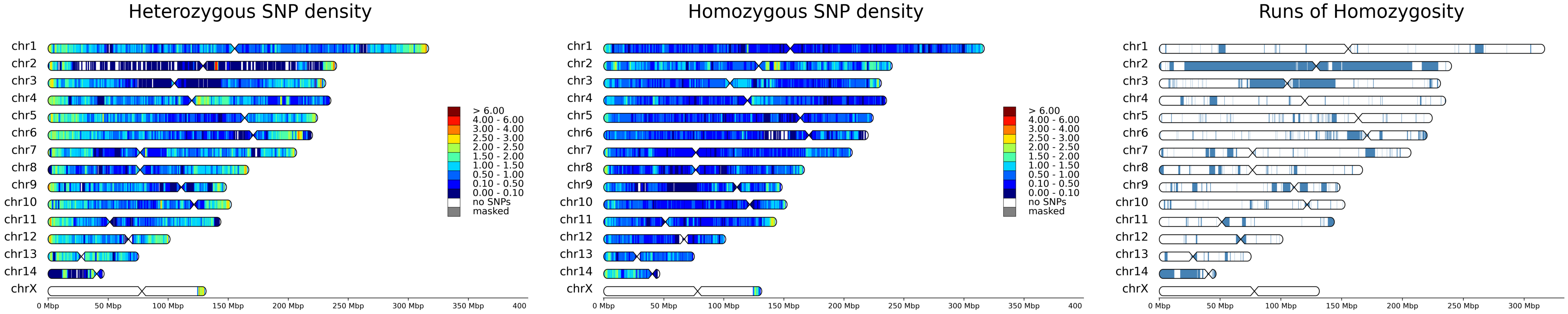

### Neogale vison: SRS11183343

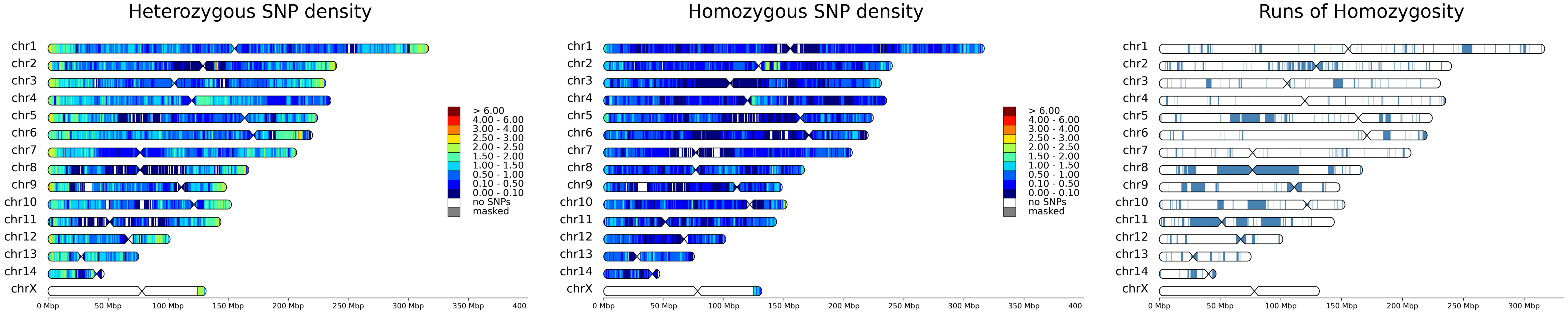
